# Deep whole genome sequencing of multiple proband tissues and parental blood reveals the complex genetic etiology of congenital diaphragmatic hernias

**DOI:** 10.1101/2020.04.03.024398

**Authors:** EL Bogenschutz, ZD Fox, A Farrell, J Wynn, B Moore, L Yu, G Aspelund, G Marth, MY Yandell, Y Shen, WK Chung, G Kardon

## Abstract

The diaphragm is a mammalian muscle critical for respiration and separation of the thoracic and abdominal cavities. Defects in the development of the diaphragm are the cause of congenital diaphragmatic hernia (CDH), a common birth defect. In CDH, weaknesses in the developing diaphragm allow abdominal contents to herniate into the thoracic cavity and impair lung development, leading to a high neonatal mortality. The genetic etiology of CDH is complex. Single nucleotide variants (SNVs), insertion/deletions (indels), and structural/copy number variants in more than 150 genes have been associated with CDH, although few genes are recurrently mutated in multiple patients and recurrently mutated genes can be incompletely penetrant. This suggests that multiple genetic variants in combination, other not yet investigated classes of variants, and/or nongenetic factors contribute to CDH susceptibility. However, to date no studies have comprehensively investigated the contribution of all possible classes of variants throughout the genome to the etiology of CDH. In our study, we used a unique cohort of four patients with isolated CDH with samples from blood, skin, and diaphragm connective tissue and parental blood samples and deep whole genome sequencing to assess germline and somatic *de novo* and inherited variants of various sizes (SNVs, indels, and structural variants) in exons, introns, UTRs, and intergenic regions. In each patient we found a different mutational landscape that included germline *de novo,* and inherited SNVs and indels in multiple genes. We also found in two patients an inherited 343 bp deletion interrupting an annotated enhancer of the CDH associated gene, *GATA4*, and we hypothesize that this common deletion (found in 1-2% of the population) acts as a sensitizing allele for CDH. Overall, our comprehensive reconstruction of the genetic architecture of four CDH individuals demonstrates that the etiology of CDH is heterogeneous and multifactorial.

**AUTHOR SUMMARY:** Deep whole genome sequencing of family trios shows that etiology of congenital diaphragmatic hernias is heterogeneous and multifactorial.

## INTRODUCTION

The diaphragm is a mammalian-specific muscle critical for respiration and separation of the abdominal and thoracic cavities [1]. Defects in diaphragm development lead to congenital diaphragmatic hernias (CDHs), a common structural birth defect [1 in 3000-3500 births; 2, 3-5] in which the barrier function of the diaphragm is compromised. In CDH, a weakness develops in the diaphragm, allowing the abdominal contents to herniate into the thoracic cavity and impede lung development. The resulting lung hypoplasia and pulmonary hypertension are important causes of the neonatal mortality and long-term morbidity associated with CDH [5–7]. The phenotype of CDH is highly variable and the clinical outcomes are diverse, in part due to other associated congenital anomalies [7, 8]. Underlying this phenotypic diversity is a complex genetic etiology [9].

Genetic variants in many chromosomal regions and over 150 genes have been implicated in CDH. Molecular cytogenetic studies of individuals with CDH have identified multiple aneuploidies, chromosomal rearrangements, and copy number variants in different chromosomal regions [9, 10]. Chromosomal abnormalities are found in 3.5-13% of CDH cases and are most frequently associated with complex cases in which hernias appear in conjunction with other comorbidities [9]. In addition, a multitude of individual genes have been identified through analyses of chromosomal regions commonly associated with CDH [e.g. 11], recent exome sequencing studies [12–15], and analyses of mouse mutants [e.g. 16, 17]. Variants in these genes can lead to either isolated or complex CDH. Altogether, the diversity of chromosomal regions and genes associated with CDH demonstrates the heterogeneous genetic origins of CDH.

Most genetic studies of the etiology of CDH have focused on the role of germline *de novo* variants. The preponderance of CDH cases that occur sporadically without a family history of CDH [7] and the low sibling recurrence rate [0.7%; 18] have argued for the importance of this class of genetic variants. Indeed studies of trios of CDH-affected children and their unaffected parents have identified *de novo* pathogenic copy number variants (affecting at least one gene) in 4% of CDH patients [19]. In addition, trio studies employing exome or genome sequencing have discovered candidate *de novo* deleterious gene variants [12–15]. Nevertheless one of these exome sequencing studies [15] estimated that only 15% of sporadic non-isolated CDH cases can be attributed to *de novo* gene-disrupting or deleterious missense variants. Particularly given the small number of cases sequenced to date, most identified genes recur in none or only a few CDH cases [20]. In addition, variants in particular CDH-associated chromosomal regions or genes are often incompletely penetrant for CDH or cause subtle subclinical diaphragm defects [11, 21]. Thus, while *de novo* copy number variants and variants in individual genes undoubtedly are important, the genetic etiology of CDH is more complex and likely polygenic and multifactorial.

Another class of variants that may contribute to CDH etiology are somatic *de novo* variants. A potential role of somatic variants has been suggested by the discordant appearance of CDH in monozygotic twins [18, 22] and the finding of tissue-specific genetic mosaicism in CDH patients [23, 24]. More recently, our functional studies using mouse conditional mutants found that development of localized muscleless regions leads to CDH and suggest that in humans somatic variants in the diaphragm may cause muscleless regions that ultimately herniate [17]. However, to date only one human study has explicitly examined whether somatic variants contribute to CDH [25].

Although less commonly investigated, inherited variants have been linked to CDH. Analyses of families with multiple members affected by CDH revealed that autosomal recessive alleles can cause CDH [26–29]. Other familial CDH cases exhibit an inheritance pattern of autosomal dominance with incomplete penetrance. For instance, two families have been reported with multiple CDH offspring that inherited either a large deletion or frameshift variant in *ZFPM2*, but with unaffected carrier parents [30]. In another case, monoallelic missense variants in *GATA4* were inherited in three generations of one family and associated with a range of diaphragm defects, but only one family member had symptomatic CDH [21]. Thus, these familial cases demonstrate that inherited variants can contribute to CDH etiology, but these genetic variants often exhibit incomplete penetrance and variable expressivity. This suggests that some inherited variants may only lead to CDH in the context of other interacting genetic variants.

While human genetics studies have been essential for identifying candidate CDH chromosomal regions and genes and providing insights into the genetic architecture underlying CDH, experiments with rodents have been critical for determining mechanistically how the diaphragm and CDH develop and explicitly testing whether candidate genes cause CDH. Embryological and genetic lineage experiments [17, 31–33] have shown that the diaphragm develops primarily from two transient embryonic tissues: the somites and the pleuroperitoneal folds (PPFs). The somites are the source of the diaphragm’s muscle, as muscle progenitors migrate from cervical somites into the nascent diaphragm [33]. The PPFs give rise to the diaphragm’s muscle connective tissue and central tendon [17]. Importantly, the PPFs regulate the development of the diaphragm’s muscle and control overall diaphragm morphogenesis, which takes place between embryonic day (E) 9.5 and E16.5 in the mouse [corresponding to E30-60 in humans; 17, 33]. Engineered mutations in the mouse of candidate CDH genes have definitively established that these genes are functionally important in CDH [17]. In addition, conditional mutagenesis experiments indicate that the PPFs are an important cellular source of CDH as inactivation of *Gata4*, *WT1*, or *β-catenin* in the PPFs results in hernias [17, 34, 35], while *Gata4* inactivation in somites does not affect diaphragm development [17]. Furthermore, these experiments established that mutations in CDH genes initiate aberrations in the earliest development of the PPFs [by E12.5 in mouse; 17, 34, 35]. In contrast, mutations in the diaphragm’s muscle lead to diaphragms that are muscle-less or with thin or aberrant muscle, but have never been found to lead to CDH [17, 36–45]. Altogether these data indicate that the PPFs are critical for diaphragm morphogenesis and a cellular source of CDH, while a direct role in CDH for genes expressed in the diaphragm’s muscle is less clear. Given the importance of the PPFs in CDH, prioritization of genes expressed in the early mouse PPFs is likely to be an effective strategy for evaluating new candidate CDH genes derived from human genetic studies.

In this study, we take a novel approach to studying the etiology of CDH. Complementing recent studies using large cohorts of CDH patients that focus on one class of possible variants - *de novo* germline variants [12–15], we comprehensively examine the genome of four CDH patients with multiple tissue samples and their unaffected parents. Using deep whole genome sequencing and a sophisticated bioinformatics toolkit, we determine the contribution of germline and somatic *de novo* and inherited variants to CDH etiology. Our analysis includes variants of different sizes – single nucleotide variants (SNVs), small insertions and deletions (indels), and larger structural variants (SVs) – in all genomic regions (exons, introns, UTR, and intergenic). We prioritize implicated genes not only based on their frequency in the general population and predicted effect on gene function, but on their expression in early PPFs and diaphragm muscle progenitors, using a newly generated mouse RNA-seq dataset. Altogether we reconstruct the diverse genetic architecture underlying isolated CDH in four individuals, revealing the heterogenous and multifactorial genetic etiology of CDH.

## RESULTS

### Strongly supported CDH genes are expressed at high levels in early mouse PPF fibroblasts or diaphragm muscle progenitors

Studies of CDH patients and mouse models have implicated many genes in the etiology of CDH. However, not all CDH-implicated genes are equally well-supported by evidence. To systematically and comprehensively analyze published CDH-implicated genes, we compiled a list of 153 CDH-implicated genes from three large cohort studies [13–15], previous literature reviews [9, 46] and recently published studies [47, 48] (**TableS1**). We included genes that either had mouse functional data or human genetic data found in more than 1 CDH individual, associated with other developmental disorders or structural birth defects that co-occur in patients with complex CDH, and/or implicated [by 13] via their interaction with known CDH genes and expression in the developing diaphragm [as determined by 46]. We ranked each gene based on the nature of the variants, penetrance, and frequency reported in mouse and human data (for details, see Materials and Methods). For mouse data, genes in which variants resulted in herniation of abdominal contents into the thoracic cavity were ranked more highly than those which simply resulted in muscleless regions or entirely muscleless diaphragms. In addition, genes in which variants lead to highly penetrant phenotypes were more highly ranked. For human data, genes in which inherited homozygous or biallelic putative deleterious (as described by original publication) variants or *de novo* deleterious variants were found in more than one CDH patient (either more than one patient in one study or across multiple studies) were ranked more highly than genes with putative deleterious variants of unknown inheritance or found in only one patient. The scores for mouse and human data were added to produce a final score and ranking. Twenty seven genes had a score of 10-19 and were deemed highly likely to contribute to CDH etiology, fifty one genes had a score of 5-9, and seventy five genes had a score of < 5.

Previous mouse studies of the development of CDH demonstrated that the PPFs are a critical cellular source of CDH, and many CDH-implicated genes are expressed and required in early PPF cells [17, 34, 35, 49]. To test whether the CDH genes identified in the literature are expressed in the PPFs or associated diaphragm muscle progenitors, we isolated E12.5 PPF fibroblasts and diaphragm muscle progenitors from wild-type mice and performed RNA sequencing. We found that nearly all 26/27 (96%) highly ranked genes are expressed at levels of at least 10 transcripts per million (TPM) reads (which includes 29% of total transcripts), while 45/51 (88%) of moderately ranked genes and 63/75 (84%) of lowly ranked genes are expressed at this level (**Fig 1A**). These data suggest that genes involved in CDH are expressed at levels of at least 10 TPM in E12.5 PPFs or diaphragm muscle progenitors. In evaluating the significance of newly identified putative CDH-genes, a finding that such genes are expressed at ≥ 10 TPM increases confidence that such genes indeed are important to CDH etiology.

**Figure 1:**
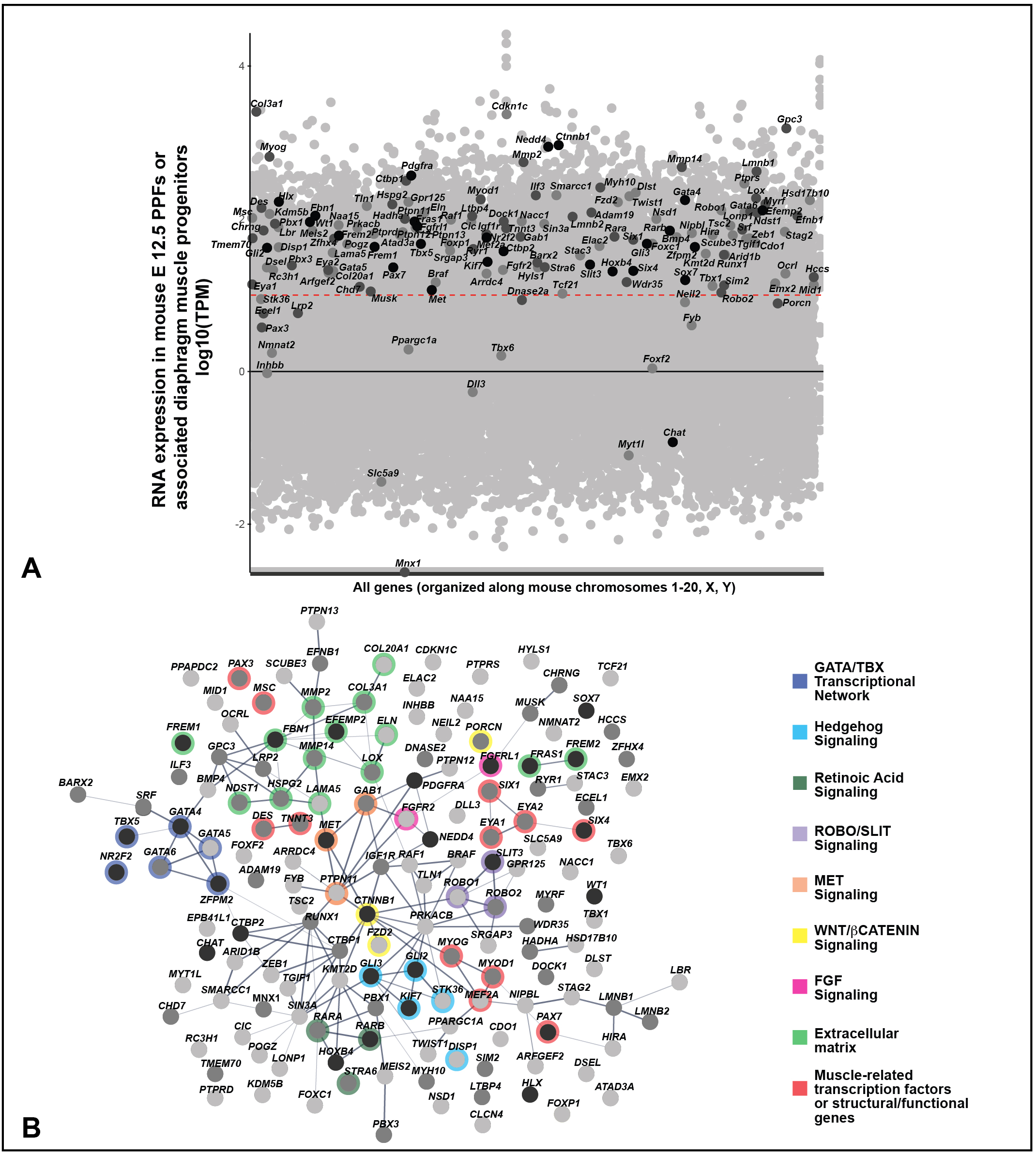
Genes associated with CDH in the literature were collected and ranked based on the amount of human CDH patient and mouse model functional evidence (see Table S1 for rankings). **A)** CDH-associated genes overlaid on all genes expressed in mouse PPFs at E12.5. RNA-seq reads normalized using TPM. Black line denotes TPM = 0 and red dashed line TPM = 10 (Log1). Black dots, CDH genes highly supported by human and mouse data; dark grey dots, genes with moderate data support; light grey dots, CDH genes with modest data support. B) STRING protein network of CDH-associated genes. Within the STRING tool all active interaction sources except text-mining was used, with a minimum required interaction score of 0.4. Edges are based on the strength of data support. Nodes without edges are genes that do not interact with any other listed genes. Thick edges, interaction score > 0.9; thinner edges, 0.9 - 0.7 interaction score; thinnest edges, interaction score < 0.4. Prominent *GATA/TBX* transcriptional network, extracellular matrix, and muscle-related genes are highlighted as well as Hedgehog, Retinoic Acid, ROBO/SLIT, MET, WNT/β-CATENIN, and FGF signaling pathways. In both RNA-seq expression and protein network genes ranked 1-27 (with a total score of ≥ 10) are shown in black, genes ranked 28-88 (with a total score 5-9) are shown in dark gray, and genes ranked 89-153 (with a total score <5) are shown in light gray. Details on rankings are described in Table S1 and Materials and Methods.

Analysis of the highly ranked CDH genes reveals gene families and pathways that likely lead to CDH. To discover protein networks, we inputted the 153 genes into STRING [50] (using all active interaction sources except text mining and requiring a minimum interaction score of 0.4) to generate a protein network (**Fig 1B**). The two highest scoring genes, *GATA4* and *ZFPM2* (*FOG2*), encode a transcription factor and a co-factor that directly interact with each other [51]. In addition, in the heart GATA4 and ZFPM2 interact with the protein encoded by the highly ranked gene, *NR2F2* [52] and GATA4 interacts with the protein encoded by the highly ranked gene, *TBX5* [53, 54]. Thus, not only are variants in *GATA4*, *GATA6*, *ZFPM2*, *NR2F2*, and *TBX5* highly implicated in CDH, but GATA4 (GATA6), ZFPM2, NR2F2, and TBX5 proteins may function together in a complex to regulate diaphragm development. Another class of highly ranked genes are those involved in the extracellular matrix (*EFEMP2, FREM1, FREM2, FRAS1, FBN1, HSPG2, COL3A1, COL20A1, LAMA5*, and *ELN*) as well as genes that modify matrix components (*MMP2*, *MMP14*, *NDST1*, and *LOX*). Also, prominent are genes involved in several critical developmental pathways: SLIT-ROBO signaling (*SLIT3, ROBO1, ROBO2*), Retinoic Acid signaling (*RARB, RARA, STRA6*), SHH signaling (*GLI2, GLI3, KIF7, DISP1, STK36*), and WNT signaling (*CTNNB1, FZD2, PORCN*), MET signaling (*MET*, *GAB1*, *PTPN11*), and FGF signaling (*FGFRL1, FGFR2*).

Present in the list of CDH-implicated genes are those expressed in myogenic cells and critical for myogenesis (*SIX4, SIX1, EYA1, EYA2, MEF2A, MSC, PAX7, PAX3, MYOD1, MYOG, DES, TNNT3*). However, while these genes are important for myogenesis and variants lead to muscle-less regions or muscle-less diaphragms, it is not clear that they lead to diaphragmatic hernias. For instance, variants in *Pax3* in mouse lead to completely muscle-less diaphragms, and while the diaphragms are highly domed, they do not allow abdominal contents to herniate into the thoracic cavity [17]. Thus, while multiple reviews have included these genes as potentially implicated in CDH [e.g. 9, 13, 46], their true role in CDH is less clear.

### Cohort of 4 CDH probands and their parents with multiple proband tissue samples enable unique genetic insights into CDH etiology

To gain greater insight into the genetic etiology of CDH, we analyzed four probands with CDH and their parents with a unique array of tissue samples (**Fig 2**). Our cohort consists of four unrelated individuals (male probands 411 and 967 and female probands 716 and 809) who had an isolated CDH in which the diaphragmatic hernias were encased by a connective tissue sac. Reflecting the increased prevalence of left versus right CDH [7], three probands (411, 809, and 967) had left CDH and one proband (716) had right CDH. Three of the probands (411, 809 and 967) had hernias < 50% of chest wall devoid of diaphragmatic tissue, and one (716) had a large hernia (> 50% of chest wall devoid of diaphragmatic tissue). From each CDH proband, the sac was surgically removed during corrective surgery and saved and skin biopsies and blood draws taken. In addition, blood samples were taken from each parent. Altogether five samples (sac, skin, and blood from CDH proband and blood from 2 parents), from each family were paired-end, whole genome sequenced to an average coverage > 50 reads across each genome. Using Peddy [55], for each pentad of samples we confirmed sample quality, sex, relatedness, and reported ancestry (**Table S2**). The DNA samples from the connective tissue sac uniquely enable us to test the hypothesis, suggested by our previous mouse functional studies [17], that somatic variants in the diaphragm’s connective tissue contribute to the etiology of CDH. In addition, the high depth of whole genome sequencing and availability of new bioinformatic tools enable us to examine a wide spectrum of potential genetic variants including SNVs, indels, and SVs in coding and non-coding regions of the genome.

**Figure 2:**
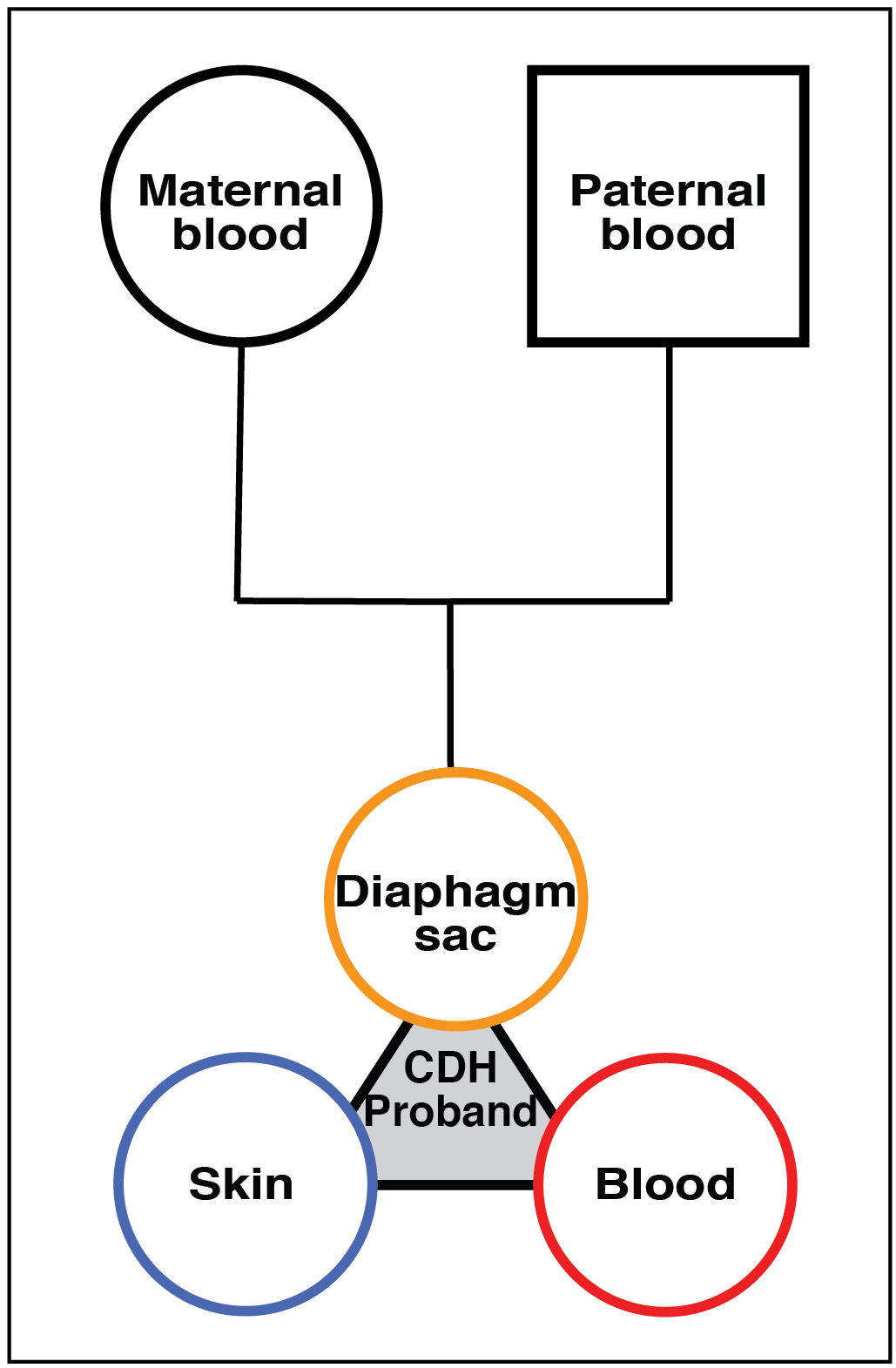
Sample cohorts analyzed in this study. Each cohort consists of DNA samples from diaphragm sac (yellow), skin (blue), and blood (red) of CDH proband and blood from proband’s mother and father.

### Analysis of germline *de novo* variants identifies *THSD7A* as a novel CDH candidate gene

Because most CDH patients have no family history of CDH, previous human genetic studies on the etiology of CDH have concentrated on *de novo* variants [12–15]. Using blood samples from CDH probands and their parents, these studies have inferred that variants present in the proband and not the parents arose in the egg or sperm giving rising to the child and thus designated the discovered variants as germline *de novo* variants. However, blood samples may also harbor somatic variants that arose later in development and such somatic variants could be erroneously designated as a germline variant. Because we have samples from three different tissues of each CDH proband as well as the parents, variants present in the proband diaphragm sac, skin, and blood but not present in the parents are confidently identified as germline (or at least early embryonic) variants. To discover such germline *de novo* variants, we analyzed whole genome sequences from the pentad of samples using the variant calling tool RUFUS (https://github.com/jandrewrfarrell/RUFUS). RUFUS subdivides each genome into a series of k-mers and aligns k-mers from samples of interest to control samples to identify unique SNVs and INDELs in the sample of interest. CDH proband genomes were compared with parental genomes, and variants present in ≥ 20% of reads (with GQ scores >20 and read depths > 15) of proband diaphragm sac, skin, and blood genomes but not in parental genomes were designated as germline *de novo* variants and confirmed by visual inspection in the Integrated Genome Viewer [56].

Germline *de novo* SNVs and INDELs were found in all four CDH probands (shown in intersection of diaphragm sac, skin, and blood of 4 CDH probands in **Fig 3A** and detailed in **Table S3**). CDH probands have 48-116 germline *de novo* variants, falling close to the 70 *de novo* SNVs expected in each newborn in an average population [57, 58]. As expected, the largest number of *de novo* germline variants were in intergenic regions (24-63 variants) and fewer were in UTR or intronic regions (20-50 variants). In the 4 probands, germline *de novo* non-synonymous coding (exon) variants in six genes (*PEX6*, *SCARB1*, *OLFM3*, *ZNF792*, *AR*, and *THSD7A*) were discovered. *De novo* coding variants impacting these genes have not been found in other CDH patients, and these genes are not located within CDH-associated chromosomal regions. All of these coding variants are rare (gnomAD frequency < 0.0001). The *PEX6* variant is a damaging frameshift variant, while the *THSD7A* variant is a damaging nonsense variant (stop gain, p.Trp260Ter). *SCARB1*, *OLFM3, ZNF792*, and *AR* are all missense variants, but only the *OLFM3* and *ZNF792* variants are predicted to be damaging by PROVEAN (which takes into account conservation of homologous sequences [59]). *THSD7A* is the only one of these four genes (*PEX6*, *THSD7A*, *OLFM3*, and *ZNF792*) with damaging variants predicted by ExAC Pli [60] to be haploinsufficient (intolerant of loss of one allele) and also substantially expressed in mouse PPFs or diaphragm muscle progenitors (TPM of 7.0; **Fig 3B**). Thus *THSD7A* is a promising new candidate gene in which variants lead to CDH. THSD7A (Thrombospondin Type I Domain Containing 7A) is a protein containing 10 thrombospondin type I repeats and through its co-localization with α_v_β_3_ integrin and paxillin has been shown to promote endothelial cell migration during development [61–63]. In diaphragm development, *THSD7A* may similarly regulate diaphragm vascularization or it may regulate PPF fibroblast migration, which is essential for diaphragm morphogenesis [17].

**Figure 3:**
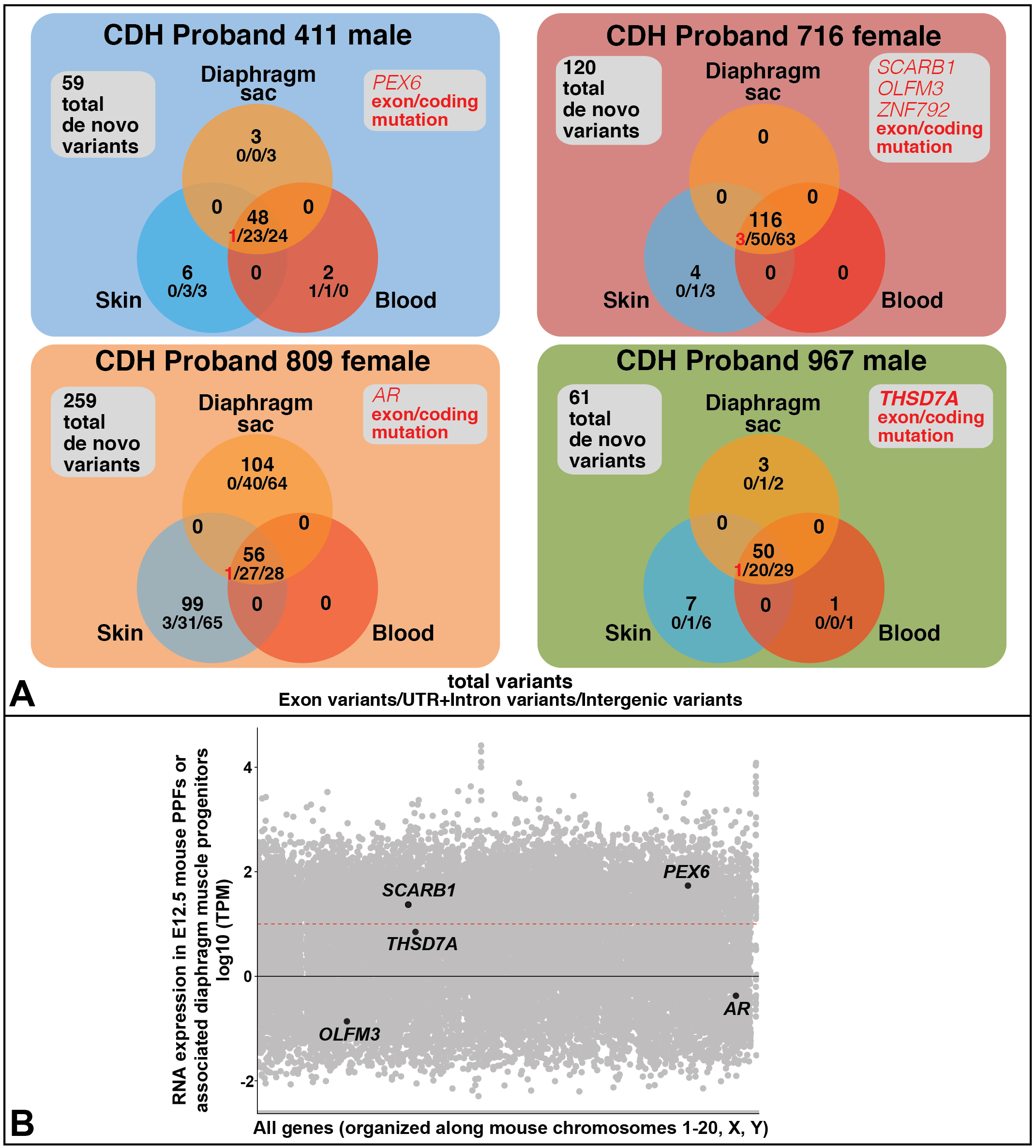
Germline and somatic *de novo* variants, identified via RUFUS, in CDH probands. **A**) *De novo* germline and somatic variants unique to a single tissue and shared between two tissues (coding, gene-associated non-coding, or non-gene associated variants) were found in the 4 CDH probands. 5 genes (*PEX6*, *SCARB1*, *OLFM3*, *ZNF792*, *AR*, and *THSD7A*) contained germline coding variants. Genes with coding variants are highlighted in red. #/#/# = exon (coding) variants/ UTR+intron variants/ intergenic variants. **B)** Five genes with germline *de novo* coding variants overlaid on all genes expressed in mouse PPFs at E12.5 (*ZNF792* has no mouse ortholog). RNA-seq reads normalized using TPM. Black line denotes TPM = 0 and red dashed line TPM = 10. Only *SCARB1*, *PEX6*, and *THSD7A* are expressed in the PPFs at approximately ≥ 10 TPM.

### Somatic *de novo* variants are present in diaphragm, skin and blood, but diaphragm somatic variants are unlikely to contribute to CDH in the probands analyzed

Our previous mouse genetic functional study of CDH [17] suggested the hypothesis that somatic variants in the PPF-derived muscle connective tissue contribute to CDH etiology. Our cohort of CDH proband samples that include the diaphragm sac, which is composed of PPF-derived connective tissue, uniquely allow us to test this hypothesis. In addition, because the samples were collected within a few weeks of birth, we are uniquely positioned to determine the background frequency of somatic variants in the skin and blood prior to exposure of any potential environmental mutagens (e.g. UV light).

To identify somatic *de novo* variants, diaphragm sac, skin, or blood CDH proband genomes were compared via RUFUS with the other 2 genomes derived from the same CDH proband, and variants present in ≥ 20% of reads (with GQ scores >20 and read depths > 15) in only 1 or 2 tissues of each proband genome and not present in parental genomes were designated as somatic *de novo* variants. Variants were only included in which all three tissue samples had at least 15X coverage and Phred quality score [64, 65] above 20. Variants were designated as tissue-specific if ≥ 20% alternate variants reads were present in one or two tissue samples and no alternate reads were present in the other samples.

Somatic *de novo* variants were found in all four CDH probands (**Fig 3**, **Table S3**). Across both individuals and tissues, the alternative allele read depth of somatic variants was significantly lower than germline *de novo* variants (**Fig S1**) [66–68]. This indicates that the somatic *de novo* variants are present in a subset of the sampled cells; in the diaphragm, this reflects either that not all PPF-derived connective cells harbor the variant and/or that the sampled tissue includes several cells types (e.g. connective tissue fibroblasts and endothelial cells) of which only one cell type (e.g. fibroblasts) harbors the variant. An analysis of the mutational spectrum (**Fig S2A**) reveals that the spectrum was generally similar across all probands (although proband 411 and 967 harbor a larger number of deletions) and, as expected, transitions (C/T or A/G changes) are more frequent than transversions (C/G, C/A, A/T, A/C changes). The mutational spectrum also did not vary widely amongst diaphragm, skin, and blood (**Fig S2B**).

Diaphragm sac, skin, and blood were all found to harbor private variants but no variants shared between two tissues (**Fig 3A**, **Table S3**). Intergenic variants were the most common class of somatic variants with variants in UTR and introns as the next most common. Somatic coding variant were only found in the blood of Proband 411 (missense variant in *MIR4717*) and in the skin of Proband 809 (missense variants in *DHX57* and *TNFAIP8L3* and a synonymous variant in *CXorf57*). In all four probands, the diaphragm contained somatic variants, but none of these were in coding regions and variants in UTR and intron regions did not overlap any annotated enhancers. Thus, while somatic variants were found in the diaphragm, none of the variants are likely to contribute to the etiology of CDH in these children.

One striking feature of our analysis was the extremely high number of somatic variants in the diaphragm sac and skin, but not blood, of proband 809. This high number of variants was not an artifact of technical issues, as the samples passed all quality control filters of the standardized Utah Genome Project pipeline (see Materials and Methods). Furthermore, not only does this child harbor more somatic variants (**Fig 3A**), but the alternate allele frequency is significantly higher than that found in the somatic variants in the other probands (**Fig S1**). This child also harbors a missense germline *de novo* in the androgen receptor gene *AR*. *AR* has been well characterized as a tumor suppressor gene, specifically in prostate cancer [69], and shown to be critical in DNA repair through activation of transcriptional targets [70, 71]. However, arguing against a causative role of *AR* is that the particular *AR* missense variant in proband 809 is not predicted by Provean to be damaging.

Our analysis of somatic variants also provides insights into how representative variants in the blood are of germline variants. Blood is the most commonly sampled human tissue and variants in the blood are typically designated as “germline” variants. However, our analysis shows that blood harbors somatic variants that would typically be erroneously tallied as germline variants. In the four children analyzed, an average of 1.5% (range of 1-4%) of variants in the blood are unique to blood. Thus our analysis suggests that in general 1.5% of variants found in the blood are not germline variants, but instead somatic variants.

### Inherited variants in genes regulating muscle structure and function potentially contribute to the etiology of CDH in one proband

Damaging variants inherited from parents may also be a genetic source of CDH. As the parents of the four probands investigated do not have CDH, it is unlikely that an inherited variant of one allele would directly cause CDH, while homozygous or biallelic variants, in which each parent contributes a gene damaging allele, are more likely to contribute to CDH. Given this, recessive homozygous and biallelic inherited variants were identified and prioritized using the GATK variant calling pipeline and the variant prioritizing tool VAAST3 [72, 73] (**Table S4**). VAAST identifies and ranks genes based on whether they are predicted to be damaging based on protein impact and rare compared against a mixed control population from 1000 Genomes phase 3 [74]. Given that the significance level VAAST assigns to a gene is limited by the size of the background population, genes were further prioritized as to their likelihood to be involved in CDH etiology. Pseudogenes [75], highly mutable genes [76, 77], and genes with multiple alleles in the variant region reported in gnomAD [78] were removed and genes expressed at higher than 10 TPM in E12.5 mouse PPFs or diaphragm muscle progenitors prioritized (**Fig 4A, B** and **Table S4**).

**Figure 4:**
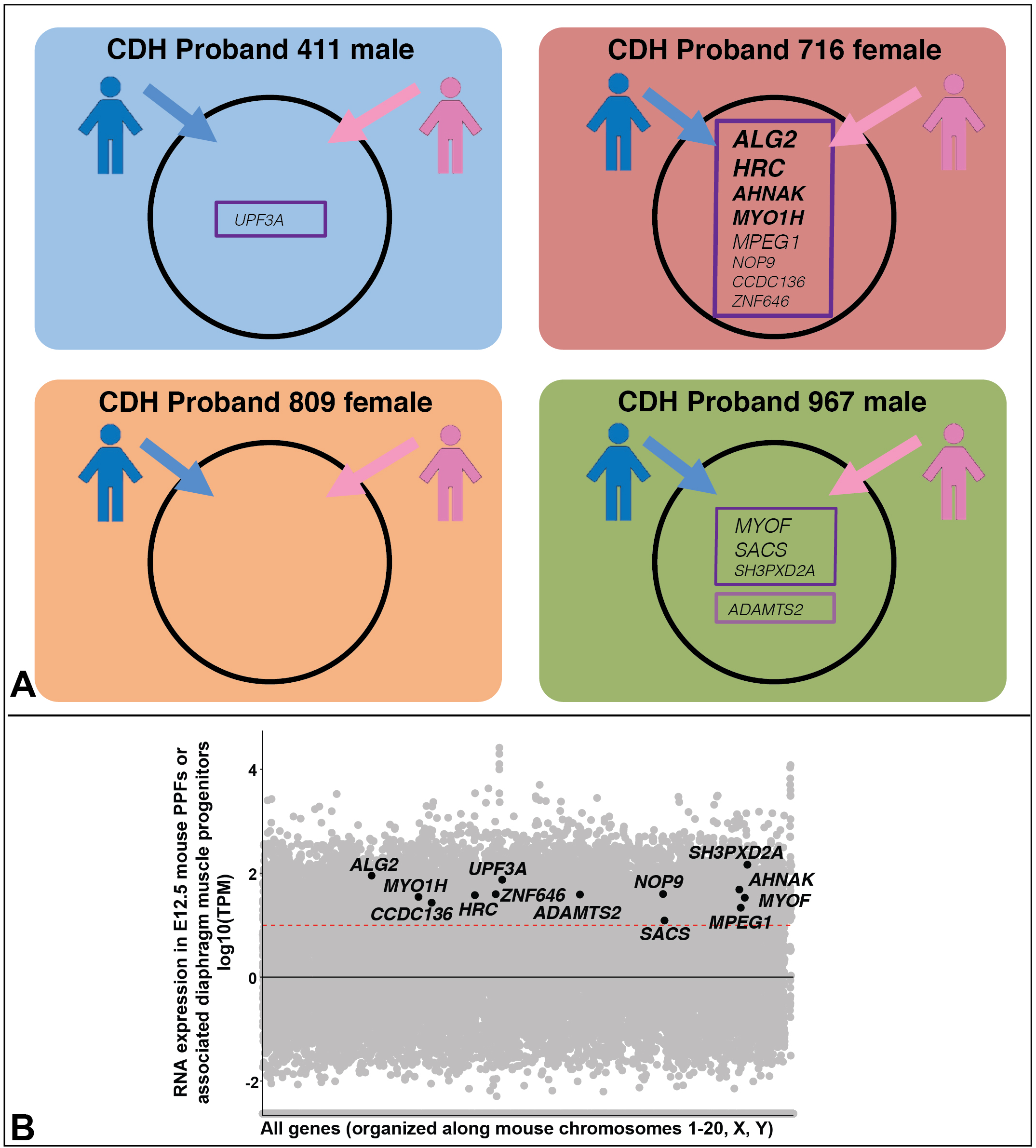
Inherited variants potentially contributing to etiology of CDH probands. **A)** Compound heterozygous (boxed in dark purple) or homozygous (boxed in light purple) inherited variants highly ranked as damaging and rare by VAAST, with pseudogenes, highly mutable genes, and genes expressed at low levels in PPFs filtered out. **B)** Candidate genes with inherited variants overlaid on all genes expressed in mouse PPFs at E12.5. RNA-seq reads normalized using TPM. Black line denotes TPM = 0 and red dashed line TPM = 10.

Using these criteria, we identified 13 genes with biallelic or homozygous variants in the three CDH probands (**Fig 4A** and **Table S4** with VAAST, CADD, PROVEAN impact, and ExAC Pli scores). None of these genes has been identified in previous CDH studies. Of these 13 genes, 3 genes (*MPEG1*, *MYOF*, *SACS*) harbor 1 deleterious frameshift allele and 1 missense allele predicted by Provean to be neutral; 5 genes (*UPF3A*, *CCDC136*, *NOP9*, *ZNF646*, *SH3PXD2A*) harbor 1 missense allele predicted to be deleterious and 1 missense allele predicted to be neutral by Provean; and 1 gene (*ADAMTS2*) is homozygous for an inherited in-frame insertion, predicted by Provean to be neutral, but predicted by ExAC to be intolerant of loss-of-function (Pli score of 0.99). Because each of these genes has at least one predicted functional allele, these genes are unlikely to directly cause CDH, but may act as sensitizing alleles that act in combination with other genetic variants produce CDH.

Proband 716, with a large right hernia, harbored four genes – *ALG2*, *HRC*, *AHNAK*, and *MYO1H* – in which both inherited maternal and paternal alleles were frameshift null or missense predicted damaging alleles and therefore more likely to contribute to CDH etiology. All four genes contain variants that are rare compared to the background population (1000 Genomes Phase 3), based on the probabilistic framework underlying VAAST, and all variants are rare (AF < 0.01) in gnomAD (**Table S4**). Interestingly, three of these genes are involved in skeletal muscle function. *ALG2* encodes an α1,3-mannosyltransferase that catalyzes early steps of asparagine-linked glycosylation and is expressed at neuromuscular junctions [79]. Human *ALG2* variants have been found to affect the function of the neuromuscular junction and mitochondria organization in myofibers, leading to congenital myasthenic syndrome (fatigable muscle weakness) and mitochondrial myopathy [79–81]. *HRC* encodes a histidine-rich calcium-binding protein that is expressed in the sarcoplasmic reticulum of skeletal, cardiac, and smooth muscle [82] and in the heart has been shown to regulate calcium cycling [83, 84]. AHNAK encodes a large nucleoprotein that acts as a structural scaffold in multi-protein complexes [85]. In particular, AHNAK interacts with dysferlin, which is a transmembrane protein critical for skeletal muscle membrane repair and loss of dysferlin causes several types of muscular dystrophy [86, 87]. AHNAK is proposed to play a role in dysferlin-mediated membrane repair [86, 87]. The finding of deleterious biallelic variants in these three genes suggests that aberrations in the diaphragm muscle’s neuromuscular junctions, mitochondria, calcium handling, or membrane integrity contributes to the development of CDH in this child. In addition, proband 716 has one predicted null and one missense deleterious allele in *MYO1H*. MYO1H is a motor protein involved in intracellular transport and vesicle trafficking, expressed in retrotrapezoid neurons critical for sensing C0_2_ and regulating respiration, and variants in *MYO1H* cause a recessive form of central hypoventilation [88]. While unlikely to directly contribute to CDH, the *MYO1H* variants’ potentially deleterious effect on neuronal regulation of respiration would have a detrimental impact on a CDH child.

### An inherited deletion in intron 2 of *Gata4* in two probands is a candidate common sensitizing allele for CDH

Another important potential source of genetic variants underlying CDH are structural variants [SVs; 47] and include ≥ 50 bp insertions, deletions, inversions, and translocations. To identify structural variants that could contribute to the etiology of the four CDH probands, we used the Lumpy smooth pipeline [89], which uses three copy number variant callers: cn.MOPS [90], CNVkit [91] and CNVnator [92]. We then determined whether identified SVs were *de novo* or inherited in the CDH probands using the tool GQT [93] and confirmed SVs visually using IGV [56]. Though *de novo* SVs were discovered in multiple probands (**Table S5**), all were common (allele frequency > 0.1) in a large SV dataset of 14,623 ancestrally diverse individuals [94] or located in an intergenic region. Thus, no discovered *de novo* SVs are likely to contribute to CDH etiology.

To discover inherited SVs potentially contributing to CDH, we analyzed variants within chromosomal regions highly associated with CDH [10]. Multiple candidate inherited SVs were identified (with allele frequencies < 0.1 and not found in intergenic or repetitive regions, **Table S5**), but in no case were the probands homozygous for the SV or biallelic with any of the identified SNVs. However, one SV, a 343 base pair deletion within the second intron of the highly ranked CDH-associated transcription factor, *GATA4* [Table S1; 16, 17, 21], was discovered in two probands (**Fig 5**) and subsequently confirmed by PCR. In proband 411 the deletion is paternally inherited (**Fig 5A**), while in proband 967 the deletion is maternally inherited (**Fig 5D**).

**Figure 5:**
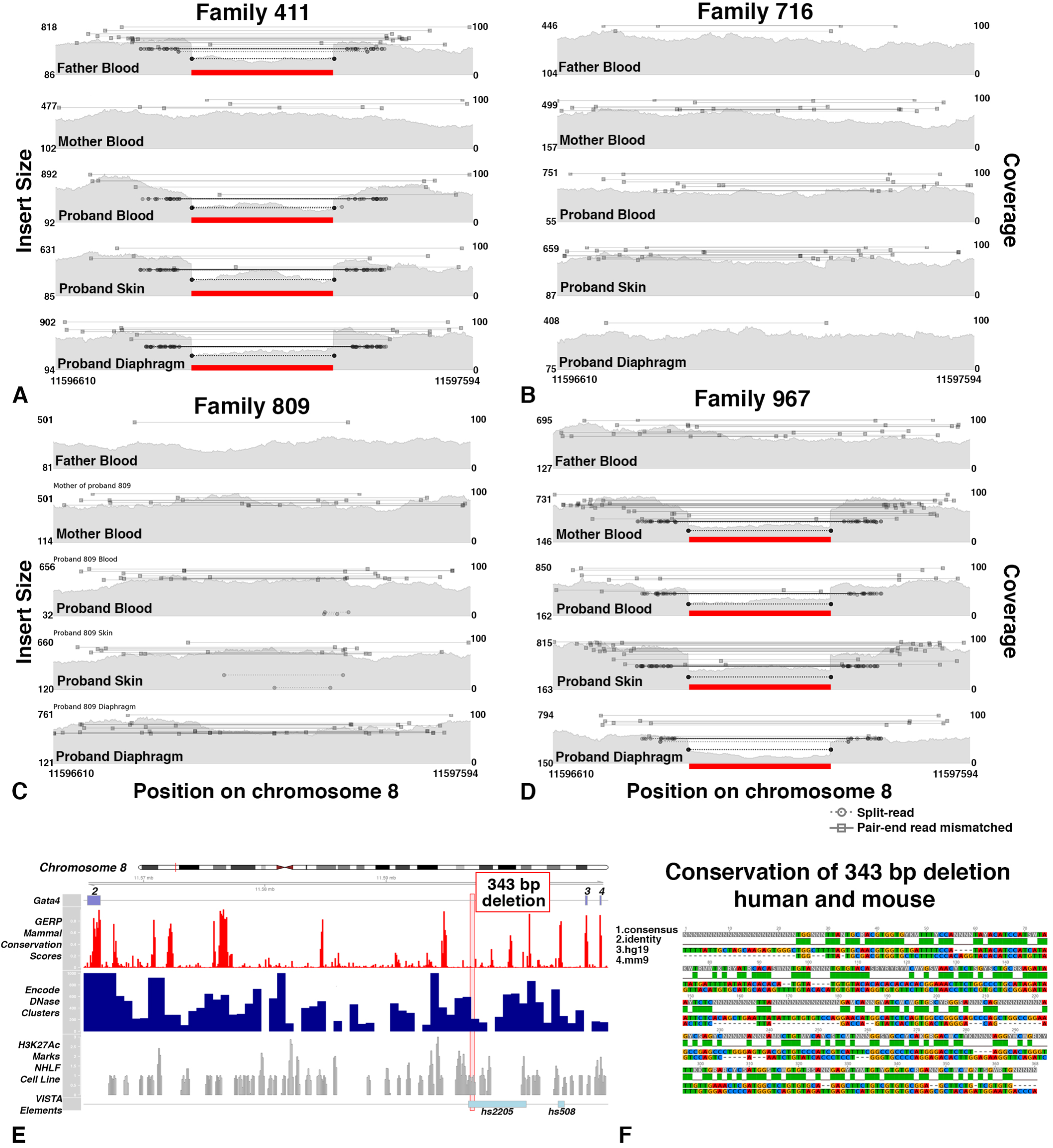
Inherited deletion in intron 2 of *GATA4*, in a region which overlaps an enhancer conserved in mouse and humans, found in CDH Probands 411 and 967. **A-D)** Samplot figures show coverage in grey (right y-axis) and Insert size between pair-end reads (left y-axis), with split and discordant reads with mis-matched insert size above 500 shown as bars. Paternal inheritance in Proband 411 (**A**) and maternal inheritance in Proband 967 (**D**) of intron 2 deletion. E) *GATA4* intron 2 and 343 bp deleted region (red box) with tracks of gene location, conservation, DNase 1 hypersensitivity and H3K27 acetylation in human lung fibroblasts. Bottom track shows enhancer element 2205 from the VISTA enhancer database within which the deletion resides. Built on the hg19 human reference genome. **F)** Pairwise alignment of 343 bp region deletion in CDH families to the homologous region in the mouse genome (mm9 reference genome). Track 1, consensus sequence; track 2, base pair identity; track 3, human (hg19) sequence; track 4, mouse minus strand sequence.

Comparison of the 343 bp deleted region in human with the orthologous region in mouse reveals that this region lies within an enhancer (element 2205) annotated in the VISTA enhancer database [Fig 5E; 95]. This enhancer drives reporter expression at E10.5 in the developing heart in a domain similar to the *GATA4* expression domain [96, 97] and demonstrates that this region is a bona fide *GATA4* enhancer. Although yet to be tested, this enhancer may also be important for *GATA4* expression in the PPF cells that are critical for diaphragm development. As a parent of proband 411 and 967 possesses the deletion and does not have any reported CDH, this deletion alone is unlikely to cause CDH in the two probands. However, this deletion may sensitize the two probands to develop CDH.

Given that two of the CDH patients in this cohort harbor the 343 bp deletion, this deletion within the *GATA4* intron 2 enhancer may be a sensitizing variant enriched in CDH individuals. To test this, we used PCR to identify the presence of the 343 bp deletion on a small cohort of 141 CDH individuals (including the 4 CDH individuals from this study) for which DNA was available. In addition, we compared the frequency of this deletion in the CDH cohort with that of a larger control population (which includes an ancestrally diverse population, but also includes individuals with common cardiovascular, neuropsychiatric, and immune-related diseases) with SVs discovered using a similar, Lumpy software-based pipeline [94]. We found that CDH individuals had an allele frequency of 0.98% (4 heterozygous individuals of 204 tested individuals, of which 141 had isolated CDH and 63 had complex CDH) and in the larger control population an allele frequency of 1.9%. These data suggest that this deletion has a roughly similar frequency of 1-2% in CDH and control populations and is a common variant in the population. This relatively common 343 bp deletion may be a sensitizing variant that, in conjunction with other variants, contributes to the genetic etiology of CDH.

## DISCUSSION

In this study, we comprehensively assessed the contribution of germline and somatic *de novo* and inherited SNVs, indels, and SVs to CDH etiology and reconstruct for the first time the genetic architecture of four individuals with isolated CDH. Our ability to perform such a comprehensive analysis of CDH, identifying genetic variants of different sizes (SNVs, indels, and SVs) in various genomic regions (exons, introns, UTR, and intergenic) with different inheritance patterns (inherited and germline and somatic *de novo*) was enabled by three factors. First, the unique collection of diaphragm sac, skin and blood samples from individuals with isolated CDH and blood from their parents allowed us to 1. confidently identify germline variants (present in all three proband samples and absent in parent samples), 2. discover *de novo* somatic variants (present in sac, but not in other samples) in the diaphragm’s connective tissue, which has previously been shown to be an important cellular source of CDH [9], and 3. determine which variants are inherited (present in all three proband samples and present in at least one parent). Second, whole genome sequencing DNA samples to an average of > 50X coverage was essential for identifying non-coding DNA regions which contribute to CDH etiology and positively calling somatic variants, which have relatively low numbers of reads. Finally, we employed a collection of computational tools that uses Illumina pair-end, whole genome sequences to discover *de novo* variants (RUFUS), identify and prioritize inherited variants (VAAST), and discover SVs (Lumpy pipeline). Altogether, the unique cohort of samples, high-depth whole genome sequences, and computational toolkit were essential to comprehensively interrogate the genetic architecture of the four CDH individuals. Our pipeline lays the groundwork for future, larger scale studies investigating the genetic etiology of CDH. In addition, our methodology should be useful for investigating other birth defects with complex genetic etiologies, such as congenital heart defects [98].

Our analysis of germline *de novo* variants revealed a potential pitfall of using blood samples to infer germline *de novo* variants and also identified a new candidate CDH-causative gene. Previous studies searching for *de novo* variants that cause CDH have used DNA from blood samples and then inferred that such variants are germline *de novo* variants [12–15]. However, while blood is the most readily available source of DNA, variants in its DNA may not have originally arisen in the germline, but instead may have arisen later in somatic cells. Our cohort, with three proband-derived tissue samples, allows us to explicitly test this alternative hypothesis. We found that an average of 1.5% of the *de novo* variants in the proband’s blood were not germline in origin (not present in all three proband tissues) and is a similar rate as found in other recent papers [58, 99]. In fact, in proband 411 a blood-specific somatic variant in *MIR4717* would have been classified as a germline variant. Thus, researchers should be cautious about inferring that all *de novo* variants in the blood are germline in origin. With DNA samples from three tissues from each CDH proband, we are able to confidently identify germline *de novo* variants because such variants will be present in all proband tissues, but not in parental samples. We found an average of 68 germline *de novo* variants per proband, and this is similar to the 70 germline *de novo* variants found in the average population [58, 100–103]. Of these, only a few are in gene-coding regions and only one of these genes, *THSD7A*, harbors a deleterious variant and is predicted to be haploinsufficient and thus is a strong candidate CDH-causative allele. A limited number of studies have investigated the function of *Thsd7a* in mice and zebrafish, focusing on its role in regulating endothelial cell migration [61–63]. As *Thsd7a* is expressed in PPF cells in mouse and migration of PPF cells is critical for normal diaphragm development and aberrant in CDH [17, 49], an intriguing hypothesis is that *Thsd7a* regulates PPF migration.

The largely discordant appearance of CDH in monozygotic twins [22, 104] and our previous mouse genetic studies [17] suggested that somatic variants in the diaphragm’s connective tissue may be a genetic feature of some CDH individuals. In particular, our mouse conditional mutagenesis experiments suggested that the connective tissue fibroblasts, which ultimately form a sac around some diaphragmatic hernias, may harbor somatic variants [17]. Previous studies of the role of somatic variants in other structural birth defects, such as congenital heart defects, have relied on blood, saliva, or skin samples and inferred that low frequency (< 30%) alternate alleles represent somatic variants that may potentially contribute to birth defect etiology [e.g. 105]. In our study, we have DNA directly derived from the proband tissue, the PPF-derived diaphragm connective sac, hypothesized to harbor somatic variants. Because we also have DNA derived from skin and blood proband samples, we were able to positively identify any alternate allele, regardless of its frequency, present in the sac, but not in skin or blood (or parental blood), as a somatic variant. Using similar logic, we were able to identify somatic variants in blood and skin. To confidently identify these somatic variants, we conservatively included alleles present in at least 20% of the reads (and not present in the other tissues). Using this strategy, we identified in three of the probands 1-7 private somatic variants in the sac, skin, and blood. Proband 809 has an aberrantly high number of somatic variants, but we currently have no mechanistic explanation (e.g. variants in DNA repair genes) for this individual’s high somatic mutational load. In all probands, the frequency of somatic alternate alleles was significantly lower than germline alleles, indicating that these variants are present in a subset of sampled cells; in the diaphragm sac, either not all PPF-derived connective tissue fibroblasts harbor the variants and/or other non-fibroblast cells are included in the sample. Because no somatic variants were shared between two of the proband tissues, all of the somatic variants must have arisen after the developmental divergence of diaphragm connective tissue, skin, and blood. Importantly, no variants are shared between blood and diaphragm. Thus, blood samples are unlikely to be informative about somatic variants in the diaphragm. Our analysis of the diaphragm revealed multiple somatic variants in the diaphragm’s connective tissue, but none in coding or annotated enhancers and so these somatic variants are unlikely to be deleterious. A previous study [25] also found no evidence of damaging somatic variants, although this study examined tissue sampled around the periphery of the herniated region and so did not specifically sample the connective tissue that mouse genetic studies [17, 34, 35] predict to be a cellular source of CDH. While our study did not find potentially damaging somatic variants in the diaphragm’s connective tissue, we have established an effective discovery strategy. A more definitive test of the role of somatic variants in CDH etiology awaits a future larger study.

The role of inherited variants in the etiology of CDH has received relatively little attention. Using VAAST, we identified multiple compound heterozygous or homozygous inherited, presumably recessive, SNVs and indel variants in all four probands. However, only a small number of these variants were rare, predicted damaging, and in genes expressed in mouse PPFs or associated diaphragm muscle progenitors. Of particular note is proband 716 who inherited multiple damaging variants, including variants in *ALG2*, *HRC*, *AHNAK*, and *MYO1H*. *AHNAK* is involved in skeletal muscle membrane structure and repair [86, 87], while *HRC* is functionally important in regulating calcium cycling in muscle [82–84]. *ALG2* regulates the function of the neuromuscular junction and organization of mitochondria in myofibers and human variants in *ALG2* have been linked to congenital myasthenic syndrome and mitochondrial myopathy [79–81]. Together, variants in these three genes may weaken muscle structure and function and lead to CDH. This is a surprising finding as mouse genetic studies have found that while variants in muscle-specific genes lead to muscle-less or diaphragms with aberrant muscle, none cause CDH (see **Table S1**). However, the inherited damaging variants in three muscle-related genes in proband 716, with an unusually large hernia, suggests that genetic alterations in muscle may lead to CDH. Another interesting aspect of the genetic profile of proband 716 is the variant in *MYO1H*. *MYO1H* regulates the function of neurons critical for sensing C02 and respiration [88], and so this *MYO1H* variant may further compound the respiratory issues introduced by CDH.

In our search for *de novo* or inherited SVs that could contribute to CDH etiology, we discovered in two probands a 343 bp deletion in intron 2 of *GATA4*, a highly ranked CDH-associated gene, that disrupts an annotated enhancer regulating *GATA4* expression [106]. We hypothesize that disruption of this enhancer leads to lower levels of *GATA4* expression. *GATA4* has notably dosage sensitive effects on heart development [107] and likely also on diaphragm development. Given that this deletion is inherited from unaffected parents and has an allele frequency of 1-2% in the general population, we hypothesize that this deletion is a relatively common SV that acts as a sensitizing allele for CDH. We hypothesize that decreased expression of *GATA4* expression resulting from the 343 bp deletion confers CDH susceptibility and in the context of other genetic variants (or environmental factors) leads to CDH. To test this hypothesis, future experiments in our lab will test in mice whether this 343bp region regulates *GATA4* expression in the PPFs and whether a deletion in this region sensitizes mice to develop CDH.

Our comprehensive analysis of the genomes of four individuals with isolated CDH allows us to reconstruct the diverse genetic architectures underlying CDH (**Fig 6**). Proband 809 is the most enigmatic of the four cases. She harbors no obvious candidate genetic variants leading to CDH, as she has no predicted damaging inherited recessive variants and one germline *de novo* missense variant in AR that is predicted to be neutral. Her genome is particularly unusual in that it contains an abnormally high somatic mutational load in her skin and diaphragm connective tissue, but the variants in the diaphragm do not affect coding or annotated enhancer regions. The source of large number of somatic mutations is unclear as she harbors no mutations in DNA repair genes. Proband 411 harbors the 343 bp intron 2 *GATA4* deletion that we hypothesize acts as a sensitizing CDH allele, but the collaborating variants that drive CDH are unclear. The germline *de novo* frameshift variant in *PEX6* is deleterious but *PEX6* is not predicted to be haploinsufficient, and only one of the inherited variants in *UPF3A* is predicted deleterious and *UPF3A* is also not predicted to be haploinsufficient. Proband 716 differs from the other probands in that she has inherited multiple rare and damaging variants in myogenic genes that likely lead to CDH. Notably, while three of the probands in our cohort have small left hernias, she is the only proband who has a large (where > 50% of the chest wall is devoid of diaphragm tissue) right hernia. Potentially, the origin of her atypical large right hernia [most hernias are found on the left side; 7] is linked to the variants in myogenic genes as opposed to genes expressed in PPF fibroblasts. Proband 967 is the individual for which we have the strongest hypothesis about the genetic origin of CDH. This individual harbors the 343 bp intron 2 *GATA4* deletion that we postulate acts as a sensitizing CDH allele and a rare germline *de novo* damaging missense variant in the haploinsufficient-intolerant gene, *THSD7A*. These two variants suggest the hypothesis that during the early development of proband 967 the PPFs of his nascent diaphragm were prone to apoptosis, unable to proliferate sufficiently [as GATA4 promotes proliferation and survival; 17], and had defects in migration (due to low expression levels *of THSD7A*) that ultimately lead to defects in diaphragm morphogenesis and CDH.

**Figure 6:**
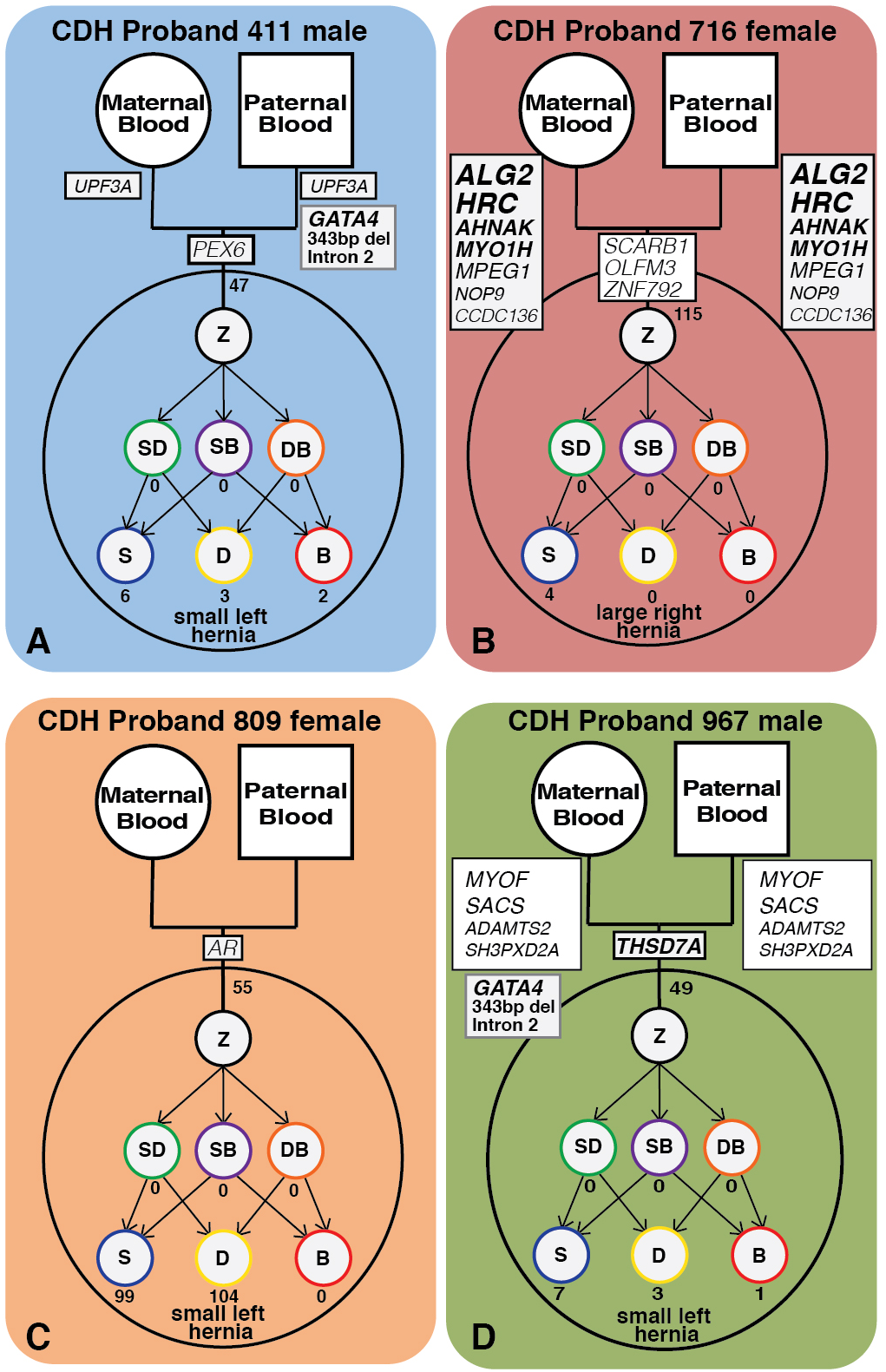
Models of four CDH probands with inherited, germline and somatic *de novo* variants, including coding, gene-associated (but non-coding), and non-gene associated SNVs, indels, and SVs. Each model highlights the genetic complexity and heterogeneity underlying CDH. Z, zygote; SD, skin-diaphragm progenitor; SB, skin-blood progenitor; DB, diaphragm-blood progenitor; S, skin cell; D, diaphragm cell; B, blood cell.

In summary, our comprehensive analysis demonstrates that the genetic etiology of CDH is heterogeneous and likely multifactorial. A challenge for future studies will be to determine whether, despite a diverse array of initiating genetic variants, a small set of molecular pathways are consistently impacted in CDH. Identification of a few key molecular pathways common to all CDH patients would be critical for designing potential *in utero* therapies to rescue or minimize the severity of herniation.

## MATERIALS AND METHODS

### Ranking of CDH-implicated genes

CDH genes reported in the literature were gathered from recent reviews [9, 46], large human patient cohort studies [12–15, 47] and recent studies [47, 48]. Original publications implicating genes in CDH were checked for the level of evidence (i.e. variants likely impacting gene function or deleterious as described in original studies) and the following ranking system used.

Mouse Data Ranking: 10 = CDH, > 80% frequency; 9 = CDH, 40-60% frequency; 8 = CDH, < 40% frequency; 7 = muscle-less patches, muscle-less diaphragm, thin diaphragm, > 80% frequency; 6 = muscle-less patches, muscle-less diaphragm, thin diaphragm, 40-60% frequency; 5 = muscle-less patches, muscle-less diaphragm, thin diaphragm, < 40% frequency. Human Data Ranking:9 = inherited compound heterozygous or homozygous deleterious variant (2 alleles in 1 gene) + de novo deleterious variant > 1 patient; 8 = inherited compound heterozygous or homozygous deleterious variant (2 alleles in 1 gene), > 1 patient; 7 = de novo deleterious variant (1 allele in 1 gene), > 1 patient; 6 = inherited compound heterozygous or homozygous deleterious variant (2 alleles in 1 gene), 1 patient; 5 = inherited deleterious variant (1 allele in 1 gene), > 1 patient; 4 = unknown inheritance deleterious variant (1 allele in 1 gene), > 1 patient; 3 = de novo deleterious variant (1 allele in 1 gene), 1 patient; 2 = inherited deleterious variant (1 allele in 1 gene), 1 patient; 1 = unknown inheritance deleterious variants (1 allele in 1 gene), 1 patient. The final ranking of each gene was determined as the sum of the mouse data and the human data ranking and constitutes the order of genes found in **Table S1**. CDH-implicated genes were queried for expression in the mouse E12.5 PPF RNA-seq dataset and expression plotted using the ggplot2 R package [108]. Gene networks within the list of CDH-associated genes were visualized using STRING [50], visualizing high and medium levels of evidence to connect gene nodes, using all evidence (except the “text mining” option was not used).

### Patient sample collection and DNA isolation

Participants were enrolled in a Columbia University IRB protocol AAAB2063 and provided informed consent/assent for participation in this study as previously described [15]. Whole genome sequencing and data analysis at the University of Utah was covered by IRB protocol 00085165. All four probands had isolated CDH. Probands 411, 809, and 967 had left hernias < 50% of chest wall devoid of diaphragmatic tissue, while proband 716 had a large (> 50% of chest wall devoid of diaphragmatic tissue) right hernia. The self-reported ancestries are probands 411 and 809 are European (non-Hispanic), proband 716 is Asian, and proband 967 is African, Hispanic, and European. The 4 CDH probands and parents are included in previous exome sequencing analysis [14].

### Whole Genome Sequencing

DNA was prepared for sequencing using TruSeq DNA PCR-free libraries (Illumina) and run on the Illumina HiSeq X Ten System at a minimum of 60× median whole-genome coverage

### Whole genome alignment, variant calling and quality checks

Genomic sequencing reads were aligned with BWA-MEM [109] against human GRCh37 reference genome (including decoy sequences from the GATK resource bundle). Aligned BAM files were de-duplicated using samblaster [110]. Base quality score recalibration and realignment of small insertions and deletions was performed with the GATK package [111]. Alignment quality was checked with samtools [112] ‘stats’ and ‘flagstats’ functions. Variants (SNVs, indels) were called with GATK Haplotypecaller [111]. Sample relatedness, sex and reported ancestry were confirmed with Peddy [55].

### Germline and somatic *de novo* SNV and indel variant analysis

*De novo* SNVs and indels were called with RUFUS (https://github.com/jandrewrfarrell/RUFUS) using standard parameters (25 length k-mers and 40 threads) and realigning k-mers to the human GRCh37 reference genome. Each proband tissue sample was run against both parents as control samples to call germline *de novo* variants, and all possible combinations of one proband tissue against the other two were run to call tissue-specific variants. Variants flagged “DeNovo” by RUFUS were retained and any variants found in all three proband tissues were called germline *de novo*. Variants were further filtered, and only variants were retained with GQ scores >20, read depths >15 at the variant site, and the variant was found in ≥ 20% of reads in the sample using annotations from the GATK haplotypecaller [111] via a python script written with the cyvcf2 package [113] and annotations in IGV [56]. Variant quality, alignment, and sample specificity were confirmed visually in IGV [56]. 5-20bp indels located at the start or end of single nucleotide repeats were filtered out, as these are potential false positives due to alignment error. Genetic locations were annotated with the UCSC Known Gene Annotation [114]. Noncoding variants were intersected against a bed file of enhancers from the VISTA Enhancer Database [106] using bedtools [115] to determine whether variants were within annotated enhancers. Coding variant predicted damage was determined by both Provean [59] and CADD scores [116] and allele frequency within a large, healthy population was determined by gnomAD [78]. Genes containing coding variants were annotated as intolerant for loss-of-function with ExAC PLi scores [60].

### Inherited SNV and indel variant analysis

The VAAST3 pipeline was used to call predicted damaging inherited recessive variants [73, 117]. The pipeline includes the following steps. First, GATK haplotypecaller [111] identified-variants were decomposed and normalized using VT [118], and variants with a GQ score < 30 or with > 25% of the samples in the VCF file not genotyped (“no-call”) were filtered out using vcftools [119]. Variant effect predicted annotations were added to the filtered VCF file using VEP [120] with version 83 of the hg19 vep-cache. As a background population, a VCF file with variants from 1000 Genomes phase 3 [74] were run through the same workflow. A VVP background database was made from the 1000 Genome filtered variants using the build_background function of VVP [121]. This background database and the filtered VCF file of variants discovered in this cohort were used as inputs to VVP to prioritize cohort-discovered variants, which were then passed to VAAST3 to be scored and ranked. Blood samples from parents and the proband were used in VAAST3’s trio recessive inherited model. Genes with a p-value > 0.05 from VAAST3 were filtered out. The remaining genes were annotated with the predicted effect from VEP, the location of the variant from VVP, and the parental origin of the allele from VVP. Ensembl IDs given in the VAAST3 output were converted to gene names using GeneCard [122]. Genes were then filtered out if 1) the genes were expressed at < 10 TPM in E12.5 mouse PPFs (human gene names were converted to mouse orthologs using OrthoRetriever (http://lighthouse.ucsf.edu/orthoretriever/)), 2) if they were a pseudogene annotated by GENCODE [75], 3) if they belonged to a highly mutable gene family, 4) if the allele called is found in > 0.01 of individuals in gnomAD, or 5) there are multiple alleles at the site in gnomAD. Variant impact was predicted by Provean [59], CADD [116] and ExAC pli scores [60], then genes were ranked based on the number and severity of damaging alleles.

### Inherited and *de novo* structural variant analysis

SVs were called with the Lumpy Smooth pipeline [89]. Regions with possible copy number variants based on read depth were called using CNVkit [91], CNVnator [92] and CN.MOPS [90] with sample BAM files. Outputs from the three tools were converted into BEDPE files and merged together as one “deletions” file (evidence of a decrease in copy number) and one “duplications” (evidence of an increase in copy number) per sample. The copy number call BEDPE files and aligned BAM files were used as input to Lumpy [89] and called with the lumpy_smooth script (in the LUMPY scripts directory). Lumpy output variants were used as inputs to SVTYPER [123] to annotate each variant as an insertion, deletion, translocation or inversion within a VCF file. *De novo* and inherited SVs were identified with GQT [124] using the LUMPY-created VCF file and a PED file describing family relations. We kept only variants with either spilt or discordant supporting read counts above 8 and but below 400 (to exclude noisy regions with high read mapping). SVs were confirmed in IGV [56], keeping variants with visible evidence in read coverage change and discordant reads. *de novo* and inherited structural variants were queried in the complete VCF file from [94] to determine allele frequency in the general population, and we filtered out SVs with an allele frequency > 10%. Variants located in intergenic regions, overlapping annotated repetitive element or elements [125] or homozygous in parental DNA were excluded. The discovered 343 bp deletion was confirmed with PCR using primers: forward, 5’-TTCCTCTACCATTGGGCGTTT-3’ and reverse, 5’-AGGTAGTACGGCTGACTTGC-3’.

### E12.5 PPF RNA sequencing

Pleuroperitoneal folds (PPFs) were isolated from wild-type E12.5 mouse embryos by cutting embryos just above the hindlimbs and below the forelimbs and removing the heart and lungs cranially, leaving the trunk with attached nascent diaphragm. The PPFs were manually dissected and stored in RNAlater (Thermo Fisher, #AM7020) at −80°c. RNA was isolated using a RNeasy Micro Kit (Qiagen, #74004), RNA quality confirmed with RNA TapeStation ScreenTape Assay (Agilent, # 5067-5576), and sequenced using TruSeq Stranded mRNA Library Preparation Kit with polyA selection (Illumina) and sequenced using HiSeq 50 Cycle Single-Read Sequencing v4 (Illumina) through the High-Throughput Genomics and Bioinformatic Analysis Shared Resource at Huntsman Cancer Institute at the University of Utah. Two biological replicates were sequenced (two pooled PPF pairs per replicated) and analyzed. Sequencing reads were aligned to the mouse genome (mm9) with STAR [126], using standard parameters. featureCounts [127] was used to count reads per gene and then normalized by TPM (transcript per million) using R.

### Figure creation and statistics

**Figures 1A, 3B, 4B**, and **S1** were created with R package ggplot2 [108]. **Fig S2** was created with Prism 7 (Graphpad). **Figures 2, 3A, 4A and 6** were created with Adobe Illustrator. **Fig 5** was created with samplot (https://github.com/ryanlayer/samplot) and Adobe Photoshop. Genome tracks were plotted using the Gviz R package [128], with UCSC Known Gene [114], GERP conservation scores [129], ENCODE DNase 1 hypersensitivity clusters, H3K27 acetylation in normal human lung fibroblasts (NHLF) cells [130] and VISTA enhancer element tracks [106]. Except for the VISTA elements (bed file downloaded from https://enhancer.lbl.gov/) all data was downloaded from the UCSC Genome Table Browser. The 343 bp deletion sequence and the homologous sequence in mouse (mm9 reference genome) were downloaded from the UCSC Genome Browser and aligned with Geneious [131]. Statistics for **Fig S1** were generated with Prism 7, using a one-way ANOVA with multiple comparisons to test differences between somatic or germline *de novo* variants found across probands or tissues and unpaired t-tests to compare somatic and germline *de novo* variants within probands or tissues.

## ACKNOWLEDGEMENTS

We would like to thank the patients and their families for their generous contribution. We are grateful for the technical assistance provided by Patricia Lanzano, Jiangyuan Hu, Jiancheng Guo, and Liyong Deng. We thank A Quinlan and RM Layer for help with LUMPY, D Neklason at Utah Genome Project for coordinating sequencing, and CY Chow, LB Jorde, A Quinlan, EM Sefton, and B Collins for critical reading of the manuscript. The support and resources from the Center for High Performance Computing at the University of Utah are gratefully acknowledged. EL Bogenschutz was supported by the University of Utah Genetics Training grant (NIH T32 GM007464). This research was supported by NIH R01HD087360, March of Dimes 6FY15203, Utah Genome Project, and Wheeler Foundation grants to GK; NIH R01GM120609 and R03HL138352 to YS; and NIH R01HD057036,UL1 RR024156, and P01HD068250 to WKC. Additional funding support was provided by grants from CHERUBS, CDHUK, and the National Greek Orthodox Ladies Philoptochos Society, Inc. and generous donations from the Williams Family, Wheeler Foundation, Vanech Family Foundation, Larsen Family, Wilke Family and many other families.

## SUPPORTING INFORMATION

Figure S1: Alternative allele read depth of somatic *de novo* variants is significantly lower than germline *de novo* variants across CDH probands (A) and tissues (B).

Figure S2: The spectrum of *de novo* variants varies between the different CDH probands (A), but does not vary between the three tissues sampled from the probands (B).

**Table S1: Genes associated with CDH in the literature.** Genes were collated from multiple studies, including family and single patient case studies, large human patient sequencing studies and functional studies using mouse genetics. A gene from human studies is included if multiple patients harbor variants in the gene, the gene is associated with other implicated genes and expressed in the developing diaphragm, and/or the gene is associated with other developmental disorders or structural birth defects that co-occur in patients with complex CDH. A gene from mouse studies is included if a variant in the gene leads to a diaphragm defect. Genes were then ranked based on the amount and type of evidence implicating their contribution to CDH (see Materials and Methods). Literature cited [11-17, 21, 23, 28, 30, 34-46, 48, 132-231]. Genes involved in GATA/TBX transcriptional network are highlighted in blue; in Hedgehog signaling in cyan; in Retinoic Acid signaling in dark green; in ROBO/SLIT signaling in purple; in MET signaling in orange; in WNT/βCATENIN signaling in yellow; and in FGF signaling in pink and ECM proteins are in light green and in muscle-related transcription factors or structural/functional genes in red.

**Table S2: DNA samples of CDH probands and their parents are of high quality and expected ancestry and relatedness.** Sheet 1 reports results of Peddy analysis of heterozygosity and indicates that sequences were of high quality and ancestry congruent with self-reported ancestry. Sheet 2 reports results of Peddy sex analysis and shows that sex of DNA samples is congruent with known sex of sample origins. Sheet 3 reports results of Peddy analysis of relatedness and shows that relatedness is as expected.

**Table S3: Germline *de novo* and somatic *de novo* variants identified by rufus in CDH probands.** True positive variants were identified by rufus and then filtered based on read depth (>15 reads total in all samples compared in the rufus call), call quality (>20 GQ score) and alternate allele frequency (sample of interest >20% alternate read frequency and other samples in comparison <20% alternate read frequency). Sheet 1: summary of all *de novo* variants identified. Sheet 2: *de novo* coding variants identified. Sheets 3-5: *de novo* gene-associated variants identified in diaphragm, skin, and blood. Sheets 6-8: *de novo* non-gene associated variants identified in diaphragm, skin, and blood.

**Table S4: Predicted gene damaging variants inherited from parents of CDH probands.** VAAST3 recessive model comparing CDH patient families to 1000 Genome phase 3 individuals was used to call predicted damaging, rare recessive variants. Genes harboring variants with a VAAST p-value <0.05 were saved as candidate genes potentially contributing to CDH (Sheet 2 shows unfiltered genes). Genes were then further filtered based on expression in PPFs (genes needed to be expressed at ≥ 10 TPM in PPFs), and pseudogenes and highly mutable genes were removed (Sheet 1 shows filtered genes).

**Table S5. Germline *de novo* and inherited structural variants possibly contributing to CDH.** Structural variants were identified using the Lumpy Smooth workflow. Germline *de novo* variants were identified by querying variants unique to the CDH proband compared to the parents, filtered based on reads that Lumpy scored as supporting the variant (>8 reads either split or discordant) and finally visually inspected in IGV to determine whether parents had evidence of the same variant. Inherited structural variants were filtered based on chromosomal regions highly associated with CDH [10]. Structural variants were visually inspected and filtered out if allele frequency > 0.1 for inherited SVs or > 0.01 for *de novo* SVs [94], if SV was located in an intergenic region, or overlapped a repetitive element [125].

