## Supplementary figures and images for "Deep whole genome sequencing of multiple proband tissues and parental blood reveals the complex genetic etiology of congenital diaphragmatic hernias"

### Supplemental FIgures

Bogenschutz et al. Fig S1

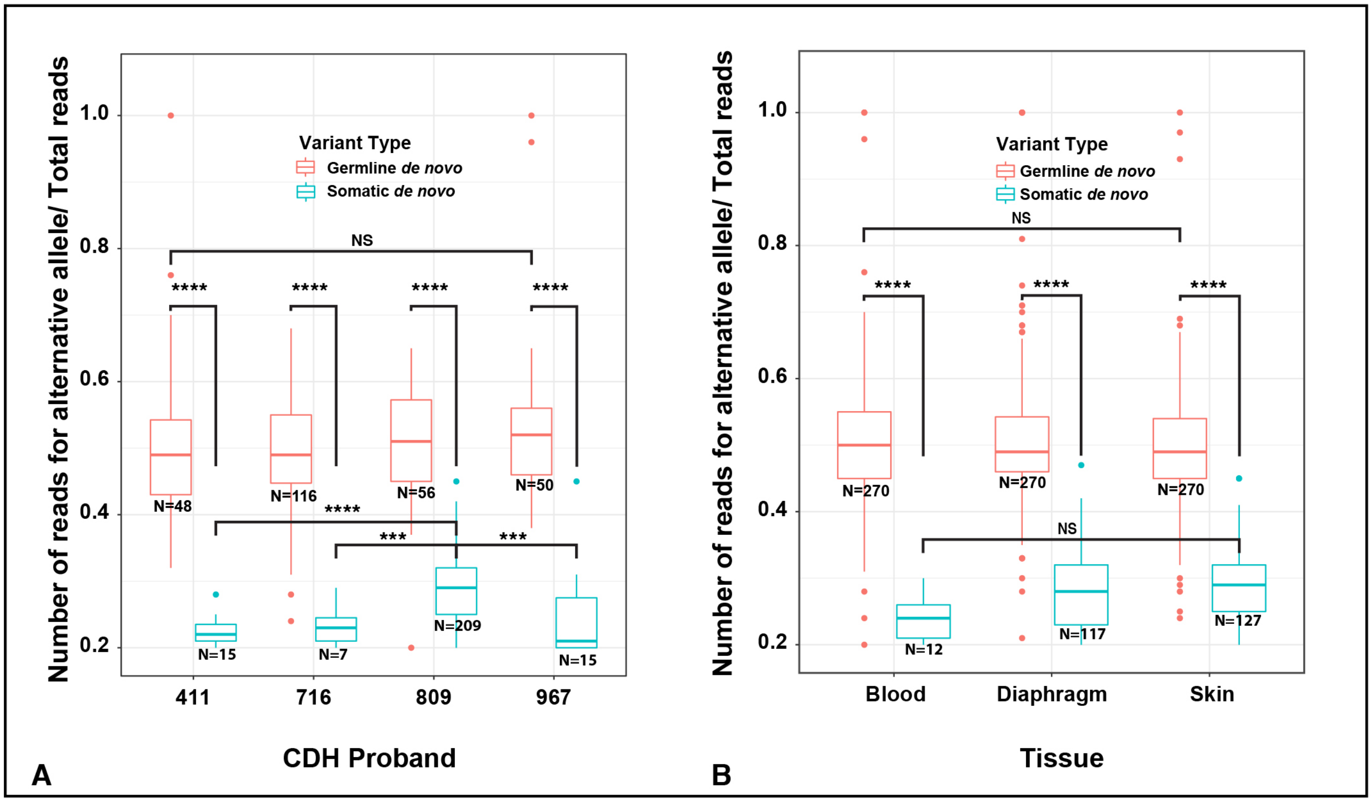

**Bogenschutz et al. Fig S2**

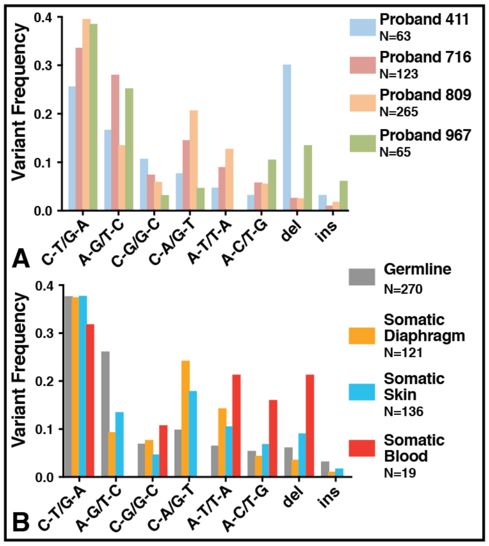
