## Supplemental Tables for "Deep whole genome sequencing of multiple proband tissues and parental blood reveals the complex genetic etiology of congenital diaphragmatic hernias"

### Table S1

CDH Implicated Genes

| | | | | | | Human Reference 1 | | | | | Human Reference 2 | | | |
|---|---|---|---|---|---|---|---|---|---|---|---|---|---|---|
| Ranking | Human Gene Name | Description | Human Genome (Hg19) coordinates | Mouse Genome (Mm9) coordinates | Isolated or Complex CDH (from human data) | Citation | De Novo Cases | Inherited Cases | Unknown inheritance | Frequency in CDH cohort | Citation | De Novo Cases | Inherited Cases | Unknown inheritance |
| 1 | ZFPM2 (FOG2) | zinc finger protein, multitype 2 | chr8<br>106331147<br>106816767 | chr15<br>40486587<br>40936138 | Isolated CDH | Longoni et al. 2015 | 1/275 | 4/275 (1 compound heterozygous patient, 2 patients with 1 paternal allelle, 1 patient with 1 maternal allele) | 3/275 | 2.9% (8/275) | Ackerman et al. 2005 | 1/30 | | |
| 2 | GATA4 | GATA binding protein 4, transcription factor | chr8<br>11561717<br>11617509 | chr14<br>63817750<br>63864108 | Isolated and Complex CDH | Yu et al., 2013 | 1/96 Isolated CDH | 1/1 (1 patient with 1 paternal allele) Complex CDH | | 1% (1/97) | Longoni et al 2012 | 1 Complex CDH | 1(1 allele inherited) Complex CDH | |
| 3 | EFEMP2 (FBLN4) | EGF containing fibulin extracellular matrix protein 2 | chr11<br>65633912<br>65640405 | chr19<br>5474734<br>5481854 | Complex CDH: Autosomal Recessive Cutis Laxa Syndrome | Hucthagowder et al 2006 | | 1 (1 homoygous patient) | | Case Study | | | | |
| 4 | FREM2 | FRAS1 Related Extracellular Matrix Protein 2 | chr13<br>39261172<br>39461267 | chr3<br>53317859<br>53461277 | Complex CDH | Jordan et al. 2018 | | 5/69 (1 compound heterozygous patient, 1 homozygous patient, 3 single inherited alleles) | | 7.2% (5/69) | | | | |
| 5 | WT1 | Wilms tumor 1 transcription factor | chr11<br>32409322<br>32457081 | chr2<br>104966685<br>105013771 | Complex CDH with Denys–Drash Syndrome, Meacham Syndrome, primary steroid-resistant nephrotic syndrome | Devriendt et al. 1995 | | | 1 | Case Study | Antonius et al. 2008 | | | 4 CDH /12 Denys-Drash Syndrome |
| 6 | SLIT3 | slit guidance ligand 3 | chr5<br>168088738<br>168728133 | chr11<br>34934957<br>35522009 | | Longoni et al. 2014 | | 2/275 (2 patients 1 paternal allele) | 1/275 | 1% (3/275) | | | | |
| 7 | KIF7 | kinesin family member 7 | chr15<br>90171200<br>90198682 | chr7<br>86842983<br>86859072 | Isolated and Complex CDH | Longoni et al. 2014 | | 1/275 (1 paternal allele; Isolated CDH) | 1/275 (Isolated CDH) | 0.7% (2/275) | Gupta et al 2016 | | | 1 (Complex CDH) |
| 8 | HLX | H2.0 like homeobox, transcription factor | chr1<br>221052743<br>221058400 | chr1<br>186551023<br>186556372 | | Slavotinek et al., 2009 | | | 4/119 CDH patients | 3% (4/119) | Longoni et al. 2014 | | | 1/275 (Complex CDH) |
| 9 | GLI2 | GLI family zinc finger 2, transcription factor | chr2<br>121554867<br>121750229 | chr1<br>120730637<br>120950196 | Isolated and Complex CDH | Longoni et al. 2014 | | 2/275 (2 patients with 1 maternal allele) | 4/275 | 2% (6/275) | | | | |
| 10 | SOX7 | SRY (sex determining region Y)-box 7, transcription factor | chr8<br>10581278<br>10588022 | chr14<br>64562542<br>64569569 | | Wat et al., 2012 | | 3/77 (1 allele from 1 parent, 2 siblings with same mutation) | 2/77 | 6.5% (5/77) | Longoni et al 2012 | | 2/226 (1 allele from 1 parent in 2 isolated CDH patients) | 1/226 (1 syndromic CDH patient) |
| 11 | FREM1 | FRAS1 related extracellular matrix protein 1 | chr9<br>14734663<br>14910993 | chr4<br>82543823<br>82698070 | Isolated and Complex CDH | Longoni et al. 2014 | | 2/275 (1 allele from 1 parent) | 2/275 | 1.5% (4/275) | | | | |
| 12 | FRAS1 | Fraser Extracellular Matrix Complex Subunit 1 | chr4<br>78978723<br>79367787 | chr5<br>96802973<br>97213747 | | Jordan et al. 2018 | | 3/69 | | 4% (3/69) | | | | |
| 13 | NR2F2 (COUPTFII) | nuclear receptor subfamily 2, group F, member 2 | chr15<br>96874111<br>96883492 | chr7<br>77496835<br>77505479 | Isolated and Complex CDH with atrial septal defect | High et al. 2016 | 1 Complex CDH | | | Case Study | Kammoun et al. 2018 | | | 1/120 |
| 14 | GLI3 | GLI family zinc finger 3, transcription factor | chr7<br>42000548<br>42276618 | chr13<br>15555555<br>15821858 | Isolated and Complex CDH with ventricular septal defect | Longoni et al. 2014 | | 1/275 (1 maternal allele; Isolated CDH) | 2/275 (1 Isolated and 1 ventricular septal defect) | 1% (3/275) | | | | |
| 15 | MET (CMET) | MET proto-oncogene, receptor tyrosine kinase | chr7<br>116312459<br>116438440 | chr6<br>17413956<br>17523980 | Isolated and Complex CDH | Longoni et al. 2014 | | 3/275 (1 patient with maternal allele, 2 patients with paternal allele: Complex CDH) | 3/275 (Isolated and Complex CDH) | 2% (6/275) | | | | |
| 16 | FGFRL1 | fibroblast growth factor receptor like 1 | chr4<br>1005760<br>1020686 | chr5<br>109123247<br>109135969 | Isolated and Complex CDH, Wolf-Hirschhorn syndrome | Tautz et al. 2010 | | | 1 | Case Study of Wolf-Hirschhorn Syndrome | van Dooren et al. 2004 | | | 1 |
| 17 | RARB | retinoic acid receptor beta | chr3<br>25469834<br>25639422 | chr14<br>17263353<br>17914845 | Complex CDH with PDAC syndrome | Srour et al., 2013 | 3/17 (2 with same mutation) with PDAC syndrome | 2/17 (2 compound heterozygous siblings) with PDAC syndrome | | 30% (5/17 with PDAC syndrome) | | | | |
| 18 | FBN1 | fibrillin 1 | chr15<br>48700503<br>48937985 | chr2<br>125126329<br>125332174 | Complex CDH with Marfan syndrome | Jacobs et al., 2002 | | | 1 | Marfan Case Study | Revencu et al., 2004 | | | 1 |
| *19* | PDGFRA | platelet derived growth factor receptor alpha | chr4<br>55095264<br>55164412 | chr5<br>75552190<br>75594231 | Isolated and Complex CDH | Bleyl et al., 2007 | | | 3/96 | 3% (3/96) | Longoni et al 2014 | | patient with maternal allele, 1 Isolated patient with paternal | 4/275 (1 Isolated, 3 Complex) |
| 20 | HOXB4 | homeobox B4, transcription factor | chr17<br>46652869<br>46655743 | chr11<br>96179580<br>96182952 | Complex CDH | Longoni et al. 2014 | | | 2/275 (Complex CDH) | 0.7% | | | | |
| 21 | NEDD4 | neural precursor cell expressed, developmentally down-regulated 4, E3 ubiquitin protein ligase | chr15<br>56119116<br>56285944 | chr9<br>72541104<br>72597657 | Complex CDH | Longoni et al. 2014 | | 3/275 (1 patient with paternal allele, 2 patients with maternal allele: Complex CDH) | 1/275 (Complex CDH) | 1.5% (4/275) | | | | |
| 22 | CTNNB1 | catenin beta 1 | chr3<br>41240941<br>41281939 | chr9<br>120842695<br>120869625 | | | | | | | | | | |
| 23 | SIX4 | sine oculis-related homeobox 4, transcription factor | chr14<br>61176255<br>61190852 | chr12<br>74209218<br>74214412 | Isolated and Complex CDH | Longoni et al. 2014 | | 2/275 (1 Complex CDH patient with paternal allele, 1 Isolated CDH with maternal allele) | | 0.7% (2/275) | | | | |
| 24 | CTBP2 | C-terminal binding protein 2 | chr10<br>126676418<br>126716453 | chr7<br>140178693<br>140206930 | Isolated and Complex CDH | Longoni et al. 2014 | | 4/275 (1 Isolated CDH with maternal allele, 1 Isolated CDH with paternal allele, 2 complex CDH with maternal allele) | 3/275 (2 Isolated, 1 Complex) | 2.5% (7/275) | Kammoun et al. 2018 | | | 13/120 (Isolated CDH) |
| 25 | PAX7 | paired box 7, transcription factor | chr1<br>18957499<br>19075360 | chr4<br>139293995<br>139388883 | Isolated and Complex CDH | Longoni et al 2014 | | 2/275 (1 Complex CDH parent with paternal allele, 1 Complex CDH parent with maternal allele, ) | 1/275 (Isolated CDH) | 1% (3/275) | | | | |
| 26 | TBX5 | T-box 5, transcription factor | chr12<br>114791734<br>114846247 | chr5<br>120284671<br>120335227 | Isolated and Complex CDH | Longoni et al. 2014 | | 1/2751 (1 Complex CDH patient with paternal allele) | 2/275 (Isolated) | 1% (3/275) | Kammoun et al 2018 | | | 1/120 (Complex) |
| 27 | CHAT | choline acetyltransferase | chr10<br>50822082<br>50873150 | chr14<br>33221388<br>33279095 | | | | | | | | | | |
| 28 | RARA | retinoic acid receptor alpha | chr17<br>38465423<br>38513895 | chr11<br>98821784<br>98836256 | Isolated CDH | Longoni et al. 2014 | | 1/275 (1 paternal allele: Isolated CH) | 2/275 (Isolated CDH) | 1% (3/275) | | | | |
| 29 | ROBO2 | roundabout guidance receptor 2 | chr3<br>77147162<br>77699114 | chr16<br>73892550<br>74353124 | Isolated and Complex CDH | Longoni et al. 2014 | | 1/275 (1 paternal allele; Complex CDH) | 2/275 (Isolated CDH) | 1% (3/275) | | | | |
| 30 | MMP2 | matrix metallopeptidase 2 | chr16<br>55513081<br>55540586 | chr8<br>95351195<br>95377320 | Isolated and Complex CDH | Longoni et al. 2014 | | 2/275 (1 Isolated CDH patient with maternal allele, 1 Complex CDH patient with maternal allele) | | 0.7% (2/275) | | | | |
| 31 | MMP14 | matrix metallopeptidase 14 | chr14<br>23305793<br>23316803 | chr14<br>55050434<br>55061293 | Isolated and Complex CDH | Longoni et al. 2014 | | 2/275 (1 Isolated CDH patient with maternal allele, 1 Complex CDH patient with paternal allele) | | 0.7% (2/275) | Kammoun et al. 2018 | | | 9/120 |
| 32 | NDST1 | N-deacetylase/N-sulfotransferase (heparan glucosaminyl) 1 | chr5<br>149877339<br>149937773 | chr18<br>60844147<br>60908074 | | | | | | | | | | |
| 33 | DNASE2A (DNASE2) | deoxyribonuclease II alpha | chr19<br>12986025<br>12992335 | chr8<br>87432522<br>87435360 | Complex CDH | Longoni et al. 2014 | | | 1/275 (Complex) | 0.4% (1/275) | | | | |
| 34 | STRA6 | stimulated by retinoic acid 6 | chr15<br>74471808<br>74495220 | chr9<br>57976894<br>58001804 | Complex CDH, PDAC (pulmonary, diaphragm, anopthalmia, and cardiac defects) | Chassaing et al., 2013 | | 1/5 Complex CDH (PDAC) with homozygous splice mutation | | 20% (1/5 PDAC) | Pasutto et al. 2007 | | homozygous, 1 compound heterozygous | |
| 35 | LTBP4 | latent transforming growth factor beta binding protein 4 | chr19<br>41103140<br>41135725 | chr7<br>28090159<br>28118667 | Complex CDH | Urban et al., 2009 | (compound heterozygous, at least 1 allele de | 2 Complex CDH (1 homozygous, 2 compound heterozygous) | | 3 Case Studies | | | | |
| 36 | EFNB1 | ephrin B1, ligand for Eph receptors | chrX<br>68048840<br>68062006 | chrX<br>96331399<br>96344361 | Complex, X-linked Craniofrontonsal Syndrome | Hogue et al., 2010 | | 1 Complex CDH with X-linked maternal allele | | Case Study | Twigg et al., 2004 | patient with craniofrontal syndrome and | | |
| 37 | PORCN | porcupine O-acyltransferase | chrX<br>48367346<br>48379202 | chrX<br>7770971<br>7783651 | Complex, X-linked Human Focal Dermal Hypoplasia Syndrome, Goltx-Gorlin Syndrome | Maas et al., 2009 | | 2 Complex CDH with Goltz-Gorlin syndrome with X-linked maternal allele | | | Smigiel et al. 2011 | with Goltz-Gorlin syndrome with de novo X-linked | | |
| 38 | MYH10 | myosin, heavy polypeptide 10, non-muscle | chr17<br>8377522<br>8534079 | chr11<br>68505416<br>68630126 | Complex CDH | Tuzovic et al., 2013 | 1 | | | Case Study | | | | |
| 39 | LRP2 | low density lipoprotein receptor-related protein 2 | chr2<br>169983619<br>170219122 | chr2<br>69262391<br>69424086 | Complex CDH, Donnai-Burrows syndrome | Kantarci et al., 2007 | | 7/7 (patients = either homozygous or compound heterozygous) | | 7 families with Donnai-Barrow syndrome children | | | | |
| 40 | EYA1 | EYA transcriptional coactivator and phosphatase 1 | chr8<br>72109667<br>72274467 | chr1<br>14159038<br>14300280 | Isolated and Complex CDH | Longoni et al., 2014 | 1/275 (Complex) | 1/275 (1Isolated CDH patient with paternal allele) | | 0.7% (2/275) | Kammoun et al. 2018 | | | 9/120 (Isolated CDH) |
| 41 | LOX | lysyl oxidase | chr5<br>121398889<br>121412918 | chr18<br>52675713<br>52689375 | | | | | | | | | | |
| 42 | MYOD1 | myogenic differentiation 1, bHLH transcription factor | chr11<br>17741109<br>17743678 | chr7<br>53631843<br>53634462 | Isolated CDH | Longoni et al. 2014 | | 1/275 (1 Isolated CDH patient with paternal allele) | 1/275 (Isolated CDH) | 0.7% (2/275) | Kammoun et al. 2018 | | | 6/120 Isolated CDH |
| 43 | ILF3 (N90) | interleukin enhancer binding factor 3 | chr19<br>10764937<br>10803095 | chr9<br>21172314<br>21209807 | Isolated and Complex CDH | Longoni et al. 2014 | | 1/275 (1 Isolated CDH with maternal allele) | 1/275 (Complex CDH) | 0.7% (2/275) | | | | |
| 44 | PAX3 | paired box 3, transcription factor | chr2<br>223064605<br>223163715 | chr1<br>78097841<br>78193711 | Isolated CDH | Longoni et al 2014 | | | 1/275 | 0.4% (1/275) | | | | |
| 45 | GPC3 | glypican 3 | chrX<br>132669776<br>133119673 | chrX<br>49625603<br>49967151 | Complex CDH, X-linked Simpson-Golabi-Behmel Syndrome | Yano et al., 2011 | | 1/4 (4 siblings with maternal allele, only 1 sibling with CDH) | | Case Study | Hughes-Benzie et al., 1996 | | Golabi-Behmel Syndrome with inherited GPC3 | |
| 46 | WDR35 | WD repeat domain 35 | chr2<br>20110023<br>20189884 | chr12<br>8980806<br>9035653 | | | | | | | | | | |
| 47 | TMEM70 | transmembrane protein 70 | chr8<br>74888376<br>74895018 | chr1<br>16655271<br>16668356 | Complex CDH | Sarajlija et al 2017 | | 1 Complex CDH homozygous mutation | | Case Study | Catteruccia et al. 2014 | | or compound heterozyotes with TMEM70 mutation | |
| 48 | EYA2 | EYA transcriptional coactivator and phosphatase 2 | chr20<br>45523262<br>45817492 | chr2<br>165420527<br>165597227 | Complex CDH | Longoni et al. 2014 | | 1/275 (1 Complex CDH patient with maternal allele) | 1/275 (Complex CDH) | 0.7% (2/275) | | | | |
| 49 | MYOG | myogeninic factor 4, bHLH transcription factor | chr1<br>203052256<br>203055166 | chr1<br>136186580<br>136189125 | | | | | | | | | | |
| 50 | LMNB1 | lamin B1 | chr5<br>126112314<br>126172712 | chr18<br>56867466<br>56913079 | | | | | | | | | | |
| 51 | LMNB2 | lamin B2 | chr19<br>2428162<br>2456966 | chr10<br>80364107<br>80369157 | | | | | | | | | | |
| 52 | MNX1 (HB9) | motor neuron and pancreas homeobox 1, transcription factor | chr7<br>156797546<br>156803347 | chr5<br>29800362<br>29805010 | | | | | | | | | | |
| 53 | HSPG2 | heparan sulfate proteoglycan 2 (perlecan) | chr1<br>22148724<br>22263790 | chr4<br>137024717<br>137126545 | | Yu et al. 2015 | 1/39 | | | 2.6% (1/39) | Longoni et al. 2017 | 1/87 | | |
| 54 | MYRF | myelin regulatory factor, transcription factor | chr11<br>61522860<br>61555990 | chr19<br>10282760<br>10315238 | Complex CDH | Qi et al. 2018 | 4/271 (Complex CDH) | | | 1.5% (4/271) | Rossetti et al. 2019 | | | 3 |
| 55 | PBX1 | PBX homeobox 1, transcription factor | chr1<br>164528596<br>164821060 | chr1<br>170049494<br>170362300 | Complex CDH | Kammoun et al. 2018 | | | 1/120 (Complex CDH) | 0.8% (1/120) | | | | |

### Table S1

| Ranking | Human Gene Name | Frequency in CDH Cohort | Human reference 3 Citation | De Novo Cases | Inherited Cases | Unknown inheritance | Frequency in CDH Cohort | Human Reference 4 Citation | De Novo Cases | Inherited Cases | Unknown inheritance | Frequency in CDH Cohort | Human Reference 5 Citation | De Novo Cases | Inherited Cases | Unknown inheritance | Frequency in CDH Cohort | Citation |
|---|---|---|---|---|---|---|---|---|---|---|---|---|---|---|---|---|---|---|
| 1 | ZFPM2 (FOG2) | 3.3% (1/30) | Bleyl et al. 2007 | | | 4/96 | 4.2% (4/96) | Qi et al. 2018 | 1/362 | | | 0.3% (1/362) | Kammoun et al. 2018 | | 2/120 | 3/120 | 4.2% (5/120) | |
| 2 | GATA4 | 2.3% 2/87 | Longoni et al. 2014 | | 1/275 (1 allele inherited) Complex | | 0.4% (1/275) | Wat et al. 2012 | | 1(1 paternal allele) | 11 | 15.6% (12/77) | Kammoun et al. 2018 | 1/120 Isolated CDH | | 1/120 Isolated CDH | 1.7% (2/120) | Arrington et al. 2012 |
| 3 | EFEMP2 (FBLN4) | | | | | | | | | | | | | | | | | |
| 4 | FREM2 | | | | | | | | | | | | | | | | | |
| 5 | WT1 | 30% (4/12 Denys-Drash) | Suri et al. 2007 | Meacham Syndrome | | 1 with Meacham Syndrome | | Denamur et al 2000 | | | primary steroid-resistant | 2.7% (1/37) primary steroid-resistant nephrotic syndrome | Qi et al. 2018 | 2/362 Isolated CDH | | | 0.6% (2/362) | |
| 6 | SLIT3 | | | | | | | | | | | | | | | | | |
| 7 | KIF7 | Case Study | | | | | | | | | | | | | | | | |
| 8 | HLX | 0.4% (1/275) | | | | | | | | | | | | | | | | |
| 9 | GLI2 | | | | | | | | | | | | | | | | | |
| 10 | SOX7 | 1.3% (3/226) | | | | | | | | | | | | | | | | |
| 11 | FREM1 | | | | | | | | | | | | | | | | | |
| 12 | FRAS1 | | | | | | | | | | | | | | | | | |
| 13 | NR2F2 (COUPTFII) | 0.8% (1/120; Isolated CDH) | | | | | | | | | | | | | | | | |
| 14 | GLI3 | | | | | | | | | | | | | | | | | |
| 15 | MET (CMET) | | | | | | | | | | | | | | | | | |
| 16 | FGFRL1 | Case Study of Wolf-Hirschhorn Syndrome | Longoni et al. 2014 | | CDH patient with maternal allele, 1 Complex CDH | | 0.7% (2/275) | | | | | | | | | | | |
| 17 | RARB | | | | | | | | | | | | | | | | | |
| 18 | FBN1 | Marfan Case Study | Stheneur et al., 2011 | | | patients with FBN1 mutation | 10% 6 CDH/60 patients with FBN1 mutation | | | | | | | | | | | |
| *19* | PDGFRA | 2% (6/275) | | | | | | | | | | | | | | | | |
| 20 | HOXB4 | | | | | | | | | | | | | | | | | |
| 21 | NEDD4 | | | | | | | | | | | | | | | | | |
| 22 | CTNNB1 | | | | | | | | | | | | | | | | | |
| 23 | SIX4 | | | | | | | | | | | | | | | | | |
| 24 | CTBP2 | 11% (13/120) | | | | | | | | | | | | | | | | |
| 25 | PAX7 | | | | | | | | | | | | | | | | | |
| 26 | TBX5 | 0.8% (1/120) | | | | | | | | | | | | | | | | |
| 27 | CHAT | | | | | | | | | | | | | | | | | |
| 28 | RARA | | | | | | | | | | | | | | | | | |
| 29 | ROBO2 | | | | | | | | | | | | | | | | | |
| 30 | MMP2 | | | | | | | | | | | | | | | | | |
| 31 | MMP14 | 7.5% (9/120) | | | | | | | | | | | | | | | | |
| 32 | NDST1 | | | | | | | | | | | | | | | | | |
| 33 | DNASE2A (DNASE2) | | | | | | | | | | | | | | | | | |
| 34 | STRA6 | 5 case studies | Longoni et al. 2014 | | 1/275 (Isolated CDH) | | 3.6% (1/285) | Chassaing et al. 2009 | | | patients with CDH, all homozygous | report, Golzio et al. 2007 and West et al. 2009 | | | | | | |
| 35 | LTBP4 | | | | | | | | | | | | | | | | | |
| 36 | EFNB1 | | Vasudevan et al., 2006 | | patient with craniofrontonasal syndrome and | | | Twigg et al., 2006 | | children with Complex CDH and | | | | | | | | |
| 37 | PORCN | | Dias et al. 2010 | | | CDH patient with Focal Dermal | Case Study | | | | | | | | | | | |
| 38 | MYH10 | | | | | | | | | | | | | | | | | |
| 39 | LRP2 | | | | | | | | | | | | | | | | | |
| 40 | EYA1 | 7.5% (9/120) | | | | | | | | | | | | | | | | |
| 41 | LOX | | | | | | | | | | | | | | | | | |
| 42 | MYOD1 | 5% (6/120) | | | | | | | | | | | | | | | | |
| 43 | ILF3 (N90) | | | | | | | | | | | | | | | | | |
| 44 | PAX3 | | | | | | | | | | | | | | | | | |
| 45 | GPC3 | | Veugelers et al., 2000 | | | Simpson-Golabi-Behmel | | Li et al., 2001 | | Simpson Golabi Behnel Syndrome | | | | | | | | |
| 46 | WDR35 | | | | | | | | | | | | | | | | | |
| 47 | TMEM70 | | | | | | | | | | | | | | | | | |
| 48 | EYA2 | | | | | | | | | | | | | | | | | |
| 49 | MYOG | | | | | | | | | | | | | | | | | |
| 50 | LMNB1 | | | | | | | | | | | | | | | | | |
| 51 | LMNB2 | | | | | | | | | | | | | | | | | |
| 52 | MNX1 (HB9) | | | | | | | | | | | | | | | | | |
| 53 | HSPG2 | 1% (1/87) | | | | | | | | | | | | | | | | |
| 54 | MYRF | | | | | | | | | | | | | | | | | |
| 55 | PBX1 | | | | | | | | | | | | | | | | | |

### Table S1

CDH Implicated Genes

| | | Human Reference 6 | | | | | Mouse data 1 | | | | |
|---|---|---|---|---|---|---|---|---|---|---|---|
| Ranking | Human Gene Name | De Novo Cases | Inherited Cases | Unknown inheritance | Frequency in CDH Cohort | Citation | Mouse Genetic Allele | Evidence | Frequency | Citation | Mouse Genetic Allele |
| 1 | ZFPM2 (FOG2) | | | | | Ackerman et al. 2005 | Zfpm2lil/lil germline LOF, posterior lateral CDH | ENU screen found point mutation leading to a splice site variant. Abnormal diaphragmatic muscularization and pulmonary hypoplasia in all mutant mice (n >25). Fig 1 | 100% | | |
| 2 | GATA4 | | 1 (GATA4 intron 1 rare variant) in 7 inter-related patients | | | Jay et al 2007 | Gata4del2/+ germline LOF, Central CDH (sac) | Diaphragm defects (frank herniation or aberrant fusion of central tendon to liver) in 6/21 (29%). Overt herniation in 3/21 (14%) Fig 6 | 43% (9/21) | Merrell et al. 2015 | $Prx1Cre^{Tg/+};Gata4^{del3-5/fl3-5}$ Conditional LOF in PPFs |
| 3 | EFEMP2 (FBLN4) | | | | | Horiguchi et al 2009 | Fbln4 Del Ex2/Del Ex2 germline LOF | Mutants die due to diaphragmatic hernias. FigS3 | 100% | | |
| 4 | FREM2 | | | | | Jordan et al. 2018 | Frem2ne/ne germline LOF | 8% of Frem2ne/ne with CDH. Fig 2- 3 | 8% | | |
| 5 | WT1 | | | | | al., 1993, Clugston et al. 2006 | WT1del1/del1germline LOF, Posterior CDH. | Kreidberg et al: 3/3 posterior diaphragm defect. Fig 2. Clugston et al. 2006 Left posterolateral defects. Fig 1 | 100% | Paris et al. 2015 | WT1del1/del1 germline LOF |
| 6 | SLIT3 | | | | | Yuan et al., 2003 | Slit3 lacZ/LacZ germline LOF | Of mutants that die before 9 months, 90% with a central CDH with liver herniated into thoracic cavity Fig. 2-3 | 90% | | |
| 7 | KIF7 | | | | | Coles and Ackerman, 2013 | Kif7dd/dd germline nonsense mutation resulting in a truncated protein | Posterior CDH 125/125. Fig 1 and 3. | 100% (125/125) | | |
| 8 | HLX | | | | | Hentsch et al., 1996 | HlxNeo/Neo germline LOF | CDH in all mutants, no numbers. Data not shown. | 100% | | |
| 9 | GLI2 | | | | | Kim et al., 2001 | Gli2Neo/Neo germline LOF and Gli2Neo/Neo;Gli3del/+ mice germline LOF | Gli2neo/neo 1/9 with CDH; Gli3del/del 2/9 with CDH; Gli2neo/neo;Gli3del/+ 2/4 with CDH. Fig 1. | 11% (1/9) | | |
| 10 | SOX7 | | | | | Wat et al. 2012 | Sox7del2/+ germline LOF | Soxdel2/+ 10/71 had CDH in ventral midline. Fig 2. | 14% (10/71) | | |
| 11 | FREM1 | | | | | Beck et al., 2013 | Frem1eyes2/eyes2 germline LOF | Frem1 eyes2/eyes2 15/32 with CDH on B6Brd/129S6 background or 6/73 on C57BL/6J background | 8.2- 46.9% (6/73 or 15/32) | | |
| 12 | FRAS1 | | | | | Jordan et al. 2018 | Fras1Q1263*/Q1263* germline | 1% of Fras1Q1263*/Q1263* ventral CDH. Fig 2-3. | 1% | | |
| 13 | NR2F2 (COUPTFII) | | | | | You et al. 2005 | Nkx3.2Cre/+;Nr2f2fl1-3/fl1-3 Conditional LOF in foregut mesentery | In E14.5–E18.5 embryos, 11/24 (45.8%) with CDH, spleen defect in 24/24 (100%). 7/9 (78%) dead with CDH; all mutants asplenic. Fig 1 -2. | 55% (18/33) | | |
| 14 | GLI3 | | | | | Kim et al., 2001 | Gli2Neo/Neo germline LOF and Gli2Neo/Neo;Gli3del/+ mice germline LOF | Gli2neo/neo 1/9 with CDH; Gli3del/del 2/9 with CDH; Gli2neo/neo;Gli3del/+ 2/4 with CDH. Fig 1. | 22% (2/9) | | |
| 15 | MET (CMET) | | | | | Babiuk and Greer, 2002 | Metneo/neo germline LOF | Muscleless diaphragm. Frequency not reported. Fig.1. | | Bladt et al. 1995 | Met-/- germline LOF |
| 16 | FGFRL1 | | | | | Baertschi et al., 2007 | Frfrl1del1-2/del1-2 germline LOF | Amuscular posterior regions and thinner diaphragm which bulged from underlying liver in mutants. No frequency reported. Fig 3. | | Catela et al., 2009 | Frfrl1del3-7/del3-7 germline LOF |
| 17 | RARB | | | | | Mendelsohn et al., 1994 | RARa neo/neo;RARb2 neo/neo germline LOF | RARαneo/neo;RAR β2+/neo 1/4 with hernias and RARαneo/neo;β2neo/neo showed 1/7 with hernias. No phenotype reported with RARB alone. Fig 1 | | | |
| 18 | FBN1 | | | | | Pereira et al., 1999 | Fbn1mgR/mgR germline hypomorph | CDH, data nor frequency shown | | | |
| *19* | PDGFRA | | | | | Bleyl et al., 2007 | Pgfradel2-4/del2-4 germline LOF, Failed or delayed PPF expansion | 7/17 mutants at E12 had diaphragmatic defects. Fig 1. | 41% (7/17) | | |
| 20 | HOXB4 | | | | | Ramirez-Solis et al., 1993 | Hoxb4Neo/Neo germline LOF | 3/25 CDH or muscleless regions in retrosternal region. Data not shown | 12% (3/25) | | |
| 21 | NEDD4 | | | | | Liu et al. 2009 | Nedd4 Gt(IRESBetageo)249Lex/Gt(IRESBetageo)249Lex germline LOF | Thin diaphragms with wavy and disorganized muscle fibers in all mutants (15/15). Fig 2. | 100% 15/15 | | |
| 22 | CTNNB1 | | | | | Paris et al., 2015 | Wt1CreERT2/+;Ctnnb1 fl2-6/fl2-6 Conditional LOF in PPFs | Posterior CDH in all mutants. Fig 3. | 100% | | |
| 23 | SIX4 | | | | | Grifone et al., 2005 | Six1tm1Mair/Six1tm1Mair; Six4tm1Mair/Six4tm1Mair germline LOF. | Double mutants amuscular diaphragm. Frequency not reported. Fig. 3. | | | |
| 24 | CTBP2 | | | | | Hildebrand et al., 2002 | Ctbp1Neo/Neo;Ctbp2Geo/+ germline LOF mutation | Many fewer myofibers in diaphragm. Frequency not reported. Fig. 5. | | | |
| 25 | PAX7 | | | | | Seale et al., 2000 | Pax7LacZ/LacZ germline LOF mice | Thin diaphrams. Frequency not reported. Fig. 3 | | | |
| 26 | TBX5 | | | | | Valasek et. al., 2011 | Prx1Cre;Tbx5fl/fl conditional LOF in PPFs | Muscleless diaphragm. Frequency not reported. Fig. 6. | | | |
| 27 | CHAT | | | | | Misgeld et al., 2002 | Chatneo/neo germline LOF deletion | Underdeveloped muscle and central tendon hernia in >85% mutants. Fig 2 | >85% | | |
| 28 | RARA | | | | | Mendelsohn et al., 1994 | RARa neo/neo;RARb2 neo/neo germline LOF | RARαneo/neo;RAR β2+/neo 1/4 with hernias and RARαneo/neo;β2neo/neo showed 1/7 with hernias. No phenotype reported with RARA alone. Fig 1 | | | |
| 29 | ROBO2 | | | | | Domyan et al., 2013 | Robo1 b-geo/b-geo;Robo2 lacZ/LacZ germline LOF | Posterior CDH in double mutants. No phenotype reported in single mutants. Frequency not reported. Fig 1. | | | |
| 30 | MMP2 | | | | | Oh et al., 2004 | Mmp14LacZ/LacZ;Mmp2Neo/Neo germline LOF | MMP2 mutants no diaphragm phenotype. Double mutants have thin diaphragms. Frequency not reported. Fig. 1- 5 | | | |
| 31 | MMP14 | | | | | Oh et al., 2004 | Mmp14LacZ/LacZ;Mmp2Neo/Neo targeted germline LOF mice present with thin diaphragms | MMP14 mutants no diaphragm phenotype. Double mutants have thin diaphragms. Frequency not reported. Fig. 1-5 | | | |
| 32 | NDST1 | | | | | Zhang et al., 2014 | Tie2CreTg/+;Ndst1fl2/fl2 Conditional LOF in endothelial cells | Postnatal central tendon CDH. Frequency 25-60%, increase with postnatal age. Fig. 1. | 25-60% | | |
| 33 | DNASE2A (DNASE2) | | | | | Krieser et al., 2002 | Dnase2aNeo/Neo germline LOF | Thin and herniated diaphragm in mutant. Frequency not reported. | | | |
| 34 | STRA6 | | | | | | | | | | |
| 35 | LTBP4 | | | | | | | | | | |
| 36 | EFNB1 | | | | | | | | | | |
| 37 | PORCN | | | | | | | | | | |
| 38 | MYH10 | | | | | Tuzovic et al., 2013 | HMHC IIB-B-/B- germline LOF | Anterior diaphragmatic hernia. No data shown. Frequency not reported. | | | |
| 39 | LRP2 | | | | | | | | | | |
| 40 | EYA1 | | | | | Grifone et al. 2007 | Eya1Neo/Neo; Eya2Neo/+ germline LOF | Double mutants have "no diaphragm". Data not shown. Frequency not reported. | | | |
| 41 | LOX | | | | | Hornstra et al., 2003 | Loxneo/neo germline LOF | Central diaphragm hernia. Frequency not reported. Fig 2. | | | |
| 42 | MYOD1 | | | | | Inanlou et al., 2003 | MyoDneo/neo; Dmdmdx/mdx germline LOF for MyoD and dystrophin | Double mutants with reduced diaphragm. Phenotype MyoD not reported. Fig. 3. | Not reported | | |
| 43 | ILF3 (N90) | | | | | Shi et al., 2005 | N90Neo/Neo germline LOF mutant mice | Thin diaphragm with fewer myofers. Frequency not reported. Fig. 3-4. | Not reported | | |
| 44 | PAX3 | | | | | Li et al., 1999 | Pax3Sp/Sp germline LOF | Amuscular diaphragm, Fig. 4. Frequency not reported. | | Merrell et al. 2015 | Pax3 SpD/SpD |
| 45 | GPC3 | | | | | | | | | | |
| 46 | WDR35 | | | | | Mill et al. 2011 | Wdr35 yeti/yeti germline LOF | Diaphragmatic hernias. Frequency not reported. Fig. 1. | | | |
| 47 | TMEM70 | | | | | | | | | | |
| 48 | EYA2 | | | | | Grifone et al. 2007 | Eya1Neo/Neo; Eya2Neo/+ germline LOF | Double mutants have "no diaphragm". Data not shown. Frequency not reported. | | | |
| 49 | MYOG | | | | | Tseng et al., 2000 | Myogneo/neo germline LOF | Amuscular diaphragms, no CDH 6/6 | 100% (6/6) | Hasty et. Al 1993 | mygml/mygml mice germline LOF |
| 50 | LMNB1 | | | | | Kim et al. 2011 | Lmnb1 tm1Yxz/Lmnb1 tm1Yxz, Lmnb2 tm1Yxz/Lmnb2 tm1Yxz single and double germline LOF | Single and double mutants thin diaphragms. Penetrant. Fig. S10 | | | |
| 51 | LMNB2 | | | | | Kim et al. 2011 | Lmnb1 tm1Yxz/Lmnb1 tm1Yxz, Lmnb2 tm1Yxz/Lmnb2 tm1Yxz single and double germline LOF | Single and double mutants thin diaphragms. Penetrant. Fig. S10 | | | |
| 52 | MNX1 (HB9) | | | | | Arber et al 1999 | HB9nlsLacZ/nlzLacZ mice germline LOF | Thin diaphragm due to innervation issues. Frequency not reported. Fig 7. " | | | |
| 53 | HSPG2 | | | | | | | | | | |
| 54 | MYRF | | | | | | | | | | |
| 55 | PBX1 | | | | | Russell et al., 2012 | Pbx1neo/neo germline LOF | Muscle-less patches in diaphragm. 3/4 homozygous mutants investigated. Fid. 3. | 75% (3/4) | | |

### Table S1

| Ranking | Human Gene Name | Mouse data 2: Evidence | Mouse data 2: Frequency | Mouse data 3: Citation | Mouse data 3: Mouse Genetic Allele | Mouse data 3: Evidence | Mouse data 3: Frequency | Wild-Type E12.5 PPF RNA seq TPM | Muscleless Pax3SpD/SpD E12.5 PPF RNA seq TPM | Human data score | Mouse data score | Total score |
|---|---|---|---|---|---|---|---|---|---|---|---|---|
| 1 | ZFPM2 (FOG2) | | | | | | | 42.64 | 29.10 | 9 | 10 | 19 |
| 2 | GATA4 | 100% of mutant mice (n > 33/33) develop multiple hernias. Size and location of hernias varied: 68% formed in posterior lateral diaphragm , 32% in anterior diaphragm. Fig. 2 | 100% (33/33) | | | | | 175.76 | 114.45 | 9 | 10 | 19 |
| 3 | EFEMP2 (FBLN4) | | | | | | | 128.92 | 77.99 | 6 | 10 | 16 |
| 4 | FREM2 | | | | | | | 60.14832 | 47.80447 | 8 | 8 | 16 |
| 5 | WT1 | 38/38 with CDH. Fig 3 | 100% (38/38) | Carmona et al. 2016 | G2-Gata4CreTg/+; WT1fl1/fl1 Conditional LOF in PPFs and Septum Transversum | 16/36 mutant embryos with CDH. Fig 2. | 16/36 | 90.70 | 114.26 | 6 | 10 | 16 |
| 6 | SLIT3 | | | | | | | 25.09 | 17.61 | 5 | 10 | 15 |
| 7 | KIF7 | | | | | | | 27.30 | 13.93 | 5 | 10 | 15 |
| 8 | HLX | | | | | | | 166.84 | 193.75 | 4 | 10 | 14 |
| 9 | GLI2 | | | | | | | 41.63 | 45.59 | 5 | 8 | 13 |
| 10 | SOX7 | | | | | | | 15.73 | 48.79 | 5 | 8 | 13 |
| 11 | FREM1 | | | | | | | 42.83 | 11.23 | 5 | 8 | 13 |
| 12 | FRAS1 | | | | | | | 91.71493 | 81.57263 | 5 | 8 | 13 |
| 13 | NR2F2 (COUPTFII) | | | | | | | 56.84 | 73.10 | 4 | 9 | 13 |
| 14 | GLI3 | | | | | | | 46.96 | 46.95 | 5 | 8 | 13 |
| 15 | MET (CMET) | Muscleless diaphragm | 100% | | | | | 11.65 | 16.94 | 6 | 7 | 13 |
| 16 | FGFRL1 | Amuscular posterior region in diaphragm that is "fully penetrant". Fig. 3 | 100% | | | | | 80.07 | 37.14 | 6 | 7 | 13 |
| 17 | RARB | | | | | | | 69.79 | 55.39 | 8 | 4 | 12 |
| 18 | FBN1 | | | | | | | 109.30 | 103.21 | 4 | 8 | 12 |
| 19 | PDGFRA | | | | | | | 368.25 | 192.69 | 6 | 6 | 12 |
| 20 | HOXB4 | | | | | | | 20.44 | 12.17 | 4 | 7 | 11 |
| 21 | NEDD4 | | | | | | | 877.34 | 449.80 | 5 | 6 | 11 |
| 22 | CTNNB1 | | | | | | | 919.19 | 444.18 | 0 | 10 | 10 |
| 23 | SIX4 | | | | | | | 20.82 | 6.56 | 5 | 5 | 10 |
| 24 | CTBP2 | | | | | | | 37.58 | 18.01 | 6 | 4 | 10 |
| 25 | PAX7 | | | | | | | 23.02 | 2.18 | 5 | 5 | 10 |
| 26 | TBX5 | | | | | | | 46.80 | 25.23 | 5 | 5 | 10 |
| 27 | CHAT | | | | | | | 0.12 | 44.07 | 0 | 10 | 10 |
| 28 | RARA | | | | | | | 58.31 | 52.44 | 5 | 4 | 9 |
| 29 | ROBO2 | | | | | | | 10.80 | 7.46 | 5 | 4 | 9 |
| 30 | MMP2 | | | | | | | 550.61 | 334.17 | 5 | 4 | 9 |
| 31 | MMP14 | | | | | | | 474.45 | 285.82 | 5 | 4 | 9 |
| 32 | NDST1 | | | | | | | 119.56 | 60.38 | 0 | 9 | 9 |
| 33 | DNASE2A (DNASE2) | | | | | | | 8.61 | 5.90 | 1 | 8 | 9 |
| 34 | STRA6 | | | | | | | 23.23 | 10.10 | 9 | 0 | 9 |
| 35 | LTBP4 | | | | | | | 108.33 | 70.67 | 9 | 0 | 9 |
| 36 | EFNB1 | | | | | | | 109.50 | 232.72 | 9 | 0 | 9 |
| 37 | PORCN | | | | | | | 7.80 | 30.46 | 9 | 0 | 9 |
| 38 | MYH10 | | | | | | | 256.19 | 154.60 | 3 | 5 | 8 |
| 39 | LRP2 | | | | | | | 5.81 | 8.46 | 8 | 0 | 8 |
| 40 | EYA1 | | | | | | | 13.83 | 7.58 | 4 | 4 | 8 |
| 41 | LOX | | | | | | | 185.51 | 121.98 | 0 | 8 | 8 |
| 42 | MYOD1 | | | | | | | 179.00 | 1.24 | 4 | 4 | 8 |
| 43 | ILF3 (N90) | | | | | | | 202.55 | 94.00 | 3 | 5 | 8 |
| 44 | PAX3 | Amuscular diaphragm. Fig. 1-3. 100% | 100% | | | | | 3.77 | 1.12 | 1 | 7 | 8 |
| 45 | GPC3 | | | | | | | 1525.60 | 963.25 | 8 | 0 | 8 |
| 46 | WDR35 | | | | | | | 14.80279 | 6.869294 | 0 | 8 | 8 |
| 47 | TMEM70 | | | | | | | 55.53796192 | 54.26057 | 8 | 0 | 8 |
| 48 | EYA2 | | | | | | | 21.29 | 8.18 | 3 | 4 | 7 |
| 49 | MYOG | Muscleless diaphragm. Frequency not reported. Fig. 3. | | | | | | 648.50 | 203.42 | 0 | 7 | 7 |
| 50 | LMNB1 | | | | | | | 417.1488 | 209.4641 | 0 | 7 | 7 |
| 51 | LMNB2 | | | | | | | 106.0142 | 37.45502 | 0 | 7 | 7 |
| 52 | MNX1 (HB9) | | | | | | | 0 | 0 | 0 | 7 | 7 |
| 53 | HSPG2 | | | | | | | 154.4014 | 76.00331 | 7 | 0 | 7 |
| 54 | MYRF | | | | | | | 156.0260142 | 143.3713261 | 7 | 0 | 7 |
| 55 | PBX1 | | | | | | | 86.08 | 119.06 | 1 | 6 | 7 |

### Table S1

CDH Implicated Genes

| | | | | | | Human Reference 1 | | | | | Human Reference 2 | | | |
|---|---|---|---|---|---|---|---|---|---|---|---|---|---|---|
| **Ranking** | **Human Gene Name** | **Description** | **Human Genome (Hg19) coordinates** | **Mouse Genome (Mm9) coordinates** | **Isolated or Complex CDH (from human data)** | **Citation** | **De Novo Cases** | **Inherited Cases** | **Unknown inheritance** | **Frequency in CDH cohort** | **Citation** | **De Novo Cases** | **Inherited Cases** | **Unknown inheritance** |
| **56** | **CHD7** | chromodomain helicase DNA binding protein 7 | chr8 61591394 61780586 | chr4 8617552 8795596 | Complex CDH, CHARGE Syndrome | Jongmans et al. 2006 | | | 2 CDH/107 CHARGE; 67/107 CHD7 mutations | | Longoni et al., 2014 | | CDH, 1 compound heterozygous, 2 maternal, 2 paternal | 1/275 |
| **57** | **CHRNG** | cholinergic receptor nicotinic gamma subunit | chr2 233404436 233411038 | chr1 89102385 89108410 | | | | | | | | | | |
| **58** | **HCCS** | holocytochrome c synthase | chrX 11129405 11141204 | chrX 165749462 165758304 | Complex CDH, Microphthalmia with linear skin defects | Qidwai et al. 2010 | | 1 (a inherited allele, CNV on other alllele) | | Case Study | | | | |
| 59 | **HADHA** | ketoacyl-CoA thiolase/enoyl-CoA hydratase (trifunctional protein), alpha subunit | chr2 26413503 26467594 | chr5 30445962 30481520 | | | | | | | | | | |
| **60** | **IGF1R** | insulin like growth factor 1 receptor | chr15 99191767 99507759 | chr7 75097142 75378553 | Isolated and Complex CDH | Longoni et al. 2014 | | 2/275 (Isolated CDH, 1 maternal allele, 1 with allele shared with unaffected sibling) | 1/275 (Complex CDH) | 1% (3/275) | | | | |
| **61** | **GATA6** | GATA binding protein 6, transcription factor | chr18 19749397 19782491 | chr18 11052507 11085633 | Complex CDH | Yu et al., 2014 | 1 | 2 siblings with 1 maternal allele (mother mild phenotype and somatic mosaicism) | | 3 Case Studies | | | | |
| **62** | **RUNX1** | runt related transcription factor 1 | chr21 36160097 36260987 | chr16 92601710 92826319 | Isolated and Complex CDH | Longoni et al. 2014 | | 2/275 (1 Complex and 1 Isolated CDH with maternal allele) | | 0.7% (2/275) | | | | |
| **63** | **ZFHX4** | zinc finger homeobox 4, transcription factor | chr8 77593514 77779521 | chr3 5218553 5415855 | Isolated CDH | Longoni et al. 2014 | | 2/275 (2 CDH with maternal allele) | 1/275 (Isolated CDH) | 1% (3/275) | | | | |
| **64** | **PBX3** | pre B cell leukemia homeobox 3, transcription factor | chr9 128509616 128729655 | chr2 34026976 34227565 | Isolated and Complex CDH | Longoni et al. 2014 | | 5/275 (3 Complex CDH patient with maternal allele; 1 Isolated CDH with paternal allele) | 1/275 (Isolated CDH) | 2% (6/275) | | | | |
| **65** | **SIX1** | sine oculis-related homeobox 1, transcription factor | chr14 61111416 61116155 | chr12 74142813 74147699 | | | | | | | | | | |
| **66** | **DES** | desmin (intermediate filament) | chr2 220283098 220291461 | chr1 75356866 75365154 | | | | | | | | | | |
| **67** | **GAB1** | growth factor receptor bound protein 2-associated binding protein 1 | chr4 144257982 144395718 | chr8 83288329 83404418 | | | | | | | | | | |
| **68** | **SIM2** | SIM bHLH transcription factor 2 | chr21 38071420 38122218 | chr16 94085504 94348638 | | | | | | | | | | |
| **69** | **CTBP1** | C-terminal binding protein 1, transcriptional repressor | chr4 1205227 1242908 | chr5 33590371 33617653 | | | | | | | | | | |
| **70** | **COL3A1** | collagen type III alpha 1 chain | chr2 189839098 189877472 | chr1 45368382 45406551 | Complex CDH, Ehlers-Danlos Syndrome | Lin et al., 2006 | | | Complex CDH, Ehlers Danos Syndrome | | | | | |
| **71** | **ADAM19** | a disintegrin and metallopeptidase domain 19 | chr5 156904311 157002831 | chr11 45868003 45960849 | | | | | | | | | | |
| **72** | **DOCK1** | dedicator of cytokinesis 1 | chr10 128593977 129250781 | chr7 141862369 142365330 | | | | | | | | | | |
| **73** | **BARX2** | BarH-like homeobox 2, transcription factor | chr11 129245880 129322174 | chr9 31653628 31720870 | | | | | | | | | | |
| **74** | **MUSK** | muscle associated receptor tyrosine kinase | chr9 113431050 113563278 | chr4 58298833 58387175 | | | | | | | | | | |
| **75** | **RYR1** | ryanodine receptor 1 | chr19 38924339 39078204 | chr7 29788358 29910170 | | | | | | | | | | |
| **76** | **SRF** | serum response factor | chr6 43140127 43149244 | chr17 46683787 46693111 | | | | | | | | | | |
| **77** | **TNNT3** | troponin T3, fast skeletal type | chr11 1940798 1959936 | chr7 149684740 149701914 | | | | | | | | | | |
| **78** | **ECEL1** | endothelin converting enzyme like 1 | chr2 233344536 233352569 | chr1 89044229 89052931 | | | | | | | | | | |
| **79** | **ROBO1** | roundabout guidance receptor 1 | chr3 78646387 79068609 | chr16 72663393 73046345 | | | | | | | | | | |
| **80** | **PTPRD** | protein tyrosine phosphatase, receptor type, D | chr9 8314245 8733946 | chr4 75587140 75990147 | | | | | | | | | | |
| **81** | **PTPRS** | protein tyrosine phosphatase, receptor type, S | chr19 5205518 5340814 | chr17 56551848 56615903 | | | | | | | | | | |
| **82** | **MSC (MYOR)** | musculin, bHLH transcription factor | chr8 72753776 72756731 | chr1 14743428 14746047 | | | | | | | | | | |
| **83** | **STAC3** | SH3 and cysteine rich domain 3 | chr12 57637235 57644976 | chr10 126938741 126945874 | | | | | | | | | | |
| **84** | **COL20A1** | collagen, type XX, alpha 1 | chr20 61924537 61962285 | chr2 180721239 180752245 | Isolated CDH | Longoni et al. 2014 | | | 2/275 (1 Isolated CDH, 1 unknown phenotype) | 1% (2/275) | | | | |
| **85** | **LBR** | Lamin B Receptor | chr1:225,589,204-225,616,627 | chr1:183,745,446-183,772,532 | Isolated CDH | Kammoun et al. 2018 | | | 12/120 Isolated CDH | 10% (12/120) | | | | |
| **86** | **PPARGC1A** | Peroxisome Proliferator-Activated Receptor Gamma Coactivator 1-Alpha | chr4:23,756,664-23,905,712 | chr5:51,845,488-51,945,160 | Isolated CDH | Kammoun et al. 2018 | | | 7/120 Isolated CDH | 6% (7/120) | | | | |
| **87** | **NSD1** | Nuclear Receptor Binding SET Domain Protein 1 | chr5:176,560,026-176,727,216 | chr13:55,311,143-55,419,686 | Isolated CDH | Kammoun et al. 2018 | | | 12/120 Isolated CDH | 10% (12/120) | | | | |
| **88** | **TLN1** | talin 1 | chr9 35697333 35732392 | chr4 43544384 43575455 | Complex CDH | Yu et al. 2015 | 1/39 Complex CDH | 1/39 Complex CDH | 1/39 Complex CDH | 7.7% (3/39) | | | | |
| **89** | **CDKN1C** | cyclin dependent kinase inhibitor 1C | chr11 2904447 2906995 | chr7 150644243 150646955 | | | | | | | | | | |
| **90** | **NMNAT2** | nicotinamide nucleotide adenylyltransferase 2 | chr1 183217371 183387634 | chr1 154802230 154966391 | | | | | | | | | | |
| **91** | **DISP1** | dispatched RND transporter family member 1 | chr1 222988422 223179337 | chr1_random 48173 185338 | Complex CDH | Kantarci et al., 2010 | 1/168 Complex CDH | | | 0.6% (1/168) | | | | |
| **92** | **SLC5A9** | solute carrier family 5, member 9 | chr1 48688356 48714316 | chr4 111547981 111575401 | Complex CDH | Yu et al. 2015 | 1/39 Complex CH | | | 2.6% (1/39) | | | | |
| **93** | **PRKACB** | protein kinase cAMP-activated catalytic subunit beta, serine/threonine protein kinase family | chr1 84629866 84704181 | chr3 146392542 146433202 | Complex CDH | Yu et al. 2015 | 1/39 Complex CH | | | 2.6% (1/39) | | | | |
| **94** | **FGFR2** | fibroblast growth factor receptor 2 | chr10 123237843 123357972 | chr7 137305964 137410322 | Complex CDH, Apert Syndrome | Bulfamante et al., 2011 | 1 Complex CDH, Apert Syndrome | | | Case Study | | | | |
| **95** | **KMT2D** | lysine methyltransferase 2D | chr12 49412757 49449107 | chr15 98662099 98701636 | Complex CDH, Kabuki Syndrome | McVeigh et al. 2015 | 1 Complex CDH, Kabuki Syndrome | | | Case Study | | | | |
| **96** | **BMP4** | bone morphogenetic protein 4 | chr14 54416454 54423609 | chr14 47003199 47010344 | Complex, SHORT Syndrome | Reis et al., 2011 | 1 Complex, SHORT Syndrome | | | | | | | |
| **97** | **DLST** | dihydrolipoamide S-succinyltransferase (E2 component of 2-oxo-glutarate complex) | chr14 75348593 75370450 | chr12 86451782 86475041 | Complex CDH | Yu et al. 2015 | 1/39 Complex CH | | | 2.6% (1/39) | | | | |
| **98** | **SIN3A** | SIN3 transcription regulator family member A | chr15 75661719 75748124 | chr9 56924182 56976175 | Complex CDH | Yu et al. 2015 | 1/39 Complex CH | | | 2.6% (1/39) | | | | |
| **99** | **TBX6** | T-box 6, transcription factor | chr16 30097114 30103205 | chr7 133924996 133929062 | Isolated CDH | Longoni et al. 2014 | 1/275 Isolated CDH | | 1/275 (1 Isolated CDH patient with 2 mutations) | 0.7% (2/275) | | | | |
| **100** | **FZD2** | frizzled class receptor 2 | chr17 42634811 42638630 | chr11 102465744 102469372 | Complex CDH | Wat et al., 2011 | 1/45 Complex CDH | | | 2% (1/45) | | | | |
| **101** | **LONP1** | lon peptidase 1, mitochondrial | chr19 5691844 5720463 | chr17 56753721 56766326 | Complex CDH | Yu et al. 2015 | 1/275 Complex CDH | | | 0.4% (1/275) | | | | |
| **102** | **INHBB** | inhibin subunit beta B | chr2 121103717 121109384 | chr1 121312039 121318825 | Complex CDH | Yu et al. 2015 | 1/39 Complex CH | | | 2.6% (1/39) | | | | |
| **103** | **ARFGEF2** | ADP ribosylation factor guanine nucleotide exchange factor 2 | chr20 47538274 47653230 | chr2 166631080 166723551 | Complex CDH | Yu et al. 2015 | 1/39 Complex CH | | | 2.6% (1/39) | | | | |
| **104** | **NIPBL** | cohesin loading factor | chr5 36876861 37065921 | chr15 8239823 8394463 | Complex CDH, Cornelia de Lange Syndrome | Wilmink et al., 2009 | 1 Complex CDH, Cornelia de Lange Syndrome | | | Case Study | Hosokawa et al. 2010 | | | CDH, Brachmann-de Lange |
| **105** | **CDO1** | cysteine dioxygenase type 1 | chr5 115140429 115152405 | chr18 46872846 46888156 | Complex CDH | Yu et al. 2015 | 1/39 Complex CDH | | | 2.6% (1/39) | | | | |
| **106** | **PTPN12** | protein tyrosine phosphatase, non-receptor type 12 | chr7 77167351 77269388 | chr5 20492462 20541065 | Complex CDH | Yu et al. 2015 | 1/39 Complex CDH | | | 2.6% (1/39) | | | | |
| **107** | **PLPP6 (PPAPDC2)** | phospholipid phosphatase 6 | chr9 4662293 4665272 | chr19 29038409 29041291 | Complex CDH | Yu et al. 2015 | 1/39 Complex CDH | | | 2.6% (1/39) | | | | |
| **108** | **CLCN4** | chloride voltage-gated channel 4 | chrX 10124984 10205699 | chr7 7235019 7251547 | Complex CDH | Yu et al. 2015 | 1/39 Complex CDH | | | 2.6% (1/39) | | | | |
| **109** | **STAG2** | stromal antigen 2, subunit of the cohesin complex | chrX 123094409 123236505 | chrX 39502588 39630363 | Complex CDH | Yu et al. 2015 | 1/39 Complex CDH | | | 2.6% (1/39) | | | | |
| **110** | **LAMA5** | laminin subunit alpha 5 | chr20:60,883,011-60,942,368 | chr2:179,911,078-179,960,564 | Isolated CDH | Qi et al. 2018 | 1/362 Isolated CDH | | | 0.3% (1/362) | | | | |

### Table S1

| | | | Human reference 3 | | | | | Human Reference 4 | | | | | Human Reference 5 | | | | | |
|---|---|---|---|---|---|---|---|---|---|---|---|---|---|---|---|---|---|---|
| Ranking | Human Gene Name | Frequency in CDH Cohort | Citation | De Novo Cases | Inherited Cases | Unknown inheritance | Frequency in CDH Cohort | Citation | De Novo Cases | Inherited Cases | Unknown inheritance | Frequency in CDH Cohort | Citation | De Novo Cases | Inherited Cases | Unknown inheritance | Frequency in CDH Cohort | Citation |
| 56 | CHD7 | 2.5% (7/275) | | | | | | | | | | | | | | | | |
| 57 | CHRNG | | | | | | | | | | | | | | | | | |
| 58 | HCCS | | | | | | | | | | | | | | | | | |
| 59 | HADHA | | | | | | | | | | | | | | | | | |
| 60 | IGF1R | | | | | | | | | | | | | | | | | |
| 61 | GATA6 | | | | | | | | | | | | | | | | | |
| 62 | RUNX1 | | | | | | | | | | | | | | | | | |
| 63 | ZFHX4 | | | | | | | | | | | | | | | | | |
| 64 | PBX3 | | | | | | | | | | | | | | | | | |
| 65 | SIX1 | | | | | | | | | | | | | | | | | |
| 66 | DES | | | | | | | | | | | | | | | | | |
| 67 | GAB1 | | | | | | | | | | | | | | | | | |
| 68 | SIM2 | | | | | | | | | | | | | | | | | |
| 69 | CTBP1 | | | | | | | | | | | | | | | | | |
| 70 | COL3A1 | | | | | | | | | | | | | | | | | |
| 71 | ADAM19 | | | | | | | | | | | | | | | | | |
| 72 | DOCK1 | | | | | | | | | | | | | | | | | |
| 73 | BARX2 | | | | | | | | | | | | | | | | | |
| 74 | MUSK | | | | | | | | | | | | | | | | | |
| 75 | RYR1 | | | | | | | | | | | | | | | | | |
| 76 | SRF | | | | | | | | | | | | | | | | | |
| 77 | TNNT3 | | | | | | | | | | | | | | | | | |
| 78 | ECEL1 | | | | | | | | | | | | | | | | | |
| 79 | ROBO1 | | | | | | | | | | | | | | | | | |
| 80 | PTPRD | | | | | | | | | | | | | | | | | |
| 81 | PTPRS | | | | | | | | | | | | | | | | | |
| 82 | MSC (MYOR) | | | | | | | | | | | | | | | | | |
| 83 | STAC3 | | | | | | | | | | | | | | | | | |
| 84 | COL20A1 | | | | | | | | | | | | | | | | | |
| 85 | LBR | | | | | | | | | | | | | | | | | |
| 86 | PPARGC1A | | | | | | | | | | | | | | | | | |
| 87 | NSD1 | | | | | | | | | | | | | | | | | |
| 88 | TLN1 | | | | | | | | | | | | | | | | | |
| 89 | CDKN1C | | | | | | | | | | | | | | | | | |
| 90 | NMNAT2 | | | | | | | | | | | | | | | | | |
| 91 | DISP1 | | | | | | | | | | | | | | | | | |
| 92 | SLC5A9 | | | | | | | | | | | | | | | | | |
| 93 | PRKACB | | | | | | | | | | | | | | | | | |
| 94 | FGFR2 | | | | | | | | | | | | | | | | | |
| 95 | KMT2D | | | | | | | | | | | | | | | | | |
| 96 | BMP4 | | | | | | | | | | | | | | | | | |
| 97 | DLST | | | | | | | | | | | | | | | | | |
| 98 | SIN3A | | | | | | | | | | | | | | | | | |
| 99 | TBX6 | | | | | | | | | | | | | | | | | |
| 100 | FZD2 | | | | | | | | | | | | | | | | | |
| 101 | LONP1 | | | | | | | | | | | | | | | | | |
| 102 | INHBB | | | | | | | | | | | | | | | | | |
| 103 | ARFGEF2 | | | | | | | | | | | | | | | | | |
| 104 | NIPBL | Case Study | | | | | | | | | | | | | | | | |
| 105 | CDO1 | | | | | | | | | | | | | | | | | |
| 106 | PTPN12 | | | | | | | | | | | | | | | | | |
| 107 | PLPP6 (PPAPDC2) | | | | | | | | | | | | | | | | | |
| 108 | CLCN4 | | | | | | | | | | | | | | | | | |
| 109 | STAG2 | | | | | | | | | | | | | | | | | |
| 110 | LAMA5 | | | | | | | | | | | | | | | | | |

### Table S1

CDH Implicated Genes

| Ranking | Human Gene Name | Human Reference 6: De Novo Cases | Inherited Cases | Unknown inheritance | Frequency in CDH Cohort | Mouse data 1: Citation | Mouse Genetic Allele | Evidence | Frequency | Citation | Mouse Genetic Allele |
|---|---|---|---|---|---|---|---|---|---|---|---|
| 56 | CHD7 | | | | | | | | | | |
| 57 | CHRNG | | | | | Pacifici et al. 2011 | Chrngtm2(Chrne)Vwi/Chrngtm2(Chrne)Vwi germline LOF | Thin diaphragm, no herniation at E17. 6/6. Fig. 6. | 100% | | |
| 58 | HCCS | | | | | | | | | | |
| 59 | HADHA | | | | | Ibdah et al. 2001 | Hadha tm1Jib/Hadha tm1Jib (Mtpa -/-) germline LOF | Degeneration of diaphragm myocytes at P0. 100%. Fig. 3 | 100% | | |
| 60 | IGF1R | | | | | | | | | | |
| 61 | GATA6 | | | | | | | | | | |
| 62 | RUNX1 | | | | | | | | | | |
| 63 | ZFHX4 | | | | | | | | | | |
| 64 | PBX3 | | | | | | | | | | |
| 65 | SIX1 | | | | | Laclef et al., 2003 | Six1LacZ/LacZ germline LOF | Amuscular diaphragm. Frequency not reported. Fig 3. | | Grifone et al., 2005 | Six1tm1Mair/Six1tm1Mair; Six4tm1Mair/Six4tm1Mair double germline LOF |
| 66 | DES | | | | | Li et al., 1996 | DesNeo/Neo germline LOF | Aberrant diaphragm muscle. Frequency not reported. Fig. 1 | | | |
| 67 | GAB1 | | | | | Sachs et al., 2000 | Gab1LacZ/LacZ germline LOF | Amuscular diaphragm. Frequency not reported. Fig 3. | | | |
| 68 | SIM2 | | | | | Goshu et al., 2002 | Sim2tm1Shmb/Sim2tm1Shmb germline LOF | Thin diaphragm. Frequency not reported. Fig. 4-5. | | | |
| 69 | CTBP1 | | | | | Hildebrand et al., 2002 | Ctbp1Neo/Neo;Ctbp2Geo/+ germline LOF | Fewer myotubes in diaphragm. Frequency not reported. Fig 5. | | | |
| 70 | COL3A1 | | | | | | | | | | |
| 71 | ADAM19 | | | | | Kurohara et | Meltrin bdMP/dMP germline LOF | Thin diaphragm. Frequency not reported. Fig. 4. | | | |
| 72 | DOCK1 | | | | | Laurin et al. 2008 | Dock1tm1.1Ysfk/Dock1tm1.1Ysfk germline LOF | Thing diaphragm. Frequency not reported. Fig. S3. | | | |
| 73 | BARX2 | | | | | Meech et al. 2012 | Barx2tm1Rsd/Barx2tm1Rsd germline LOF. | Thin diaphragm. Frequency not reported. Fig. 3. | | | |
| 74 | MUSK | | | | | Chevessier et al 2008 | muskV789M/- germline LOF | Thing diaphragm. Frequency not reported. Data not shown. | | | |
| 75 | RYR1 | | | | | Zvaritch et al 2007 | Ryr1 I4895T/I4895T germline LOF mice have myotube misspatterning and thin diaphragm muscle | Mis-patterned myotubes and thin diaphragms.Frequency not reported. Fig 1 and 6. | | | |
| 76 | SRF | | | | | Li et al 2005 | Myo-Cre TG; Srf flex1/flex1 mice conditional LOF in myocytes | Diaphragm with thinner myofibers. Frequency not reported. Fig. 2. | | | |
| 77 | TNNT3 | | | | | Ju et al 2013 | Tnnt 3tm2a(KOMP)Wtsi/ Tnnt3 tm2a(KOMP)Wtsi germline LOF | Markedly thinner diaphragms. Frequency not reported. Fig. 3 and 5. | | | |
| 78 | ECEL1 | | | | | Hagata et al. 2010 | Ecel 1tm1Hiki/Ecel 1tm1Hiki germline LOF | Mutant diaphragms with decreased NMJs and thinner. Frequency not reported. Fig. 1 and 5. | | | |
| 79 | ROBO1 | | | | | Domyan et al., 2013 | Robo1 b-geo/b-geo;Robo2 lacZ/LacZ germline LOF | Posterior CDH in double mutants. No phenotype reported in single mutants. Frequency not reported. Fig 1. | | Xian et al., 2001 | Robo1Neo/Neo targeted germline LOF mutation present with incompletely pentrant CDH |
| 80 | PTPRD | | | | | Uetani et al. 2006 | Ptprd tm1Yiw/Ptprd tm1Yiw; Ptprs tm1Mtr/Ptprs tm1Mtr germline LOF. | Thin diaphragms in double mutants due to innervation issues. 100% Fig. 3. | 100% in double mutants | | |
| 81 | PTPRS | | | | | Uetani et al. 2006 | Ptprd tm1Yiw/Ptprd tm1Yiw; Ptprs tm1Mtr/Ptprs tm1Mtr germline LOF. | Thin diaphragms in double mutants due to innervation issues. 100% Fig. 3. | 100% in double mutants | | |
| 82 | MSC (MYOR) | | | | | Lu et al., 2002 | MyoR(Msc)Neo/Neo;capsulin(Tcf21)Neo/Neo germline LOF | CDH. Frequency not reported. Fig.1. | Not reported | | |
| 83 | STAC3 | | | | | Reinholt et al. 2013 | Stac3 tm1a(KOMP)Wtsi/Stac3 tm1a(KOMP)Wtsi germline LOF | Diaphragms slightly thinner. Frequency not reported. Fig. 3. | | | |
| 84 | COL20A1 | | | | | | | | | | |
| 85 | LBR | | | | | | | | | | |
| 86 | PPARGC1A | | | | | | | | | | |
| 87 | NSD1 | | | | | | | | | | |
| 88 | TLN1 | | | | | | | | | | |
| 89 | CDKN1C | | | | | Zhang et al., 1997 | Cdkn1ctm1Sje/Cdkn1ctm1Sje germline LOF | Umbilical hernia. Frequency not reported. Fig. 2 | | | |
| 90 | NMNAT2 | | | | | Hicks et al. 2012 | Nmnat2blad/blad germline LOF | Underdeveloped diaphragm. Frequency not reported. Fig. 3. | | | |
| 91 | DISP1 | | | | | | | | | | |
| 92 | SLC5A9 | | | | | | | | | | |
| 93 | PRKACB | | | | | | | | | | |
| 94 | FGFR2 | | | | | | | | | | |
| 95 | KMT2D | | | | | | | | | | |
| 96 | BMP4 | | | | | | | | | | |
| 97 | DLST | | | | | | | | | | |
| 98 | SIN3A | | | | | | | | | | |
| 99 | TBX6 | | | | | | | | | | |
| 100 | FZD2 | | | | | | | | | | |
| 101 | LONP1 | | | | | | | | | | |
| 102 | INHBB | | | | | | | | | | |
| 103 | ARFGEF2 | | | | | | | | | | |
| 104 | NIPBL | | | | | | | | | | |
| 105 | CDO1 | | | | | | | | | | |
| 106 | PTPN12 | | | | | | | | | | |
| 107 | PLPP6 (PPAPDC2) | | | | | | | | | | |
| 108 | CLCN4 | | | | | | | | | | |
| 109 | STAG2 | | | | | | | | | | |
| 110 | LAMA5 | | | | | | | | | | |

### Table S1

| | | Mouse data 2 | | Mouse data 3 | | | | | | | | |
|---|---|---|---|---|---|---|---|---|---|---|---|---|
| Ranking | Human Gene Name | Evidence | Frequency | Citation | Mouse Genetic Allele | Evidence | Frequency | Wild-Type E12.5 PPF RNA seq TPM | Muscleless Pax3SpD/SpD E12.5 PPF RNA seq TPM | Human data score | Mouse data score | Total score |
| 56 | CHD7 | | | | | | | 12.86 | 4.95 | 7 | 0 | 7 |
| 57 | CHRNG | | | | | | | 70.17358 | 22.00626 | 0 | 7 | 7 |
| 58 | HCCS | | | | | | | 17.20 | 29.66 | 6 | 0 | 6 |
| 59 | HADHA | | | | | | | 99.71124 | 87.44172 | 0 | 5 | 5 |
| 60 | IGF1R | | | | | | | 62.53 | 52.06 | 5 | 0 | 5 |
| 61 | GATA6 | | | | | | | 125.28 | 101.77 | 5 | 0 | 5 |
| 62 | RUNX1 | | | | | | | 33.69 | 20.70 | 5 | 0 | 5 |
| 63 | ZFHX4 | | | | | | | 54.86 | 27.18 | 5 | 0 | 5 |
| 64 | PBX3 | | | | | | | 24.45 | 29.27 | 5 | 0 | 5 |
| 65 | SIX1 | Amuscular diaphragm. Frequency not reported. Fig 3. | | | | | | 53.06 | 16.88 | 0 | 5 | 5 |
| 66 | DES | | | | | | | 138.85 | 57.21 | 0 | 5 | 5 |
| 67 | GAB1 | | | | | | | 58.04 | 36.82 | | 5 | 5 |
| 68 | SIM2 | | | | | | | 13.42345 | 15.23161 | 0 | 5 | 5 |
| 69 | CTBP1 | | | | | | | 314.26 | 174.06 | 0 | 5 | 5 |
| 70 | COL3A1 | | | | | | | 2520.70 | 2558.60 | 5 | 0 | 5 |
| 71 | ADAM19 | | | | | | | 120.8468 | 55.06543 | 0 | 5 | 5 |
| 72 | DOCK1 | | | | | | | 92.41859 | 46.94166 | 0 | 5 | 5 |
| 73 | BARX2 | | | | | | | 26.6715 | 12.36687 | 0 | 5 | 5 |
| 74 | MUSK | | | | | | | 11.15899 | 0.1380624 | 0 | 5 | 5 |
| 75 | RYR1 | | | | | | | 40.12508 | 1.367034 | 0 | 5 | 5 |
| 76 | SRF | | | | | | | 71.95743 | 91.07074 | 0 | 5 | 5 |
| 77 | TNNT3 | | | | | | | 74.82668 | 3.342672 | 0 | 5 | 5 |
| 78 | ECEL1 | | | | | | | 5.74572 | 8.709288 | 0 | 5 | 5 |
| 79 | ROBO1 | 3/? Mice reported to have hernias, data not shown | 3/? | | | | | 128.90 | 97.21 | 0 | 4 | 4 |
| 80 | PTPRD | | | | | | | 63.25875 | 74.12381 | 0 | 4 | 4 |
| 81 | PTPRS | | | | | | | 289.2762 | 165.5851 | 0 | 4 | 4 |
| 82 | MSC (MYOR) | | | | | | | 102.45 | 25.85 | 0 | 4 | 4 |
| 83 | STAC3 | | | | | | | 33.10312 | 14.4892 | 0 | 4 | 4 |
| 84 | COL20A1 | | | | | | | 20.66 | 7.28 | 4 | 0 | 4 |
| 85 | LBR | | | | | | | 77.48212 | 97.98092 | 4 | 0 | 4 |
| 86 | PPARGC1A | | | | | | | 1.942315 | 9.737457 | 4 | 0 | 4 |
| 87 | NSD1 | | | | | | | 99.37538 | 50.39356 | 4 | 0 | 4 |
| 88 | TLN1 | | | | | | | 146.94 | 77.63 | 4 | 0 | 4 |
| 89 | CDKN1C | | | | | | | 2334.16 | 952.79 | 0 | 3 | 3 |
| 90 | NMNAT2 | | | | | | | 1.750923 | 1.377123 | 0 | 3 | 3 |
| 91 | DISP1 | | | | | | | 42.35 | 41.81 | 3 | 0 | 3 |
| 92 | SLC5A9 | | | | | | | 0.04 | 0.01 | 3 | 0 | 3 |
| 93 | PRKACB | | | | | | | 76.06 | 105.34 | 3 | 0 | 3 |
| 94 | FGFR2 | | | | | | | 21.37 | 16.57 | 3 | 0 | 3 |
| 95 | KMT2D | | | | | | | 34.78 | 32.46 | 3 | 0 | 3 |
| 96 | BMP4 | | | | | | | 57.31 | 37.44 | 3 | 0 | 3 |
| 97 | DLST | | | | | | | 219.75 | 1270.56 | 3 | 0 | 3 |
| 98 | SIN3A | | | | | | | 85.51 | 41.84 | 3 | 0 | 3 |
| 99 | TBX6 | | | | | | | 1.61 | 0.81 | 3 | 0 | 3 |
| 100 | FZD2 | | | | | | | 164.7291 | 180.6991 | 3 | 0 | 3 |
| 101 | LONP1 | | | | | | | 120.95 | 155.94 | 3 | 0 | 3 |
| 102 | INHBB | | | | | | | 0.95 | 0.93 | 3 | 0 | 3 |
| 103 | ARFGEF2 | | | | | | | 19.65 | 16.80 | 3 | 0 | 3 |
| 104 | NIPBL | | | | | | | 64.87 | 34.48 | 3 | 0 | 3 |
| 105 | CDO1 | | | | | | | 48.33 | 664.10 | 3 | 0 | 3 |
| 106 | PTPN12 | | | | | | | 66.58 | 32.95 | 3 | 0 | 3 |
| 107 | PLPP6 (PPAPDC2) | | | | | | | 8.26 | 20.64 | 3 | 0 | 3 |
| 108 | CLCN4 | | | | | | | 53.04 | 35.01 | 3 | 0 | 3 |
| 109 | STAG2 | | | | | | | 67.46 | 33.79 | 3 | 0 | 3 |
| 110 | LAMA5 | | | | | | | 35.964582 | 43.33068 | 3 | 0 | 3 |

### Table S1

CDH Implicated Genes

| | | | | | | Human Reference 1 | | | | | Human Reference 2 | | | |
|---|---|---|---|---|---|---|---|---|---|---|---|---|---|---|
| Ranking | Human Gene Name | Description | Human Genome (Hg19) coordinates | Mouse Genome (Mm9) coordinates | Isolated or Complex CDH (from human data) | Citation | De Novo Cases | Inherited Cases | Unknown inheritance | Frequency in CDH cohort | Citation | De Novo Cases | Inherited Cases | Unknown inheritance |
| 111 | **MEIS2** | Meis homeobox 2, transcription factor | chr15:37,181,406-37,393,504 | chr2:115,687,000-115,890,794 | Isolated CDH | Qi et al. 2018 | 1/362 Isolated CDH | | | 0.3% (1/362) | | | | |
| 112 | **ARID1B** | AT-rich interaction domain 1B | chr6:157,099,063-157,531,913 | chr17:4,995,074-5,347,656 | Complex CDH | Qi et al. 2018 | 1/362 Complex CDH | | | 0.3% (1/362) | | | | |
| 113 | **NAA15** | N(Alpha)-Acetyltransferase 15, NatA Auxiliary Subunit | chr4:140,222,609-140,341,187 | chr3:51,219,938-51,279,907 | Isolated CDH | Qi et al. 2018 | 1/362 Isolated CDH | | | 0.3% (1/362) | | | | |
| 114 | **PTPN11** | Protein Tyrosine Phosphatase, Non-Receptor Type 11 | chr12:112,856,155-112,947,717 | chr5:121,580,542-121,641,406 | Complex CDH | Qi et al. 2018 | 1/362 Complex CDH | | | 0.3% (1/362) | | | | |
| 115 | **BRAF** | B-Raf proto-oncogene, serine/threonine kinase | chr7:140,419,127-140,624,564 | chr6:39,553,237-39,675,462 | Complex CDH | Qi et al. 2018 | 1/362 Complex CDH | | | 0.3% (1/362) | | | | |
| 116 | **KDM5B** | lysine demethylase 5B | chr1:202,696,526-202,778,598 | chr1:136,456,755-136,529,455 | Isolated CDH | Qi et al. 2018 | 1/362 Isolated CDH | | | 0.3% (1/362) | | | | |
| 117 | **POGZ** | pogo transposable element with ZNF domain | chr1:151,375,200-151,431,941 | chr3:94,641,489-94,687,489 | Isolated CDH | Qi et al. 2018 | 1/362 Isolated CDH | | | 0.3% (1/362) | | | | |
| 118 | **RAF1** | Raf-1 proto-oncogene, serine/threonine kinase | chr3:12,625,100-12,705,725 | chr6:115,568,591-115,626,653 | Complex CDH patient | Qi et al. 2018 | 1/362 Complex CDH | | | 0.3% (1/362) | | | | |
| 119 | **ATAD3A** | ATPase family, AAA domain containing 3A | chr1:1,447,523-1,470,067 | chr4:155,114,749-155,135,207 | Isolated CDH | Qi et al. 2018 | 1/362 Isolated CDH | | | 0.3% (1/362) | | | | |
| 120 | **CIC** | Capicua Transcriptional Repressor | chr19:42,772,689-42,799,949 | chr7:26,052,718-26,079,167 | Isolated CDH | Qi et al. 2018 | 1/362 Isolated CDH | | | 0.3% (1/362) | | | | |
| 121 | **EMX2** | empty spiracles homeobox 2, transcription factor | chr10:119,301,955-119,309,057 | chr19:59,533,180-59,539,847 | Isolated CDH | Qi et al. 2018 | 1/362 Isolated CDH | | | 0.3% (1/362) | | | | |
| 122 | **FOXP1** | forkhead box P1, transcription factor | chr3:71,003,844-71,633,140 | chr6:98,875,336-99,385,339 | Isolated CDH | Qi et al. 2018 | 1/362 Isolated CDH | | | 0.3% (1/362) | | | | |
| 123 | **MYT1L** | myelin transcription factor 1 like | chr2:1,792,885-2,335,045 | chr12:30,213,249-30,608,074 | Isolated CDH | Qi et al. 2018 | 1/362 Isolated CDH | | | 0.3% (1/362) | | | | |
| 124 | **EPB41L1** | erythrocyte membrane protein band 4.1 like 1 | chr20:34,679,426-34,820,721 | Chr2:156420909-156543214 | Complex CDH | Qi et al. 2018 | 1/362 Complex CDH | | | 0.3% (1/362) | | | | |
| 125 | **HSD17B10** | hydroxysteroid 17-beta dehydrogenase 10 | chrX:53,458,206-53,461,323 | chrX:148,436,439-148,438,985 | Isolated CDH | Qi et al. 2018 | 1/362 Isolated CDH | | | 0.3% (1/362) | | | | |
| 126 | **NACC1** | nucleus accumbens associated 1 | chr19:13,228,917-13,251,959 | chr8:87,194,378-87,211,761 | Complex CDH | Qi et al. 2018 | 1/362 Isolated CDH | | | 0.3% (1/362) | | | | |
| 127 | **SRGAP3** | SLIT-ROBO Rho GTPase activating protein 3 | chr3:9,022,275-9,404,737 | chr6:112,667,966-112,897,260 | Isolated CDH | Qi et al. 2018 | 1/362 Isolated CDH | | | 0.3% (1/362) | | | | |
| 128 | **ARRDC4** | arrestin domain containing 4 | chr15 98503932 98517068 | chr7 75881879 75894124 | Complex CDH | Longoni et al. 2014 | | 1/275 (Complex CDH with 1 paternal allele) | 1/275 (Complex CDH) | 0.7% (2/275) | | | | |
| 129 | **TGIF1** | TGFB induced factor homeobox, transcription factor | chr18 3449410 3458406 | chr17 71193544 71202872 | Human CDH patient | Longoni et al. 2014 | | 1/275 (Complex CDH with 1 paternal allele) | 1/275 (Isolated CDH) | 0.7% (2/275) | | | | |
| 130 | **FOXF2** | forkhead box F2, transcription factor | chr6 1389810 1395838 | chr13 31717684 31723275 | Isolated CDH | Longoni et al. 2014 | | 1/275 (Isolated CDH with 1 paternal allele) | 1/275 (Isolated CDH) | 0.7% (2/275) | | | | |
| 131 | **NEIL2** | nei like DNA glycosylase 2 | chr8 11627133 11644864 | chr14 63801281 63812362 | Isolated CDH | Longoni et al 2012 | | 1/226 (Isolated CDH with 1 maternal allele) | 1/226 (Isolated CDH) | 0.9% (2/226) | | | | |
| 132 | **MEF2A** | myocyte enhancer factor 2A, transcription factor | chr15 100106132 100256693 | chr7 74376048 74517744 | Isolated CDH | Longoni et al. 2014 | | 1/275 (Isolated CDH with 1 paternal allele) | | 0.4% (1/275) | | | | |
| 133 | **TWIST1** | twist family bHLH transcription factor 1 | chr7 19155090 19157295 | chr12 34642535 34644696 | Complex CDH, Coronal craniosynostosis and radial ray hypoplasia | Piard et al., 2012 | | 1/1 (Complex CDH with 1 paternal allele) | | Case Study | | | | |
| 134 | **ELN** | elastin | chr7 73442118 73484236 | chr5 135178464 135223193 | Complex CDH, cutis laxa | Graul-Neumann et al., 2008 | | 1/1 (Complex CDH with 1 paternal allele) | | Case Study | | | | |
| 135 | **TBX1** | T-Box 1, transcription factor | chr22:19,744,226-19,771,116 | chr16:18,581,806-18,587,062 | Isolated CDH | Kammoun et al. 2018 | | 1/120 (Isolated CDH with 1 maternal allele) | | 0.8% (1/120) | | | | |
| 136 | **RC3H1** | RING CCCH (C3H) domains 1 | chr1 173900221 173962210 | chr1 162836541 162905107 | Human CDH patient | Longoni et al. 2014 | | | 1/275 (Complex CDH) | 0.4% (1/275) | | | | |
| 137 | **ZEB1** | zinc finger E-box binding homeobox 1, transcription factor | chr10 31610063 31818742 | chr18 5592644 5775466 | Human CDH patient | Longoni et al. 2014 | | | 1/275 (Isolated CDH) | 0.4% (1/275) | | | | |
| 138 | **TSC2** | TSC complex subunit 2 | chr16 2097895 2138721 | chr17 24732760 24769572 | Complex CDH, Tuberous sclerosis complex | Niemi et al., 2011 | | | Tuberous sclerosis complex | Case Study | | | | |
| 139 | **ELAC2 (ELAC)** | elaC ribonuclease Z 2 | chr17 12894928 12921381 | chr11 64792536 64815576 | Complex CDH | Longoni et al. 2014 | | | 1/275 (Complex CDH) | 0.4% (1/275) | | | | |
| 140 | **DSEI** | dermatan sulfate epimerase like | chr18 65173818 65183967 | chr1 113755278 113761495 | Isolated CDH | Zayed et al., 2010 | | | 1/125 | 0.8% (1/125) | | | | |
| 141 | **DLL3** | delta like canonical Notch ligand 3 | chr19 39989556 39999121 | chr7 29078573 29086804 | Complex, spondylocostal dysostosis | Bulman et al., 2000 | | | 1 (Complex CDH) | | | | | |
| 142 | **STK36** | serine/threonine kinase 36 | chr2 219536748 219567440 | chr1 74648028 74683467 | Complex CDH | Longoni et al. 2014 | | | 1/275 (Complex CDH) | 0.4% (1/275) | | | | |
| 143 | **SMARCC1** | SWI/SNF related, matrix associated, actin dependent regulator of chromatin subfamily c member 1 | chr3 47627377 47823405 | chr9 110034527 110142682 | Isolated CDH | Longoni et al. 2014 | | | 1/275 (Isolated CDH) | 0.4% (1/275) | | | | |
| 144 | **GPR125 (ADGRA3)** | G protein-coupled receptor 125 | chr4 22388996 22517677 | chr5 50351189 50450235 | Complex CDH | Longoni et al. 2014 | | | 1/275 (Complex CDH) | 0.4% (1/275) | | | | |
| 145 | **PTPN13** | protein tyrosine phosphatase, non-receptor type 13 | chr4 87515467 87736328 | chr5 103854210 104027380 | Complex CDH | Longoni et al. 2014 | | | 1/275 (Complex CDH) | 0.4% (1/275) | | | | |
| 146 | **FYB (FYB1)** | FYN binding protein | chr5 39105353 39270759 | chr15 6529846 6615608 | Isolated CDH | Longoni et al. 2014 | | | 1/275 (Isolated CDH) | 0.4% (1/275) | | | | |
| 147 | **FOXC1** | forkhead box C1, transcription factor | chr6 1610680 1614129 | chr13 31898514 31902504 | Complex CDH, Tetralogy of Fallot | Longoni et al. 2014 | | | 1/275 (Complex CDH) | 0.4% (1/275) | | | | |
| 148 | **SCUBE3** | signal peptide, CUB domain and EGF like domain containing 3 | chr6 35181838 35220856 | chr17 28279470 28308290 | Isolated CDH | Longoni et al. 2014 | | | 1/275 (Isolated CDH) | 0.4% (1/275) | | | | |
| 149 | **OCRL** | inositol polyphosphate-5-phosphatase | chrX 128674251 128726530 | chrX 45265632 45319043 | Complex CDH, Lowe syndrome | Race et al., 2010 | | | 1 (Complex CDH) | Case Study | | | | |
| 150 | **HYLS1** | centriolar and ciliogenesis associated | chr11 125753508 125770541 | chr9 35368405 35377654 | Complex CDH, Hydrolethalus syndrome | Salonen et al., 1990 | | | 1 (Complex CDH) | Case Study | | | | |
| 151 | **MID1** | midline 1 | chrX 10413349 10588674 | chrX 166266090 166428729 | Complex CDH | Taylor and Aftimos 2010 | | | 1 (Complex CDH) | Case Study | | | | |
| 152 | **HIRA** | histone cell cycle regulator | chr22 19318223 19419219 | chr16 18876842 18970401 | Complex CDH | Gupta et al 2016 | | | 1 (Complex CDH) | Case Study | | | | |
| 153 | **GATA5** | GATA binding protein 5, transcription factor | chr20:61,038,553-61,051,026 | chr2:180,059,793-180,069,384 | Isolated CDH | Kammoun et al. 2018 | | | 1/120 (Isolated CDH) | 0.8% (1/120) | | | | |

### Table S1

CDH Implicated Genes

| | | | Human reference 3 | | | | | Human Reference 4 | | | | | Human Reference 5 | | | | | |
|---|---|---|---|---|---|---|---|---|---|---|---|---|---|---|---|---|---|---|
| Ranking | Human Gene Name | Frequency in CDH Cohort | Citation | De Novo Cases | Inherited Cases | Unknown inheritance | Frequency in CDH Cohort | Citation | De Novo Cases | Inherited Cases | Unknown inheritance | Frequency in CDH Cohort | Citation | De Novo Cases | Inherited Cases | Unknown inheritance | Frequency in CDH Cohort | Citation |
| 111 | MEIS2 | | | | | | | | | | | | | | | | | |
| 112 | ARID1B | | | | | | | | | | | | | | | | | |
| 113 | NAA15 | | | | | | | | | | | | | | | | | |
| 114 | PTPN11 | | | | | | | | | | | | | | | | | |
| 115 | BRAF | | | | | | | | | | | | | | | | | |
| 116 | KDM5B | | | | | | | | | | | | | | | | | |
| 117 | POGZ | | | | | | | | | | | | | | | | | |
| 118 | RAF1 | | | | | | | | | | | | | | | | | |
| 119 | ATAD3A | | | | | | | | | | | | | | | | | |
| 120 | CIC | | | | | | | | | | | | | | | | | |
| 121 | EMX2 | | | | | | | | | | | | | | | | | |
| 122 | FOXP1 | | | | | | | | | | | | | | | | | |
| 123 | MYT1L | | | | | | | | | | | | | | | | | |
| 124 | EPB41L1 | | | | | | | | | | | | | | | | | |
| 125 | HSD17B10 | | | | | | | | | | | | | | | | | |
| 126 | NACC1 | | | | | | | | | | | | | | | | | |
| 127 | SRGAP3 | | | | | | | | | | | | | | | | | |
| 128 | ARRDC4 | | | | | | | | | | | | | | | | | |
| 129 | TGIF1 | | | | | | | | | | | | | | | | | |
| 130 | FOXF2 | | | | | | | | | | | | | | | | | |
| 131 | NEIL2 | | | | | | | | | | | | | | | | | |
| 132 | MEF2A | | | | | | | | | | | | | | | | | |
| 133 | TWIST1 | | | | | | | | | | | | | | | | | |
| 134 | ELN | | | | | | | | | | | | | | | | | |
| 135 | TBX1 | | | | | | | | | | | | | | | | | |
| 136 | RC3H1 | | | | | | | | | | | | | | | | | |
| 137 | ZEB1 | | | | | | | | | | | | | | | | | |
| 138 | TSC2 | | | | | | | | | | | | | | | | | |
| 139 | ELAC2 (ELAC) | | | | | | | | | | | | | | | | | |
| 140 | DSEI | | | | | | | | | | | | | | | | | |
| 141 | DLL3 | | | | | | | | | | | | | | | | | |
| 142 | STK36 | | | | | | | | | | | | | | | | | |
| 143 | SMARCC1 | | | | | | | | | | | | | | | | | |
| 144 | GPR125 (ADGRA3) | | | | | | | | | | | | | | | | | |
| 145 | PTPN13 | | | | | | | | | | | | | | | | | |
| 146 | FYB (FYB1) | | | | | | | | | | | | | | | | | |
| 147 | FOXC1 | | | | | | | | | | | | | | | | | |
| 148 | SCUBE3 | | | | | | | | | | | | | | | | | |
| 149 | OCRL | | | | | | | | | | | | | | | | | |
| 150 | HYLS1 | | | | | | | | | | | | | | | | | |
| 151 | MID1 | | | | | | | | | | | | | | | | | |
| 152 | HIRA | | | | | | | | | | | | | | | | | |
| 153 | GATA5 | | | | | | | | | | | | | | | | | |

### Table S1



| | | Human Reference 6 | | | | | Mouse data 1 | | | | |
|---|---|---|---|---|---|---|---|---|---|---|---|
| Ranking | Human Gene Name | De Novo Cases | Inherited Cases | Unknown inheritance | Frequency in CDH Cohort | Citation | Mouse Genetic Allele | Evidence | Frequency | Citation | Mouse Genetic Allele |
| 111 | MEIS2 | | | | | | | | | | |
| 112 | ARID1B | | | | | | | | | | |
| 113 | NAA15 | | | | | | | | | | |
| 114 | PTPN11 | | | | | | | | | | |
| 115 | BRAF | | | | | | | | | | |
| 116 | KDM5B | | | | | | | | | | |
| 117 | POGZ | | | | | | | | | | |
| 118 | RAF1 | | | | | | | | | | |
| 119 | ATAD3A | | | | | | | | | | |
| 120 | CIC | | | | | | | | | | |
| 121 | EMX2 | | | | | | | | | | |
| 122 | FOXP1 | | | | | | | | | | |
| 123 | MYT1L | | | | | | | | | | |
| 124 | EPB41L1 | | | | | | | | | | |
| 125 | HSD17B10 | | | | | | | | | | |
| 126 | NACC1 | | | | | | | | | | |
| 127 | SRGAP3 | | | | | | | | | | |
| 128 | ARRDC4 | | | | | | | | | | |
| 129 | TGIF1 | | | | | | | | | | |
| 130 | FOXF2 | | | | | | | | | | |
| 131 | NEIL2 | | | | | | | | | | |
| 132 | MEF2A | | | | | | | | | | |
| 133 | TWIST1 | | | | | | | | | | |
| 134 | ELN | | | | | | | | | | |
| 135 | TBX1 | | | | | | | | | | |
| 136 | RC3H1 | | | | | | | | | | |
| 137 | ZEB1 | | | | | | | | | | |
| 138 | TSC2 | | | | | | | | | | |
| 139 | ELAC2 (ELAC) | | | | | | | | | | |
| 140 | DSEI | | | | | | | | | | |
| 141 | DLL3 | | | | | | | | | | |
| 142 | STK36 | | | | | | | | | | |
| 143 | SMARCC1 | | | | | | | | | | |
| 144 | GPR125 (ADGRA3) | | | | | | | | | | |
| 145 | PTPN13 | | | | | | | | | | |
| 146 | FYB (FYB1) | | | | | | | | | | |
| 147 | FOXC1 | | | | | | | | | | |
| 148 | SCUBE3 | | | | | | | | | | |
| 149 | OCRL | | | | | | | | | | |
| 150 | HYLS1 | | | | | | | | | | |
| 151 | MID1 | | | | | | | | | | |
| 152 | HIRA | | | | | | | | | | |
| 153 | GATA5 | | | | | | | | | | |

### Table S1

| Ranking | Human Gene Name | Mouse data 2: Evidence | Frequency | Mouse data 3: Citation | Mouse Genetic Allele | Evidence | Frequency | Wild-Type E12.5 PPF RNA seq TPM | Muscleless Pax3SpD/SpD E12.5 PPF RNA seq TPM | Human data score | Mouse data score | Total score |
|---|---|---|---|---|---|---|---|---|---|---|---|---|
| 111 | MEIS2 | | | | | | | 69.82003 | 82.11509 | 3 | 0 | 3 |
| 112 | ARID1B | | | | | | | 33.76292 | 36.81079 | 3 | 0 | 3 |
| 113 | NAA15 | | | | | | | 82.55439 | 77.45732 | 3 | 0 | 3 |
| 114 | PTPN11 | | | | | | | 111.2246 | 105.8143 | 3 | 0 | 3 |
| 115 | BRAF | | | | | | | 14.69511 | 15.69106 | 3 | 0 | 3 |
| 116 | KDM5B | | | | | | | 115.5003 | 115.7965 | 3 | 0 | 3 |
| 117 | POGZ | | | | | | | 46.65302 | 47.51435 | 3 | 0 | 3 |
| 118 | RAF1 | | | | | | | 97.2209382 | 98.4986101 | 3 | 0 | 3 |
| 119 | ATAD3A | | | | | | | 61.27298 | 52.13447 | 3 | 0 | 3 |
| 120 | CIC | | | | | | | 91.17046 | 97.6336 | 3 | 0 | 3 |
| 121 | EMX2 | | | | | | | 14.269812 | 18.985659 | 3 | 0 | 3 |
| 122 | FOXP1 | | | | | | | 46.39696 | 60.03954 | 3 | 0 | 3 |
| 123 | MYT1L | | | | | | | 0.07966889 | 0.1212797 | 3 | 0 | 3 |
| 124 | EPB41L1 | | | | | | | 22.614645 | 23.901433 | 3 | 0 | 3 |
| 125 | HSD17B10 | | | | | | | 175.3691 | 166.9034 | 3 | 0 | 3 |
| 126 | NACC1 | | | | | | | 92.35968 | 101.6267 | 3 | 0 | 3 |
| 127 | SRGAP3 | | | | | | | 38.1786 | 34.93193 | 3 | 0 | 3 |
| 128 | ARRDC4 | | | | | | | 19.11 | 30.77 | 3 | 0 | 3 |
| 129 | TGIF1 | | | | | | | 54.10 | 56.27 | 3 | 0 | 3 |
| 130 | FOXF2 | | | | | | | 1.10 | 0.68 | 3 | 0 | 3 |
| 131 | NEIL2 | | | | | | | 8.07 | 1.86 | 3 | 0 | 3 |
| 132 | MEF2A | | | | | | | 55.99 | 22.24 | 2 | 0 | 2 |
| 133 | TWIST1 | | | | | | | 197.02 | 205.11 | 2 | 0 | 2 |
| 134 | ELN | | | | | | | 120.33 | 64.35 | 2 | 0 | 2 |
| 135 | TBX1 | | | | | | | 13.1794823 | 2.29196216 | 2 | 0 | 2 |
| 136 | RC3H1 | | | | | | | 21.36 | 23.33 | 1 | 0 | 1 |
| 137 | ZEB1 | | | | | | | 55.39 | 18.63 | 1 | 0 | 1 |
| 138 | TSC2 | | | | | | | 81.29 | 44.44 | 1 | 0 | 1 |
| 139 | ELAC2 (ELAC) | | | | | | | 11.36 | 5.46 | 1 | 0 | 1 |
| 140 | DSEI | | | | | | | 23.04 | 24.23 | 1 | 0 | 1 |
| 141 | DLL3 | | | | | | | 0.54 | 0.32 | 1 | 0 | 1 |
| 142 | STK36 | | | | | | | 8.86 | 8.98 | 1 | 0 | 1 |
| 143 | SMARCC1 | | | | | | | 203.15 | 97.36 | 1 | 0 | 1 |
| 144 | GPR125 (ADGRA3) | | | | | | | 165.88 | 67.71 | 1 | 0 | 1 |
| 145 | PTPN13 | | | | | | | 76.29 | 46.36 | 1 | 0 | 1 |
| 146 | FYB (FYB1) | | | | | | | 4.02 | 3.54 | 1 | 0 | 1 |
| 147 | FOXC1 | | | | | | | 47.11 | 25.70 | 1 | 0 | 1 |
| 148 | SCUBE3 | | | | | | | 60.76 | 47.67 | 1 | 0 | 1 |
| 149 | OCRL | | | | | | | 19.14 | 10.50 | 1 | 0 | 1 |
| 150 | HYLS1 | | | | | | | 23.09 | 16.92 | 1 | 0 | 1 |
| 151 | MID1 | | | | | | | 13.60897 | 8.930501 | 1 | 0 | 1 |
| 152 | HIRA | | | | | | | 61.13702 | 34.94552 | 1 | 0 | 1 |
| 153 | GATA5 | | | | | | | 19.5886 | 34.5876 | 1 | 0 | 1 |

#### TABLE S2

Heterozygosity and Ancestry

| Sample | Read Depth lower than expected | Count of Heterozygous sites sampled | Ratio of heterozygous/ homozygous calls | Mean depth of sites sampled | Median depth of sites sampled | Total number of sites sampled | Percentage of callable sites | Ancestry prediction | Ancestry probability | Self-reported ancestry |
|---|---|---|---|---|---|---|---|---|---|---|
| Father 411 | FALSE | 7726 | 0.3542 | 49.55 | 48 | 21780 | 0.9997 | EUR | 0.9068 | White, non-hispanic |
| Mother 411 | FALSE | 7681 | 0.3522 | 47.75 | 46 | 21791 | 0.9996 | EUR | 0.9833 | White, non-hispanic |
| Proband Blood 411 | FALSE | 7632 | 0.3499 | 54.13 | 53 | 21782 | 0.9995 | EUR | 0.9792 | White, non-hispanic |
| Proband Skin 411 | FALSE | 7626 | 0.3497 | 51.19 | 50 | 21781 | 0.9996 | EUR | 0.979 | White, non-hispanic |
| Proband Diaphragm 411 | FALSE | 7632 | 0.3499 | 62.99 | 62 | 21784 | 0.9996 | EUR | 0.9792 | White, non-hispanic |
| Father 716 | FALSE | 7278 | 0.3337 | 61.21 | 60 | 21794 | 0.9996 | UNKNOWN | 0.4144 | Asian |
| Mother 716 | FALSE | 7344 | 0.3367 | 62.5 | 61 | 21798 | 0.9996 | UNKNOWN | 0.4933 | Asian |
| Proband Blood 716 | FALSE | 7310 | 0.3352 | 57.6 | 56 | 21790 | 0.9994 | UNKNOWN | 0.4762 | Asian |
| Proband Skin 716 | FALSE | 7318 | 0.3355 | 64.43 | 63 | 21794 | 0.9995 | UNKNOWN | 0.4777 | Asian |
| Proband Diaphragm 716 | FALSE | 7305 | 0.3349 | 60.57 | 59 | 21782 | 0.9994 | UNKNOWN | 0.4748 | Asian |
| Father 809 | FALSE | 7885 | 0.3615 | 47.83 | 46 | 21789 | 0.9996 | EUR | 0.9943 | White, non-hispanic |
| Mother 809 | FALSE | 7299 | 0.3347 | 65.44 | 64 | 21800 | 0.9995 | EUR | 0.9857 | White, non-hispanic |
| Proband Blood 809 | FALSE | 7647 | 0.3506 | 61.26 | 60 | 21804 | 0.9995 | EUR | 0.8451 | White, non-hispanic |
| Proband Skin 809 | FALSE | 7646 | 0.3506 | 60.34 | 59 | 21799 | 0.9994 | EUR | 0.8335 | White, non-hispanic |
| Proband Diaphragm 809 | FALSE | 7645 | 0.3505 | 64.86 | 64 | 21803 | 0.9993 | EUR | 0.8355 | White, non-hispanic |
| Father 967 | FALSE | 7820 | 0.3586 | 55.13 | 54 | 21797 | 0.9996 | AFR | 0.9739 | black and hispanic white |
| Mother 967 | FALSE | 7361 | 0.3375 | 58 | 57 | 21801 | 0.9996 | AFR | 0.9956 | black non-hispanic |
| Proband Blood 967 | FALSE | 7439 | 0.3411 | 46.33 | 45 | 21787 | 0.9997 | AFR | 0.9862 | |
| Proband Skin 967 | FALSE | 7440 | 0.3411 | 59.93 | 59 | 21803 | 0.9998 | AFR | 0.9852 | |
| Proband Diaphragm 967 | FALSE | 7428 | 0.3406 | 40.19 | 40 | 21776 | 0.9995 | AFR | 0.9852 | |

**TABLE S2**

| Sample | Incorrect sex? | Count of heterozygous sites sampled | Ratio of heterozygous/ homozygous calls | Count of homozygous alternative sites | Count of homozygous reference sites | Sex of sample given | Predicted Sex |
|---|---|---|---|---|---|---|---|
| Father 411 | FALSE | 926 | 0.03606 | 25677 | 72541 | male | male |
| Mother 411 | FALSE | 25214 | 1.806 | 13959 | 60383 | female | female |
| Proband Blood 411 | FALSE | 1311 | 0.05019 | 26120 | 71644 | male | male |
| Proband Skin 411 | FALSE | 1250 | 0.04792 | 26085 | 71679 | male | male |
| Proband Diaphragm 411 | FALSE | 1538 | 0.05879 | 26162 | 71487 | male | male |
| Father 716 | FALSE | 1015 | 0.04011 | 25307 | 72917 | male | male |
| Mother 716 | FALSE | 22510 | 1.367 | 16468 | 60741 | female | female |
| Proband Blood 716 | FALSE | 23893 | 1.579 | 15127 | 60667 | female | female |
| Proband Skin 716 | FALSE | 24017 | 1.587 | 15134 | 60534 | female | female |
| Proband Diaphragm 716 | FALSE | 24005 | 1.593 | 15067 | 60497 | female | female |
| Father 809 | FALSE | 864 | 0.03322 | 26012 | 72094 | male | male |
| Mother 809 | FALSE | 27672 | 2.229 | 12414 | 59666 | female | female |
| Proband Blood 809 | FALSE | 27156 | 2.109 | 12877 | 59679 | female | female |
| Proband Skin 809 | FALSE | 27194 | 2.114 | 12863 | 59598 | female | female |
| Proband Diaphragm 809 | FALSE | 27260 | 2.115 | 12887 | 59569 | female | female |
| Father 967 | FALSE | 950 | 0.0346 | 27459 | 70883 | male | male |
| Mother 967 | FALSE | 40982 | 2.334 | 17558 | 41044 | female | female |
| Proband Blood 967 | FALSE | 917 | 0.0246 | 37276 | 60550 | male | male |
| Proband Skin 967 | FALSE | 1108 | 0.02955 | 37491 | 60504 | male | male |
| Proband Diaphragm 967 | FALSE | 830 | 0.02243 | 37005 | 60545 | male | male |

### TABLE S2

| Sample A | Sample B | Number of sites sampled | Shared heterozygous sites | Heterozygous sites in sample A | Identity by state 2 | Relatedness coefficent | Heterozygous sites in sample B | Identity by state 0 | Sample A reported as parent of sample B | Pedigree Relatedness | Predicted Parents | Parent Error | Sample Duplication | |
|---|---|---|---|---|---|---|---|---|---|---|---|---|---|---|
| Father 411 | Mother 411 | 23069 | 3035 | 7709 | 12331 | 0.01635 | 7646 | 1455 | FALSE | 0 | FALSE | FALSE | FALSE | |
| Father 411 | Proband Blood 411 | 23075 | 3876 | 7709 | 15502 | 0.5064 | 7610 | 11 | TRUE | 0.5 | TRUE | FALSE | FALSE | |
| Father 411 | Proband Skin 411 | 23073 | 3876 | 7709 | 15504 | 0.5072 | 7611 | 8 | TRUE | 0.5 | TRUE | FALSE | FALSE | |
| Father 411 | Proband Diaphragm 411 | 23076 | 3879 | 7709 | 15504 | 0.5076 | 7615 | 7 | TRUE | 0.5 | TRUE | FALSE | FALSE | |
| Father 411 | Father 716 | 23071 | 2615 | 7709 | 11733 | -0.08225 | 7258 | 1606 | FALSE | 0 | FALSE | FALSE | FALSE | |
| Father 411 | Mother 716 | 23068 | 2702 | 7709 | 11739 | -0.09578 | 7329 | 1702 | FALSE | 0 | FALSE | FALSE | FALSE | |
| Father 411 | Proband Blood 716 | 23063 | 2688 | 7709 | 11733 | -0.09893 | 7298 | 1705 | FALSE | 0 | FALSE | FALSE | FALSE | |
| Father 411 | Proband Skin 716 | 23066 | 2691 | 7709 | 11734 | -0.09767 | 7300 | 1702 | FALSE | 0 | FALSE | FALSE | FALSE | |
| Father 411 | Proband Diaphragm 716 | 23064 | 2689 | 7709 | 11739 | -0.09893 | 7288 | 1705 | FALSE | 0 | FALSE | FALSE | FALSE | |
| Father 411 | Father 809 | 23073 | 3040 | 7709 | 12132 | 0.0205 | 7858 | 1441 | FALSE | 0 | FALSE | FALSE | FALSE | |
| Father 411 | Mother 809 | 23068 | 2670 | 7709 | 11803 | -0.07723 | 7277 | 1616 | FALSE | 0 | FALSE | FALSE | FALSE | |
| Father 411 | Proband Blood 809 | 23068 | 2936 | 7709 | 12063 | -0.01888 | 7626 | 1540 | FALSE | 0 | FALSE | FALSE | FALSE | |
| Father 411 | Proband Skin 809 | 23070 | 2938 | 7709 | 12070 | -0.01861 | 7629 | 1540 | FALSE | 0 | FALSE | FALSE | FALSE | |
| Father 411 | Proband Diaphragm 809 | 23062 | 2939 | 7709 | 12064 | -0.01822 | 7630 | 1539 | FALSE | 0 | FALSE | FALSE | FALSE | |
| Father 411 | Father 967 | 23068 | 2863 | 7709 | 11656 | -0.05124 | 7801 | 1629 | FALSE | 0 | FALSE | FALSE | FALSE | |
| Father 411 | Mother 967 | 23068 | 2752 | 7709 | 11959 | -0.05179 | 7337 | 1566 | FALSE | 0 | FALSE | FALSE | FALSE | |
| Father 411 | Proband Blood 967 | 23071 | 2738 | 7709 | 11843 | -0.05606 | 7420 | 1577 | FALSE | 0 | FALSE | FALSE | FALSE | |
| Father 411 | Proband Skin 967 | 23074 | 2740 | 7709 | 11845 | -0.05522 | 7425 | 1575 | FALSE | 0 | FALSE | FALSE | FALSE | |
| Father 411 | Proband Diaphragm 967 | 23065 | 2737 | 7709 | 11835 | -0.05655 | 7409 | 1578 | FALSE | 0 | FALSE | FALSE | FALSE | |
| Mother 411 | Proband Blood 411 | 23064 | 3808 | 7646 | 15429 | 0.4996 | 7610 | 3 | TRUE | 0.5 | TRUE | FALSE | FALSE | |
| Mother 411 | Proband Skin 411 | 23059 | 3807 | 7646 | 15421 | 0.4994 | 7611 | 3 | TRUE | 0.5 | TRUE | FALSE | FALSE | |
| Mother 411 | Proband Diaphragm 411 | 23066 | 3808 | 7646 | 15432 | 0.4995 | 7615 | 2 | TRUE | 0.5 | TRUE | FALSE | FALSE | |
| Mother 411 | Father 716 | 23062 | 2761 | 7646 | 12110 | -0.05304 | 7258 | 1573 | FALSE | 0 | FALSE | FALSE | FALSE | |
| Mother 411 | Mother 716 | 23060 | 2648 | 7646 | 11694 | -0.1004 | 7329 | 1692 | FALSE | 0 | FALSE | FALSE | FALSE | |
| Mother 411 | Proband Blood 716 | 23055 | 2726 | 7646 | 11921 | -0.07947 | 7298 | 1653 | FALSE | 0 | FALSE | FALSE | FALSE | |
| Mother 411 | Proband Skin 716 | 23060 | 2726 | 7646 | 11917 | -0.07863 | 7300 | 1650 | FALSE | 0 | FALSE | FALSE | FALSE | |
| Mother 411 | Proband Diaphragm 716 | 23057 | 2726 | 7646 | 11922 | -0.07876 | 7288 | 1650 | FALSE | 0 | FALSE | FALSE | FALSE | |
| Mother 411 | Father 809 | 23064 | 2982 | 7646 | 12162 | 0.03244 | 7858 | 1367 | FALSE | 0 | FALSE | FALSE | FALSE | |
| Mother 411 | Mother 809 | 23060 | 2704 | 7646 | 12054 | -0.04205 | 7277 | 1505 | FALSE | 0 | FALSE | FALSE | FALSE | |
| Mother 411 | Proband Blood 809 | 23059 | 2917 | 7646 | 12202 | 0.008523 | 7626 | 1426 | FALSE | 0 | FALSE | FALSE | FALSE | |
| Mother 411 | Proband Skin 809 | 23063 | 2920 | 7646 | 12203 | 0.008913 | 7629 | 1426 | FALSE | 0 | FALSE | FALSE | FALSE | |
| Mother 411 | Proband Diaphragm 809 | 23053 | 2919 | 7646 | 12199 | 0.008781 | 7630 | 1426 | FALSE | 0 | FALSE | FALSE | FALSE | |
| Mother 411 | Father 967 | 23062 | 2755 | 7646 | 11602 | -0.0412 | 7801 | 1535 | FALSE | 0 | FALSE | FALSE | FALSE | |
| Mother 411 | Mother 967 | 23060 | 2723 | 7646 | 11867 | -0.08137 | 7337 | 1660 | FALSE | 0 | FALSE | FALSE | FALSE | |
| Mother 411 | Proband Blood 967 | 23063 | 2759 | 7646 | 11893 | -0.06968 | 7420 | 1638 | FALSE | 0 | FALSE | FALSE | FALSE | |
| Mother 411 | Proband Skin 967 | 23065 | 2763 | 7646 | 11893 | -0.06855 | 7425 | 1636 | FALSE | 0 | FALSE | FALSE | FALSE | |
| Mother 411 | Proband Diaphragm 967 | 23060 | 2759 | 7646 | 11887 | -0.07032 | 7409 | 1640 | FALSE | 0 | FALSE | FALSE | FALSE | |
| Proband Blood 411 | Proband Skin 411 | 23079 | 7601 | 7610 | 23067 | 0.9986 | 7611 | 1 | FALSE | 0.5 | TRUE | TRUE | TRUE | different tissue samples from same proband |
| Proband Blood 411 | Proband Diaphragm 411 | 23082 | 7606 | 7610 | 23069 | 0.9995 | 7615 | 0 | FALSE | 0.5 | TRUE | TRUE | TRUE | different tissue samples from same proband |
| Proband Blood 411 | Father 716 | 23076 | 2677 | 7610 | 11988 | -0.06655 | 7258 | 1580 | FALSE | 0 | FALSE | FALSE | FALSE | |
| Proband Blood 411 | Mother 716 | 23072 | 2709 | 7610 | 11796 | -0.1139 | 7329 | 1772 | FALSE | 0 | FALSE | FALSE | FALSE | |
| Proband Blood 411 | Proband Blood 716 | 23068 | 2695 | 7610 | 11842 | -0.1002 | 7298 | 1713 | FALSE | 0 | FALSE | FALSE | FALSE | |
| Proband Blood 411 | Proband Skin 716 | 23071 | 2694 | 7610 | 11844 | -0.09973 | 7300 | 1711 | FALSE | 0 | FALSE | FALSE | FALSE | |
| Proband Blood 411 | Proband Diaphragm 716 | 23070 | 2692 | 7610 | 11853 | -0.1007 | 7288 | 1713 | FALSE | 0 | FALSE | FALSE | FALSE | |
| Proband Blood 411 | Father 809 | 23078 | 2955 | 7610 | 12063 | 0.007227 | 7858 | 1450 | FALSE | 0 | FALSE | FALSE | FALSE | |
| Proband Blood 411 | Mother 809 | 23072 | 2767 | 7610 | 12161 | -0.04906 | 7277 | 1562 | FALSE | 0 | FALSE | FALSE | FALSE | |
| Proband Blood 411 | Proband Blood 809 | 23074 | 2837 | 7610 | 12000 | -0.02378 | 7626 | 1509 | FALSE | 0 | FALSE | FALSE | FALSE | |
| Proband Blood 411 | Proband Skin 809 | 23076 | 2841 | 7610 | 12010 | -0.023 | 7629 | 1508 | FALSE | 0 | FALSE | FALSE | FALSE | |
| Proband Blood 411 | Proband Diaphragm 809 | 23066 | 2839 | 7610 | 11999 | -0.02326 | 7630 | 1508 | FALSE | 0 | FALSE | FALSE | FALSE | |
| Proband Blood 411 | Father 967 | 23074 | 2825 | 7610 | 11633 | -0.0703 | 7801 | 1680 | FALSE | 0 | FALSE | FALSE | FALSE | |
| Proband Blood 411 | Mother 967 | 23072 | 2773 | 7610 | 12030 | -0.06965 | 7337 | 1642 | FALSE | 0 | FALSE | FALSE | FALSE | |
| Proband Blood 411 | Proband Blood 967 | 23076 | 2733 | 7610 | 11878 | -0.07264 | 7420 | 1636 | FALSE | 0 | FALSE | FALSE | FALSE | |
| Proband Blood 411 | Proband Skin 967 | 23078 | 2732 | 7610 | 11876 | -0.07219 | 7425 | 1634 | FALSE | 0 | FALSE | FALSE | FALSE | |
| Proband Blood 411 | Proband Diaphragm 967 | 23072 | 2733 | 7610 | 11868 | -0.0741 | 7409 | 1641 | FALSE | 0 | FALSE | FALSE | FALSE | |
| Proband Skin 411 | Proband Diaphragm 411 | 23083 | 7602 | 7611 | 23067 | 0.9988 | 7615 | 0 | FALSE | 0.5 | TRUE | TRUE | TRUE | different tissue samples from same proband |
| Proband Skin 411 | Father 716 | 23081 | 2677 | 7611 | 11987 | -0.06682 | 7258 | 1581 | FALSE | 0 | FALSE | FALSE | FALSE | |
| Proband Skin 411 | Mother 716 | 23076 | 2713 | 7611 | 11796 | -0.1134 | 7329 | 1772 | FALSE | 0 | FALSE | FALSE | FALSE | |
| Proband Skin 411 | Proband Blood 716 | 23072 | 2693 | 7611 | 11840 | -0.1002 | 7298 | 1712 | FALSE | 0 | FALSE | FALSE | FALSE | |
| Proband Skin 411 | Proband Skin 716 | 23075 | 2693 | 7611 | 11838 | -0.09986 | 7300 | 1711 | FALSE | 0 | FALSE | FALSE | FALSE | |
| Proband Skin 411 | Proband Diaphragm 716 | 23073 | 2691 | 7611 | 11847 | -0.1006 | 7288 | 1712 | FALSE | 0 | FALSE | FALSE | FALSE | |
| Proband Skin 411 | Father 809 | 23080 | 2952 | 7611 | 12060 | 0.006307 | 7858 | 1452 | FALSE | 0 | FALSE | FALSE | FALSE | |

### TABLE S2

| | | | | | | | | | | | | | | |
|---|---|---|---|---|---|---|---|---|---|---|---|---|---|---|
| Proband Skin 411 | Mother 809 | 23075 | 2770 | 7611 | 12168 | -0.04865 | 7277 | 1562 | FALSE | 0 | FALSE | FALSE | FALSE | |
| Proband Skin 411 | Proband Blood 809 | 23075 | 2838 | 7611 | 12001 | -0.02391 | 7626 | 1510 | FALSE | 0 | FALSE | FALSE | FALSE | |
| Proband Skin 411 | Proband Skin 809 | 23077 | 2841 | 7611 | 12009 | -0.02326 | 7629 | 1509 | FALSE | 0 | FALSE | FALSE | FALSE | |
| Proband Skin 411 | Proband Diaphragm 809 | 23068 | 2839 | 7611 | 11998 | -0.02326 | 7630 | 1508 | FALSE | 0 | FALSE | FALSE | FALSE | |
| Proband Skin 411 | Father 967 | 23077 | 2825 | 7611 | 11632 | -0.07056 | 7801 | 1681 | FALSE | 0 | FALSE | FALSE | FALSE | |
| Proband Skin 411 | Mother 967 | 23078 | 2771 | 7611 | 12034 | -0.07046 | 7337 | 1644 | FALSE | 0 | FALSE | FALSE | FALSE | |
| Proband Skin 411 | Proband Blood 967 | 23079 | 2730 | 7611 | 11879 | -0.07358 | 7420 | 1638 | FALSE | 0 | FALSE | FALSE | FALSE | |
| Proband Skin 411 | Proband Skin 967 | 23081 | 2731 | 7611 | 11878 | -0.07286 | 7425 | 1636 | FALSE | 0 | FALSE | FALSE | FALSE | |
| Proband Skin 411 | Proband Diaphragm 967 | 23072 | 2730 | 7611 | 11866 | -0.07477 | 7409 | 1642 | FALSE | 0 | FALSE | FALSE | FALSE | |
| Proband Diaphragm 411 | Father 716 | 23085 | 2681 | 7615 | 11996 | -0.06627 | 7258 | 1581 | FALSE | 0 | FALSE | FALSE | FALSE | |
| Proband Diaphragm 411 | Mother 716 | 23078 | 2712 | 7615 | 11799 | -0.1135 | 7329 | 1772 | FALSE | 0 | FALSE | FALSE | FALSE | |
| Proband Diaphragm 411 | Proband Blood 716 | 23076 | 2696 | 7615 | 11848 | -0.1 | 7298 | 1713 | FALSE | 0 | FALSE | FALSE | FALSE | |
| Proband Diaphragm 411 | Proband Skin 716 | 23079 | 2697 | 7615 | 11852 | -0.09959 | 7300 | 1712 | FALSE | 0 | FALSE | FALSE | FALSE | |
| Proband Diaphragm 411 | Proband Diaphragm 716 | 23077 | 2693 | 7615 | 11856 | -0.1003 | 7288 | 1712 | FALSE | 0 | FALSE | FALSE | FALSE | |
| Proband Diaphragm 411 | Father 809 | 23085 | 2955 | 7615 | 12065 | 0.007223 | 7858 | 1450 | FALSE | 0 | FALSE | FALSE | FALSE | |
| Proband Diaphragm 411 | Mother 809 | 23080 | 2770 | 7615 | 12172 | -0.04837 | 7277 | 1561 | FALSE | 0 | FALSE | FALSE | FALSE | |
| Proband Diaphragm 411 | Proband Blood 809 | 23082 | 2840 | 7615 | 12006 | -0.0239 | 7626 | 1511 | FALSE | 0 | FALSE | FALSE | FALSE | |
| Proband Diaphragm 411 | Proband Skin 809 | 23084 | 2843 | 7615 | 12016 | -0.02324 | 7629 | 1510 | FALSE | 0 | FALSE | FALSE | FALSE | |
| Proband Diaphragm 411 | Proband Diaphragm 809 | 23075 | 2842 | 7615 | 12006 | -0.02337 | 7630 | 1510 | FALSE | 0 | FALSE | FALSE | FALSE | |
| Proband Diaphragm 411 | Father 967 | 23081 | 2826 | 7615 | 11640 | -0.07039 | 7801 | 1681 | FALSE | 0 | FALSE | FALSE | FALSE | |
| Proband Diaphragm 411 | Mother 967 | 23080 | 2775 | 7615 | 12032 | -0.06937 | 7337 | 1642 | FALSE | 0 | FALSE | FALSE | FALSE | |
| Proband Diaphragm 411 | Proband Blood 967 | 23084 | 2736 | 7615 | 11890 | -0.07278 | 7420 | 1638 | FALSE | 0 | FALSE | FALSE | FALSE | |
| Proband Diaphragm 411 | Proband Skin 967 | 23086 | 2736 | 7615 | 11888 | -0.07192 | 7425 | 1635 | FALSE | 0 | FALSE | FALSE | FALSE | |
| Proband Diaphragm 411 | Proband Diaphragm 967 | 23078 | 2735 | 7615 | 11877 | -0.0741 | 7409 | 1642 | FALSE | 0 | FALSE | FALSE | FALSE | |
| Father 716 | Mother 716 | 23075 | 2805 | 7258 | 12804 | 0.02935 | 7329 | 1296 | FALSE | 0 | FALSE | FALSE | FALSE | |
| Father 716 | Proband Blood 716 | 23077 | 3728 | 7258 | 15986 | 0.5123 | 7298 | 5 | TRUE | 0.5 | TRUE | FALSE | FALSE | |
| Father 716 | Proband Skin 716 | 23080 | 3734 | 7258 | 15990 | 0.5131 | 7300 | 5 | TRUE | 0.5 | TRUE | FALSE | FALSE | |
| Father 716 | Proband Diaphragm 716 | 23078 | 3725 | 7258 | 15987 | 0.5118 | 7288 | 5 | TRUE | 0.5 | TRUE | FALSE | FALSE | |
| Father 716 | Father 809 | 23081 | 2805 | 7258 | 12039 | -0.03541 | 7858 | 1531 | FALSE | 0 | FALSE | FALSE | FALSE | |
| Father 716 | Mother 809 | 23077 | 2486 | 7258 | 11754 | -0.1414 | 7277 | 1756 | FALSE | 0 | FALSE | FALSE | FALSE | |
| Father 716 | Proband Blood 809 | 23077 | 2617 | 7258 | 11822 | -0.0806 | 7626 | 1601 | FALSE | 0 | FALSE | FALSE | FALSE | |
| Father 716 | Proband Skin 809 | 23080 | 2617 | 7258 | 11834 | -0.08143 | 7629 | 1604 | FALSE | 0 | FALSE | FALSE | FALSE | |
| Father 716 | Proband Diaphragm 809 | 23070 | 2615 | 7258 | 11823 | -0.08088 | 7630 | 1601 | FALSE | 0 | FALSE | FALSE | FALSE | |
| Father 716 | Father 967 | 23077 | 2848 | 7258 | 12219 | -0.02067 | 7801 | 1499 | FALSE | 0 | FALSE | FALSE | FALSE | |
| Father 716 | Mother 967 | 23079 | 2617 | 7258 | 12064 | -0.09493 | 7337 | 1653 | FALSE | 0 | FALSE | FALSE | FALSE | |
| Father 716 | Proband Blood 967 | 23081 | 2646 | 7258 | 12183 | -0.05291 | 7420 | 1515 | FALSE | 0 | FALSE | FALSE | FALSE | |
| Father 716 | Proband Skin 967 | 23082 | 2647 | 7258 | 12184 | -0.05277 | 7425 | 1515 | FALSE | 0 | FALSE | FALSE | FALSE | |
| Father 716 | Proband Diaphragm 967 | 23074 | 2642 | 7258 | 12175 | -0.05456 | 7409 | 1519 | FALSE | 0 | FALSE | FALSE | FALSE | |
| Mother 716 | Proband Blood 716 | 23074 | 3756 | 7329 | 15962 | 0.5136 | 7298 | 4 | TRUE | 0.5 | TRUE | FALSE | FALSE | |
| Mother 716 | Proband Skin 716 | 23079 | 3760 | 7329 | 15974 | 0.5142 | 7300 | 3 | TRUE | 0.5 | TRUE | FALSE | FALSE | |
| Mother 716 | Proband Diaphragm 716 | 23076 | 3756 | 7329 | 15964 | 0.5145 | 7288 | 3 | TRUE | 0.5 | TRUE | FALSE | FALSE | |
| Mother 716 | Father 809 | 23080 | 2766 | 7329 | 11815 | -0.06276 | 7858 | 1613 | FALSE | 0 | FALSE | FALSE | FALSE | |
| Mother 716 | Mother 809 | 23077 | 2508 | 7329 | 11692 | -0.1476 | 7277 | 1791 | FALSE | 0 | FALSE | FALSE | FALSE | |
| Mother 716 | Proband Blood 809 | 23076 | 2702 | 7329 | 11859 | -0.0876 | 7626 | 1672 | FALSE | 0 | FALSE | FALSE | FALSE | |
| Mother 716 | Proband Skin 809 | 23079 | 2699 | 7329 | 11851 | -0.08773 | 7629 | 1671 | FALSE | 0 | FALSE | FALSE | FALSE | |
| Mother 716 | Proband Diaphragm 809 | 23070 | 2704 | 7329 | 11851 | -0.08705 | 7630 | 1671 | FALSE | 0 | FALSE | FALSE | FALSE | |
| Mother 716 | Father 967 | 23078 | 2776 | 7329 | 11857 | -0.06931 | 7801 | 1642 | FALSE | 0 | FALSE | FALSE | FALSE | |
| Mother 716 | Mother 967 | 23078 | 2451 | 7329 | 11612 | -0.1314 | 7337 | 1707 | FALSE | 0 | FALSE | FALSE | FALSE | |
| Mother 716 | Proband Blood 967 | 23080 | 2681 | 7329 | 12007 | -0.09838 | 7420 | 1701 | FALSE | 0 | FALSE | FALSE | FALSE | |
| Mother 716 | Proband Skin 967 | 23082 | 2687 | 7329 | 12006 | -0.09728 | 7425 | 1700 | FALSE | 0 | FALSE | FALSE | FALSE | |
| Mother 716 | Proband Diaphragm 967 | 23075 | 2676 | 7329 | 11996 | -0.09906 | 7409 | 1701 | FALSE | 0 | FALSE | FALSE | FALSE | |
| Proband Blood 716 | Proband Skin 716 | 23086 | 7292 | 7298 | 23074 | 0.9992 | 7300 | 0 | FALSE | 0.5 | TRUE | TRUE | TRUE | different tissue samples from same proband |
| Proband Blood 716 | Proband Diaphragm 716 | 23085 | 7281 | 7298 | 23069 | 0.999 | 7288 | 0 | FALSE | 0.5 | TRUE | TRUE | TRUE | different tissue samples from same proband |
| Proband Blood 716 | Father 809 | 23082 | 2797 | 7298 | 11981 | -0.03796 | 7858 | 1537 | FALSE | 0 | FALSE | FALSE | FALSE | |
| Proband Blood 716 | Mother 809 | 23077 | 2534 | 7298 | 11741 | -0.1536 | 7277 | 1826 | FALSE | 0 | FALSE | FALSE | FALSE | |
| Proband Blood 716 | Proband Blood 809 | 23077 | 2689 | 7298 | 11906 | -0.07632 | 7626 | 1623 | FALSE | 0 | FALSE | FALSE | FALSE | |
| Proband Blood 716 | Proband Skin 809 | 23079 | 2688 | 7298 | 11907 | -0.07591 | 7629 | 1621 | FALSE | 0 | FALSE | FALSE | FALSE | |
| Proband Blood 716 | Proband Diaphragm 809 | 23070 | 2689 | 7298 | 11904 | -0.07577 | 7630 | 1621 | FALSE | 0 | FALSE | FALSE | FALSE | |
| Proband Blood 716 | Father 967 | 23078 | 2718 | 7298 | 11892 | -0.04549 | 7801 | 1525 | FALSE | 0 | FALSE | FALSE | FALSE | |
| Proband Blood 716 | Mother 967 | 23079 | 2589 | 7298 | 11942 | -0.1056 | 7337 | 1680 | FALSE | 0 | FALSE | FALSE | FALSE | |
| Proband Blood 716 | Proband Blood 967 | 23083 | 2663 | 7298 | 12100 | -0.07029 | 7420 | 1588 | FALSE | 0 | FALSE | FALSE | FALSE | |
| Proband Blood 716 | Proband Skin 967 | 23084 | 2664 | 7298 | 12100 | -0.0707 | 7425 | 1590 | FALSE | 0 | FALSE | FALSE | FALSE | |
| Proband Blood 716 | Proband Diaphragm 967 | 23076 | 2662 | 7298 | 12096 | -0.0718 | 7409 | 1593 | FALSE | 0 | FALSE | FALSE | FALSE | |
| Proband Skin 716 | Proband Diaphragm 716 | 23081 | 7283 | 7300 | 23071 | 0.9993 | 7288 | 0 | FALSE | 0.5 | TRUE | TRUE | TRUE | different tissue samples from same proband |
| Proband Skin 716 | Father 809 | 23078 | 2800 | 7300 | 11986 | -0.03726 | 7858 | 1536 | FALSE | 0 | FALSE | FALSE | FALSE | |

### TABLE S2

| | | | | | | | | | | | | | | |
|---|---|---|---|---|---|---|---|---|---|---|---|---|---|---|
| Proband Skin 716 | Mother 809 | 23075 | 2534 | 7300 | 11741 | -0.1525 | 7277 | 1822 | FALSE | 0 | FALSE | FALSE | FALSE | |
| Proband Skin 716 | Proband Blood 809 | 23074 | 2689 | 7300 | 11908 | -0.07603 | 7626 | 1622 | FALSE | 0 | FALSE | FALSE | FALSE | |
| Proband Skin 716 | Proband Skin 809 | 23077 | 2689 | 7300 | 11909 | -0.07548 | 7629 | 1620 | FALSE | 0 | FALSE | FALSE | FALSE | |
| Proband Skin 716 | Proband Diaphragm 809 | 23068 | 2689 | 7300 | 11904 | -0.07548 | 7630 | 1620 | FALSE | 0 | FALSE | FALSE | FALSE | |
| Proband Skin 716 | Father 967 | 23076 | 2718 | 7300 | 11888 | -0.04575 | 7801 | 1526 | FALSE | 0 | FALSE | FALSE | FALSE | |
| Proband Skin 716 | Mother 967 | 23076 | 2587 | 7300 | 11939 | -0.1053 | 7337 | 1678 | FALSE | 0 | FALSE | FALSE | FALSE | |
| Proband Skin 716 | Proband Blood 967 | 23079 | 2664 | 7300 | 12101 | -0.06986 | 7420 | 1587 | FALSE | 0 | FALSE | FALSE | FALSE | |
| Proband Skin 716 | Proband Skin 967 | 23080 | 2665 | 7300 | 12101 | -0.07 | 7425 | 1588 | FALSE | 0 | FALSE | FALSE | FALSE | |
| Proband Skin 716 | Proband Diaphragm 967 | 23074 | 2663 | 7300 | 12098 | -0.0711 | 7409 | 1591 | FALSE | 0 | FALSE | FALSE | FALSE | |
| Proband Diaphragm 716 | Father 809 | 23078 | 2799 | 7288 | 11991 | -0.03801 | 7858 | 1538 | FALSE | 0 | FALSE | FALSE | FALSE | |
| Proband Diaphragm 716 | Mother 809 | 23074 | 2534 | 7288 | 11747 | -0.1534 | 7277 | 1825 | FALSE | 0 | FALSE | FALSE | FALSE | |
| Proband Diaphragm 716 | Proband Blood 809 | 23073 | 2686 | 7288 | 11909 | -0.07739 | 7626 | 1625 | FALSE | 0 | FALSE | FALSE | FALSE | |
| Proband Diaphragm 716 | Proband Skin 809 | 23075 | 2686 | 7288 | 11908 | -0.07684 | 7629 | 1623 | FALSE | 0 | FALSE | FALSE | FALSE | |
| Proband Diaphragm 716 | Proband Diaphragm 809 | 23066 | 2688 | 7288 | 11906 | -0.07629 | 7630 | 1622 | FALSE | 0 | FALSE | FALSE | FALSE | |
| Proband Diaphragm 716 | Father 967 | 23074 | 2718 | 7288 | 11890 | -0.04555 | 7801 | 1525 | FALSE | 0 | FALSE | FALSE | FALSE | |
| Proband Diaphragm 716 | Mother 967 | 23075 | 2584 | 7288 | 11941 | -0.1062 | 7337 | 1679 | FALSE | 0 | FALSE | FALSE | FALSE | |
| Proband Diaphragm 716 | Proband Blood 967 | 23078 | 2664 | 7288 | 12109 | -0.06998 | 7420 | 1587 | FALSE | 0 | FALSE | FALSE | FALSE | |
| Proband Diaphragm 716 | Proband Skin 967 | 23079 | 2665 | 7288 | 12111 | -0.07039 | 7425 | 1589 | FALSE | 0 | FALSE | FALSE | FALSE | |
| Proband Diaphragm 716 | Proband Diaphragm 967 | 23072 | 2663 | 7288 | 12107 | -0.07176 | 7409 | 1593 | FALSE | 0 | FALSE | FALSE | FALSE | |
| Father 809 | Mother 809 | 23072 | 2890 | 7858 | 12247 | -0.007146 | 7277 | 1471 | FALSE | 0 | FALSE | FALSE | FALSE | |
| Father 809 | Proband Blood 809 | 23075 | 3976 | 7858 | 15550 | 0.5206 | 7626 | 3 | TRUE | 0.5 | TRUE | FALSE | FALSE | |
| Father 809 | Proband Skin 809 | 23079 | 3976 | 7858 | 15556 | 0.5204 | 7629 | 3 | TRUE | 0.5 | TRUE | FALSE | FALSE | |
| Father 809 | Proband Diaphragm 809 | 23070 | 3976 | 7858 | 15547 | 0.5203 | 7630 | 3 | TRUE | 0.5 | TRUE | FALSE | FALSE | |
| Father 809 | Father 967 | 23071 | 2865 | 7858 | 11682 | -0.009614 | 7801 | 1470 | FALSE | 0 | FALSE | FALSE | FALSE | |
| Father 809 | Mother 967 | 23071 | 2776 | 7858 | 11726 | -0.08777 | 7337 | 1710 | FALSE | 0 | FALSE | FALSE | FALSE | |
| Father 809 | Proband Blood 967 | 23075 | 2769 | 7858 | 11739 | -0.05863 | 7420 | 1602 | FALSE | 0 | FALSE | FALSE | FALSE | |
| Father 809 | Proband Skin 967 | 23077 | 2771 | 7858 | 11740 | -0.05832 | 7425 | 1602 | FALSE | 0 | FALSE | FALSE | FALSE | |
| Father 809 | Proband Diaphragm 967 | 23069 | 2765 | 7858 | 11735 | -0.05952 | 7409 | 1603 | FALSE | 0 | FALSE | FALSE | FALSE | |
| Mother 809 | Proband Blood 809 | 23074 | 3578 | 7277 | 15327 | 0.4895 | 7626 | 8 | TRUE | 0.5 | TRUE | FALSE | FALSE | |
| Mother 809 | Proband Skin 809 | 23077 | 3579 | 7277 | 15330 | 0.4894 | 7629 | 9 | TRUE | 0.5 | TRUE | FALSE | FALSE | |
| Mother 809 | Proband Diaphragm 809 | 23070 | 3583 | 7277 | 15327 | 0.4899 | 7630 | 9 | TRUE | 0.5 | TRUE | FALSE | FALSE | |
| Mother 809 | Father 967 | 23073 | 2755 | 7277 | 11751 | -0.1049 | 7801 | 1759 | FALSE | 0 | FALSE | FALSE | FALSE | |
| Mother 809 | Mother 967 | 23073 | 2453 | 7277 | 11682 | -0.1257 | 7337 | 1684 | FALSE | 0 | FALSE | FALSE | FALSE | |
| Mother 809 | Proband Blood 967 | 23076 | 2574 | 7277 | 11764 | -0.1308 | 7420 | 1763 | FALSE | 0 | FALSE | FALSE | FALSE | |
| Mother 809 | Proband Skin 967 | 23079 | 2577 | 7277 | 11770 | -0.131 | 7425 | 1765 | FALSE | 0 | FALSE | FALSE | FALSE | |
| Mother 809 | Proband Diaphragm 967 | 23070 | 2573 | 7277 | 11763 | -0.1321 | 7409 | 1767 | FALSE | 0 | FALSE | FALSE | FALSE | |
| Proband Blood 809 | Proband Skin 809 | 23084 | 7620 | 7626 | 23070 | 0.999 | 7629 | 1 | FALSE | 0.5 | TRUE | TRUE | TRUE | different tissue samples from same proband |
| Proband Blood 809 | Proband Diaphragm 809 | 23075 | 7617 | 7626 | 23063 | 0.9988 | 7630 | 0 | FALSE | 0.5 | TRUE | TRUE | TRUE | different tissue samples from same proband |
| Proband Blood 809 | Father 967 | 23072 | 2840 | 7626 | 11737 | -0.04563 | 7801 | 1594 | FALSE | 0 | FALSE | FALSE | FALSE | |
| Proband Blood 809 | Mother 967 | 23072 | 2705 | 7626 | 11860 | -0.08409 | 7337 | 1661 | FALSE | 0 | FALSE | FALSE | FALSE | |
| Proband Blood 809 | Proband Blood 967 | 23076 | 2744 | 7626 | 11863 | -0.07682 | 7420 | 1657 | FALSE | 0 | FALSE | FALSE | FALSE | |
| Proband Blood 809 | Proband Skin 967 | 23077 | 2745 | 7626 | 11867 | -0.07663 | 7425 | 1657 | FALSE | 0 | FALSE | FALSE | FALSE | |
| Proband Blood 809 | Proband Diaphragm 967 | 23069 | 2738 | 7626 | 11858 | -0.07801 | 7409 | 1658 | FALSE | 0 | FALSE | FALSE | FALSE | |
| Proband Skin 809 | Proband Diaphragm 809 | 23080 | 7620 | 7629 | 23071 | 0.9988 | 7630 | 0 | FALSE | 0.5 | TRUE | TRUE | TRUE | different tissue samples from same proband |
| Proband Skin 809 | Father 967 | 23079 | 2842 | 7629 | 11745 | -0.0464 | 7801 | 1598 | FALSE | 0 | FALSE | FALSE | FALSE | |
| Proband Skin 809 | Mother 967 | 23079 | 2706 | 7629 | 11864 | -0.08478 | 7337 | 1664 | FALSE | 0 | FALSE | FALSE | FALSE | |
| Proband Skin 809 | Proband Blood 967 | 23082 | 2747 | 7629 | 11871 | -0.07642 | 7420 | 1657 | FALSE | 0 | FALSE | FALSE | FALSE | |
| Proband Skin 809 | Proband Skin 967 | 23084 | 2747 | 7629 | 11873 | -0.07636 | 7425 | 1657 | FALSE | 0 | FALSE | FALSE | FALSE | |
| Proband Skin 809 | Proband Diaphragm 967 | 23076 | 2740 | 7629 | 11863 | -0.07774 | 7409 | 1658 | FALSE | 0 | FALSE | FALSE | FALSE | |
| Proband Diaphragm 809 | Father 967 | 23074 | 2843 | 7630 | 11739 | -0.04574 | 7801 | 1596 | FALSE | 0 | FALSE | FALSE | FALSE | |
| Proband Diaphragm 809 | Mother 967 | 23072 | 2707 | 7630 | 11862 | -0.08355 | 7337 | 1660 | FALSE | 0 | FALSE | FALSE | FALSE | |
| Proband Diaphragm 809 | Proband Blood 967 | 23075 | 2746 | 7630 | 11862 | -0.07601 | 7420 | 1655 | FALSE | 0 | FALSE | FALSE | FALSE | |
| Proband Diaphragm 809 | Proband Skin 967 | 23078 | 2748 | 7630 | 11866 | -0.07542 | 7425 | 1654 | FALSE | 0 | FALSE | FALSE | FALSE | |
| Proband Diaphragm 809 | Proband Diaphragm 967 | 23068 | 2740 | 7630 | 11856 | -0.07693 | 7409 | 1655 | FALSE | 0 | FALSE | FALSE | FALSE | |
| Father 967 | Mother 967 | 23076 | 2760 | 7801 | 12039 | -0.01009 | 7337 | 1417 | FALSE | 0 | FALSE | FALSE | FALSE | |
| Father 967 | Proband Blood 967 | 23081 | 3818 | 7801 | 15499 | 0.5135 | 7420 | 4 | TRUE | 0.5 | TRUE | FALSE | FALSE | |
| Father 967 | Proband Skin 967 | 23082 | 3821 | 7801 | 15505 | 0.5133 | 7425 | 5 | TRUE | 0.5 | TRUE | FALSE | FALSE | |
| Father 967 | Proband Diaphragm 967 | 23077 | 3818 | 7801 | 15497 | 0.5132 | 7409 | 8 | TRUE | 0.5 | TRUE | FALSE | FALSE | |
| Mother 967 | Proband Blood 967 | 23076 | 3598 | 7337 | 15520 | 0.4885 | 7420 | 7 | TRUE | 0.5 | TRUE | FALSE | FALSE | |
| Mother 967 | Proband Skin 967 | 23078 | 3601 | 7337 | 15523 | 0.4892 | 7425 | 6 | TRUE | 0.5 | TRUE | FALSE | FALSE | |
| Mother 967 | Proband Diaphragm 967 | 23069 | 3593 | 7337 | 15516 | 0.4878 | 7409 | 7 | TRUE | 0.5 | TRUE | FALSE | FALSE | |
| Proband Blood 967 | Proband Skin 967 | 23088 | 7415 | 7420 | 23079 | 0.9993 | 7425 | 0 | FALSE | 0.5 | TRUE | TRUE | TRUE | different tissue samples from same proband |
| Proband Blood 967 | Proband Diaphragm 967 | 23082 | 7396 | 7420 | 23055 | 0.9982 | 7409 | 0 | FALSE | 0.5 | TRUE | TRUE | TRUE | different tissue samples from same proband |
| Proband Skin 967 | Proband Diaphragm 967 | 23086 | 7402 | 7425 | 23065 | 0.9991 | 7409 | 0 | FALSE | 0.5 | TRUE | TRUE | TRUE | different tissue samples from same proband |

### TABLE S3

| Proband | Total Variants | De Novo Exon Variants in Germline (in Diaphragm, Skin, and Blood) | De Novo UTR + Intron Variants in Germline (in Diaphragm, Skin, + Blood) | De Novo Intergenic Variants in Germline (in Diaphragm, Skin, + Blood) | Total De Novo Variants in Germline (in Diaphragm, Skin, + Blood) | De Novo Exon Variants in Diaphragm+Blood | De Novo UTR + Intron Variants in Diaphragm+Blood | De Novo Intergenic Variants in Diaphragm+Blood | Total De Novo Variants in Diaphragm+Blood | De Novo Exon Variants in Diaphragm+Skin | De Novo UTR + Intron Variants in Diaphragm+Skin | De Novo Intergenic Variants in Diaphragm+Skin | Total De Novo Variants in Diaphragm+Skin | De Novo Exon Variants in Skin+Blood | De Novo UTR + Intron Variants in Skin+Blood | De Novo Intergenic Variants in Skin+Blood |
|---|---|---|---|---|---|---|---|---|---|---|---|---|---|---|---|---|
| **411** | **63** | 1 | 23 | 24 | **48** | 0 | 0 | 0 | **0** | 0 | 0 | 0 | **0** | 0 | 0 | 0 |
| **716** | **123** | 3 | 50 | 63 | **116** | 0 | 0 | 0 | **0** | 0 | 0 | 0 | **0** | 0 | 0 | 0 |
| **809** | **265** | 1 | 27 | 28 | **56** | 0 | 0 | 0 | **0** | 0 | 0 | 0 | **0** | 0 | 0 | 0 |
| **967** | **65** | 1 | 20 | 29 | **50** | 0 | 0 | 0 | **0** | 0 | 0 | 0 | **0** | 0 | 0 | 0 |

### TABLE S3

| Proband | Total Variants | Total De Novo Variants in Skin+Blood | De Novo Exon Variants in Diaphragm | De Novo UTR + Intron Variants in Diaphragm | De Novo Intergenic Variants in Diaphragm | Total De Novo Variants in Diaphragm | De Novo Exon Variants in Skin | De Novo UTR + Intron Variants in Skin | De Novo Intergenic Variants in Skin | Total De Novo Variants in Skin | De Novo Exon Variants in Blood | De Novo UTR + Intron Variants in Blood | De Novo Intergenic Variants in Blood | Total De Novo Variants in Blood |
|---|---|---|---|---|---|---|---|---|---|---|---|---|---|---|
| 411 | 63 | 0 | 0 | 0 | 3 | 3 | 0 | 3 | 3 | 6 | 1 | 1 | 0 | 2 |
| 716 | 123 | 0 | 0 | 0 | 1 | 1 | 0 | 1 | 3 | 4 | 0 | 0 | 0 | 0 |
| 809 | 265 | 0 | 0 | 40 | 64 | 104 | 3 | 31 | 65 | 99 | 0 | 0 | 0 | 0 |
| 967 | 65 | 0 | 0 | 1 | 2 | 3 | 0 | 1 | 6 | 7 | 0 | 0 | 1 | 1 |

### TABLE S3

| CDH Proband Family | Sex | Gene | Gene full name | Location in Gene | Chromosome | Position HG19 | Reference allele | Alternate allele | Resulting change | PROVEAN predicted effect | ExAC Pli 1= intolerant 0= tolerant of haploinsufficiency |
|---|---|---|---|---|---|---|---|---|---|---|---|
| 411 | Male | PEX6 | peroxisomal biogenesis factor 6 | exon 10 | 6 | 42934352 | CA | C | **Frameshift** | **N/A -Deleterious** | 0 |
| 411 | Male | MIR4717 | MicroRNA 4717 | exon 1 | 16 | 2324626 | [illegible]GTGGCCTAA | T | Missense | N/A | ? |
| 716 | Female | OLFM3 | olfactomedin 3 | exon 6 | 1 | 102270146 | G | C | **Missense** | **Deleterious** | 0 |
| 716 | Female | SCARB1 | scavenger receptor class B: member 1 | exon 1 | 12 | 125348217 | G | T | Missense | Neutral | 0.08 |
| 716 | Female | ZNF792 | zinc finger protein 792 | exon 4 | 19 | 35449333 | G | A | **Missense** | **Deleterious** | 0 |
| 809 | Female | AR | androgen receptor | exon 7 | X | 66942720 | A | G | Missense | Neutral | **0.99** |
| 809 | Female | DHX57 | DEAH (Asp-Glu-Ala-Asp/His) box polypeptide 57 | exon 14 | 2 | 39055578 | C | A | Missense | Deleterious | 0 |
| 809 | Female | TNFAIP8L3 | tumor necrosis factor: alpha-induced protein 8-like 3 | exon 1 | 15 | 51397258 | G | A | Missense | Neutral | 0 |
| 809 | Female | CXorf57 | Chromosome X open reading frame 57 | exon 11 | X | 105905393 | A | G | Synonymous | Neutral | 0.19 |
| 967 | Male | **THSD7A** | thrombospondin: type I: domain containing 7A | exon 2 | 7 | 11675999 | C | T | **Nonsense** | **N/A Deleterious** | **1** |

### TABLE S3

| CDH Proband Family | Sex | Gene | gnomAD frequency | CADD score, 10= top 10% damaging mutations, 40= 0.01% damaging mutatons | Protein Amino Acid Change | Tissue found in | Father | Mother | Proband Blood | Proband Skin | Proband Diaphragm |
|---|---|---|---|---|---|---|---|---|---|---|---|
| 411 | Male | PEX6 | Not found in gnomAD | NA | p.Cys668TrpfsTer46 | All three | 0/39 | 0/42 | 28/54 | 29/59 | 38/80 |
| 411 | Male | MIR4717 | Not found in gnomAD | NA | NA | Blood | 0/38 | 0/29 | 15/73 | 0/36 | 0/46 |
| 716 | Female | OLFM3 | Not found in gnomAD | 24.3 | p.Thr362Ser | All three | 0/55 | 0/61 | 42/79 | 48/81 | 33/68 |
| 716 | Female | SCARB1 | Not found in gnomAD | 9.987 | p.Ala17Glu | All three | 0/44 | 0/43 | 35/73 | 37/63 | 45/80 |
| 716 | Female | ZNF792 | 0.00001606 | 23.3 | p.Arg476Trp | All three | 0/64 | 0/50 | 24/56 | 34/77 | 29/81 |
| 809 | Female | AR | Not found in gnomAD | 23 | p.Asn834Ser | All three | 0/29 | 0/55 | 31/57 | 16/48 | 34/61 |
| 809 | Female | DHX57 | Not found in gnomAD | 34 | p.Gly848Val | Skin | 0/41 | 0/70 | 0/46 | 20/72 | 0/38 |
| 809 | Female | TNFAIP8L3 | 0.00009942 | 1.356 | p.Thr39Met | Skin | 0/37 | 0/54 | 0/54 | 20/58 | 0/47 |
| 809 | Female | CXorf57 | Not found in gnomAD | 9.28 | p.Lys612Lys | Skin | 0/39 | 0/58 | 0/51 | 25/63 | 0/56 |
| 967 | Male | **THSD7A** | Not found in gnomAD | 40 | p.Trp260Ter (STOP) | All three | 0/46 | 0/53 | 28/47 | 29/71 | 18/29 |

**TABLE S3**

| CDH Proband Family | Sex | Gene | Gene full name | Location | Chromosome | Position | Reference allele | Alternate allele |
|---|---|---|---|---|---|---|---|---|
| 411 | Male | KIF1B | kinesin family member 1B | intron 32 | 1 | 10405687 | C | T |
| 411 | Male | ZRANB3 | zinc finger: RAN-binding domain containing 3 | intron 2 | 2 | 136149974 | T | C |
| 411 | Male | SUMF1 | Formylglycine-generating enzyme | intron 8 | 3 | 4114688 | G | A |
| 411 | Male | HRH1 | histamine receptor H1 | intron 1 | 3 | 11223160 | T | A |
| 411 | male | SYN2 | synapsin II | intron 2 | 3 | 12146936 | T | A |
| 411 | Male | ADAMTS9 | l metallopeptidase with thrombospondin type 1 m | intron 29 | 3 | 64549044 | C | T |
| 411 | Male | LOC339975 | | intron 1 | 4 | 188264225 | AAG | A |
| 411 | male | LPCAT1 | lysophosphatidylcholine acyltransferase 1 | intron 6 | 5 | 1482348 | C | T |
| 411 | Male | FARS2 | phenylalanyl-tRNA synthetase 2: mitochondrial | intron 4 | 6 | 5465769 | G | A |
| 411 | Male | AY927641 | | intron 2 | 6 | 88581123 | A | G |
| 411 | Male | UBR5 | ubiquitin protein ligase E3 component n-recognin 5 | intron 47 | 8 | 103286470 | C | CA |
| 411 | Male | ADARB2 | enosine deaminase: RNA-specific: B2 (non-functior | intron 1 | 10 | 1769515 | C | T |
| 411 | Male | ZNF503-AS2 | ZNF503 antisense RNA 2 | intron 1 | 10 | 77163901 | A | G |
| 411 | Male | SBF2 | SET binding factor 2 | intron 16 | 11 | 9924092 | A | G |
| 411 | Male | LRRC4C | leucine rich repeat containing 4C | intron 1 | 11 | 41217824 | G | A |
| 411 | Male | PLA2G16 | HRAS-like suppressor 3 | intron 2 | 11 | 63371811 | TAAA | T |
| 411 | Male | OPCML | opioid binding protein/cell adhesion molecule-like | intron 2 | 11 | 132804548 | C | G |
| 411 | Male | PTPRO | protein tyrosine phosphatase. receptor type. O | Intron 21 | 12 | 15734228 | C | A |
| 411 | Male | DBX2 | developing brain homeobox 2 | intron 3 | 12 | 45413075 | A | G |
| 411 | Male | PTPRR | Protein Tyrosine Phosphatase Receptor Type R | intron 2 | 12 | 71250628 | GGA | G |
| 411 | Male | ANKS1B | rin repeat and sterile alpha motif domain containir | intron 11 | 12 | 99830616 | C | G |
| 411 | Male | CRY1 | cryptoome 1 (photolyase-like) | intron 1 | 12 | 107417286 | G | C |
| 411 | Male | SETD1B | SET domain containing 1B | intron 16 | 12 | 122267613 | C | T |
| 411 | Male | LINC01599 | | intron 2 | 14 | 50529288 | ATC | A |
| 411 | Male | GABRA5 | amma-aminobutyric acid (GABA) A receptor: alpha | intron 7 | 15 | 27178074 | C | G |
| 411 | Male | AK021563 | | intron 1 | 16 | 72735949 | TG | T |
| 411 | male | NLRP1 | NLR family: pyrin domain containing 1 | intron 7 | 17 | 5442308 | C | T |
| 411 | Male | RUNX1 | runt-related transcription factor 1 | intron 4 | 21 | 37165065 | C | G |
| 411 | Male | PTCHD1-AS | PTCHD1 Antisense RNA | intron 4 | X | 22444899 | GGT | G |
| 716 | Female | PPP1R12B | protein phosphatase 1: regulatory subunit 12B | intron 11 | 1 | 202411485 | G | T |
| 716 | Female | TON1-GTF2A1 | STON1-GTF2A1L readthrough | intron 10 | 2 | 48912220 | G | T |
| 716 | Female | GFPT1 | glutamine--fructose-6-phosphate transaminase 1 | intron 3 | 2 | 69591482 | G | C |
| 716 | Female | SESTD1 | SEC14 and spectrin domains 1 | intron 1 | 2 | 180104838 | A | G |
| 716 | Female | ZNF804A | zinc finger protein 804A | intron 2 | 2 | 185793971 | C | T |

**TABLE S3**

| CDH Proband Family | Tissue found in | Father | Mother | Proband Blood | Proband Skin | Proband Diaphragm | Variants with 0 reads in other tissues |
|---|---|---|---|---|---|---|---|
| 411 | All three | 0/56 | 0/50 | 46/88 | 27/58 | 41/85 | |
| 411 | All three | 0/51 | 0/54 | 16/29 | 14/30 | 26/46 | |
| 411 | All three | 0/44 | 0/50 | 37/70 | 34/66/ | 55/90 | |
| 411 | All three | 0/46 | 0/41 | 24/44 | 33/66 | 21/56 | |
| 411 | Skin+Blood | 0/31 | 0/44 | 9/44 | 21/73 | 5/54 | |
| 411 | All three | 0/52 | 0/58 | 33/70 | 30/71 | 51/83 | |
| 411 | Skin | 0/57 | 0/45 | 0/49 | 6/28 | 0/51 | Skin |
| 411 | Skin | 0/46 | 0/48 | 10/55 | 18/64 | 5/67 | |
| 411 | All three | 0/54 | 0/51 | 27/58 | 32/65 | 32/66 | |
| 411 | All three | 0/42 | 0/41 | 33/63 | 29/47 | 28/56 | |
| 411 | All three | 0/47 | 0/43 | 26/66 | 30/48 | 39/84 | |
| 411 | All three | 0/46 | 0/41 | 29/53 | 30/49 | 29/70 | |
| 411 | All three | 0/43 | 0/38 | 21/65 | 17/60 | 22/74 | |
| 411 | All three | 0/52 | 0/47 | 38/61 | 33/65 | 48/98 | |
| 411 | All three | 0/51 | 0/46 | 36/67 | 33/67 | 38/78 | |
| 411 | All three | 0/43 | 0/39 | 50/95 | 26/54 | 41/86 | |
| 411 | All three | 0/43 | 0/48 | 32/60 | 39/69 | 47/89 | |
| 411 | All three | 0/62 | 0/53 | 6/16 | 9/38 | 23/50 | |
| 411 | All three | 0/57 | 0/47 | 30/43 | 37/69 | 40/73 | |
| 411 | Blood | 0/45 | 0/56 | 8/34 | 0/53 | 0/66 | Blood |
| 411 | All three | 0/47 | 0/47 | 29/60 | 44/70 | 35/65 | |
| 411 | All three | 0/61 | 0/45 | 36/63 | 27/55 | 39/78 | |
| 411 | All three | 0/48 | 0/51 | 36/84 | 41/63 | 35/73 | |
| 411 | Skin | 0/62 | 0/42 | 0/50 | 11/52 | 0/51 | Skin |
| 411 | All three | 0/44 | 0/50 | 20/49 | 31/57 | 41/83 | |
| 411 | All three | 0/48 | 0/44 | 31/58 | 26/41 | 39/66 | |
| 411 | All three | 0/60 | 0/61 | 29/57 | 32/65 | 34/68 | |
| 411 | All three | 0/58 | 0/40 | 37/68 | 23/60 | 64/94 | |
| 411 | Skin | 3/32 | 0/59 | 0/37 | 7/33 | 0/51 | Skin |
| 716 | All three | 0/55 | 0/44 | 33/72 | 38/81 | 46/97 | |
| 716 | All three | 0/58 | 0/60 | 34/61 | 34/70 | 34/73 | |
| 716 | All three | 0/55 | 0/54 | 30/59 | 35/74 | 26/61 | |
| 716 | All three | 0/49 | 0/58 | 32/54 | 31/63 | 44/72 | |
| 716 | All three | 0/47 | 0/67 | 22/49 | 36/77 | 34/67 | |

**TABLE S3**

| CDH Proband Family | Sex | Gene | Gene full name | Location | Chromosome | Position | Reference allele | Alternate allele |
|---|---|---|---|---|---|---|---|---|
| 716 | Female | RQCD1 | 1 required for cell differentiation1 homolog (S. por | intron 1 | 2 | 219436257 | C | T |
| 716 | Female | GPR55 | G protein-coupled receptor 55 | intron 1 | 2 | 231787570 | C | A |
| 716 | Female | LINC00880 | long intergenic non-protein coding RNA 880 | intron 1 | 3 | 156823715 | T | C |
| 716 | Female | ZBBX | zinc finger B-box domain containing | intron 18 | 3 | 167012263 | G | T |
| 716 | Female | EIF4G1 | eukaryotic translation initiation factor 4 gamma: 1 | intron 7 | 3 | 184040109 | T | A |
| 716 | Female | STX18 | syntaxin 18 | intron 5 | 4 | 4457642 | T | C |
| 716 | Female | FLJ13197 | | intron 1 | 4 | 38664228 | T | C |
| 716 | Female | STPG2 | sperm-tail PG-rich repeat containing 2 | intron 10 | 4 | 98532709 | T | C |
| 716 | Female | CXXC4 | CXXC finger protein 4 | 3 utr | 4 | 105390032 | T | C |
| 716 | Female | CDH18 | cadherin 18 | intron 2 | 5 | 20088379 | T | G |
| 716 | Female | NR_073113 | | intron 1 | 5 | 43598985 | T | C |
| 716 | Female | ANKRD31 | ankyrin repeat domain 31 | intron 4 | 5 | 74504516 | T | C |
| 716 | Female | MCC | mutated in colorectal cancers | intron 1 | 5 | 112734744 | G | A |
| 716 | Female | KCNIP1 | Kv channel interacting protein 1 | intron 1 | 5 | 170030091 | GGATCCCACC | T |
| 716 | Female | ANKS1A | in repeat and sterile alpha motif domain containir | intron 1 | 6 | 34925031 | T | C |
| 716 | Female | HMGCLL1 | 3-hydroxymethyl-3-methylglutaryl-CoA lyase-like 1 | intron 2 | 6 | 55434220 | T | C |
| 716 | Female | THEMIS | thymocyte selection associated | intron 4 | 6 | 128042029 | A | G |
| 716 | Female | AIG1 | androgen-induced 1 | intron 3 | 6 | 143506521 | C | T |
| 716 | Female | GRM8 | glutamate receptor: metabotropic 8 | intron 2 | 7 | 126737917 | A | G |
| 716 | Female | NEIL2 | nei endonuclease VIII-like 2 (E. coli) | 3 utr | 8 | 11643801 | C | T |
| 716 | Female | ANGPT1 | angiopoietin 1 | intron 4 | 8 | 108328503 | T | C |
| 716 | Female | KHDRBS3 | n containing: RNA binding: signal transduction ass | intron 2 | 8 | 136543376 | G | T |
| 716 | Female | DOCK8 | dedicator of cytokinesis 8 | intron 18 | 9 | 375626 | C | T |
| 716 | Female | ERCC6L2 | omplementing rodent repair deficiency: compleme | intron 13 | 9 | 98720169 | T | A |
| 716 | Female | MIR3134 | microRNA 3134 | intron 1 | 9 | 114934634 | G | A |
| 716 | Female | ADAM12 | ADAM metallopeptidase domain 12 | intron 9 | 10 | 127787410 | A | G |
| 716 | female | NELL1 | NEL-like 1 (chicken) | intron 5 | 11 | 20910939 | A | C |
| 716 | Female | LRRC4C | leucine rich repeat containing 4C | intron 1 | 11 | 40832247 | C | T |
| 716 | Female | CD6 | CD6 molecule | intron 2 | 11 | 60774521 | G | C |
| 716 | Female | PDS5B | gulator of cohesion maintenance: homolog B (S. ce | intron 1 | 13 | 33169492 | A | G |
| 716 | Female | CLYBL | citrate lyase beta like | intron 2 | 13 | 100501587 | G | A |
| 716 | Female | abParts | | intron 7 | 15 | 22421168 | C | A |
| 716 | Female | PEAK1 | pseudopodium-enriched atypical kinase 1 | intron 1 | 15 | 77658986 | C | T |
| 716 | Female | ACSBG1 | acyl-CoA synthetase bubblegum family member 1 | intron 2 | 15 | 78499714 | T | C |

**TABLE S3**

| CDH Proband Family | Tissue found in | Father | Mother | Proband Blood | Proband Skin | Proband Diaphragm | Variants with 0 reads in other tissues |
|---|---|---|---|---|---|---|---|
| 716 | All three | 0/63 | 0/59 | 15/37 | 21/47 | 24/50 | |
| 716 | All three | 0/48 | 0/46 | 37/60 | 22/54 | 32/61 | |
| 716 | All three | 0/40 | 0/59 | 46/82 | 35/76 | 43/90 | |
| 716 | All three | 0/59 | 0/64 | 24/65 | 23/62 | 25/64 | |
| 716 | All three | 0/58 | 0/47 | 39/72 | 34/78 | 45/90 | |
| 716 | All three | 0/40 | 0/42 | 35/67 | 26/59 | 32/64 | |
| 716 | All three | 0/46 | 0/53 | 28/64 | 33/67 | 29/63 | |
| 716 | All three | 0/48 | 0/50 | 32/65 | 39/68 | 35/57 | |
| 716 | All three | 0/56 | 0/47 | 34/71 | 30/65 | 22/61 | |
| 716 | All three | 0/62 | 0/67 | 32/74 | 35/73 | 37/79 | |
| 716 | All three | 0/44 | 0/44 | 36/69 | 33/73 | 39/70 | |
| 716 | All three | 0/47 | 0/54 | 26/60 | 36/69 | 27/63 | |
| 716 | All three | 0/48 | 0/53 | 22/49 | 41/76 | 27/64 | |
| 716 | All three | 0/51 | 0/66 | 29/59 | 30/76 | 32/58 | |
| 716 | All three | 0/48 | 0/56 | 41/85 | 30/80 | 29/59 | |
| 716 | All three | 0/55 | 0/61 | 30/65 | 41/76 | 34/63 | |
| 716 | All three | 0/43 | 0/48 | 42/71 | 41/75 | 43/68 | |
| 716 | All three | 0/41 | 0/63 | 35/63 | 35/68 | 39/81 | |
| 716 | All three | 0/57 | 0/66 | 46/77 | 25/61 | 35/66 | |
| 716 | All three | 0/59 | 0/55 | 28/55 | 36/69 | 22/54 | |
| 716 | All three | 0/53 | 0/55 | 26/68 | 47/80 | 42/73 | |
| 716 | All three | 0/55 | 0/60 | 40/67 | 23/57 | 34/66 | |
| 716 | All three | 0/51 | 0/57 | 37/66 | 35/64 | 36/68 | |
| 716 | All three | 0/53 | 0/59 | 27/51 | 31/65 | 38/69 | |
| 716 | All three | 0/47 | 0/56 | 41/74 | 35/65 | 28/66 | |
| 716 | All three | 0/61 | 0/60 | 30/71 | 32/72 | 33/68 | |
| 716 | Blood | 0/64 | 0/75 | 19/94 | 4/62 | 4/71 | |
| 716 | All three | 0/49 | 0/46 | 27/61 | 34/62 | 29/57 | |
| 716 | All three | 0/53 | 0/61 | 26/53 | 26/63 | 32/68 | |
| 716 | All three | 0/54 | 0/50 | 35/80 | 24/54 | 39/66 | |
| 716 | All three | 0/52 | 0/53 | 33/59 | 23/58 | 41/73 | |
| 716 | All three | 0/122 | 0/97 | 33/107 | 37/117 | 44/116 | |
| 716 | All three | 0/50 | 0/50 | 38/63 | 37/69 | 34/58 | |
| 716 | All three | 0/66 | 0/65 | 34/56 | 41/82 | 35/69 | |

**TABLE S3**

| CDH Proband Family | Sex | Gene | Gene full name | Location | Chromosome | Position | Reference allele | Alternate allele |
|---|---|---|---|---|---|---|---|---|
| 716 | Female | SLCO3A1 | e carrier organic anion transporter family: membe | intron 5 | 15 | 92664705 | T | C |
| 716 | female | LRRK1 | leucine-rich repeat kinase 1 | 3 utr | 15 | 101611011 | A | T |
| 716 | Female | PRKCB | protein kinase C: beta | intron 3 | 16 | 24033310 | C | A |
| 716 | Female | CKM | creatine kinase: muscle | intron 4 | 19 | 45818459 | C | T |
| 716 | Female | ZNF256 | zinc finger protein 256 | intron 1 | 19 | 58458152 | T | C |
| 716 | Female | SIRPA | signal-regulatory protein alpha | intron 2 | 20 | 1890924 | T | G |
| 716 | Female | PTPRT | protein tyrosine phosphatase: receptor type: T | intron 2 | 20 | 41496654 | A | G |
| 716 | Female | APCDD1L | adenomatosis polyposis coli down-regulated 1-like | intron 1 | 20 | 57048292 | C | CT |
| 716 | Female | SYCP2 | synaptonemal complex protein 2 | intron 11 | 20 | 58488059 | TCA | T |
| 716 | Female | SGSM1 | small G protein signaling modulator 1 | intron 2 | 22 | 25212931 | T | C |
| 716 | Female | TBC1D22A | TBC1 domain family: member 22A | intron 12 | 22 | 47565909 | T | C |
| 716 | Female | FRMPD4 | FERM And PDZ Domain Containing 4 | intron 4 | X | 12663028 | G | A |
| 716 | Female | HEPH | hephaestin | intron 2 | X | 65391234 | T | C |
| 809 | Female | SORCS3 | ortilin-related VPS10 domain containing receptor | intron 5 | 1 | 10687743 | C | T |
| 809 | Female | CHIA | chitinase: acidic | intron 2 | 1 | 111854262 | C | T |
| 809 | Female | SYCP1 | Synaptonemal Complex Protein 1 | intron 27 | 1 | 115498719 | C | T |
| 809 | Female | BRINP3 | phogenetic protein/retinoic acid inducible neural- | intron 6 | 1 | 190140359 | T | A |
| 809 | Female | TRAPPC12 | Trafficking Protein Particle Complex 12 | intron 10 | 2 | 3485756 | C | T |
| 809 | Female | ALK | anaplastic lymphoma receptor tyrosine kinase | intron 5 | 2 | 29563357 | C | T |
| 809 | Female | FLJ30838 | | intron 2 | 2 | 59058824 | G | A |
| 809 | Female | AFTPH | aftiphilin | intron 7 | 2 | 64808306 | C | T |
| 809 | Female | SLC4A5 | rier family 4 (sodium bicarbonate cotransporter): | intron 8 | 2 | 74496034 | C | A |
| 809 | Female | SFTPB | surfactant protein B | 3 utr | 2 | 85884898 | T | G |
| 809 | Female | BIN1 | Bridging integrator 1 | intron 1 | 2 | 127840765 | G | A |
| 809 | Female | PLA2R1 | phospholipase A2 receptor 1: 180kDa | intron 11 | 2 | 160851223 | G | A |
| 809 | Female | BOLL | bol: boule-like (Drosophila) | intron 2 | 2 | 198644842 | C | A |
| 809 | Female | LINC00607 | long intergenic non-protein coding RNA 607 | intron 5 | 2 | 216549095 | G | A |
| 809 | Female | ULK4 | unc-51 like kinase 4 | intron 25 | 3 | 41755470 | CTG | C |
| 809 | Female | FHIT | fragile histidine triad | intron 5 | 3 | 60152652 | G | T |
| 809 | Female | PPP2R3A | protein phosphatase 2: regulatory subunit B'': alpha | intron 6 | 3 | 135794102 | C | A |
| 809 | Female | SLC9A9 | mily 9: subfamily A (NHE9: cation proton antiporte | intron 4 | 3 | 143433614 | T | C |
| 809 | Female | VEPH1 | ventricular zone expressed PH domain-containing 1 | intron 9 | 3 | 157074940 | A | G |
| 809 | Female | NLGN1 | Neuroligin 1 | intron 4 | 3 | 173983838 | T | A |
| 809 | Female | PEX5L | peroxisomal biogenesis factor 5-like | intron 1 | 3 | 179719642 | C | T |

De Novo UTR+Intron All

**TABLE S3**

| CDH Proband Family | Tissue found in | Father | Mother | Proband Blood | Proband Skin | Proband Diaphragm | Variants with 0 reads in other tissues |
|---|---|---|---|---|---|---|---|
| 716 | All three | 0/40 | 0/46 | 39/74 | 35/76 | 41/73 | |
| 716 | All three | 0/64 | 0/53 | 25/55 | 31/66 | 23/65 | |
| 716 | All three | 0/39 | 0/62 | 25/68 | 35/76 | 33/62 | |
| 716 | All three | 0/58 | 0/52 | 30/66 | 34/85 | 36/61 | |
| 716 | All three | 0/50 | 0/55 | 42/85 | 45/87 | 30/72 | |
| 716 | All three | 0/47 | 0/53 | 34/56 | 31/59 | 37/76 | |
| 716 | All three | 0/55 | 0/50 | 31/62 | 42/82 | 32/66 | |
| 716 | All three | 0/58 | 0/46 | 33/58 | 38/76 | 36/56 | |
| 716 | All three | 0/47 | 0/57 | 38/69 | 39/71 | 32/57 | |
| 716 | All three | 0/62 | 0/57 | 39/79 | 39/69 | 34/67 | |
| 716 | All three | 0/54 | 0/47 | 39/67 | 32/61 | 33/68 | |
| 716 | Skin | 0/36 | 0/56 | 0/54 | 17/59 | 0/47 | Skin |
| 716 | All three | 0/36 | 0/54 | 30/61 | 34/75 | 22/51 | |
| 809 | All three | 0/43 | 0/47 | 44/68 | 30/59 | 36/75 | |
| 809 | Skin | 0/53 | 0/73 | 0/63 | 19/61 | 0/61 | Skin |
| 809 | Diaphragm | 0/50 | 0/53 | 0/65 | 0/60 | 19/94 | Diaphragm |
| 809 | Diaphragm | 0/53 | 0/64 | 0/58 | 0/52 | 26/80 | Diaphragm |
| 809 | Diaphragm | 0/41 | 0/57 | 0/43 | 0/51 | 12/60 | Diaphragm |
| 809 | Skin | 0/54 | 0/64 | 0/59 | 19/65 | 0/53 | Skin |
| 809 | Skin | 0/49 | 0/73 | 0/58 | 16/53 | 0/74 | Skin |
| 809 | All three | 0/42 | 0/55 | 30/68 | 37/72 | 47/87 | |
| 809 | Diaphragm | 0/47 | 0/57 | 0/77 | 0/65 | 26/63 | Diaphragm |
| 809 | All three | 0/46 | 0/61 | 29/58 | 29/59 | 38/73 | |
| 809 | Diaphragm | 0/43 | 0/67 | 0/64 | 0/58 | 10/49 | Diaphragm |
| 809 | All three | 0/45 | 0/65 | 39/67 | 37/85 | 44/85 | |
| 809 | Diaphragm | 0/38 | 0/69 | 0/55 | 0/38 | 20/70 | Diaphragm |
| 809 | All three | 0/50 | 0/64 | 38/85 | 40/69 | 30/66 | |
| 809 | Diaphragm | 0/45 | 0/68 | 0/55 | 0/53 | 22/87 | Diaphragm |
| 809 | Diaphragm | 0/42 | 0/53 | 0/61 | 0/56 | 27/78 | Diaphragm |
| 809 | Diaphragm | 0/39 | 0/59 | 0/44 | 0/51 | 16/71 | Diaphragm |
| 809 | All three | 0/47 | 0/57 | 38/70 | 32/59 | 36/72 | |
| 809 | Skin | 0/39 | 0/50 | 0/52 | 22/72 | 0/57 | Skin |
| 809 | Diaphragm | 0/42 | 0/51 | 0/50 | 0/51 | 13/42 | Diaphragm |
| 809 | Diaphragm | 0/46 | 0/55 | 0/51 | 0/54 | 21/58 | Diaphragm |

**TABLE S3**

| CDH Proband Family | Sex | Gene | Gene full name | Location | Chromosome | Position | Reference allele | Alternate allele |
|---|---|---|---|---|---|---|---|---|
| 809 | Female | MB21D2 | Mab-21 domain containing 2 | intron 1 | 3 | 192557614 | G | A |
| 809 | female | LMLN | leishmanolysin-like (metallopeptidase M8 family) | 5 utr | 3 | 197765899 | C | A |
| 809 | Female | ZFYVE28 | zinc finger: FYVE domain containing 28 | intron 3 | 4 | 2341715 | C | T |
| 809 | Female | STK32B | serine/threonine kinase 32B | intron 3 | 4 | 5257751 | C | G |
| 809 | Female | KCTD8 | :assium channel tetramerization domain containin | intron 1 | 4 | 44359274 | G | A |
| 809 | Female | FSTL5 | follistatin-like 5 | intron 2 | 4 | 163016217 | G | T |
| 809 | Female | ENPP6 | :tonucleotide pyrophosphatase/phosphodiesterase | intron 1 | 4 | 185077386 | G | A |
| 809 | Female | LOC285692 | | intron 4 | 5 | 9878319 | G | A |
| 809 | female | EDIL3 | EGF-like repeats and discoidin I-like domains 3 | intron 6 | 5 | 83397673 | G | A |
| 809 | Female | GPR98 | G protein-coupled receptor 98 | intron 59 | 5 | 90064793 | C | A |
| 809 | Female | FBXL17 | F-box and leucine-rich repeat protein 17 | intron 6 | 5 | 107485066 | G | A |
| 809 | Female | GRAMD3 | GRAM domain containing 3 | intron 12 | 5 | 125823743 | T | C |
| 809 | Female | TRPC7 | t receptor potential cation channel: subfamily C: m | intron 4 | 5 | 135592775 | C | T |
| 809 | Female | KCTD16 | assium channel tetramerization domain containing | intron 2 | 5 | 143580957 | T | G |
| 809 | Female | STK32A | serine/threonine kinase 32A | intron 4 | 5 | 146670218 | A | T |
| 809 | Female | CAMK2A | ılcium/calmodulin-dependent protein kinase II alpl | intron 7 | 5 | 149632443 | G | A |
| 809 | Female | EBF1 | early B-cell factor 1 | intron 9 | 5 | 158210489 | A | G |
| 809 | Female | TDP2 | tyrosyl-DNA phosphodiesterase 2 | intron 3 | 6 | 24658388 | T | A |
| 809 | Female | EYS | eyes shut homolog (Drosophila) | intron 12 | 6 | 65845477 | G | T |
| 809 | Female | COL12A1 | collagen: type XII: alpha 1 | intron 45 | 6 | 75829061 | C | T |
| 809 | Female | PREP | prolyl endopeptidase | intron 10 | 6 | 105750309 | T | C |
| 809 | Female | FAM184A | family with sequence similarity 184: member A | intron 1 | 6 | 119395836 | C | G |
| 809 | Female | TARID | F21 Antisense RNA Inducing Promoter Demethylat | intron 5 | 6 | 133975860 | C | A |
| 809 | Female | TAB2 | -Beta Activated Kinase 1 (MAP3K7) Binding Protei | intron 1 | 6 | 149608630 | A | T |
| 809 | Female | STEAP1B | STEAP family member 1B | intron 5 | 7 | 22475570 | TG | T |
| 809 | Female | ELMO1 | engulfment and cell motility 1 | intron 2 | 7 | 37358037 | A | T |
| 809 | Female | CACNA2D1 | ım channel: voltage-dependent: alpha 2/delta subı | intron 3 | 7 | 81834306 | G | A |
| 809 | Female | SEMA3A | ınoglobulin domain (Ig): short basic domain: secret | intron 2 | 7 | 83763857 | T | C |
| 809 | Female | DGKI | diacylglycerol kinase: iota | intron 28 | 7 | 137143548 | G | T |
| 809 | Female | CNTNAP2 | Contactin Associated Protein Like 2 | intron 1 | 7 | 146201971 | T | C |
| 809 | Female | KMT2C | lysine (K)-specific methyltransferase 2C | intron 40 | 7 | 151866844 | C | T |
| 809 | Female | CSMD1 | CUB And Sushi Multiple Domains 1 | intron 27 | 8 | 3014345 | C | T |
| 809 | Female | DLC1 | deleted in liver cancer 1 | intron 2 | 8 | 13287693 | C | T |
| 809 | Female | ANK1 | ankyrin 1: erythrocytic | intron 1 | 8 | 41676846 | C | A |

**TABLE S3**

| CDH Proband Family | Tissue found in | Father | Mother | Proband Blood | Proband Skin | Proband Diaphragm | Variants with 0 reads in other tissues |
|---|---|---|---|---|---|---|---|
| 809 | Diaphragm | 0/47 | 0/63 | 0/52 | 0/49 | 19/72 | Diaphragm |
| 809 | Diaphragm | 0/78 | 0/72 | 0/75 | 0/63 | 33/95 | Diaphragm |
| 809 | Skin | 0/36 | 0/42 | 0/43 | 19/62 | 0/48 | Skin |
| 809 | Diaphragm | 0/38 | 0/57 | 0/59 | 0/47 | 19/67 | Diaphragm |
| 809 | All three | 0/52 | 0/66 | 45/77 | 35/83 | 32/80 | |
| 809 | Skin | 0/43 | 0/59 | 0/55 | 27/73 | 0/61 | Skin |
| 809 | Diaphragm | 0/48 | 0/56 | 0/45 | 0/57 | 22/79 | Diaphragm |
| 809 | Skin | 0/47 | 0/68 | 0/62 | 18/65 | 0/57 | Skin |
| 809 | All three | 0/62 | 0/81 | 37/96 | 37/82 | 29/57 | |
| 809 | Skin | 0/43 | 0/58 | 0/64 | 15/64 | 0/60 | Skin |
| 809 | All three | 0/41 | 0/62 | 30/74 | 41/76 | 40/82 | |
| 809 | All three | 0/51 | 0/61 | 41/64 | 35/64 | 35/88 | |
| 809 | Diaphragm | 0/37 | 0/57 | 0/54 | 0/49 | 23/77 | Diaphragm |
| 809 | Skin | 0/45 | 0/66 | 0/59 | 30/75 | 0/55 | Skin |
| 809 | Diaphragm | 0/45 | 0/75 | 0/74 | 0/79 | 21/88 | Diaphragm |
| 809 | Skin | 0/49 | 0/52 | 0/55 | 21/76 | 0/53 | Skin |
| 809 | All three | 0/42 | 0/53 | 37/78 | 32/74 | 49/88 | |
| 809 | All three | 0/44 | 0/58 | 40/62 | 34/76 | 31/71 | |
| 809 | All three | 0/51 | 0/76 | 37/83 | 28/59 | 36/81 | |
| 809 | All three | 0/44 | 0/59 | 37/89 | 33/65 | 33/68 | |
| 809 | All three | 0/47 | 0/56 | 37/77 | 34/60 | 31/76 | |
| 809 | Skin+Blood | 0/55 | 0/68 | 19/79 | 25/81 | 11/77 | |
| 809 | Skin | 0/54 | 0/56 | 0/58 | 29/84 | 0/57 | Skin |
| 809 | Diaphragm | 0/56 | 0/60 | 0/76 | 0/65 | 11/47 | Diaphragm |
| 809 | All three | 0/45 | 0/34 | 27/58 | 46/81 | 39/71 | |
| 809 | Skin | 0/38 | 0/66 | 0/52 | 20/71 | 0/57 | Skin |
| 809 | All three | 0/54 | 0/65 | 28/56 | 32/69 | 25/55 | |
| 809 | Diaphragm | 0/43 | 0/56 | 0/57 | 0/45 | 21/85 | Diaphragm |
| 809 | Skin | 0/43 | 0/51 | 0/51 | 15/72 | 0/60 | Skin |
| 809 | Diaphragm | 0/49 | 0/60 | 0/58 | 0/46 | 17/78 | Diaphragm |
| 809 | Skin | 0/42 | 0/53 | 0/48 | 23/77 | 0/53 | Skin |
| 809 | Diaphragm | 0/44 | 0/48 | 0/56 | 0/41 | 14/58 | Diaphragm |
| 809 | Skin | 0/38 | 0/73 | 0/63 | 21/70 | 0/68 | Skin |
| 809 | Skin | 0/48 | 0/52 | 0/54 | 23/72 | 0/61 | Skin |

**TABLE S3**

| CDH Proband Family | Sex | Gene | Gene full name | Location | Chromosome | Position | Reference allele | Alternate allele |
|---|---|---|---|---|---|---|---|---|
| 809 | Female | NECAB1 | N-Terminal EF-Hand Calcium Binding Protein 1 | intron 4 | 8 | 91890081 | T | C |
| 809 | Female | TRAPPC9 | Trafficking Protein Particle Complex 9 | intron 19 | 8 | 140943579 | G | A |
| 809 | Female | PTPRD | protein tyrosine phosphatase: receptor type: D | intron 8 | 9 | 9448343 | G | T |
| 809 | Female | PTPRD | protein tyrosine phosphatase: receptor type: D | intron 7 | 9 | 9600668 | G | A |
| 809 | Female | PTPRD | Protein Tyrosine Phosphatase Receptor Type D | intron 5 | 9 | 9830793 | G | A |
| 809 | Female | PTPRD | protein tyrosine phosphatase: receptor type: D | intron 3 | 9 | 10048095 | A | C |
| 809 | Female | LINGO2 | leucine rich repeat and Ig domain containing 2 | intron 5 | 9 | 28265953 | G | T |
| 809 | Female | TRPM3 | receptor potential cation channel: subfamily M: m | intron 2 | 9 | 73481507 | A | G |
| 809 | Female | ROR2 | receptor tyrosine kinase-like orphan receptor 2 | intron 1 | 9 | 94578709 | C | A |
| 809 | Female | UGCG | UDP-glucose ceramide glucosyltransferase | intron 1 | 9 | 114666708 | C | T |
| 809 | Female | DBC1 | cell cycle and apoptosis regulator 2 | intron 7 | 9 | 121956986 | A | T |
| 809 | Female | ZCCHC24 | zinc finger: CCHC domain containing 24 | intron 2 | 10 | 81168284 | C | T |
| 809 | Female | SLC16A12 | Solute Carrier Family 16 Member 12 | intron 3 | 10 | 91220525 | A | G |
| 809 | Female | TACC2 | ransforming: acidic coiled-coil containing protein 2 | intron 5 | 10 | 123855269 | C | T |
| 809 | Female | ST5 | suppression of tumorigenicity 5 | intron 1 | 11 | 8822290 | T | G |
| 809 | Female | SOX6 | SRY (sex determining region Y)-box 6 | intron 12 | 11 | 16061906 | A | T |
| 809 | Female | LRRC4C | leucine rich repeat containing 4C | intron 1 | 11 | 40749101 | AT | A |
| 809 | Female | DLG2 | 11 83978757 | intron 1 | 11 | 83978757 | TC | T |
| 809 | Female | MAML2 | mastermind-like 2 (Drosophila) | intron 2 | 11 | 95731095 | C | CAT |
| 809 | Female | ALG9 | ALG9: alpha-1:2-mannosyltransferase | intron 9 | 11 | 111723240 | G | A |
| 809 | Female | KIRREL3 | kin of IRRE like 3 (Drosophila) | intron 1 | 11 | 126775175 | G | A |
| 809 | Female | GLB1L2 | galactosidase: beta 1-like 2 | intron 6 | 11 | 134227263 | A | T |
| 809 | Female | BCAT1 | ranched chain amino-acid transaminase 1. cytosoli | intron 4 | 12 | 25029237 | G | C |
| 809 | Female | NELL2 | NEL-like 2 (chicken) | intron 16 | 12 | 44969449 | G | A |
| 809 | Female | BTBD11 | BTB (POZ) domain containing 11 | intron 1 | 12 | 107743246 | C | A |
| 809 | Female | ATP2A2 | Pase: Ca++ transporting: cardiac muscle: slow twitc | intron 5 | 12 | 110734883 | C | T |
| 809 | Female | ZMYM2 | Zinc Finger MYM-Type Containing 2 | intron 15 | 13 | 20632248 | G | C |
| 809 | Female | NPAS3 | neuronal PAS domain protein 3 | intron 3 | 14 | 33833460 | A | G |
| 809 | Female | GABRB3 | amma-aminobutyric acid (GABA) A receptor: beta | intron 3 | 15 | 26992702 | T | C |
| 809 | Female | IQGAP1 | IQ motif containing GTPase activating protein 1 | intron 15 | 15 | 91003597 | C | T |
| 809 | female | SLC7A6 | mily 7 (amino acid transporter light chain. y+L syst | intron 4 | 16 | 68311346 | G | C |
| 809 | Female | RAB11FIP4 | RAB11 family interacting protein 4 (class II) | intron 3 | 17 | 29765052 | G | A |
| 809 | Female | SLC9A3R1 | 9: subfamily A (NHE3: cation proton antiporter 3): r | intron 1 | 17 | 72753883 | C | T |
| 809 | Female | ANKRD30B | Ankyrin Repeat Domain 30B | intron 3 | 18 | 14753321 | T | C |

**TABLE S3**

| CDH Proband Family | Tissue found in | Father | Mother | Proband Blood | Proband Skin | Proband Diaphragm | Variants with 0 reads in other tissues |
|---|---|---|---|---|---|---|---|
| 809 | Diaphragm | 0/54 | 0/73 | 0/64 | 0/65 | 18/73 | Diaphragm |
| 809 | Diaphragm | 0/46 | 0/57 | 0/44 | 0/45 | 14/63 | Diaphragm |
| 809 | All three | 0/47 | 0/65 | 39/66 | 40/63 | 40/77 | |
| 809 | All three | 0/44 | 0/67 | 45/76 | 41/79 | 35/66 | |
| 809 | Diaphragm | 0/53 | 0/59 | 0/54 | 0/49 | 18/85 | Diaphragm |
| 809 | Skin | 0/50 | 0/58 | 0/68 | 29/86 | 0/66 | Skin |
| 809 | Diaphragm | 0/40 | 0/60 | 0/68 | 0/56 | 27/79 | Diaphragm |
| 809 | Skin | 0/50 | 0/57 | 0/63 | 20/81 | 0/57 | Skin |
| 809 | Diaphragm | 0/39 | 0/48 | 0/47 | 0/44 | 21/73 | Diaphragm |
| 809 | All three | 0/37 | 0/54 | 47/87 | 37/76 | 38/63 | |
| 809 | Diaphragm | 0/42 | 0/54 | 0/52 | 0/47 | 24/77 | Diaphragm |
| 809 | Diaphragm | 0/45 | 0/53 | 0/67 | 0/65 | 23/75 | Diaphragm |
| 809 | Skin | 0/40 | 0/73 | 0/48 | 7/26 | 0/68 | Skin |
| 809 | Diaphragm | 0/38 | 0/67 | 0/64 | 0/63 | 23/75 | Diaphragm |
| 809 | All three | 0/50 | 0/54 | 32/73 | 43/78 | 38/79 | |
| 809 | Skin | 0/36 | 0/53 | 0/55 | 21/75 | 0/43 | Skin |
| 809 | All three | 0/38 | 0/61 | 49/85 | 30/59 | 43/69 | |
| 809 | Skin | 0/51 | 0/73 | 0/69 | 18/74 | 0/65 | Skin |
| 809 | Diaphragm | 0/40 | 0/73 | 0/59 | 0/53 | 21/75 | Diaphragm |
| 809 | All three | 0/52 | 0/55 | 30/58 | 32/60 | 37/71 | |
| 809 | All three | 0/60 | 0/64 | 31/61 | 35/67 | 36/73 | |
| 809 | Skin | 0/45 | 0/63 | 0/52 | 24/71 | 0/55 | Skin |
| 809 | Skin | 0/45 | 0/59 | 0/50 | 10/32 | 0/67 | Skin |
| 809 | Skin | 0/48 | 0/63 | 0/62 | 22/73 | 0/56 | Skin |
| 809 | Diaphragm | 0/55 | 0/65 | 0/63 | 0/63 | 16/64 | Diaphragm |
| 809 | All three | 0/42 | 0/69 | 26/67 | 37/67 | 36/60 | |
| 809 | Diaphragm | 0/46 | 0/65 | 0/55 | 0/47 | 13/64 | Diaphragm |
| 809 | All three | 0/42 | 0/52 | 39/71 | 27/67 | 30/60 | |
| 809 | Skin | 0/51 | 0/66 | 0/61 | 19/61 | 0/65 | Skin |
| 809 | Skin | 0/42 | 0/60 | 0/49 | 18/64 | 0/53 | Skin |
| 809 | Diaphragm | 0/51 | 0/72 | 0/73 | 0/60 | 23/68 | Diaphragm |
| 809 | Skin | 0/39 | 0/48 | 0/46 | 19/65 | 0/53 | Skin |
| 809 | Diaphragm | 0/41 | 0/50 | 0/51 | 0/54 | 21/72 | Diaphragm |
| 809 | Diaphragm | 0/51 | 0/66 | 0/61 | 0/58 | 13/61 | Diaphragm |

**TABLE S3**

De Novo UTR+Intron All

| CDH Proband Family | Sex | Gene | Gene full name | Location | Chromosome | Position | Reference allele | Alternate allele |
|---|---|---|---|---|---|---|---|---|
| 809 | Female | LINC00907 | long intergenic non-protein coding RNA 907 | intron 4 | 18 | 39871960 | C | A |
| 809 | Female | DCC | deleted in colorectal carcinoma | intron 4 | 18 | 50450951 | G | T |
| 809 | Female | DCC | deleted in colorectal carcinoma | intron 5 | 18 | 50470480 | T | C |
| 809 | Female | ZNF812P | 19 9808127 | intron 1 | 19 | 9808127 | T | A |
| 809 | Female | GAPDHS | ;raldehyde-3-phosphate dehydrogenase: spermato; | intron 9 | 19 | 36035048 | G | A |
| 809 | Female | PRKD2 | Protein Kinase D2 | intron 2 | 19 | 47216261 | T | TTTTCTTTTT( |
| 809 | Female | NR_040084 | | intron 1 | 21 | 37474586 | A | C |
| 809 | Female | CNKSR2 | Connector Enhancer Of Kinase Suppressor Of Ras 2 | intron 4 | X | 21476844 | T | G |
| 809 | Female | KLHL4 | kelch-like family member 4 | intron 1 | X | 86862155 | C | A |
| 809 | Female | ZCCHC16 | zinc finger: CCHC domain containing 16 | intron 2 | X | 111601842 | C | T |
| 967 | Male | SPATA6 | spermatogenesis associated 6 | intron 11 | 1 | 48785688 | G | A |
| 967 | Male | LRRC8C | leucine rich repeat containing 8 family: member C | intron 1 | 1 | 90115215 | C | T |
| 967 | Male | ZNF695 | Zinc Finger Protein 695 | intron 5 | 1 | 247110379 | AAAGAGTAA/ | T |
| 967 | Male | MERTK | c-mer proto-oncogene tyrosine kinase | intron 8 | 2 | 112744068 | C | T |
| 967 | Male | THSD7B | thrombospondin: type I: domain containing 7B | intron 13 | 2 | 138202217 | CCT | C |
| 967 | Male | SCHIP1 | Schwannomin Interacting Protein 1 | intron 2 | 3 | 159529771 | T | TTG |
| 967 | Male | NAALADL2 | N-acetylated alpha-linked acidic dipeptidase-like 2 | intron 9 | 3 | 175240124 | C | T |
| 967 | Male | BC005018 | | intron 4 | 4 | 84733093 | C | T |
| 967 | Male | PDZD2 | PDZ domain containing 2 | intron 2 | 5 | 31946007 | T | C |
| 967 | Male | DQ515898 | | intron 1 | 8 | 128336350 | T | C |
| 967 | Male | ANTD3-TMEI | MSANTD3-TMEFF1 readthrough | intron 1 | 9 | 103248140 | T | G |
| 967 | Male | NELFB | negative elongation factor complex member B | intron 4 | 9 | 140151750 | C | T |
| 967 | Male | RSU1 | Ras suppressor protein 1 | intron 1 | 10 | 16830733 | G | A |
| 967 | Male | SUFU | suppressor of fused homolog (Drosophila) | intron 2 | 10 | 104283460 | G | A |
| 967 | Male | OR8U8 | olfactory receptor family 8 subfamily U member 8 | intron 1 | 11 | 56202375 | A | G |
| 967 | male | DISC1FP1 | DISC1 fusion partner 1 (non-protein coding) | intron 1 | 11 | 90243703 | TATGC | T |
| 967 | Male | SPATA13 | Spermatogenesis Associated 13 | intron 3 | 13 | 24682941 | A | C |
| 967 | Male | UGGT2 | UDP-glucose glycoprotein glucosyltransferase 2 | intron 4 | 13 | 96673108 | A | G |
| 967 | Male | MDGA2 | domain containing glycosylphosphatidylinositol an | intron 1 | 14 | 47897999 | T | C |
| 967 | Male | nCG_2003567 | | intron 1 | 15 | 66926125 | A | C |
| 967 | Male | CD33 | CD33 molecule | intron 7 | 19 | 51739471 | G | T |
| 967 | Male | TBX1 | T-box 1 | intron 8 | 22 | 19768601 | G | A |
| 967 | male | TENM1 | teneurin transmembrane protein 1 | intron 1 | X | 124078891 | A | C |

**TABLE S3**

| CDH Proband Family | Tissue found in | Father | Mother | Proband Blood | Proband Skin | Proband Diaphragm | Variants with 0 reads in other tissues |
|---|---|---|---|---|---|---|---|
| 809 | Diaphragm | 0/41 | 0/65 | 0/72 | 0/55 | 21/78 | Diaphragm |
| 809 | Diaphragm | 0/40 | 0/65 | 0/60 | 0/53 | 27/71 | Diaphragm |
| 809 | Skin | 0/46 | 0/55 | 0/50 | 22/71 | 0/55 | Skin |
| 809 | Diaphragm | 0/39 | 0/52 | 0/51 | 0/44 | 27/68 | Diaphragm |
| 809 | Skin | 0/41 | 0/56 | 0/51 | 19/66 | 0/62 | Skin |
| 809 | All three | 0/48 | 0/69 | 15/30 | 13/36 | 29/43 | |
| 809 | Skin | 0/47 | 0/67 | 0/57 | 21/93 | 0/74 | Skin |
| 809 | Diaphragm | 0/18 | 0/58 | 0/52 | 0/46 | 17/66 | Diaphragm |
| 809 | Diaphragm | 0/35 | 0/74 | 0/61 | 0/59 | 26/69 | Diaphragm |
| 809 | Skin | 0/31 | 0/63 | 0/49 | 24/65 | 0/54 | Skin |
| 967 | All three | 0/47 | 0/75 | 18/38 | 25/55 | 27/48 | |
| 967 | All three | 0/47 | 0/58 | 29/52 | 23/54 | 35/52 | |
| 967 | Skin | 0/55 | 5/66 | 0/44 | 23/111 | 0/52 | Skin |
| 967 | All three | 0/69 | 0/60 | 28/67 | 38/78 | 16/48 | |
| 967 | All three | 0/54 | 0/55 | 21/40 | 25/65 | 14/30 | |
| 967 | All three | 0/56 | 0/69 | 26/40 | 35/59 | 19/28 | |
| 967 | All three | 0/47 | 0/62 | 33/63 | 22/67 | 24/57 | |
| 967 | All three | 0/42 | 0/53 | 27/49 | 25/55 | 24/45 | |
| 967 | All three | 0/43 | 0/43 | 22/49 | 26/53 | 29/47 | |
| 967 | All three | 0/52 | 0/64 | 24/52 | 27/66 | 13/36 | |
| 967 | All three | 0/54 | 0/59 | 23/43 | 39/65 | 16/46 | |
| 967 | All three | 0/54 | 0/44 | 26/57 | 31/69 | 20/40 | |
| 967 | All three | 0/44 | 0/58 | 35/64 | 36/65 | 22/48 | |
| 967 | All three | 0/53 | 0/57 | 20/43 | 26/64 | 24/50 | |
| 967 | All three | 0/42 | 0/49 | 23/49 | 34/65 | 12/36 | |
| 967 | All three | 1/25 | 0/43 | 12/22 | 10/24 | 13/16 | |
| 967 | Diaphragm | 0/46 | 0/58 | 0/41 | 0/60 | 6/22 | Diaphragm |
| 967 | All three | 0/50 | 0/59 | 42/68 | 22/58 | 32/60 | |
| 967 | All three | 0/62 | 0/52 | 31/60 | 30/59 | 22/47 | |
| 967 | All three | 0/42 | 0/54 | 14/36 | 36/71 | 20/45 | |
| 967 | All three | 0/50 | 0/62 | 20/49 | 38/61 | 18/47 | |
| 967 | All three | 0/55 | 0/50 | 20/40 | 28/54 | 20/41 | |
| 967 | Diaphragm | 0/36 | 0/78 | 2/20 | 3/24 | 10/32 | |

**TABLE S3**

| CDH Proband Family | Sex | Location | Chromosome | Position | Reference allele | Alternate allele | Tissue found in | Father |
|---|---|---|---|---|---|---|---|---|
| 411 | Male | Intergenic | 2 | 6448411 | G | A | All three | 0/51 |
| 411 | Male | Intergenic | 2 | 15979430 | C | T | All three | 0/46 |
| 411 | Male | Intergenic | 2 | 23607772 | C | A | All three | 0/34 |
| 411 | Male | Intergenic | 2 | 23607823 | G | A | All three | 0/34 |
| 411 | Male | Intergenic | 2 | 48488807 | CTTTGAAAG | T | All three | 0/55 |
| 411 | male | Intergenic | 4 | 12542452 | ACAAAAAT | A | All three | 0/61 |
| 411 | Male | Intergenic | 4 | 40268400 | A | G | All three | 0/59 |
| 411 | Male | Intergenic | 4 | 130485536 | C | T | All three | 0/39 |
| 411 | Male | Intergenic | 4 | 135941340 | ATAATACATG | T | Skin | 0/36 |
| 411 | Male | Intergenic | 5 | 30972421 | G | T | All three | 0/54 |
| 411 | Male | Intergenic | 5 | 43876751 | A | T | All three | 0/43 |
| 411 | Male | Intergenic | 7 | 63891470 | TTGGCCGGG | A | Blood | 0/44 |
| 411 | Male | Intergenic | 7 | 121415957 | T | G | All three | 0/42 |
| 411 | Male | Intergenic | 8 | 49656228 | G | C | All three | 0/45 |
| 411 | Male | Intergenic | 9 | 24573667 | A | C | All three | 0/57 |
| 411 | Male | Intergenic | 10 | 45696971 | C | T | All three | 0/44 |
| 411 | male | Intergenic | 10 | 107240148 | T | C | All three | 0/54 |
| 411 | Male | Intergenic | 11 | 48522238 | T | C | All three | 0/53 |
| 411 | Male | Intergenic | 11 | 68646620 | CCT | C | Diaphragm | 0/44 |
| 411 | Male | Intergenic | 12 | 51286910 | T | C | All three | 0/46 |
| 411 | Male | Intergenic | 13 | 63562946 | G | GA | All three | 0/63 |
| 411 | Male | Intergenic | 15 | 38941044 | C | T | All three | 0/45 |
| 411 | Male | Intergenic | 15 | 46407038 | C | T | All three | 0/55 |
| 411 | Male | Intergenic | 16 | 75543055 | C | A | All three | 0/46 |
| 411 | Male | Intergenic | 17 | 404337 | C | T | All three | 0/55 |
| 411 | Male | Intergenic | 20 | 5206920 | C | A | All three | 0/45 |
| 411 | Male | Intergenic | 20 | 48832820 | CAG | C | Skin | 0/45 |
| 411 | Male | Intergenic | X | 3355438 | CTG | C | Skin | 0/38 |
| 411 | Male | Intergenic | X | 50778530 | ACATTTCTCA | G | Diaphragm | 0/18 |
| 411 | Male | Intergenic | X | 117269505 | CGT | C | Diaphragm | 0/43 |

**TABLE S3**

| CDH Proband Family | Mother | Proband Blood | Proband Skin | Proband Diaphragm | Variants with 0 reads in other tissues |
|---|---|---|---|---|---|
| 411 | 0/49 | 29/59 | 21/54 | 51/89 | |
| 411 | 0/38 | 16/47 | 23/50 | 36/63 | |
| 411 | 0/39 | 16/26 | 25/40 | 23/56 | |
| 411 | 0/47 | 16/21 | 17/25 | 17/49 | |
| 411 | 0/46 | 23/50 | 22/60 | 32/70 | |
| 411 | 0/67 | 23/54 | 23/50 | 32/86 | |
| 411 | 0/56 | 36/75 | 30/55 | 38/78 | |
| 411 | 0/55 | 34/67 | 23/58 | 36/75 | |
| 411 | 0/46 | 0/49 | 19/86 | 0/40 | Skin |
| 411 | 0/52 | 28/65 | 34/65 | 41/99 | |
| 411 | 0/43 | 37/78 | 32/59 | 21/55 | |
| 411 | 0/35 | 17/67 | 0/57 | 4/82 | |
| 411 | 0/50 | 25/62 | 26/61 | 42/81 | |
| 411 | 0/47 | 31/57 | 21/58 | 42/75 | |
| 411 | 0/45 | 24/69 | 18/48 | 32/78 | |
| 411 | 0/59 | 33/70 | 31/65 | 29/60 | |
| 411 | 0/58 | 32/79 | 25/58 | 33/71 | |
| 411 | 0/41 | 26/61 | 24/69 | 35/85 | |
| 411 | 0/50 | 0/53 | 0/44 | 7/35 | Diaphragm |
| 411 | 0/47 | 57/97 | 34/58 | 35/80 | |
| 411 | 0/51 | 19/48 | 20/40 | 33/68 | |
| 411 | 0/48 | 28/58 | 31/72 | 37/75 | |
| 411 | 0/43 | 34/58 | 22/57 | 36/77 | |
| 411 | 0/43 | 22/56 | 30/53 | 41/79 | |
| 411 | 0/50 | 36/73 | 28/49 | 38/77 | |
| 411 | 0/43 | 26/57 | 26/67 | 35/77 | |
| 411 | 0/44 | 0/45 | 6/27 | 0/62 | Skin |
| 411 | 0/56 | 0/30 | 4/18 | 0/44 | Skin |
| 411 | 0/39 | 0/25 | 0/19 | 16/71 | Diaphragm |
| 411 | 0/46 | 0/35 | 0/27 | 7/35 | Diaphragm |

**TABLE S3**

| CDH Proband Family | Sex | Location | Chromosome | Position | Reference allele | Alternate allele | Tissue found in | Father |
|---|---|---|---|---|---|---|---|---|
| 411 | Male | Intergenic | Y | 21994006 | CAG | C | Skin | 0/29 |
| 411 | Male | Intergenic | Y | 23424401 | T | C | All three | 0/37 |
| 716 | Female | Intergenic | 1 | 25520943 | G | A | All three | 0/53 |
| 716 | Female | Intergenic | 1 | 108550027 | G | A | All three | 0/55 |
| 716 | Female | Intergenic | 1 | 188119862 | C | T | All three | 0/60 |
| 716 | female | Intergenic | 1 | 190952170 | A | G | Diaphragm+Skin | 0/20 |
| 716 | Female | Intergenic | 1 | 219133910 | G | A | All three | 0/54 |
| 716 | Female | Intergenic | 1 | 241538742 | A | G | All three | 0/64 |
| 716 | Female | Intergenic | 2 | 60131252 | T | C | All three | 0/56 |
| 716 | Female | Intergenic | 2 | 237733339 | C | T | All three | 0/48 |
| 716 | Female | Intergenic | 2 | 241273592 | G | T | All three | 0/48 |
| 716 | Female | Intergenic | 3 | 31317266 | C | T | All three | 0/65 |
| 716 | Female | Intergenic | 3 | 70707930 | T | C | All three | 0/51 |
| 716 | Female | Intergenic | 3 | 87755171 | C | T | All three | 0/51 |
| 716 | Female | Intergenic | 3 | 96134789 | G | A | All three | 0/58 |
| 716 | Female | Intergenic | 3 | 116969700 | C | T | All three | 0/61 |
| 716 | Female | Intergenic | 4 | 76738805 | A | C | All three | 0/55 |
| 716 | Female | Intergenic | 4 | 90360867 | C | G | All three | 0/45 |
| 716 | Female | Intergenic | 4 | 92765694 | T | A | All three | 0/55 |
| 716 | Female | Intergenic | 4 | 165797047 | C | T | All three | 0/49 |
| 716 | Female | Intergenic | 5 | 13079144 | A | T | All three | 0/45 |
| 716 | Female | Intergenic | 5 | 57641900 | G | T | Skin | 0/48 |
| 716 | Female | Intergenic | 5 | 92381616 | C | G | All three | 0/62 |
| 716 | Female | Intergenic | 6 | 18809404 | A | C | All three | 0/66 |
| 716 | Female | Intergenic | 6 | 50605045 | A | G | All three | 0/57 |
| 716 | Female | Intergenic | 6 | 69243495 | C | G | All three | 0/41 |
| 716 | Female | Intergenic | 6 | 78773081 | C | T | All three | 0/62 |
| 716 | Female | Intergenic | 7 | 2657275 | C | T | All three | 0/47 |
| 716 | Female | Intergenic | 7 | 6718899 | A | G | All three | 0/68 |
| 716 | Female | Intergenic | 7 | 19285491 | C | T | All three | 0/50 |

De Novo Intergenic All

| CDH Proband Family | Mother | Proband Blood | Proband Skin | Proband Diaphragm | Variants with 0 reads in other tissues |
|---|---|---|---|---|---|
| 411 | 0/0 | 4/44 | 6/24 | 0/36 | |
| 411 | 0/0 | 35/35 | 35/36 | 31/31 | |
| 716 | 0/58 | 24/50 | 40/84 | 23/51 | |
| 716 | 0/58 | 38/66 | 36/64 | 33/66 | |
| 716 | 0/65 | 27/61 | 38/66 | 23/56 | |
| 716 | 0/44 | 3/25 | 7/27 | 4/16 | |
| 716 | 0/50 | 37/69 | 28/56 | 29/66 | |
| 716 | 0/58 | 40/85 | 46/78 | 33/67 | |
| 716 | 0/57 | 28/55 | 43/90 | 49/79 | |
| 716 | 0/51 | 29/68 | 32/76 | 39/74 | |
| 716 | 0/54 | 42/89 | 30/77 | 26/55 | |
| 716 | 0/56 | 34/61 | 26/59 | 39/77 | |
| 716 | 0/50 | 26/62 | 32/61 | 35/66 | |
| 716 | 0/59 | 35/82 | 34/67 | 38/66 | |
| 716 | 0/60 | 31/58 | 46/81 | 23/58 | |
| 716 | 0/72 | 35/79 | 37/71 | 23/52 | |
| 716 | 0/56 | 26/54 | 21/45 | 45/67 | |
| 716 | 0/61 | 29/59 | 25/53 | 29/60 | |
| 716 | 0/46 | 35/69 | 40/63 | 31/63 | |
| 716 | 0/57 | 39/68 | 34/69 | 39/72 | |
| 716 | 0/59 | 27/60 | 36/63 | 28/51 | |
| 716 | 0/53 | 0/56 | 14/68 | 0/48 | Skin |
| 716 | 0/59 | 40/72 | 27/57 | 32/57 | |
| 716 | 0/68 | 30/68 | 33/77 | 22/47 | |
| 716 | 0/53 | 39/75 | 28/58 | 36/56 | |
| 716 | 0/61 | 26/53 | 31/72 | 35/69 | |
| 716 | 0/58 | 38/79 | 36/62 | 38/76 | |
| 716 | 0/57 | 30/58 | 23/54 | 29/67 | |
| 716 | 0/55 | 34/75 | 35/78 | 34/68 | |
| 716 | 0/63 | 36/70 | 38/73 | 41/77 | |

De Novo Intergenic All

| CDH Proband Family | Sex | Location | Chromosome | Position | Reference allele | Alternate allele | Tissue found in | Father |
|---|---|---|---|---|---|---|---|---|
| 716 | Female | Intergenic | 7 | 96000892 | G | T | All three | 0/46 |
| 716 | Female | Intergenic | 8 | 23549713 | A | G | All three | 0/50 |
| 716 | Female | Intergenic | 8 | 50342185 | C | A | All three | 0/55 |
| 716 | Female | Intergenic | 8 | 53690025 | G | A | All three | 0/65 |
| 716 | Female | Intergenic | 8 | 121394186 | C | G | All three | 0/43 |
| 716 | Female | Intergenic | 9 | 37870079 | C | A | All three | 0/52 |
| 716 | Female | Intergenic | 9 | 89980836 | A | G | Diaphragm | 0/52 |
| 716 | Female | Intergenic | 9 | 92940311 | C | T | All three | 0/62 |
| 716 | Female | Intergenic | 10 | 21531548 | A | T | All three | 0/61 |
| 716 | female | Intergenic | 10 | 78610342 | GACAA | G | All three | 0/79 |
| 716 | Female | Intergenic | 10 | 102805027 | C | T | All three | 0/49 |
| 716 | Female | Intergenic | 10 | 111937754 | A | G | All three | 0/55 |
| 716 | Female | Intergenic | 11 | 68910808 | C | T | All three | 0/52 |
| 716 | Female | Intergenic | 11 | 68917454 | A | T | All three | 0/63 |
| 716 | Female | Intergenic | 11 | 85940710 | C | T | All three | 0/52 |
| 716 | Female | Intergenic | 12 | 2894972 | G | A | All three | 0/71 |
| 716 | Female | Intergenic | 12 | 20253658 | G | A | All three | 0/60 |
| 716 | Female | Intergenic | 12 | 61786788 | C | T | All three | 0/56 |
| 716 | female | Intergenic | 12 | 73894281 | T | C | All three | 0/67 |
| 716 | Female | Intergenic | 12 | 81133670 | T | C | All three | 0/44 |
| 716 | Female | Intergenic | 12 | 128640569 | T | C | All three | 0/60 |
| 716 | Female | Intergenic | 13 | 38770384 | A | T | All three | 0/56 |
| 716 | Female | Intergenic | 13 | 53424768 | G | T | Skin | 0/49 |
| 716 | Female | Intergenic | 14 | 94631627 | C | T | All three | 0/47 |
| 716 | Female | Intergenic | 14 | 98723131 | C | T | All three | 0/57 |
| 716 | Female | Intergenic | 15 | 53418167 | A | G | All three | 0/59 |
| 716 | Female | Intergenic | 18 | 26978994 | C | T | All three | 0/53 |
| 716 | Female | Intergenic | 18 | 48313755 | G | T | All three | 0/60 |
| 716 | Female | Intergenic | 18 | 63851839 | G | C | All three | 0/55 |
| 716 | Female | Intergenic | 19 | 583802 | C | T | Skin | 0/53 |

**TABLE S3**

| CDH Proband Family | Mother | Proband Blood | Proband Skin | Proband Diaphragm | Variants with 0 reads in other tissues |
|---|---|---|---|---|---|
| 716 | 0/52 | 23/83 | 19/64 | 29/71 | |
| 716 | 0/47 | 17/55 | 34/68 | 37/76 | |
| 716 | 0/55 | 21/47 | 42/77 | 48/85 | |
| 716 | 0/54 | 25/44 | 26/66 | 30/58 | |
| 716 | 0/43 | 20/50 | 38/72 | 31/54 | |
| 716 | 0/52 | 29/68 | 31/71 | 35/66 | |
| 716 | 0/59 | 0/54 | 7/59 | 18/78 | |
| 716 | 0/56 | 28/66 | 37/73 | 38/77 | |
| 716 | 0/56 | 33/61 | 37/73 | 30/61 | |
| 716 | 0/80 | 38/79 | 35/88 | 45/91 | |
| 716 | 0/49 | 40/77 | 45/80 | 36/73 | |
| 716 | 0/47 | 38/77 | 43/81 | 43/68 | |
| 716 | 0/59 | 34/72 | 36/71 | 27/59 | |
| 716 | 0/52 | 40/72 | 35/69 | 36/65 | |
| 716 | 0/53 | 28/63 | 38/76 | 31/57 | |
| 716 | 0/58 | 47/80 | 31/61 | 38/76 | |
| 716 | 0/64 | 39/77 | 29/50 | 43/61 | |
| 716 | 0/74 | 36/76 | 34/67 | 36/61 | |
| 716 | 0/77 | 43/81 | 40/73 | 33/71 | |
| 716 | 0/63 | 37/65 | 38/72 | 30/70 | |
| 716 | 0/64 | 22/57 | 38/73 | 35/76 | |
| 716 | 0/42 | 28/57 | 30/63 | 32/61 | |
| 716 | 0/48 | 0/44 | 11/53 | 0/45 | Skin |
| 716 | 0/61 | 30/55 | 31/66 | 23/44 | |
| 716 | 0/55 | 34/65 | 32/66 | 30/67 | |
| 716 | 0/60 | 24/67 | 30/66 | 35/66 | |
| 716 | 0/49 | 28/62 | 34/74 | 38/74 | |
| 716 | 0/53 | 35/64 | 33/71 | 40/74 | |
| 716 | 0/54 | 38/76 | 41/83 | 33/67 | |
| 716 | 0/42 | 0/43 | 14/60 | 0/50 | Skin |

**TABLE S3**

| CDH Proband Family | Sex | Location | Chromosome | Position | Reference allele | Alternate allele | Tissue found in | Father |
|---|---|---|---|---|---|---|---|---|
| 716 | Female | Intergenic | 19 | 12407418 | A | C | All three | 0/78 |
| 716 | Female | Intergenic | 20 | 40498713 | A | T | All three | 0/41 |
| 716 | Female | Intergenic | 21 | 26441156 | G | T | All three | 0/53 |
| 716 | Female | Intergenic | 21 | 29172156 | A | T | All three | 0/59 |
| 716 | Female | Intergenic | X | 39689788 | C | G | All three | 0/31 |
| 716 | Female | Intergenic | X | 65140289 | G | A | All three | 0/25 |
| 716 | Female | Intergenic | X | 82221769 | C | T | All three | 0/30 |
| 716 | Female | Intergenic | X | 88110135 | C | T | All three | 0/34 |
| 716 | Female | Intergenic | X | 89554886 | C | T | All three | 0/25 |
| 716 | Female | Intergenic | X | 105821982 | G | T | All three | 0/35 |
| 809 | Female | Intergenic | 1 | 107022943 | G | T | Diaphragm | 0/38 |
| 809 | Female | Intergenic | 1 | 159684393 | C | T | Diaphragm | 0/36 |
| 809 | Female | Intergenic | 1 | 229943691 | T | A | All three | 0/41 |
| 809 | Female | Intergenic | 1 | 248631521 | G | A | All three | 0/47 |
| 809 | female | Intergenic | 2 | 52087068 | T | A | Diaphragm+Blood | 1/49 |
| 809 | Female | Intergenic | 2 | 52678867 | T | G | Skin | 0/39 |
| 809 | Female | Intergenic | 2 | 53069390 | G | A | Diaphragm | 0/44 |
| 809 | Female | Intergenic | 2 | 81121345 | A | G | Skin | 0/47 |
| 809 | Female | Intergenic | 2 | 106320315 | C | G | Diaphragm | 0/36 |
| 809 | Female | Intergenic | 2 | 111473446 | G | A | Skin | 0/43 |
| 809 | Female | Intergenic | 2 | 115175328 | G | A | Skin | 0/43 |
| 809 | Female | Intergenic | 2 | 118456374 | T | C | All three | 0/46 |
| 809 | Female | Intergenic | 2 | 125969667 | T | A | Skin | 0/48 |
| 809 | Female | Intergenic | 2 | 126870837 | T | A | Diaphragm | 0/46 |
| 809 | female | Intergenic | 2 | 165063447 | G | A | Skin+Blood | 0/64 |
| 809 | Female | Intergenic | 2 | 170283195 | C | T | Diaphragm | 0/47 |
| 809 | Female | Intergenic | 2 | 184254426 | A | T | Diaphragm | 0/42 |
| 809 | Female | Intergenic | 2 | 185100776 | C | A | Diaphragm | 0/44 |
| 809 | Female | Intergenic | 2 | 212004910 | C | A | Diaphragm | 0/46 |
| 809 | Female | Intergenic | 2 | 214077949 | ATTAC | A | Skin | 0/41 |

**TABLE S3**

| CDH Proband Family | Mother | Proband Blood | Proband Skin | Proband Diaphragm | Variants with 0 reads in other tissues |
|---|---|---|---|---|---|
| 716 | 0/74 | 46/68 | 28/54 | 44/62 | |
| 716 | 0/46 | 28/68 | 37/68 | 25/59 | |
| 716 | 0/58 | 27/62 | 41/73 | 28/59 | |
| 716 | 0/62 | 44/74 | 34/84 | 30/58 | |
| 716 | 0/57 | 31/66 | 24/54 | 34/81 | |
| 716 | 0/47 | 40/79 | 32/68 | 31/70 | |
| 716 | 0/55 | 39/68 | 37/69 | 34/54 | |
| 716 | 0/63 | 35/64 | 36/69 | 35/68 | |
| 716 | 0/44 | 34/72 | 25/57 | 35/82 | |
| 716 | 0/48 | 35/68 | 39/66 | 33/61 | |
| 809 | 0/60 | 0/68 | 0/57 | 22/90 | Diaphragm |
| 809 | 0/58 | 0/59 | 0/65 | 18/68 | Diaphragm |
| 809 | 0/51 | 27/73 | 43/70 | 29/61 | |
| 809 | 0/52 | 26/60 | 35/61 | 40/78 | |
| 809 | 0/88 | 19/64 | 4/48 | 37/79 | |
| 809 | 0/66 | 0/63 | 18/59 | 0/64 | Skin |
| 809 | 0/60 | 0/52 | 0/53 | 15/69 | Diaphragm |
| 809 | 0/61 | 0/60 | 18/61 | 0/49 | Skin |
| 809 | 0/65 | 0/60 | 0/62 | 24/77 | Diaphragm |
| 809 | 0/53 | 0/47 | 16/73 | 0/53 | Skin |
| 809 | 0/67 | 0/49 | 17/57 | 0/62 | Skin |
| 809 | 0/56 | 26/65 | 35/71 | 39/71 | |
| 809 | 0/47 | 0/61 | 18/65 | 0/61 | Skin |
| 809 | 0/56 | 0/53 | 0/42 | 20/73 | Diaphragm |
| 809 | 0/78 | 19/83 | 26/93 | 12/82 | |
| 809 | 0/50 | 0/41 | 0/51 | 18/69 | Diaphragm |
| 809 | 0/59 | 0/65 | 0/57 | 31/87 | Diaphragm |
| 809 | 0/62 | 0/58 | 0/61 | 24/89 | Diaphragm |
| 809 | 0/71 | 0/67 | 0/68 | 27/74 | Diaphragm |
| 809 | 0/63 | 0/54 | 17/58 | 0/54 | Skin |

**TABLE S3**

| CDH Proband Family | Sex | Location | Chromosome | Position | Reference allele | Alternate allele | Tissue found in | Father |
|---|---|---|---|---|---|---|---|---|
| 809 | Female | Intergenic | 2 | 235795397 | C | T | Skin | 0/40 |
| 809 | Female | Intergenic | 3 | 5446160 | G | A | Skin | 0/39 |
| 809 | female | Intergenic | 3 | 35466073 | C | T | Skin+Blood | 0/53 |
| 809 | Female | Intergenic | 3 | 74259109 | A | G | All three | 0/39 |
| 809 | Female | Intergenic | 3 | 74801621 | G | A | Diaphragm | 0/42 |
| 809 | Female | Intergenic | 3 | 96502668 | T | A | Diaphragm | 0/45 |
| 809 | Female | Intergenic | 3 | 99014368 | G | C | Skin | 0/48 |
| 809 | Female | Intergenic | 3 | 117747604 | G | T | Skin | 0/46 |
| 809 | Female | Intergenic | 3 | 118151660 | G | T | Diaphragm | 0/58 |
| 809 | Female | Intergenic | 3 | 137898799 | C | CAT | Skin | 0/44 |
| 809 | Female | Intergenic | 3 | 162105735 | C | G | Diaphragm | 0/43 |
| 809 | Female | Intergenic | 3 | 172322331 | C | T | Diaphragm | 0/48 |
| 809 | female | Intergenic | 4 | 4112769 | C | T | Skin | 0/52 |
| 809 | Female | Intergenic | 4 | 8715501 | G | A | All three | 0/44 |
| 809 | female | Intergenic | 4 | 12826901 | A | T | Skin | 0/54 |
| 809 | Female | Intergenic | 4 | 23219247 | C | T | Diaphragm | 0/48 |
| 809 | Female | Intergenic | 4 | 30105753 | G | T | Diaphragm | 0/35 |
| 809 | Female | Intergenic | 4 | 30258855 | G | T | Skin | 0/44 |
| 809 | Female | Intergenic | 4 | 36849853 | G | A | Skin | 0/62 |
| 809 | Female | Intergenic | 4 | 59397258 | C | T | All three | 0/48 |
| 809 | Female | Intergenic | 4 | 64527308 | T | G | Diaphragm | 0/42 |
| 809 | Female | Intergenic | 4 | 75368244 | G | T | Skin | 0/45 |
| 809 | Female | Intergenic | 4 | 118404267 | C | T | Diaphragm | 0/50 |
| 809 | Female | Intergenic | 4 | 139479057 | C | G | Diaphragm | 0/43 |
| 809 | Female | Intergenic | 4 | 176517204 | G | T | Diaphragm | 0/49 |
| 809 | Female | Intergenic | 4 | 187684926 | T | C | Skin | 0/41 |
| 809 | Female | Intergenic | 5 | 4375226 | C | T | Diaphragm | 0/42 |
| 809 | Female | Intergenic | 5 | 13302657 | G | A | Diaphragm | 0/46 |
| 809 | Female | Intergenic | 5 | 24990895 | C | T | Diaphragm | 0/44 |
| 809 | Female | Intergenic | 5 | 45771487 | G | A | Diaphragm | 0/39 |

De Novo Intergenic All

| CDH Proband Family | Mother | Proband Blood | Proband Skin | Proband Diaphragm | Variants with 0 reads in other tissues |
|---|---|---|---|---|---|
| 809 | 0/72 | 0/58 | 23/69 | 0/67 | Skin |
| 809 | 0/62 | 0/61 | 22/67 | 0/62 | Skin |
| 809 | 0/66 | 19/89 | 19/68 | 11/80 | |
| 809 | 0/59 | 34/70 | 33/59 | 30/64 | |
| 809 | 0/52 | 0/54 | 0/58 | 23/73 | Diaphragm |
| 809 | 0/59 | 0/64 | 0/59 | 24/82 | Diaphragm |
| 809 | 0/73 | 0/54 | 25/76 | 0/70 | Skin |
| 809 | 0/59 | 0/60 | 19/61 | 0/63 | Skin |
| 809 | 0/72 | 0/62 | 0/64 | 19/85 | Diaphragm |
| 809 | 0/61 | 0/61 | 15/50 | 0/56 | Skin |
| 809 | 0/57 | 0/66 | 0/72 | 25/65 | Diaphragm |
| 809 | 0/48 | 0/52 | 0/48 | 24/73 | Diaphragm |
| 809 | 1/74 | 0/79 | 24/67 | 0/91 | Skin |
| 809 | 0/58 | 42/69 | 40/76 | 33/73 | |
| 809 | 1/54 | 0/60 | 20/51 | 0/66 | Skin |
| 809 | 0/73 | 0/51 | 0/79 | 13/65 | Diaphragm |
| 809 | 0/49 | 0/41 | 0/50 | 24/75 | Diaphragm |
| 809 | 0/61 | 0/49 | 22/62 | 0/45 | Skin |
| 809 | 0/66 | 0/75 | 19/54 | 0/66 | Skin |
| 809 | 0/77 | 41/70 | 43/67 | 31/58 | |
| 809 | 0/64 | 0/59 | 0/51 | 22/70 | Diaphragm |
| 809 | 0/66 | 0/81 | 29/70 | 0/67 | Skin |
| 809 | 0/79 | 0/60 | 0/66 | 22/69 | Diaphragm |
| 809 | 0/65 | 0/54 | 0/54 | 16/74 | Diaphragm |
| 809 | 0/63 | 0/65 | 0/59 | 24/74 | Diaphragm |
| 809 | 0/59 | 0/51 | 24/79 | 0/56 | Skin |
| 809 | 0/64 | 0/68 | 0/53 | 20/76 | Diaphragm |
| 809 | 0/68 | 0/53 | 0/52 | 21/69 | Diaphragm |
| 809 | 0/60 | 0/50 | 0/53 | 20/71 | Diaphragm |
| 809 | 0/56 | 0/55 | 0/53 | 18/67 | Diaphragm |

**TABLE S3**

| CDH Proband Family | Sex | Location | Chromosome | Position | Reference allele | Alternate allele | Tissue found in | Father |
|---|---|---|---|---|---|---|---|---|
| 809 | Female | Intergenic | 5 | 117288082 | G | T | Diaphragm | 0/43 |
| 809 | female | Intergenic | 5 | 132509196 | T | C | All three | 0/59 |
| 809 | Female | Intergenic | 5 | 144084315 | G | GA | All three | 0/40 |
| 809 | Female | Intergenic | 5 | 144352082 | A | T | Skin | 0/47 |
| 809 | Female | Intergenic | 5 | 161028079 | G | A | Diaphragm | 0/53 |
| 809 | Female | Intergenic | 5 | 163064447 | G | A | Diaphragm | 0/46 |
| 809 | Female | Intergenic | 5 | 175131810 | G | T | Skin | 0/43 |
| 809 | Female | Intergenic | 6 | 9617082 | G | T | Skin | 0/38 |
| 809 | Female | Intergenic | 6 | 57726168 | G | A | Skin | 0/45 |
| 809 | Female | Intergenic | 6 | 84499729 | T | C | All three | 0/43 |
| 809 | Female | Intergenic | 6 | 118084618 | A | T | Skin | 0/55 |
| 809 | Female | Intergenic | 6 | 142319520 | A | G | Skin | 0/40 |
| 809 | Female | Intergenic | 7 | 46969823 | T | A | All three | 0/39 |
| 809 | Female | Intergenic | 7 | 56693729 | A | T | Diaphragm | 0/45 |
| 809 | Female | Intergenic | 7 | 79427437 | G | A | Diaphragm | 0/43 |
| 809 | Female | Intergenic | 7 | 93895381 | G | T | Diaphragm | 0/40 |
| 809 | Female | Intergenic | 7 | 134456152 | T | A | Skin | 0/44 |
| 809 | Female | Intergenic | 7 | 155849943 | G | T | Diaphragm | 0/42 |
| 809 | Female | Intergenic | 8 | 24834454 | C | T | All three | 0/50 |
| 809 | Female | Intergenic | 8 | 29547994 | C | A | Skin | 0/41 |
| 809 | Female | Intergenic | 8 | 30173080 | C | T | Diaphragm | 0/56 |
| 809 | Female | Intergenic | 8 | 51862888 | C | T | Skin | 0/41 |
| 809 | Female | Intergenic | 8 | 60040127 | C | T | All three | 0/47 |
| 809 | Female | Intergenic | 8 | 112091216 | T | A | Skin | 0/48 |
| 809 | Female | Intergenic | 8 | 135335680 | C | T | Skin | 0/48 |
| 809 | female | Intergenic | 8 | 138869304 | T | G | Skin+Blood | 1/62 |
| 809 | Female | Intergenic | 9 | 23952880 | C | A | Skin | 0/41 |
| 809 | Female | Intergenic | 9 | 24832115 | G | A | Diaphragm | 0/45 |
| 809 | Female | Intergenic | 9 | 30064518 | T | C | Skin | 0/43 |
| 809 | Female | Intergenic | 9 | 90691068 | C | T | Skin | 0/38 |

De Novo Intergenic All

| CDH Proband Family | Mother | Proband Blood | Proband Skin | Proband Diaphragm | Variants with 0 reads in other tissues |
|---|---|---|---|---|---|
| 809 | 0/67 | 0/56 | 0/75 | 20/88 | Diaphragm |
| 809 | 0/73 | 46/74 | 38/78 | 41/89 | |
| 809 | 0/66 | 14/70 | 27/53 | 12/58 | |
| 809 | 0/51 | 0/63 | 19/76 | 0/52 | Skin |
| 809 | 0/63 | 0/66 | 0/53 | 22/65 | Diaphragm |
| 809 | 0/57 | 0/61 | 0/56 | 16/73 | Diaphragm |
| 809 | 0/65 | 0/52 | 17/83 | 0/56 | Skin |
| 809 | 0/59 | 0/56 | 19/61 | 0/55 | Skin |
| 809 | 0/50 | 0/66 | 21/68 | 0/66 | Skin |
| 809 | 0/50 | 48/75 | 29/59 | 42/91 | |
| 809 | 0/63 | 0/63 | 30/80 | 0/60 | Skin |
| 809 | 0/62 | 0/57 | 17/65 | 0/68 | Skin |
| 809 | 0/57 | 39/82 | 31/49 | 35/73 | |
| 809 | 0/65 | 0/75 | 0/62 | 18/76 | Diaphragm |
| 809 | 0/49 | 0/56 | 0/57 | 28/88 | Diaphragm |
| 809 | 0/55 | 0/58 | 0/56 | 23/72 | Diaphragm |
| 809 | 0/51 | 0/65 | 22/75 | 0/61 | Skin |
| 809 | 0/68 | 0/60 | 0/50 | 21/70 | Diaphragm |
| 809 | 0/62 | 14/29 | 13/29 | 18/35 | |
| 809 | 0/52 | 0/53 | 20/75 | 0/57 | Skin |
| 809 | 0/56 | 0/66 | 0/59 | 23/63 | Diaphragm |
| 809 | 0/50 | 0/59 | 28/70 | 0/61 | Skin |
| 809 | 0/55 | 27/65 | 37/80 | 43/88 | |
| 809 | 0/73 | 0/57 | 21/72 | 0/57 | Skin |
| 809 | 0/56 | 0/49 | 14/70 | 0/57 | Skin |
| 809 | 0/68 | 23/77 | 26/74 | 12/74 | |
| 809 | 0/56 | 0/62 | 24/62 | 0/54 | Skin |
| 809 | 0/58 | 0/51 | 0/53 | 19/66 | Diaphragm |
| 809 | 0/67 | 0/53 | 18/71 | 0/43 | Skin |
| 809 | 0/58 | 0/53 | 17/58 | 0/49 | Skin |

De Novo Intergenic All

| CDH Proband Family | Sex | Location | Chromosome | Position | Reference allele | Alternate allele | Tissue found in | Father |
|---|---|---|---|---|---|---|---|---|
| 809 | Female | Intergenic | 9 | 99837526 | G | A | Skin | 0/38 |
| 809 | Female | Intergenic | 10 | 4733230 | A | G | All three | 0/44 |
| 809 | female | Intergenic | 10 | 9768107 | G | C | Diaphragm | 0/68 |
| 809 | Female | Intergenic | 10 | 29015321 | T | C | Diaphragm | 0/49 |
| 809 | Female | Intergenic | 10 | 42969976 | G | T | Skin | 0/44 |
| 809 | female | Intergenic | 10 | 47607914 | G | A | Skin | 0/72 |
| 809 | Female | Intergenic | 10 | 52428713 | T | C | Diaphragm | 0/44 |
| 809 | Female | Intergenic | 10 | 58006434 | C | A | Skin | 0/41 |
| 809 | Female | Intergenic | 10 | 66550830 | T | C | Skin | 0/46 |
| 809 | Female | Intergenic | 10 | 91874832 | T | A | Diaphragm | 0/55 |
| 809 | Female | Intergenic | 11 | 15343128 | AT | A | All three | 0/39 |
| 809 | Female | Intergenic | 11 | 37638703 | G | C | All three | 0/55 |
| 809 | Female | Intergenic | 11 | 38424427 | C | A | Diaphragm | 0/40 |
| 809 | Female | Intergenic | 11 | 39193391 | C | T | Diaphragm | 0/40 |
| 809 | Female | Intergenic | 11 | 61888225 | G | A | Diaphragm | 0/44 |
| 809 | Female | Intergenic | 11 | 73309897 | C | A | Diaphragm | 0/35 |
| 809 | Female | Intergenic | 11 | 80793572 | G | A | Skin | 0/42 |
| 809 | Female | Intergenic | 11 | 112170508 | T | C | All three | 0/45 |
| 809 | Female | Intergenic | 11 | 116677972 | C | T | Skin | 0/48 |
| 809 | Female | Intergenic | 11 | 123118311 | G | C | Diaphragm | 0/40 |
| 809 | Female | Intergenic | 12 | 19007497 | G | A | All three | 0/40 |
| 809 | Female | Intergenic | 12 | 71618878 | G | A | Skin | 0/47 |
| 809 | Female | Intergenic | 12 | 82273075 | G | A | Skin | 0/49 |
| 809 | Female | Intergenic | 12 | 82620453 | G | A | Skin | 0/49 |
| 809 | Female | Intergenic | 12 | 110872268 | A | C | All three | 0/55 |
| 809 | Female | Intergenic | 12 | 113182276 | C | T | Diaphragm | 0/45 |
| 809 | Female | Intergenic | 12 | 113896611 | C | A | Diaphragm | 0/47 |
| 809 | Female | Intergenic | 13 | 25735290 | C | T | Skin | 0/54 |
| 809 | Female | Intergenic | 13 | 26669122 | G | T | All three | 0/45 |
| 809 | female | Intergenic | 13 | 35502281 | A | T | Diaphragm | 0/45 |

De Novo Intergenic All

| CDH Proband Family | Mother | Proband Blood | Proband Skin | Proband Diaphragm | Variants with 0 reads in other tissues |
|---|---|---|---|---|---|
| 809 | 0/59 | 0/48 | 16/52 | 0/56 | Skin |
| 809 | 0/59 | 34/60 | 21/73 | 21/58 | |
| 809 | 0/82 | 0/71 | 0/75 | 24/68 | Diaphragm |
| 809 | 0/81 | 0/63 | 0/66 | 15/71 | Diaphragm |
| 809 | 0/43 | 0/62 | 20/68 | 0/56 | Skin |
| 809 | 0/81 | 0/89 | 22/66 | 0/70 | Skin |
| 809 | 0/65 | 0/52 | 0/49 | 23/69 | Diaphragm |
| 809 | 0/58 | 0/48 | 24/90 | 0/47 | Skin |
| 809 | 0/61 | 0/50 | 20/79 | 0/48 | Skin |
| 809 | 0/63 | 0/51 | 0/55 | 22/75 | Diaphragm |
| 809 | 0/64 | 35/78 | 34/67 | 42/70 | |
| 809 | 0/44 | 38/70 | 35/71 | 27/55 | |
| 809 | 0/73 | 0/55 | 0/55 | 15/62 | Diaphragm |
| 809 | 0/57 | 0/50 | 0/52 | 17/84 | Diaphragm |
| 809 | 0/64 | 0/72 | 0/59 | 17/81 | Diaphragm |
| 809 | 0/47 | 0/48 | 0/46 | 23/77 | Diaphragm |
| 809 | 0/57 | 0/72 | 18/61 | 0/53 | Skin |
| 809 | 0/67 | 30/55 | 35/67 | 23/69 | |
| 809 | 0/62 | 0/69 | 20/63 | 0/69 | Skin |
| 809 | 0/56 | 0/50 | 0/56 | 12/58 | Diaphragm |
| 809 | 0/64 | 40/73 | 32/76 | 36/66 | |
| 809 | 0/69 | 0/61 | 18/77 | 0/49 | Skin |
| 809 | 0/63 | 0/66 | 23/73 | 0/55 | Skin |
| 809 | 0/56 | 0/53 | 20/67 | 0/61 | Skin |
| 809 | 0/58 | 25/50 | 32/67 | 43/81 | |
| 809 | 0/61 | 0/55 | 0/59 | 24/87 | Diaphragm |
| 809 | 0/66 | 0/52 | 0/58 | 23/92 | Diaphragm |
| 809 | 0/54 | 0/61 | 19/63 | 0/62 | Skin |
| 809 | 0/65 | 30/70 | 34/51 | 38/71 | |
| 809 | 0/60 | 0/58 | 0/62 | 26/98 | Diaphragm |

De Novo Intergenic All

| CDH Proband Family | Sex | Location | Chromosome | Position | Reference allele | Alternate allele | Tissue found in | Father |
|---|---|---|---|---|---|---|---|---|
| 809 | Female | Intergenic | 13 | 57081014 | T | G | Skin | 0/52 |
| 809 | Female | Intergenic | 13 | 62968767 | G | A | Skin | 0/45 |
| 809 | Female | Intergenic | 13 | 69271119 | T | C | All three | 0/43 |
| 809 | Female | Intergenic | 13 | 80553307 | T | C | Skin | 0/39 |
| 809 | Female | Intergenic | 14 | 34632380 | C | T | All three | 0/48 |
| 809 | Female | Intergenic | 14 | 47032376 | T | C | Skin | 0/38 |
| 809 | Female | Intergenic | 14 | 67992825 | G | T | Skin | 0/47 |
| 809 | Female | Intergenic | 15 | 47132793 | C | T | All three | 0/44 |
| 809 | Female | Intergenic | 16 | 1290232 | C | T | All three | 0/24 |
| 809 | Female | Intergenic | 16 | 5365225 | C | T | Diaphragm | 0/54 |
| 809 | Female | Intergenic | 16 | 9322325 | G | A | Skin | 0/46 |
| 809 | Female | Intergenic | 16 | 13426591 | C | G | Skin | 0/44 |
| 809 | Female | Intergenic | 16 | 49287467 | C | A | Diaphragm | 0/45 |
| 809 | Female | Intergenic | 16 | 76211179 | C | A | Skin | 0/41 |
| 809 | Female | Intergenic | 17 | 14274664 | T | A | All three | 0/41 |
| 809 | Female | Intergenic | 17 | 21723862 | T | A | Diaphragm | 0/47 |
| 809 | Female | Intergenic | 17 | 52194918 | G | T | Skin | 0/51 |
| 809 | Female | Intergenic | 17 | 52357758 | A | T | Skin | 0/47 |
| 809 | Female | Intergenic | 18 | 12059828 | T | G | Skin | 0/52 |
| 809 | Female | Intergenic | 18 | 25950365 | G | T | Skin | 0/46 |
| 809 | Female | Intergenic | 18 | 37760582 | G | A | Diaphragm | 0/43 |
| 809 | Female | Intergenic | 18 | 49692265 | G | A | Skin | 0/43 |
| 809 | Female | Intergenic | 18 | 61185856 | A | T | Diaphragm | 0/45 |
| 809 | Female | Intergenic | 18 | 68778863 | C | T | Diaphragm | 0/48 |
| 809 | Female | Intergenic | 18 | 70286882 | C | T | Skin | 0/42 |
| 809 | Female | Intergenic | 19 | 28034043 | A | T | Diaphragm | 0/55 |
| 809 | Female | Intergenic | 19 | 33836836 | G | A | Diaphragm | 0/43 |
| 809 | Female | Intergenic | 20 | 38137398 | G | A | Skin | 0/45 |
| 809 | Female | Intergenic | 20 | 44965441 | C | A | Skin | 0/44 |
| 809 | female | Intergenic | 20 | 46859236 | C | T | Skin+Blood | 0/58 |

De Novo Intergenic All

| CDH Proband Family | Mother | Proband Blood | Proband Skin | Proband Diaphragm | Variants with 0 reads in other tissues |
|---|---|---|---|---|---|
| 809 | 0/66 | 0/60 | 26/77 | 0/64 | Skin |
| 809 | 0/57 | 0/45 | 18/78 | 0/49 | Skin |
| 809 | 0/49 | 39/74 | 33/74 | 34/71 | |
| 809 | 0/61 | 0/59 | 17/64 | 0/60 | Skin |
| 809 | 0/55 | 36/63 | 28/57 | 26/58 | |
| 809 | 0/57 | 0/58 | 22/68 | 0/57 | Skin |
| 809 | 0/62 | 0/54 | 21/59 | 0/62 | Skin |
| 809 | 0/55 | 32/63 | 32/66 | 43/91 | |
| 809 | 0/46 | 34/52 | 32/50 | 31/56 | |
| 809 | 0/77 | 0/64 | 0/50 | 20/66 | Diaphragm |
| 809 | 0/62 | 0/68 | 21/82 | 0/59 | Skin |
| 809 | 0/65 | 0/60 | 16/65 | 0/62 | Skin |
| 809 | 0/62 | 0/53 | 0/46 | 27/82 | Diaphragm |
| 809 | 0/52 | 0/58 | 15/60 | 0/54 | Skin |
| 809 | 0/62 | 43/70 | 34/73 | 38/70 | |
| 809 | 0/54 | 0/44 | 0/44 | 16/72 | Diaphragm |
| 809 | 0/70 | 0/55 | 21/70 | 0/53 | Skin |
| 809 | 0/57 | 0/53 | 22/72 | 0/63 | Skin |
| 809 | 0/80 | 0/70 | 22/67 | 0/70 | Skin |
| 809 | 0/72 | 0/57 | 23/75 | 0/63 | Skin |
| 809 | 0/54 | 0/54 | 0/40 | 23/85 | Diaphragm |
| 809 | 0/54 | 0/70 | 20/68 | 0/65 | Skin |
| 809 | 0/72 | 0/56 | 0/50 | 12/57 | Diaphragm |
| 809 | 0/59 | 0/57 | 0/55 | 27/65 | Diaphragm |
| 809 | 0/48 | 0/45 | 28/86 | 0/58 | Skin |
| 809 | 0/73 | 0/65 | 0/70 | 25/78 | Diaphragm |
| 809 | 0/73 | 0/53 | 0/59 | 20/88 | Diaphragm |
| 809 | 0/59 | 0/58 | 20/71 | 0/60 | Skin |
| 809 | 0/58 | 0/50 | 24/70 | 0/69 | Skin |
| 809 | 0/62 | 20/81 | 24/64 | 12/99 | |

De Novo Intergenic All

| CDH Proband Family | Sex | Location | Chromosome | Position | Reference allele | Alternate allele | Tissue found in | Father |
|---|---|---|---|---|---|---|---|---|
| 809 | Female | Intergenic | 20 | 53041363 | T | C | Skin | 0/36 |
| 809 | Female | Intergenic | 21 | 16549913 | C | A | Skin | 0/41 |
| 809 | Female | Intergenic | 21 | 18844527 | G | C | All three | 0/56 |
| 809 | Female | Intergenic | 21 | 20700243 | C | T | All three | 0/44 |
| 809 | Female | Intergenic | 21 | 28403305 | T | A | All three | 0/60 |
| 809 | Female | Intergenic | X | 455196 | C | T | All three | 0/49 |
| 809 | Female | Intergenic | X | 18180676 | G | C | Diaphragm | 0/27 |
| 809 | Female | Intergenic | X | 21286505 | T | C | Skin | 0/26 |
| 809 | Female | Intergenic | X | 35733096 | C | T | Diaphragm | 0/34 |
| 809 | female | Intergenic | X | 66349522 | C | A | Diaphragm | 0/34 |
| 809 | Female | Intergenic | X | 69433109 | C | A | Diaphragm | 0/28 |
| 809 | Female | Intergenic | X | 80911975 | G | A | Diaphragm | 0/30 |
| 809 | Female | Intergenic | X | 84394392 | A | G | Diaphragm | 0/26 |
| 809 | Female | Intergenic | X | 88218323 | C | A | Diaphragm | 0/21 |
| 809 | Female | Intergenic | X | 89240172 | A | T | Skin | 0/28 |
| 809 | Female | Intergenic | X | 89570314 | G | A | Diaphragm | 0/34 |
| 809 | Female | Intergenic | X | 94175321 | C | A | Diaphragm | 0/20 |
| 809 | Female | Intergenic | X | 120808757 | C | A | Skin | 0/21 |
| 809 | Female | Intergenic | X | 125153870 | T | G | Diaphragm | 0/34 |
| 809 | Female | Intergenic | X | 127078243 | T | G | Skin | 0/23 |
| 809 | Female | Intergenic | X | 145177018 | T | A | Skin | 0/28 |
| 809 | Female | Intergenic | X | 145921844 | G | T | Diaphragm | 0/34 |
| 967 | Male | Intergenic | 1 | 163727963 | ATCAGGTGG | G | Skin | 0/28 |
| 967 | Male | Intergenic | 2 | 58718375 | T | C | All three | 0/50 |
| 967 | Male | Intergenic | 2 | 121280003 | G | C | All three | 0/41 |
| 967 | Male | Intergenic | 2 | 158071768 | G | A | All three | 0/42 |
| 967 | Male | Intergenic | 2 | 205132597 | C | A | All three | 0/54 |
| 967 | Male | Intergenic | 3 | 42294565 | G | A | All three | 0/45 |
| 967 | Male | Intergenic | 3 | 75864306 | C | T | All three | 0/53 |
| 967 | Male | Intergenic | 3 | 80519543 | T | G | All three | 0/51 |

De Novo Intergenic All

| CDH Proband Family | Mother | Proband Blood | Proband Skin | Proband Diaphragm | Variants with 0 reads in other tissues |
|---|---|---|---|---|---|
| 809 | 0/40 | 0/57 | 31/69 | 0/62 | Skin |
| 809 | 0/55 | 0/65 | 20/69 | 0/68 | Skin |
| 809 | 0/57 | 26/69 | 35/68 | 38/83 | |
| 809 | 0/66 | 40/73 | 39/66 | 36/76 | |
| 809 | 0/65 | 33/65 | 29/60 | 30/59 | |
| 809 | 0/65 | 36/69 | 26/59 | 27/75 | |
| 809 | 0/60 | 0/51 | 0/49 | 16/60 | Diaphragm |
| 809 | 0/56 | 0/54 | 31/77 | 0/49 | Skin |
| 809 | 0/53 | 0/53 | 0/50 | 26/75 | Diaphragm |
| 809 | 0/73 | 0/70 | 0/71 | 21/82 | Diaphragm |
| 809 | 0/48 | 0/58 | 0/48 | 23/69 | Diaphragm |
| 809 | 0/67 | 0/60 | 0/62 | 25/74 | Diaphragm |
| 809 | 0/61 | 0/58 | 0/78 | 23/79 | Diaphragm |
| 809 | 0/58 | 0/52 | 0/52 | 19/59 | Diaphragm |
| 809 | 0/46 | 0/40 | 20/74 | 0/47 | Skin |
| 809 | 0/67 | 0/46 | 0/50 | 13/65 | Diaphragm |
| 809 | 0/74 | 0/64 | 0/54 | 24/72 | Diaphragm |
| 809 | 0/60 | 0/60 | 16/59 | 0/58 | Skin |
| 809 | 0/56 | 0/68 | 0/59 | 14/70 | Diaphragm |
| 809 | 0/51 | 0/54 | 19/80 | 0/57 | Skin |
| 809 | 0/58 | 0/54 | 24/77 | 0/60 | Skin |
| 809 | 0/66 | 0/50 | 0/62 | 28/78 | Diaphragm |
| 967 | 0/27 | 0/38 | 14/70 | 0/24 | Skin |
| 967 | 0/48 | 32/57 | 40/79 | 24/52 | |
| 967 | 0/50 | 19/41 | 31/64 | 19/35 | |
| 967 | 0/66 | 19/47 | 34/67 | 33/58 | |
| 967 | 0/58 | 27/52 | 29/64 | 29/45 | |
| 967 | 0/51 | 34/53 | 28/60 | 25/35 | |
| 967 | 0/53 | 33/64 | 37/75 | 19/43 | |
| 967 | 0/56 | 31/56 | 40/83 | 38/61 | |

De Novo Intergenic All

| CDH Proband Family | Sex | Location | Chromosome | Position | Reference allele | Alternate allele | Tissue found in | Father |
|---|---|---|---|---|---|---|---|---|
| 967 | Male | Intergenic | 4 | 3793114 | T | C | All three | 0/46 |
| 967 | Male | Intergenic | 4 | 45403029 | A | G | All three | 0/43 |
| 967 | Male | Intergenic | 6 | 70264933 | ATC | A | Skin | 0/60 |
| 967 | Male | Intergenic | 6 | 148384255 | T | C | All three | 0/49 |
| 967 | Male | Intergenic | 7 | 42518541 | G | A | All three | 0/59 |
| 967 | Male | Intergenic | 8 | 18093640 | C | T | Diaphragm | 0/52 |
| 967 | Male | Intergenic | 8 | 29283236 | C | T | All three | 0/49 |
| 967 | Male | Intergenic | 8 | 35788470 | T | C | All three | 0/46 |
| 967 | Male | Intergenic | 8 | 109375130 | G | A | All three | 0/51 |
| 967 | Male | Intergenic | 8 | 137476647 | C | T | Skin | 0/44 |
| 967 | Male | Intergenic | 9 | 87643765 | A | G | All three | 0/52 |
| 967 | Male | Intergenic | 10 | 9433109 | CCTT | C | All three | 0/57 |
| 967 | Male | Intergenic | 10 | 20971557 | T | G | All three | 0/55 |
| 967 | Male | Intergenic | 10 | 35283939 | A | G | All three | 0/52 |
| 967 | Male | Intergenic | 10 | 114676907 | A | AT | All three | 0/50 |
| 967 | Male | Intergenic | 11 | 35124820 | C | T | All three | 0/58 |
| 967 | Male | Intergenic | 11 | 80173545 | G | A | All three | 0/51 |
| 967 | male | Intergenic | 14 | 24359361 | G | A | Diaphragm | 0/58 |
| 967 | Male | Intergenic | 14 | 101628801 | GA | G | All three | 0/49 |
| 967 | male | Intergenic | 15 | 25739074 | C | T | Diaphragm+Skin | 0/64 |
| 967 | Male | Intergenic | 15 | 73336781 | C | A | All three | 0/49 |
| 967 | Male | Intergenic | 17 | 39613324 | T | C | All three | 0/41 |
| 967 | Male | Intergenic | 17 | 51582010 | A | G | All three | 0/41 |
| 967 | Male | Intergenic | 22 | 25083212 | TACTC | T | All three | 0/47 |
| 967 | Male | Intergenic | X | 25378699 | CAT | C | Blood | 0/34 |
| 967 | Male | Intergenic | X | 35867136 | G | A | All three | 0/23 |
| 967 | male | Intergenic | X | 68010485 | C | T | Diaphragm+Skin | 0/29 |
| 967 | Male | Intergenic | X | 90667481 | T | TA | Skin | 0/36 |
| 967 | Male | Intergenic | X | 128842994 | A | G | Skin | 0/38 |
| 967 | Male | Intergenic | X | 137019473 | C | G | Skin | 0/33 |

**TABLE S3**

| CDH Proband Family | Mother | Proband Blood | Proband Skin | Proband Diaphragm | Variants with 0 reads in other tissues |
|---|---|---|---|---|---|
| 967 | 0/51 | 30/51 | 40/79 | 16/34 | |
| 967 | 0/36 | 30/54 | 18/34 | 15/38 | |
| 967 | 0/60 | 0/61 | 8/40 | 0/42 | Skin |
| 967 | 0/56 | 26/47 | 38/74 | 25/48 | |
| 967 | 0/54 | 34/62 | 35/75 | 27/52 | |
| 967 | 0/61 | 0/56 | 0/46 | 7/34 | Diaphragm |
| 967 | 0/38 | 28/60 | 34/60 | 15/36 | |
| 967 | 0/43 | 30/59 | 34/71 | 30/61 | |
| 967 | 0/52 | 34/84 | 31/54 | 33/58 | |
| 967 | 0/53 | 0/64 | 18/73 | 0/53 | Skin |
| 967 | 0/46 | 25/54 | 20/49 | 34/46 | |
| 967 | 0/63 | 31/62 | 36/67 | 14/36 | |
| 967 | 0/47 | 31/54 | 39/64 | 18/42 | |
| 967 | 0/53 | 17/38 | 35/68 | 13/32 | |
| 967 | 0/61 | 13/29 | 20/35 | 11/20 | |
| 967 | 0/62 | 30/60 | 28/51 | 22/44 | |
| 967 | 0/59 | 22/50 | 32/68 | 25/50 | |
| 967 | 0/79 | 10/58 | 12/72 | 9/43 | |
| 967 | 0/58 | 37/63 | 32/69 | 31/52 | |
| 967 | 0/83 | 6/60 | 16/82 | 15/58 | |
| 967 | 0/48 | 24/59 | 34/61 | 30/57 | |
| 967 | 0/64 | 32/58 | 40/67 | 23/52 | |
| 967 | 0/47 | 31/53 | 37/67 | 22/38 | |
| 967 | 0/64 | 23/60 | 32/56 | 24/48 | |
| 967 | 0/58 | 5/17 | 0/38 | 0/20 | Blood |
| 967 | 0/47 | 29/29 | 37/37 | 13/13 | |
| 967 | 0/60 | 2/30 | 18/40 | 5/22 | |
| 967 | 4/59 | 0/27 | 5/25 | 0/17 | Skin |
| 967 | 0/44 | 0/32 | 10/44 | 0/21 | Skin |
| 967 | 0/50 | 0/32 | 5/25 | 0/25 | Skin |

**TABLE S3**

| CDH Proband Family | Sex | Location | Chromosome | Position | Reference allele | Alternate allele | Tissue found in | Father |
|---|---|---|---|---|---|---|---|---|
| 967 | Male | Intergenic | X | 139661068 | A | G | Diaphragm | 0/23 |
| 967 | Male | Intergenic | Y | 13317016 | G | A | All three | 0/36 |
| 967 | Male | Intergenic | Y | 17597506 | G | GA | All three | 0/32 |

**TABLE S3**

De Novo Intergenic All

| CDH Proband Family | Mother | Proband Blood | Proband Skin | Proband Diaphragm | Variants with 0 reads in other tissues |
|---|---|---|---|---|---|
| 967 | 0/52 | 0/33 | 0/35 | 5/18 | Diaphragm |
| 967 | 0/28 | 23/36 | 33/48 | 16/38 | |
| 967 | 0/0 | 24/25 | 27/29 | 15/15 | |

**TABLE S3**

| CDH Proband Family | Sex | Gene | Gene full name | Location | Chromosome | Position | Reference allele |
|---|---|---|---|---|---|---|---|
| 411 | Male | KIF1B | kinesin family member 1B | intron 32 | 1 | 10405687 | C |
| 411 | Male | ZRANB3 | zinc finger: RAN-binding domain containing 3 | intron 2 | 2 | 136149974 | T |
| 411 | Male | SUMF1 | Formylglycine-generating enzyme | intron 8 | 3 | 4114688 | G |
| 411 | Male | HRH1 | histamine receptor H1 | intron 1 | 3 | 11223160 | T |
| 411 | Male | ADAMTS9 | l metallopeptidase with thrombospondin type 1 m | intron 29 | 3 | 64549044 | C |
| 411 | Male | FARS2 | phenylalanyl-tRNA synthetase 2: mitochondrial | intron 4 | 6 | 5465769 | G |
| 411 | Male | AY927641 | | intron 2 | 6 | 88581123 | A |
| 411 | Male | UBR5 | ubiquitin protein ligase E3 component n-recognin 5 | intron 47 | 8 | 103286470 | C |
| 411 | Male | ADARB2 | enosine deaminase: RNA-specific: B2 (non-functior | intron 1 | 10 | 1769515 | C |
| 411 | Male | ZNF503-AS2 | ZNF503 antisense RNA 2 | intron 1 | 10 | 77163901 | A |
| 411 | Male | SBF2 | SET binding factor 2 | intron 16 | 11 | 9924092 | A |
| 411 | Male | LRRC4C | leucine rich repeat containing 4C | intron 1 | 11 | 41217824 | G |
| 411 | Male | PLA2G16 | HRAS-like suppressor 3 | intron 2 | 11 | 63371811 | TAAA |
| 411 | Male | OPCML | opioid binding protein/cell adhesion molecule-like | intron 2 | 11 | 132804548 | C |
| 411 | Male | PTPRO | protein tyrosine phosphatase. receptor type. O | Intron 21 | 12 | 15734228 | C |
| 411 | Male | DBX2 | developing brain homeobox 2 | intron 3 | 12 | 45413075 | A |
| 411 | Male | ANKS1B | in repeat and sterile alpha motif domain containir | intron 11 | 12 | 99830616 | C |
| 411 | Male | CRY1 | cryptoome 1 (photolyase-like) | intron 1 | 12 | 107417286 | G |
| 411 | Male | SETD1B | SET domain containing 1B | intron 16 | 12 | 122267613 | C |
| 411 | Male | GABRA5 | mma-aminobutyric acid (GABA) A receptor: alpha | intron 7 | 15 | 27178074 | C |
| 411 | Male | AK021563 | | intron 1 | 16 | 72735949 | TG |
| 411 | male | NLRP1 | NLR family: pyrin domain containing 1 | intron 7 | 17 | 5442308 | C |
| 411 | Male | RUNX1 | runt-related transcription factor 1 | intron 4 | 21 | 37165065 | C |
| 716 | Female | PPP1R12B | protein phosphatase 1: regulatory subunit 12B | intron 11 | 1 | 202411485 | G |
| 716 | Female | TON1-GTF2A1 | STON1-GTF2A1L readthrough | intron 10 | 2 | 48912220 | G |
| 716 | Female | GFPT1 | glutamine--fructose-6-phosphate transaminase 1 | intron 3 | 2 | 69591482 | G |
| 716 | Female | SESTD1 | SEC14 and spectrin domains 1 | intron 1 | 2 | 180104838 | A |
| 716 | Female | ZNF804A | zinc finger protein 804A | intron 2 | 2 | 185793971 | C |
| 716 | Female | RQCD1 | 1 required for cell differentiation1 homolog (S. por | intron 1 | 2 | 219436257 | C |
| 716 | Female | GPR55 | G protein-coupled receptor 55 | intron 1 | 2 | 231787570 | C |
| 716 | Female | LINC00880 | long intergenic non-protein coding RNA 880 | intron 1 | 3 | 156823715 | T |

**TABLE S3**

| CDH Proband Family | Alternate allele | Tissue found in | Father | Mother | Proband Blood | Proband Skin | Proband Diaphragm |
|---|---|---|---|---|---|---|---|
| 411 | T | All three | 0/56 | 0/50 | 46/88 | 27/58 | 41/85 |
| 411 | C | All three | 0/51 | 0/54 | 16/29 | 14/30 | 26/46 |
| 411 | A | All three | 0/44 | 0/50 | 37/70 | 34/66/ | 55/90 |
| 411 | A | All three | 0/46 | 0/41 | 24/44 | 33/66 | 21/56 |
| 411 | T | All three | 0/52 | 0/58 | 33/70 | 30/71 | 51/83 |
| 411 | A | All three | 0/54 | 0/51 | 27/58 | 32/65 | 32/66 |
| 411 | G | All three | 0/42 | 0/41 | 33/63 | 29/47 | 28/56 |
| 411 | CA | All three | 0/47 | 0/43 | 26/66 | 30/48 | 39/84 |
| 411 | T | All three | 0/46 | 0/41 | 29/53 | 30/49 | 29/70 |
| 411 | G | All three | 0/43 | 0/38 | 21/65 | 17/60 | 22/74 |
| 411 | G | All three | 0/52 | 0/47 | 38/61 | 33/65 | 48/98 |
| 411 | A | All three | 0/51 | 0/46 | 36/67 | 33/67 | 38/78 |
| 411 | T | All three | 0/43 | 0/39 | 50/95 | 26/54 | 41/86 |
| 411 | G | All three | 0/43 | 0/48 | 32/60 | 39/69 | 47/89 |
| 411 | A | All three | 0/62 | 0/53 | 6/16 | 9/38 | 23/50 |
| 411 | G | All three | 0/57 | 0/47 | 30/43 | 37/69 | 40/73 |
| 411 | G | All three | 0/47 | 0/47 | 29/60 | 44/70 | 35/65 |
| 411 | C | All three | 0/61 | 0/45 | 36/63 | 27/55 | 39/78 |
| 411 | T | All three | 0/48 | 0/51 | 36/84 | 41/63 | 35/73 |
| 411 | G | All three | 0/44 | 0/50 | 20/49 | 31/57 | 41/83 |
| 411 | T | All three | 0/48 | 0/44 | 31/58 | 26/41 | 39/66 |
| 411 | T | All three | 0/60 | 0/61 | 29/57 | 32/65 | 34/68 |
| 411 | G | All three | 0/58 | 0/40 | 37/68 | 23/60 | 64/94 |
| 716 | T | All three | 0/55 | 0/44 | 33/72 | 38/81 | 46/97 |
| 716 | T | All three | 0/58 | 0/60 | 34/61 | 34/70 | 34/73 |
| 716 | C | All three | 0/55 | 0/54 | 30/59 | 35/74 | 26/61 |
| 716 | G | All three | 0/49 | 0/58 | 32/54 | 31/63 | 44/72 |
| 716 | T | All three | 0/47 | 0/67 | 22/49 | 36/77 | 34/67 |
| 716 | T | All three | 0/63 | 0/59 | 15/37 | 21/47 | 24/50 |
| 716 | A | All three | 0/48 | 0/46 | 37/60 | 22/54 | 32/61 |
| 716 | C | All three | 0/40 | 0/59 | 46/82 | 35/76 | 43/90 |

**TABLE S3**

| CDH Proband Family | Sex | Gene | Gene full name | Location | Chromosome | Position | Reference allele |
|---|---|---|---|---|---|---|---|
| 716 | Female | ZBBX | zinc finger B-box domain containing | intron 18 | 3 | 167012263 | G |
| 716 | Female | EIF4G1 | eukaryotic translation initiation factor 4 gamma: 1 | intron 7 | 3 | 184040109 | T |
| 716 | Female | STX18 | syntaxin 18 | intron 5 | 4 | 4457642 | T |
| 716 | Female | FLJ13197 | | intron 1 | 4 | 38664228 | T |
| 716 | Female | STPG2 | sperm-tail PG-rich repeat containing 2 | intron 10 | 4 | 98532709 | T |
| 716 | Female | CXXC4 | CXXC finger protein 4 | 3 utr | 4 | 105390032 | T |
| 716 | Female | CDH18 | cadherin 18 | intron 2 | 5 | 20088379 | T |
| 716 | Female | NR_073113 | | intron 1 | 5 | 43598985 | T |
| 716 | Female | ANKRD31 | ankyrin repeat domain 31 | intron 4 | 5 | 74504516 | T |
| 716 | Female | MCC | mutated in colorectal cancers | intron 1 | 5 | 112734744 | G |
| 716 | Female | KCNIP1 | Kv channel interacting protein 1 | intron 1 | 5 | 170030091 | GGATCCCACC |
| 716 | Female | ANKS1A | in repeat and sterile alpha motif domain containir | intron 1 | 6 | 34925031 | T |
| 716 | Female | HMGCLL1 | 3-hydroxymethyl-3-methylglutaryl-CoA lyase-like 1 | intron 2 | 6 | 55434220 | T |
| 716 | Female | THEMIS | thymocyte selection associated | intron 4 | 6 | 128042029 | A |
| 716 | Female | AIG1 | androgen-induced 1 | intron 3 | 6 | 143506521 | C |
| 716 | Female | GRM8 | glutamate receptor: metabotropic 8 | intron 2 | 7 | 126737917 | A |
| 716 | Female | NEIL2 | nei endonuclease VIII-like 2 (E. coli) | 3 utr | 8 | 11643801 | C |
| 716 | Female | ANGPT1 | angiopoietin 1 | intron 4 | 8 | 108328503 | T |
| 716 | Female | KHDRBS3 | n containing: RNA binding: signal transduction ass | intron 2 | 8 | 136543376 | G |
| 716 | Female | DOCK8 | dedicator of cytokinesis 8 | intron 18 | 9 | 375626 | C |
| 716 | Female | ERCC6L2 | omplementing rodent repair deficiency: complem | intron 13 | 9 | 98720169 | T |
| 716 | Female | MIR3134 | microRNA 3134 | intron 1 | 9 | 114934634 | G |
| 716 | Female | ADAM12 | ADAM metallopeptidase domain 12 | intron 9 | 10 | 127787410 | A |
| 716 | Female | LRRC4C | leucine rich repeat containing 4C | intron 1 | 11 | 40832247 | C |
| 716 | Female | CD6 | CD6 molecule | intron 2 | 11 | 60774521 | G |
| 716 | Female | PDS5B | gulator of cohesion maintenance: homolog B (S. ce | intron 1 | 13 | 33169492 | A |
| 716 | Female | CLYBL | citrate lyase beta like | intron 2 | 13 | 100501587 | G |
| 716 | Female | abParts | | intron 7 | 15 | 22421168 | C |
| 716 | Female | PEAK1 | pseudopodium-enriched atypical kinase 1 | intron 1 | 15 | 77658986 | C |
| 716 | Female | ACSBG1 | acyl-CoA synthetase bubblegum family member 1 | intron 2 | 15 | 78499714 | T |
| 716 | Female | SLCO3A1 | e carrier organic anion transporter family: membe | intron 5 | 15 | 92664705 | T |

| CDH Proband Family | Alternate allele | Tissue found in | Father | Mother | Proband Blood | Proband Skin | Proband Diaphragm |
|---|---|---|---|---|---|---|---|
| 716 | T | All three | 0/59 | 0/64 | 24/65 | 23/62 | 25/64 |
| 716 | A | All three | 0/58 | 0/47 | 39/72 | 34/78 | 45/90 |
| 716 | C | All three | 0/40 | 0/42 | 35/67 | 26/59 | 32/64 |
| 716 | C | All three | 0/46 | 0/53 | 28/64 | 33/67 | 29/63 |
| 716 | C | All three | 0/48 | 0/50 | 32/65 | 39/68 | 35/57 |
| 716 | C | All three | 0/56 | 0/47 | 34/71 | 30/65 | 22/61 |
| 716 | G | All three | 0/62 | 0/67 | 32/74 | 35/73 | 37/79 |
| 716 | C | All three | 0/44 | 0/44 | 36/69 | 33/73 | 39/70 |
| 716 | C | All three | 0/47 | 0/54 | 26/60 | 36/69 | 27/63 |
| 716 | A | All three | 0/48 | 0/53 | 22/49 | 41/76 | 27/64 |
| 716 | T | All three | 0/51 | 0/66 | 29/59 | 30/76 | 32/58 |
| 716 | C | All three | 0/48 | 0/56 | 41/85 | 30/80 | 29/59 |
| 716 | C | All three | 0/55 | 0/61 | 30/65 | 41/76 | 34/63 |
| 716 | G | All three | 0/43 | 0/48 | 42/71 | 41/75 | 43/68 |
| 716 | T | All three | 0/41 | 0/63 | 35/63 | 35/68 | 39/81 |
| 716 | G | All three | 0/57 | 0/66 | 46/77 | 25/61 | 35/66 |
| 716 | T | All three | 0/59 | 0/55 | 28/55 | 36/69 | 22/54 |
| 716 | C | All three | 0/53 | 0/55 | 26/68 | 47/80 | 42/73 |
| 716 | T | All three | 0/55 | 0/60 | 40/67 | 23/57 | 34/66 |
| 716 | T | All three | 0/51 | 0/57 | 37/66 | 35/64 | 36/68 |
| 716 | A | All three | 0/53 | 0/59 | 27/51 | 31/65 | 38/69 |
| 716 | A | All three | 0/47 | 0/56 | 41/74 | 35/65 | 28/66 |
| 716 | G | All three | 0/61 | 0/60 | 30/71 | 32/72 | 33/68 |
| 716 | T | All three | 0/49 | 0/46 | 27/61 | 34/62 | 29/57 |
| 716 | C | All three | 0/53 | 0/61 | 26/53 | 26/63 | 32/68 |
| 716 | G | All three | 0/54 | 0/50 | 35/80 | 24/54 | 39/66 |
| 716 | A | All three | 0/52 | 0/53 | 33/59 | 23/58 | 41/73 |
| 716 | A | All three | 0/122 | 0/97 | 33/107 | 37/117 | 44/116 |
| 716 | T | All three | 0/50 | 0/50 | 38/63 | 37/69 | 34/58 |
| 716 | C | All three | 0/66 | 0/65 | 34/56 | 41/82 | 35/69 |
| 716 | C | All three | 0/40 | 0/46 | 39/74 | 35/76 | 41/73 |

De Novo UTR+Intron Diaphragm

| CDH Proband Family | Sex | Gene | Gene full name | Location | Chromosome | Position | Reference allele |
|---|---|---|---|---|---|---|---|
| 716 | female | LRRK1 | leucine-rich repeat kinase 1 | 3 utr | 15 | 101611011 | A |
| 716 | Female | PRKCB | protein kinase C: beta | intron 3 | 16 | 24033310 | C |
| 716 | Female | CKM | creatine kinase: muscle | intron 4 | 19 | 45818459 | C |
| 716 | Female | ZNF256 | zinc finger protein 256 | intron 1 | 19 | 58458152 | T |
| 716 | Female | SIRPA | signal-regulatory protein alpha | intron 2 | 20 | 1890924 | T |
| 716 | Female | PTPRT | protein tyrosine phosphatase: receptor type: T | intron 2 | 20 | 41496654 | A |
| 716 | Female | APCDD1L | adenomatosis polyposis coli down-regulated 1-like | intron 1 | 20 | 57048292 | C |
| 716 | Female | SYCP2 | synaptonemal complex protein 2 | intron 11 | 20 | 58488059 | TCA |
| 716 | Female | SGSM1 | small G protein signaling modulator 1 | intron 2 | 22 | 25212931 | T |
| 716 | Female | TBC1D22A | TBC1 domain family: member 22A | intron 12 | 22 | 47565909 | T |
| 716 | Female | HEPH | hephaestin | intron 2 | X | 65391234 | T |
| 809 | Female | SORCS3 | ortilin-related VPS10 domain containing receptor [illegible] | intron 5 | 1 | 10687743 | C |
| 809 | Female | SYCP1 | Synaptonemal Complex Protein 1 | intron 27 | 1 | 115498719 | C |
| 809 | Female | BRINP3 | phogenetic protein/retinoic acid inducible neural-[illegible] | intron 6 | 1 | 190140359 | T |
| 809 | Female | TRAPPC12 | Trafficking Protein Particle Complex 12 | intron 10 | 2 | 3485756 | C |
| 809 | Female | AFTPH | aftiphilin | intron 7 | 2 | 64808306 | C |
| 809 | Female | SLC4A5 | rier family 4 (sodium bicarbonate cotransporter): [illegible] | intron 8 | 2 | 74496034 | C |
| 809 | Female | SFTPB | surfactant protein B | 3 utr | 2 | 85884898 | T |
| 809 | Female | BIN1 | Bridging integrator 1 | intron 1 | 2 | 127840765 | G |
| 809 | Female | PLA2R1 | phospholipase A2 receptor 1: 180kDa | intron 11 | 2 | 160851223 | G |
| 809 | Female | BOLL | bol: boule-like (Drosophila) | intron 2 | 2 | 198644842 | C |
| 809 | Female | LINC00607 | long intergenic non-protein coding RNA 607 | intron 5 | 2 | 216549095 | G |
| 809 | Female | ULK4 | unc-51 like kinase 4 | intron 25 | 3 | 41755470 | CTG |
| 809 | Female | FHIT | fragile histidine triad | intron 5 | 3 | 60152652 | G |
| 809 | Female | PPP2R3A | rotein phosphatase 2: regulatory subunit B'': alpha | intron 6 | 3 | 135794102 | C |
| 809 | Female | SLC9A9 | mily 9: subfamily A (NHE9: cation proton antiporte | intron 4 | 3 | 143433614 | T |
| 809 | Female | NLGN1 | Neuroligin 1 | intron 4 | 3 | 173983838 | T |
| 809 | Female | PEX5L | peroxisomal biogenesis factor 5-like | intron 1 | 3 | 179719642 | C |
| 809 | Female | MB21D2 | Mab-21 domain containing 2 | intron 1 | 3 | 192557614 | G |
| 809 | female | LMLN | leishmanolysin-like (metallopeptidase M8 family) | 5 utr | 3 | 197765899 | C |
| 809 | Female | STK32B | serine/threonine kinase 32B | intron 3 | 4 | 5257751 | C |

De Novo UTR+Intron Diaphragm

**TABLE S3**

| CDH Proband Family | Alternate allele | Tissue found in | Father | Mother | Proband Blood | Proband Skin | Proband Diaphragm |
|---|---|---|---|---|---|---|---|
| 716 | T | All three | 0/64 | 0/53 | 25/55 | 31/66 | 23/65 |
| 716 | A | All three | 0/39 | 0/62 | 25/68 | 35/76 | 33/62 |
| 716 | T | All three | 0/58 | 0/52 | 30/66 | 34/85 | 36/61 |
| 716 | C | All three | 0/50 | 0/55 | 42/85 | 45/87 | 30/72 |
| 716 | G | All three | 0/47 | 0/53 | 34/56 | 31/59 | 37/76 |
| 716 | G | All three | 0/55 | 0/50 | 31/62 | 42/82 | 32/66 |
| 716 | CT | All three | 0/58 | 0/46 | 33/58 | 38/76 | 36/56 |
| 716 | T | All three | 0/47 | 0/57 | 38/69 | 39/71 | 32/57 |
| 716 | C | All three | 0/62 | 0/57 | 39/79 | 39/69 | 34/67 |
| 716 | C | All three | 0/54 | 0/47 | 39/67 | 32/61 | 33/68 |
| 716 | C | All three | 0/36 | 0/54 | 30/61 | 34/75 | 22/51 |
| 809 | T | All three | 0/43 | 0/47 | 44/68 | 30/59 | 36/75 |
| 809 | T | Diaphragm | 0/50 | 0/53 | 0/65 | 0/60 | 19/94 |
| 809 | A | Diaphragm | 0/53 | 0/64 | 0/58 | 0/52 | 26/80 |
| 809 | T | Diaphragm | 0/41 | 0/57 | 0/43 | 0/51 | 12/60 |
| 809 | T | All three | 0/42 | 0/55 | 30/68 | 37/72 | 47/87 |
| 809 | A | Diaphragm | 0/47 | 0/57 | 0/77 | 0/65 | 26/63 |
| 809 | G | All three | 0/46 | 0/61 | 29/58 | 29/59 | 38/73 |
| 809 | A | Diaphragm | 0/43 | 0/67 | 0/64 | 0/58 | 10/49 |
| 809 | A | All three | 0/45 | 0/65 | 39/67 | 37/85 | 44/85 |
| 809 | A | Diaphragm | 0/38 | 0/69 | 0/55 | 0/38 | 20/70 |
| 809 | A | All three | 0/50 | 0/64 | 38/85 | 40/69 | 30/66 |
| 809 | C | Diaphragm | 0/45 | 0/68 | 0/55 | 0/53 | 22/87 |
| 809 | T | Diaphragm | 0/42 | 0/53 | 0/61 | 0/56 | 27/78 |
| 809 | A | Diaphragm | 0/39 | 0/59 | 0/44 | 0/51 | 16/71 |
| 809 | C | All three | 0/47 | 0/57 | 38/70 | 32/59 | 36/72 |
| 809 | A | Diaphragm | 0/42 | 0/51 | 0/50 | 0/51 | 13/42 |
| 809 | T | Diaphragm | 0/46 | 0/55 | 0/51 | 0/54 | 21/58 |
| 809 | A | Diaphragm | 0/47 | 0/63 | 0/52 | 0/49 | 19/72 |
| 809 | A | Diaphragm | 0/78 | 0/72 | 0/75 | 0/63 | 33/95 |
| 809 | G | Diaphragm | 0/38 | 0/57 | 0/59 | 0/47 | 19/67 |

**TABLE S3**

| CDH Proband Family | Sex | Gene | Gene full name | Location | Chromosome | Position | Reference allele |
|---|---|---|---|---|---|---|---|
| 809 | Female | KCTD8 | :assium channel tetramerization domain containin | intron 1 | 4 | 44359274 | G |
| 809 | Female | ENPP6 | :tonucleotide pyrophosphatase/phosphodiesterase | intron 1 | 4 | 185077386 | G |
| 809 | female | EDIL3 | EGF-like repeats and discoidin I-like domains 3 | intron 6 | 5 | 83397673 | G |
| 809 | Female | FBXL17 | F-box and leucine-rich repeat protein 17 | intron 6 | 5 | 107485066 | G |
| 809 | Female | GRAMD3 | GRAM domain containing 3 | intron 12 | 5 | 125823743 | T |
| 809 | Female | TRPC7 | t receptor potential cation channel: subfamily C: m | intron 4 | 5 | 135592775 | C |
| 809 | Female | STK32A | serine/threonine kinase 32A | intron 4 | 5 | 146670218 | A |
| 809 | Female | EBF1 | early B-cell factor 1 | intron 9 | 5 | 158210489 | A |
| 809 | Female | TDP2 | tyrosyl-DNA phosphodiesterase 2 | intron 3 | 6 | 24658388 | T |
| 809 | Female | EYS | eyes shut homolog (Drosophila) | intron 12 | 6 | 65845477 | G |
| 809 | Female | COL12A1 | collagen: type XII: alpha 1 | intron 45 | 6 | 75829061 | C |
| 809 | Female | PREP | prolyl endopeptidase | intron 10 | 6 | 105750309 | T |
| 809 | Female | TAB2 | -Beta Activated Kinase 1 (MAP3K7) Binding Protei | intron 1 | 6 | 149608630 | A |
| 809 | Female | STEAP1B | STEAP family member 1B | intron 5 | 7 | 22475570 | TG |
| 809 | Female | CACNA2D1 | ım channel: voltage-dependent: alpha 2/delta subı | intron 3 | 7 | 81834306 | G |
| 809 | Female | SEMA3A | ınoglobulin domain (Ig): short basic domain: secret | intron 2 | 7 | 83763857 | T |
| 809 | Female | CNTNAP2 | Contactin Associated Protein Like 2 | intron 1 | 7 | 146201971 | T |
| 809 | Female | CSMD1 | CUB And Sushi Multiple Domains 1 | intron 27 | 8 | 3014345 | C |
| 809 | Female | NECAB1 | N-Terminal EF-Hand Calcium Binding Protein 1 | intron 4 | 8 | 91890081 | T |
| 809 | Female | TRAPPC9 | Trafficking Protein Particle Complex 9 | intron 19 | 8 | 140943579 | G |
| 809 | Female | PTPRD | protein tyrosine phosphatase: receptor type: D | intron 8 | 9 | 9448343 | G |
| 809 | Female | PTPRD | protein tyrosine phosphatase: receptor type: D | intron 7 | 9 | 9600668 | G |
| 809 | Female | PTPRD | Protein Tyrosine Phosphatase Receptor Type D | intron 5 | 9 | 9830793 | G |
| 809 | Female | LINGO2 | leucine rich repeat and Ig domain containing 2 | intron 5 | 9 | 28265953 | G |
| 809 | Female | ROR2 | receptor tyrosine kinase-like orphan receptor 2 | intron 1 | 9 | 94578709 | C |
| 809 | Female | UGCG | UDP-glucose ceramide glucosyltransferase | intron 1 | 9 | 114666708 | C |
| 809 | Female | DBC1 | cell cycle and apoptosis regulator 2 | intron 7 | 9 | 121956986 | A |
| 809 | Female | ZCCHC24 | zinc finger: CCHC domain containing 24 | intron 2 | 10 | 81168284 | C |
| 809 | Female | TACC2 | :ransforming: acidic coiled-coil containing protein : | intron 5 | 10 | 123855269 | C |
| 809 | Female | ST5 | suppression of tumorigenicity 5 | intron 1 | 11 | 8822290 | T |
| 809 | Female | LRRC4C | leucine rich repeat containing 4C | intron 1 | 11 | 40749101 | AT |

De Novo UTR+Intron Diaphragm

**TABLE S3**

| CDH Proband Family | Alternate allele | Tissue found in | Father | Mother | Proband Blood | Proband Skin | Proband Diaphragm |
|---|---|---|---|---|---|---|---|
| 809 | A | All three | 0/52 | 0/66 | 45/77 | 35/83 | 32/80 |
| 809 | A | Diaphragm | 0/48 | 0/56 | 0/45 | 0/57 | 22/79 |
| 809 | A | All three | 0/62 | 0/81 | 37/96 | 37/82 | 29/57 |
| 809 | A | All three | 0/41 | 0/62 | 30/74 | 41/76 | 40/82 |
| 809 | C | All three | 0/51 | 0/61 | 41/64 | 35/64 | 35/88 |
| 809 | T | Diaphragm | 0/37 | 0/57 | 0/54 | 0/49 | 23/77 |
| 809 | T | Diaphragm | 0/45 | 0/75 | 0/74 | 0/79 | 21/88 |
| 809 | G | All three | 0/42 | 0/53 | 37/78 | 32/74 | 49/88 |
| 809 | A | All three | 0/44 | 0/58 | 40/62 | 34/76 | 31/71 |
| 809 | T | All three | 0/51 | 0/76 | 37/83 | 28/59 | 36/81 |
| 809 | T | All three | 0/44 | 0/59 | 37/89 | 33/65 | 33/68 |
| 809 | C | All three | 0/47 | 0/56 | 37/77 | 34/60 | 31/76 |
| 809 | T | Diaphragm | 0/56 | 0/60 | 0/76 | 0/65 | 11/47 |
| 809 | T | All three | 0/45 | 0/34 | 27/58 | 46/81 | 39/71 |
| 809 | A | All three | 0/54 | 0/65 | 28/56 | 32/69 | 25/55 |
| 809 | C | Diaphragm | 0/43 | 0/56 | 0/57 | 0/45 | 21/85 |
| 809 | C | Diaphragm | 0/49 | 0/60 | 0/58 | 0/46 | 17/78 |
| 809 | T | Diaphragm | 0/44 | 0/48 | 0/56 | 0/41 | 14/58 |
| 809 | C | Diaphragm | 0/54 | 0/73 | 0/64 | 0/65 | 18/73 |
| 809 | A | Diaphragm | 0/46 | 0/57 | 0/44 | 0/45 | 14/63 |
| 809 | T | All three | 0/47 | 0/65 | 39/66 | 40/63 | 40/77 |
| 809 | A | All three | 0/44 | 0/67 | 45/76 | 41/79 | 35/66 |
| 809 | A | Diaphragm | 0/53 | 0/59 | 0/54 | 0/49 | 18/85 |
| 809 | T | Diaphragm | 0/40 | 0/60 | 0/68 | 0/56 | 27/79 |
| 809 | A | Diaphragm | 0/39 | 0/48 | 0/47 | 0/44 | 21/73 |
| 809 | T | All three | 0/37 | 0/54 | 47/87 | 37/76 | 38/63 |
| 809 | T | Diaphragm | 0/42 | 0/54 | 0/52 | 0/47 | 24/77 |
| 809 | T | Diaphragm | 0/45 | 0/53 | 0/67 | 0/65 | 23/75 |
| 809 | T | Diaphragm | 0/38 | 0/67 | 0/64 | 0/63 | 23/75 |
| 809 | G | All three | 0/50 | 0/54 | 32/73 | 43/78 | 38/79 |
| 809 | A | All three | 0/38 | 0/61 | 49/85 | 30/59 | 43/69 |

**TABLE S3**

| CDH Proband Family | Sex | Gene | Gene full name | Location | Chromosome | Position | Reference allele |
|---|---|---|---|---|---|---|---|
| 809 | Female | MAML2 | mastermind-like 2 (Drosophila) | intron 2 | 11 | 95731095 | C |
| 809 | Female | ALG9 | ALG9: alpha-1:2-mannosyltransferase | intron 9 | 11 | 111723240 | G |
| 809 | Female | KIRREL3 | kin of IRRE like 3 (Drosophila) | intron 1 | 11 | 126775175 | G |
| 809 | Female | BTBD11 | BTB (POZ) domain containing 11 | intron 1 | 12 | 107743246 | C |
| 809 | Female | ATP2A2 | Pase: Ca++ transporting: cardiac muscle: slow twit | intron 5 | 12 | 110734883 | C |
| 809 | Female | ZMYM2 | Zinc Finger MYM-Type Containing 2 | intron 15 | 13 | 20632248 | G |
| 809 | Female | NPAS3 | neuronal PAS domain protein 3 | intron 3 | 14 | 33833460 | A |
| 809 | female | SLC7A6 | mily 7 (amino acid transporter light chain. y+L syst | intron 4 | 16 | 68311346 | G |
| 809 | Female | SLC9A3R1 | 9: subfamily A (NHE3: cation proton antiporter 3): r | intron 1 | 17 | 72753883 | C |
| 809 | Female | ANKRD30B | Ankyrin Repeat Domain 30B | intron 3 | 18 | 14753321 | T |
| 809 | Female | LINC00907 | long intergenic non-protein coding RNA 907 | intron 4 | 18 | 39871960 | C |
| 809 | Female | DCC | deleted in colorectal carcinoma | intron 4 | 18 | 50450951 | G |
| 809 | Female | ZNF812P | 19 9808127 | intron 1 | 19 | 9808127 | T |
| 809 | Female | PRKD2 | Protein Kinase D2 | intron 2 | 19 | 47216261 | T |
| 809 | Female | CNKSR2 | Connector Enhancer Of Kinase Suppressor Of Ras 2 | intron 4 | X | 21476844 | T |
| 809 | Female | KLHL4 | kelch-like family member 4 | intron 1 | X | 86862155 | C |
| 967 | Male | SPATA6 | spermatogenesis associated 6 | intron 11 | 1 | 48785688 | G |
| 967 | Male | LRRC8C | leucine rich repeat containing 8 family: member C | intron 1 | 1 | 90115215 | C |
| 967 | Male | MERTK | c-mer proto-oncogene tyrosine kinase | intron 8 | 2 | 112744068 | C |
| 967 | Male | THSD7B | thrombospondin: type I: domain containing 7B | intron 13 | 2 | 138202217 | CCT |
| 967 | Male | SCHIP1 | Schwannomin Interacting Protein 1 | intron 2 | 3 | 159529771 | T |
| 967 | Male | NAALADL2 | N-acetylated alpha-linked acidic dipeptidase-like 2 | intron 9 | 3 | 175240124 | C |
| 967 | Male | BC005018 | | intron 4 | 4 | 84733093 | C |
| 967 | Male | PDZD2 | PDZ domain containing 2 | intron 2 | 5 | 31946007 | T |
| 967 | Male | DQ515898 | | intron 1 | 8 | 128336350 | T |
| 967 | Male | ANTD3-TMEF | MSANTD3-TMEFF1 readthrough | intron 1 | 9 | 103248140 | T |
| 967 | Male | NELFB | negative elongation factor complex member B | intron 4 | 9 | 140151750 | C |
| 967 | Male | RSU1 | Ras suppressor protein 1 | intron 1 | 10 | 16830733 | G |
| 967 | Male | SUFU | suppressor of fused homolog (Drosophila) | intron 2 | 10 | 104283460 | G |
| 967 | Male | OR8U8 | olfactory receptor family 8 subfamily U member 8 | intron 1 | 11 | 56202375 | A |
| 967 | male | DISC1FP1 | DISC1 fusion partner 1 (non-protein coding) | intron 1 | 11 | 90243703 | TATGC |

De Novo UTR+Intron Diaphragm

**TABLE S3**

| CDH Proband Family | Alternate allele | Tissue found in | Father | Mother | Proband Blood | Proband Skin | Proband Diaphragm |
|---|---|---|---|---|---|---|---|
| 809 | CAT | Diaphragm | 0/40 | 0/73 | 0/59 | 0/53 | 21/75 |
| 809 | A | All three | 0/52 | 0/55 | 30/58 | 32/60 | 37/71 |
| 809 | A | All three | 0/60 | 0/64 | 31/61 | 35/67 | 36/73 |
| 809 | A | Diaphragm | 0/55 | 0/65 | 0/63 | 0/63 | 16/64 |
| 809 | T | All three | 0/42 | 0/69 | 26/67 | 37/67 | 36/60 |
| 809 | C | Diaphragm | 0/46 | 0/65 | 0/55 | 0/47 | 13/64 |
| 809 | G | All three | 0/42 | 0/52 | 39/71 | 27/67 | 30/60 |
| 809 | C | Diaphragm | 0/51 | 0/72 | 0/73 | 0/60 | 23/68 |
| 809 | T | Diaphragm | 0/41 | 0/50 | 0/51 | 0/54 | 21/72 |
| 809 | C | Diaphragm | 0/51 | 0/66 | 0/61 | 0/58 | 13/61 |
| 809 | A | Diaphragm | 0/41 | 0/65 | 0/72 | 0/55 | 21/78 |
| 809 | T | Diaphragm | 0/40 | 0/65 | 0/60 | 0/53 | 27/71 |
| 809 | A | Diaphragm | 0/39 | 0/52 | 0/51 | 0/44 | 27/68 |
| 809 | [illegible]TTTTCTTTTT[illegible] | All three | 0/48 | 0/69 | 15/30 | 13/36 | 29/43 |
| 809 | G | Diaphragm | 0/18 | 0/58 | 0/52 | 0/46 | 17/66 |
| 809 | A | Diaphragm | 0/35 | 0/74 | 0/61 | 0/59 | 26/69 |
| 967 | A | All three | 0/47 | 0/75 | 18/38 | 25/55 | 27/48 |
| 967 | T | All three | 0/47 | 0/58 | 29/52 | 23/54 | 35/52 |
| 967 | T | All three | 0/69 | 0/60 | 28/67 | 38/78 | 16/48 |
| 967 | C | All three | 0/54 | 0/55 | 21/40 | 25/65 | 14/30 |
| 967 | TTG | All three | 0/56 | 0/69 | 26/40 | 35/59 | 19/28 |
| 967 | T | All three | 0/47 | 0/62 | 33/63 | 22/67 | 24/57 |
| 967 | T | All three | 0/42 | 0/53 | 27/49 | 25/55 | 24/45 |
| 967 | C | All three | 0/43 | 0/43 | 22/49 | 26/53 | 29/47 |
| 967 | C | All three | 0/52 | 0/64 | 24/52 | 27/66 | 13/36 |
| 967 | G | All three | 0/54 | 0/59 | 23/43 | 39/65 | 16/46 |
| 967 | T | All three | 0/54 | 0/44 | 26/57 | 31/69 | 20/40 |
| 967 | A | All three | 0/44 | 0/58 | 35/64 | 36/65 | 22/48 |
| 967 | A | All three | 0/53 | 0/57 | 20/43 | 26/64 | 24/50 |
| 967 | G | All three | 0/42 | 0/49 | 23/49 | 34/65 | 12/36 |
| 967 | T | All three | 1/25 | 0/43 | 12/22 | 10/24 | 13/16 |

**TABLE S3**

De Novo UTR+Intron Diaphragm

| CDH Proband Family | Sex | Gene | Gene full name | Location | Chromosome | Position | Reference allele |
|---|---|---|---|---|---|---|---|
| 967 | Male | SPATA13 | Spermatogenesis Associated 13 | intron 3 | 13 | 24682941 | A |
| 967 | Male | UGGT2 | UDP-glucose glycoprotein glucosyltransferase 2 | intron 4 | 13 | 96673108 | A |
| 967 | Male | MDGA2 | domain containing glycosylphosphatidylinositol an | intron 1 | 14 | 47897999 | T |
| 967 | Male | hCG_2003567 | | intron 1 | 15 | 66926125 | A |
| 967 | Male | CD33 | CD33 molecule | intron 7 | 19 | 51739471 | G |
| 967 | Male | TBX1 | T-box 1 | intron 8 | 22 | 19768601 | G |
| 967 | male | TENM1 | teneurin transmembrane protein 1 | intron 1 | X | 124078891 | A |

**TABLE S3**

De Novo UTR+Intron Diaphragm

| CDH Proband Family | Alternate allele | Tissue found in | Father | Mother | Proband Blood | Proband Skin | Proband Diaphragm |
|---|---|---|---|---|---|---|---|
| 967 | C | Diaphragm | 0/46 | 0/58 | 0/41 | 0/60 | 6/22 |
| 967 | G | All three | 0/50 | 0/59 | 42/68 | 22/58 | 32/60 |
| 967 | C | All three | 0/62 | 0/52 | 31/60 | 30/59 | 22/47 |
| 967 | C | All three | 0/42 | 0/54 | 14/36 | 36/71 | 20/45 |
| 967 | T | All three | 0/50 | 0/62 | 20/49 | 38/61 | 18/47 |
| 967 | A | All three | 0/55 | 0/50 | 20/40 | 28/54 | 20/41 |
| 967 | C | Diaphragm | 0/36 | 0/78 | 2/20 | 3/24 | 10/32 |

**TABLE S3**

| Gene full name | Location | Chromosome | Position | Reference allele | Alternate allele | Tissue found in | Father |
|---|---|---|---|---|---|---|---|
| kinesin family member 1B | intron 32 | 1 | 10405687 | C | T | All three | 0/56 |
| zinc finger: RAN-binding domain containing 3 | intron 2 | 2 | 136149974 | T | C | All three | 0/51 |
| Formylglycine-generating enzyme | intron 8 | 3 | 4114688 | G | A | All three | 0/44 |
| histamine receptor H1 | intron 1 | 3 | 11223160 | T | A | All three | 0/46 |
| synapsin II | intron 2 | 3 | 12146936 | T | A | Skin+Blood | 0/31 |
| ADAM metallopeptidase with thrombospondin type 1 motif: [illegible] | intron 29 | 3 | 64549044 | C | T | All three | 0/52 |
| | intron 1 | 4 | 188264225 | AAG | A | Skin | 0/57 |
| lysophosphatidylcholine acyltransferase 1 | intron 6 | 5 | 1482348 | C | T | Skin | 0/46 |
| phenylalanyl-tRNA synthetase 2: mitochondrial | intron 4 | 6 | 5465769 | G | A | All three | 0/54 |
| | intron 2 | 6 | 88581123 | A | G | All three | 0/42 |
| ubiquitin protein ligase E3 component n-recognin 5 | intron 47 | 8 | 103286470 | C | CA | All three | 0/47 |
| adenosine deaminase: RNA-specific: B2 (non-functional) | intron 1 | 10 | 1769515 | C | T | All three | 0/46 |
| ZNF503 antisense RNA 2 | intron 1 | 10 | 77163901 | A | G | All three | 0/43 |
| SET binding factor 2 | intron 16 | 11 | 9924092 | A | G | All three | 0/52 |
| leucine rich repeat containing 4C | intron 1 | 11 | 41217824 | G | A | All three | 0/51 |
| HRAS-like suppressor 3 | intron 2 | 11 | 63371811 | TAAA | T | All three | 0/43 |
| opioid binding protein/cell adhesion molecule-like | intron 2 | 11 | 132804548 | C | G | All three | 0/43 |
| protein tyrosine phosphatase. receptor type. O | Intron 21 | 12 | 15734228 | C | A | All three | 0/62 |
| developing brain homeobox 2 | intron 3 | 12 | 45413075 | A | G | All three | 0/57 |
| ankyrin repeat and sterile alpha motif domain containing 1B | intron 11 | 12 | 99830616 | C | G | All three | 0/47 |
| cryptoome 1 (photolyase-like) | intron 1 | 12 | 107417286 | G | C | All three | 0/61 |
| SET domain containing 1B | intron 16 | 12 | 122267613 | C | T | All three | 0/48 |
| | intron 2 | 14 | 50529288 | ATC | A | Skin | 0/62 |
| gamma-aminobutyric acid (GABA) A receptor: alpha 5 | intron 7 | 15 | 27178074 | C | G | All three | 0/44 |
| | intron 1 | 16 | 72735949 | TG | T | All three | 0/48 |
| NLR family: pyrin domain containing 1 | intron 7 | 17 | 5442308 | C | T | All three | 0/60 |
| runt-related transcription factor 1 | intron 4 | 21 | 37165065 | C | G | All three | 0/58 |
| PTCHD1 Antisense RNA | intron 4 | X | 22444899 | GGT | G | Skin | 3/32 |
| protein phosphatase 1: regulatory subunit 12B | intron 11 | 1 | 202411485 | G | T | All three | 0/55 |
| STON1-GTF2A1L readthrough | intron 10 | 2 | 48912220 | G | T | All three | 0/58 |
| glutamine--fructose-6-phosphate transaminase 1 | intron 3 | 2 | 69591482 | G | C | All three | 0/55 |
| SEC14 and spectrin domains 1 | intron 1 | 2 | 180104838 | A | G | All three | 0/49 |
| zinc finger protein 804A | intron 2 | 2 | 185793971 | C | T | All three | 0/47 |
| RCD1 required for cell differentiation1 homolog (S. pombe) | intron 1 | 2 | 219436257 | C | T | All three | 0/63 |
| G protein-coupled receptor 55 | intron 1 | 2 | 231787570 | C | A | All three | 0/48 |
| long intergenic non-protein coding RNA 880 | intron 1 | 3 | 156823715 | T | C | All three | 0/40 |
| zinc finger B-box domain containing | intron 18 | 3 | 167012263 | G | T | All three | 0/59 |
| eukaryotic translation initiation factor 4 gamma: 1 | intron 7 | 3 | 184040109 | T | A | All three | 0/58 |
| syntaxin 18 | intron 5 | 4 | 4457642 | T | C | All three | 0/40 |

**TABLE S3**

| Gene full name | Mother | Proband Blood | Proband Skin | Proband Diaphragm |
|---|---|---|---|---|
| kinesin family member 1B | 0/50 | 46/88 | 27/58 | 41/85 |
| zinc finger: RAN-binding domain containing 3 | 0/54 | 16/29 | 14/30 | 26/46 |
| Formylglycine-generating enzyme | 0/50 | 37/70 | 34/66 | 55/90 |
| histamine receptor H1 | 0/41 | 24/44 | 33/66 | 21/56 |
| synapsin II | 0/44 | 9/44 | 21/73 | 5/54 |
| ADAM metallopeptidase with thrombospondin type 1 motif: 9 | 0/58 | 33/70 | 30/71 | 51/83 |
| | 0/45 | 0/49 | 6/28 | 0/51 |
| lysophosphatidylcholine acyltransferase 1 | 0/48 | 10/55 | 18/64 | 5/67 |
| phenylalanyl-tRNA synthetase 2: mitochondrial | 0/51 | 27/58 | 32/65 | 32/66 |
| | 0/41 | 33/63 | 29/47 | 28/56 |
| ubiquitin protein ligase E3 component n-recognin 5 | 0/43 | 26/66 | 30/48 | 39/84 |
| adenosine deaminase: RNA-specific: B2 (non-functional) | 0/41 | 29/53 | 30/49 | 29/70 |
| ZNF503 antisense RNA 2 | 0/38 | 21/65 | 17/60 | 22/74 |
| SET binding factor 2 | 0/47 | 38/61 | 33/65 | 48/98 |
| leucine rich repeat containing 4C | 0/46 | 36/67 | 33/67 | 38/78 |
| HRAS-like suppressor 3 | 0/39 | 50/95 | 26/54 | 41/86 |
| opioid binding protein/cell adhesion molecule-like | 0/48 | 32/60 | 39/69 | 47/89 |
| protein tyrosine phosphatase. receptor type. O | 0/53 | 6/16 | 9/38 | 23/50 |
| developing brain homeobox 2 | 0/47 | 30/43 | 37/69 | 40/73 |
| ankyrin repeat and sterile alpha motif domain containing 1B | 0/47 | 29/60 | 44/70 | 35/65 |
| cryptoome 1 (photolyase-like) | 0/45 | 36/63 | 27/55 | 39/78 |
| SET domain containing 1B | 0/51 | 36/84 | 41/63 | 35/73 |
| | 0/42 | 0/50 | 11/52 | 0/51 |
| gamma-aminobutyric acid (GABA) A receptor: alpha 5 | 0/50 | 20/49 | 31/57 | 41/83 |
| | 0/44 | 31/58 | 26/41 | 39/66 |
| NLR family: pyrin domain containing 1 | 0/61 | 29/57 | 32/65 | 34/68 |
| runt-related transcription factor 1 | 0/40 | 37/68 | 23/60 | 64/94 |
| PTCHD1 Antisense RNA | 0/59 | 0/37 | 7/33 | 0/51 |
| protein phosphatase 1: regulatory subunit 12B | 0/44 | 33/72 | 38/81 | 46/97 |
| STON1-GTF2A1L readthrough | 0/60 | 34/61 | 34/70 | 34/73 |
| glutamine--fructose-6-phosphate transaminase 1 | 0/54 | 30/59 | 35/74 | 26/61 |
| SEC14 and spectrin domains 1 | 0/58 | 32/54 | 31/63 | 44/72 |
| zinc finger protein 804A | 0/67 | 22/49 | 36/77 | 34/67 |
| RCD1 required for cell differentiation1 homolog (S. pombe) | 0/59 | 15/37 | 21/47 | 24/50 |
| G protein-coupled receptor 55 | 0/46 | 37/60 | 22/54 | 32/61 |
| long intergenic non-protein coding RNA 880 | 0/59 | 46/82 | 35/76 | 43/90 |
| zinc finger B-box domain containing | 0/64 | 24/65 | 23/62 | 25/64 |
| eukaryotic translation initiation factor 4 gamma: 1 | 0/47 | 39/72 | 34/78 | 45/90 |
| syntaxin 18 | 0/42 | 35/67 | 26/59 | 32/64 |

**TABLE S3**

| Gene full name | Location | Chromosome | Position | Reference allele | Alternate allele | Tissue found in | Father |
|---|---|---|---|---|---|---|---|
| | intron 1 | 4 | 38664228 | T | C | All three | 0/46 |
| sperm-tail PG-rich repeat containing 2 | intron 10 | 4 | 98532709 | T | C | All three | 0/48 |
| CXXC finger protein 4 | 3 utr | 4 | 105390032 | T | C | All three | 0/56 |
| cadherin 18 | intron 2 | 5 | 20088379 | T | G | All three | 0/62 |
| | intron 1 | 5 | 43598985 | T | C | All three | 0/44 |
| ankyrin repeat domain 31 | intron 4 | 5 | 74504516 | T | C | All three | 0/47 |
| mutated in colorectal cancers | intron 1 | 5 | 112734744 | G | A | All three | 0/48 |
| Kv channel interacting protein 1 | intron 1 | 5 | 170030091 | TGGGATCCCACCCA | T | All three | 0/51 |
| ankyrin repeat and sterile alpha motif domain containing 1A | intron 1 | 6 | 34925031 | T | C | All three | 0/48 |
| 3-hydroxymethyl-3-methylglutaryl-CoA lyase-like 1 | intron 2 | 6 | 55434220 | T | C | All three | 0/55 |
| thymocyte selection associated | intron 4 | 6 | 128042029 | A | G | All three | 0/43 |
| androgen-induced 1 | intron 3 | 6 | 143506521 | C | T | All three | 0/41 |
| glutamate receptor: metabotropic 8 | intron 2 | 7 | 126737917 | A | G | All three | 0/57 |
| nei endonuclease VIII-like 2 (E. coli) | 3 utr | 8 | 11643801 | C | T | All three | 0/59 |
| angiopoietin 1 | intron 4 | 8 | 108328503 | T | C | All three | 0/53 |
| lomain containing: RNA binding: signal transduction associat | intron 2 | 8 | 136543376 | G | T | All three | 0/55 |
| dedicator of cytokinesis 8 | intron 18 | 9 | 375626 | C | T | All three | 0/51 |
| oss-complementing rodent repair deficiency: complementati | intron 13 | 9 | 98720169 | T | A | All three | 0/53 |
| microRNA 3134 | intron 1 | 9 | 114934634 | G | A | All three | 0/47 |
| ADAM metallopeptidase domain 12 | intron 9 | 10 | 127787410 | A | G | All three | 0/61 |
| leucine rich repeat containing 4C | intron 1 | 11 | 40832247 | C | T | All three | 0/49 |
| CD6 molecule | intron 2 | 11 | 60774521 | G | C | All three | 0/53 |
| 5: regulator of cohesion maintenance: homolog B (S. cerevis | intron 1 | 13 | 33169492 | A | G | All three | 0/54 |
| citrate lyase beta like | intron 2 | 13 | 100501587 | G | A | All three | 0/52 |
| | intron 7 | 15 | 22421168 | C | A | All three | 0/122 |
| pseudopodium-enriched atypical kinase 1 | intron 1 | 15 | 77658986 | C | T | All three | 0/50 |
| acyl-CoA synthetase bubblegum family member 1 | intron 2 | 15 | 78499714 | T | C | All three | 0/66 |
| solute carrier organic anion transporter family: member 3A1 | intron 5 | 15 | 92664705 | T | C | All three | 0/40 |
| leucine-rich repeat kinase 1 | 3 utr | 15 | 101611011 | A | T | All three | 0/64 |
| protein kinase C: beta | intron 3 | 16 | 24033310 | C | A | All three | 0/39 |
| creatine kinase: muscle | intron 4 | 19 | 45818459 | C | T | All three | 0/58 |
| zinc finger protein 256 | intron 1 | 19 | 58458152 | T | C | All three | 0/50 |
| signal-regulatory protein alpha | intron 2 | 20 | 1890924 | T | G | All three | 0/47 |
| protein tyrosine phosphatase: receptor type: T | intron 2 | 20 | 41496654 | A | G | All three | 0/55 |
| adenomatosis polyposis coli down-regulated 1-like | intron 1 | 20 | 57048292 | C | CT | All three | 0/58 |
| synaptonemal complex protein 2 | intron 11 | 20 | 58488059 | TCA | T | All three | 0/47 |
| small G protein signaling modulator 1 | intron 2 | 22 | 25212931 | T | C | All three | 0/62 |
| TBC1 domain family: member 22A | intron 12 | 22 | 47565909 | T | C | All three | 0/54 |
| FERM And PDZ Domain Containing 4 | intron 4 | X | 12663028 | G | A | Skin | 0/36 |

**TABLE S3**

| Gene full name | Mother | Proband Blood | Proband Skin | Proband Diaphragm |
|---|---|---|---|---|
| | 0/53 | 28/64 | 33/67 | 29/63 |
| sperm-tail PG-rich repeat containing 2 | 0/50 | 32/65 | 39/68 | 35/57 |
| CXXC finger protein 4 | 0/47 | 34/71 | 30/65 | 22/61 |
| cadherin 18 | 0/67 | 32/74 | 35/73 | 37/79 |
| | 0/44 | 36/69 | 33/73 | 39/70 |
| ankyrin repeat domain 31 | 0/54 | 26/60 | 36/69 | 27/63 |
| mutated in colorectal cancers | 0/53 | 22/49 | 41/76 | 27/64 |
| Kv channel interacting protein 1 | 0/66 | 29/59 | 30/76 | 32/58 |
| ankyrin repeat and sterile alpha motif domain containing 1A | 0/56 | 41/85 | 30/80 | 29/59 |
| 3-hydroxymethyl-3-methylglutaryl-CoA lyase-like 1 | 0/61 | 30/65 | 41/76 | 34/63 |
| thymocyte selection associated | 0/48 | 42/71 | 41/75 | 43/68 |
| androgen-induced 1 | 0/63 | 35/63 | 35/68 | 39/81 |
| glutamate receptor: metabotropic 8 | 0/66 | 46/77 | 25/61 | 35/66 |
| nei endonuclease VIII-like 2 (E. coli) | 0/55 | 28/55 | 36/69 | 22/54 |
| angiopoietin 1 | 0/55 | 26/68 | 47/80 | 42/73 |
| lomain containing: RNA binding: signal transduction associat | 0/60 | 40/67 | 23/57 | 34/66 |
| dedicator of cytokinesis 8 | 0/57 | 37/66 | 35/64 | 36/68 |
| oss-complementing rodent repair deficiency: complementati | 0/59 | 27/51 | 31/65 | 38/69 |
| microRNA 3134 | 0/56 | 41/74 | 35/65 | 28/66 |
| ADAM metallopeptidase domain 12 | 0/60 | 30/71 | 32/72 | 33/68 |
| leucine rich repeat containing 4C | 0/46 | 27/61 | 34/62 | 29/57 |
| CD6 molecule | 0/61 | 26/53 | 26/63 | 32/68 |
| 5: regulator of cohesion maintenance: homolog B (S. cerevis | 0/50 | 35/80 | 24/54 | 39/66 |
| citrate lyase beta like | 0/53 | 33/59 | 23/58 | 41/73 |
| | 0/97 | 33/107 | 37/117 | 44/116 |
| pseudopodium-enriched atypical kinase 1 | 0/50 | 38/63 | 37/69 | 34/58 |
| acyl-CoA synthetase bubblegum family member 1 | 0/65 | 34/56 | 41/82 | 35/69 |
| solute carrier organic anion transporter family: member 3A1 | 0/46 | 39/74 | 35/76 | 41/73 |
| leucine-rich repeat kinase 1 | 0/53 | 25/55 | 31/66 | 23/65 |
| protein kinase C: beta | 0/62 | 25/68 | 35/76 | 33/62 |
| creatine kinase: muscle | 0/52 | 30/66 | 34/85 | 36/61 |
| zinc finger protein 256 | 0/55 | 42/85 | 45/87 | 30/72 |
| signal-regulatory protein alpha | 0/53 | 34/56 | 31/59 | 37/76 |
| protein tyrosine phosphatase: receptor type: T | 0/50 | 31/62 | 42/82 | 32/66 |
| adenomatosis polyposis coli down-regulated 1-like | 0/46 | 33/58 | 38/76 | 36/56 |
| synaptonemal complex protein 2 | 0/57 | 38/69 | 39/71 | 32/57 |
| small G protein signaling modulator 1 | 0/57 | 39/79 | 39/69 | 34/67 |
| TBC1 domain family: member 22A | 0/47 | 39/67 | 32/61 | 33/68 |
| FERM And PDZ Domain Containing 4 | 0/56 | 0/54 | 17/59 | 0/47 |

**TABLE S3**

| Gene full name | Location | Chromosome | Position | Reference allele | Alternate allele | Tissue found in | Father |
|---|---|---|---|---|---|---|---|
| hephaestin | intron 2 | X | 65391234 | T | C | All three | 0/36 |
| sortilin-related VPS10 domain containing receptor 3 | intron 5 | 1 | 10687743 | C | T | All three | 0/43 |
| chitinase: acidic | intron 2 | 1 | 111854262 | C | T | Skin | 0/53 |
| anaplastic lymphoma receptor tyrosine kinase | intron 5 | 2 | 29563357 | C | T | Skin | 0/54 |
| | intron 2 | 2 | 59058824 | G | A | Skin | 0/49 |
| aftiphilin | intron 7 | 2 | 64808306 | C | T | All three | 0/42 |
| surfactant protein B | 3 utr | 2 | 85884898 | T | G | All three | 0/46 |
| phospholipase A2 receptor 1: 180kDa | intron 11 | 2 | 160851223 | G | A | All three | 0/45 |
| long intergenic non-protein coding RNA 607 | intron 5 | 2 | 216549095 | G | A | All three | 0/50 |
| ier family 9: subfamily A (NHE9: cation proton antiporter 9): | intron 4 | 3 | 143433614 | T | C | All three | 0/47 |
| ventricular zone expressed PH domain-containing 1 | intron 9 | 3 | 157074940 | A | G | Skin | 0/39 |
| zinc finger: FYVE domain containing 28 | intron 3 | 4 | 2341715 | C | T | Skin | 0/36 |
| potassium channel tetramerization domain containing 8 | intron 1 | 4 | 44359274 | G | A | All three | 0/52 |
| follistatin-like 5 | intron 2 | 4 | 163016217 | G | T | Skin | 0/43 |
| | intron 4 | 5 | 9878319 | G | A | Skin | 0/47 |
| EGF-like repeats and discoidin I-like domains 3 | intron 6 | 5 | 83397673 | G | A | All three | 0/62 |
| G protein-coupled receptor 98 | intron 59 | 5 | 90064793 | C | A | Skin | 0/43 |
| F-box and leucine-rich repeat protein 17 | intron 6 | 5 | 107485066 | G | A | All three | 0/41 |
| GRAM domain containing 3 | intron 12 | 5 | 125823743 | T | C | All three | 0/51 |
| potassium channel tetramerization domain containing 16 | intron 2 | 5 | 143580957 | T | G | Skin | 0/45 |
| calcium/calmodulin-dependent protein kinase II alpha | intron 7 | 5 | 149632443 | G | A | Skin | 0/49 |
| early B-cell factor 1 | intron 9 | 5 | 158210489 | A | G | All three | 0/42 |
| tyrosyl-DNA phosphodiesterase 2 | intron 3 | 6 | 24658388 | T | A | All three | 0/44 |
| eyes shut homolog (Drosophila) | intron 12 | 6 | 65845477 | G | T | All three | 0/51 |
| collagen: type XII: alpha 1 | intron 45 | 6 | 75829061 | C | T | All three | 0/44 |
| prolyl endopeptidase | intron 10 | 6 | 105750309 | T | C | All three | 0/47 |
| family with sequence similarity 184: member A | intron 1 | 6 | 119395836 | C | G | Skin+Blood | 0/55 |
| TCF21 Antisense RNA Inducing Promoter Demethylation | intron 5 | 6 | 133975860 | C | A | Skin | 0/54 |
| STEAP family member 1B | intron 5 | 7 | 22475570 | TG | T | All three | 0/45 |
| engulfment and cell motility 1 | intron 2 | 7 | 37358037 | A | T | Skin | 0/38 |
| calcium channel: voltage-dependent: alpha 2/delta subunit 1 | intron 3 | 7 | 81834306 | G | A | All three | 0/54 |
| diacylglycerol kinase: iota | intron 28 | 7 | 137143548 | G | T | Skin | 0/43 |
| lysine (K)-specific methyltransferase 2C | intron 40 | 7 | 151866844 | C | T | Skin | 0/42 |
| deleted in liver cancer 1 | intron 2 | 8 | 13287693 | C | T | Skin | 0/38 |
| ankyrin 1: erythrocytic | intron 1 | 8 | 41676846 | C | A | Skin | 0/48 |
| protein tyrosine phosphatase: receptor type: D | intron 8 | 9 | 9448343 | G | T | All three | 0/47 |
| protein tyrosine phosphatase: receptor type: D | intron 7 | 9 | 9600668 | G | A | All three | 0/44 |
| protein tyrosine phosphatase: receptor type: D | intron 3 | 9 | 10048095 | A | C | Skin | 0/50 |
| isient receptor potential cation channel: subfamily M: memb | intron 2 | 9 | 73481507 | A | G | Skin | 0/50 |

**TABLE S3**

| Gene full name | Mother | Proband Blood | Proband Skin | Proband Diaphragm |
|---|---|---|---|---|
| hephaestin | 0/54 | 30/61 | 34/75 | 22/51 |
| sortilin-related VPS10 domain containing receptor 3 | 0/47 | 44/68 | 30/59 | 36/75 |
| chitinase: acidic | 0/73 | 0/63 | 19/61 | 0/61 |
| anaplastic lymphoma receptor tyrosine kinase | 0/64 | 0/59 | 19/65 | 0/53 |
| | 0/73 | 0/58 | 16/53 | 0/74 |
| aftiphilin | 0/55 | 30/68 | 37/72 | 47/87 |
| surfactant protein B | 0/61 | 29/58 | 29/59 | 38/73 |
| phospholipase A2 receptor 1: 180kDa | 0/65 | 39/67 | 37/85 | 44/85 |
| long intergenic non-protein coding RNA 607 | 0/64 | 38/85 | 40/69 | 30/66 |
| ier family 9: subfamily A (NHE9: cation proton antiporter 9): | 0/57 | 38/70 | 32/59 | 36/72 |
| ventricular zone expressed PH domain-containing 1 | 0/50 | 0/52 | 22/72 | 0/57 |
| zinc finger: FYVE domain containing 28 | 0/42 | 0/43 | 19/62 | 0/48 |
| potassium channel tetramerization domain containing 8 | 0/66 | 45/77 | 35/83 | 32/80 |
| follistatin-like 5 | 0/59 | 0/55 | 27/73 | 0/61 |
| | 0/68 | 0/62 | 18/65 | 0/57 |
| EGF-like repeats and discoidin I-like domains 3 | 0/81 | 37/96 | 37/82 | 29/57 |
| G protein-coupled receptor 98 | 0/58 | 0/64 | 15/64 | 0/60 |
| F-box and leucine-rich repeat protein 17 | 0/62 | 30/74 | 41/76 | 40/82 |
| GRAM domain containing 3 | 0/61 | 41/64 | 35/64 | 35/88 |
| potassium channel tetramerization domain containing 16 | 0/66 | 0/59 | 30/75 | 0/55 |
| calcium/calmodulin-dependent protein kinase II alpha | 0/52 | 0/55 | 21/76 | 0/53 |
| early B-cell factor 1 | 0/53 | 37/78 | 32/74 | 49/88 |
| tyrosyl-DNA phosphodiesterase 2 | 0/58 | 40/62 | 34/76 | 31/71 |
| eyes shut homolog (Drosophila) | 0/76 | 37/83 | 28/59 | 36/81 |
| collagen: type XII: alpha 1 | 0/59 | 37/89 | 33/65 | 33/68 |
| prolyl endopeptidase | 0/56 | 37/77 | 34/60 | 31/76 |
| family with sequence similarity 184: member A | 0/68 | 19/79 | 25/81 | 11/77 |
| TCF21 Antisense RNA Inducing Promoter Demethylation | 0/56 | 0/58 | 29/84 | 0/57 |
| STEAP family member 1B | 0/34 | 27/58 | 46/81 | 39/71 |
| engulfment and cell motility 1 | 0/66 | 0/52 | 20/71 | 0/57 |
| calcium channel: voltage-dependent: alpha 2/delta subunit 1 | 0/65 | 28/56 | 32/69 | 25/55 |
| diacylglycerol kinase: iota | 0/51 | 0/51 | 15/72 | 0/60 |
| lysine (K)-specific methyltransferase 2C | 0/53 | 0/48 | 23/77 | 0/53 |
| deleted in liver cancer 1 | 0/73 | 0/63 | 21/70 | 0/68 |
| ankyrin 1: erythrocytic | 0/52 | 0/54 | 23/72 | 0/61 |
| protein tyrosine phosphatase: receptor type: D | 0/65 | 39/66 | 40/63 | 40/77 |
| protein tyrosine phosphatase: receptor type: D | 0/67 | 45/76 | 41/79 | 35/66 |
| protein tyrosine phosphatase: receptor type: D | 0/58 | 0/68 | 29/86 | 0/66 |
| isient receptor potential cation channel: subfamily M: memb | 0/57 | 0/63 | 20/81 | 0/57 |

**TABLE S3**

| Gene full name | Location | Chromosome | Position | Reference allele | Alternate allele | Tissue found in | Father |
|---|---|---|---|---|---|---|---|
| UDP-glucose ceramide glucosyltransferase | intron 1 | 9 | 114666708 | C | T | All three | 0/37 |
| Solute Carrier Family 16 Member 12 | intron 3 | 10 | 91220525 | A | G | Skin | 0/40 |
| suppression of tumorigenicity 5 | intron 1 | 11 | 8822290 | T | G | All three | 0/50 |
| SRY (sex determining region Y)-box 6 | intron 12 | 11 | 16061906 | A | T | Skin | 0/36 |
| leucine rich repeat containing 4C | intron 1 | 11 | 40749101 | AT | A | All three | 0/38 |
| 11 83978757 | intron 1 | 11 | 83978757 | TC | T | Skin | 0/51 |
| ALG9: alpha-1:2-mannosyltransferase | intron 9 | 11 | 111723240 | G | A | All three | 0/52 |
| kin of IRRE like 3 (Drosophila) | intron 1 | 11 | 126775175 | G | A | All three | 0/60 |
| galactosidase: beta 1-like 2 | intron 6 | 11 | 134227263 | A | T | Skin | 0/45 |
| branched chain amino-acid transaminase 1. cytosolic | intron 4 | 12 | 25029237 | G | C | Skin | 0/45 |
| NEL-like 2 (chicken) | intron 16 | 12 | 44969449 | G | A | Skin | 0/48 |
| ATPase: Ca++ transporting: cardiac muscle: slow twitch 2 | intron 5 | 12 | 110734883 | C | T | All three | 0/42 |
| neuronal PAS domain protein 3 | intron 3 | 14 | 33833460 | A | G | All three | 0/42 |
| gamma-aminobutyric acid (GABA) A receptor: beta 3 | intron 3 | 15 | 26992702 | T | C | Skin | 0/51 |
| IQ motif containing GTPase activating protein 1 | intron 15 | 15 | 91003597 | C | T | Skin | 0/42 |
| RAB11 family interacting protein 4 (class II) | intron 3 | 17 | 29765052 | G | A | Skin | 0/39 |
| deleted in colorectal carcinoma | intron 5 | 18 | 50470480 | T | C | Skin | 0/46 |
| glyceraldehyde-3-phosphate dehydrogenase: spermatogenic | intron 9 | 19 | 36035048 | G | A | Skin | 0/41 |
| Protein Kinase D2 | intron 2 | 19 | 47216261 | T | TTTTCTTTTT( | All three | 0/48 |
| | intron 1 | 21 | 37474586 | A | C | Skin | 0/47 |
| zinc finger: CCHC domain containing 16 | intron 2 | X | 111601842 | C | T | Skin | 0/31 |
| spermatogenesis associated 6 | intron 11 | 1 | 48785688 | G | A | All three | 0/47 |
| leucine rich repeat containing 8 family: member C | intron 1 | 1 | 90115215 | C | T | All three | 0/47 |
| Zinc Finger Protein 695 | intron 5 | 1 | 247110379 | AAAAATCAAAGAGTAAATGGGGT( | T | Skin | 0/55 |
| c-mer proto-oncogene tyrosine kinase | intron 8 | 2 | 112744068 | C | T | All three | 0/69 |
| thrombospondin: type I: domain containing 7B | intron 13 | 2 | 138202217 | CCT | C | All three | 0/54 |
| Schwannomin Interacting Protein 1 | intron 2 | 3 | 159529771 | T | TTG | All three | 0/56 |
| N-acetylated alpha-linked acidic dipeptidase-like 2 | intron 9 | 3 | 175240124 | C | T | All three | 0/47 |
| | intron 4 | 4 | 84733093 | C | T | All three | 0/42 |
| PDZ domain containing 2 | intron 2 | 5 | 31946007 | T | C | All three | 0/43 |
| | intron 1 | 8 | 128336350 | T | C | All three | 0/52 |
| MSANTD3-TMEFF1 readthrough | intron 1 | 9 | 103248140 | T | G | All three | 0/54 |
| negative elongation factor complex member B | intron 4 | 9 | 140151750 | C | T | All three | 0/54 |
| Ras suppressor protein 1 | intron 1 | 10 | 16830733 | G | A | All three | 0/44 |
| suppressor of fused homolog (Drosophila) | intron 2 | 10 | 104283460 | G | A | All three | 0/53 |
| olfactory receptor family 8 subfamily U member 8 | intron 1 | 11 | 56202375 | A | G | All three | 0/42 |
| DISC1 fusion partner 1 (non-protein coding) | intron 1 | 11 | 90243703 | TATGC | T | All three | 1/25 |
| UDP-glucose glycoprotein glucosyltransferase 2 | intron 4 | 13 | 96673108 | A | G | All three | 0/50 |
| MAM domain containing glycosylphosphatidylinositol anchor [illegible] | intron 1 | 14 | 47897999 | T | C | All three | 0/62 |

**TABLE S3**

| Gene full name | Mother | Proband Blood | Proband Skin | Proband Diaphragm |
|---|---|---|---|---|
| UDP-glucose ceramide glucosyltransferase | 0/54 | 47/87 | 37/76 | 38/63 |
| Solute Carrier Family 16 Member 12 | 0/73 | 0/48 | 7/26 | 0/68 |
| suppression of tumorigenicity 5 | 0/54 | 32/73 | 43/78 | 38/79 |
| SRY (sex determining region Y)-box 6 | 0/53 | 0/55 | 21/75 | 0/43 |
| leucine rich repeat containing 4C | 0/61 | 49/85 | 30/59 | 43/69 |
| 11 83978757 | 0/73 | 0/69 | 18/74 | 0/65 |
| ALG9: alpha-1:2-mannosyltransferase | 0/55 | 30/58 | 32/60 | 37/71 |
| kin of IRRE like 3 (Drosophila) | 0/64 | 31/61 | 35/67 | 36/73 |
| galactosidase: beta 1-like 2 | 0/63 | 0/52 | 24/71 | 0/55 |
| branched chain amino-acid transaminase 1. cytosolic | 0/59 | 0/50 | 10/32 | 0/67 |
| NEL-like 2 (chicken) | 0/63 | 0/62 | 22/73 | 0/56 |
| ATPase: Ca++ transporting: cardiac muscle: slow twitch 2 | 0/69 | 26/67 | 37/67 | 36/60 |
| neuronal PAS domain protein 3 | 0/52 | 39/71 | 27/67 | 30/60 |
| gamma-aminobutyric acid (GABA) A receptor: beta 3 | 0/66 | 0/61 | 19/61 | 0/65 |
| IQ motif containing GTPase activating protein 1 | 0/60 | 0/49 | 18/64 | 0/53 |
| RAB11 family interacting protein 4 (class II) | 0/48 | 0/46 | 19/65 | 0/53 |
| deleted in colorectal carcinoma | 0/55 | 0/50 | 22/71 | 0/55 |
| glyceraldehyde-3-phosphate dehydrogenase: spermatogenic | 0/56 | 0/51 | 19/66 | 0/62 |
| Protein Kinase D2 | 0/69 | 15/30 | 13/36 | 29/43 |
| | 0/67 | 0/57 | 21/93 | 0/74 |
| zinc finger: CCHC domain containing 16 | 0/63 | 0/49 | 24/65 | 0/54 |
| spermatogenesis associated 6 | 0/75 | 18/38 | 25/55 | 27/48 |
| leucine rich repeat containing 8 family: member C | 0/58 | 29/52 | 23/54 | 35/52 |
| Zinc Finger Protein 695 | 5/66 | 0/44 | 23/111 | 0/52 |
| c-mer proto-oncogene tyrosine kinase | 0/60 | 28/67 | 38/78 | 16/48 |
| thrombospondin: type I: domain containing 7B | 0/55 | 21/40 | 25/65 | 14/30 |
| Schwannomin Interacting Protein 1 | 0/69 | 26/40 | 35/59 | 19/28 |
| N-acetylated alpha-linked acidic dipeptidase-like 2 | 0/62 | 33/63 | 22/67 | 24/57 |
| | 0/53 | 27/49 | 25/55 | 24/45 |
| PDZ domain containing 2 | 0/43 | 22/49 | 26/53 | 29/47 |
| | 0/64 | 24/52 | 27/66 | 13/36 |
| MSANTD3-TMEFF1 readthrough | 0/59 | 23/43 | 39/65 | 16/46 |
| negative elongation factor complex member B | 0/44 | 26/57 | 31/69 | 20/40 |
| Ras suppressor protein 1 | 0/58 | 35/64 | 36/65 | 22/48 |
| suppressor of fused homolog (Drosophila) | 0/57 | 20/43 | 26/64 | 24/50 |
| olfactory receptor family 8 subfamily U member 8 | 0/49 | 23/49 | 34/65 | 12/36 |
| DISC1 fusion partner 1 (non-protein coding) | 0/43 | 12/22 | 10/24 | 13/16 |
| UDP-glucose glycoprotein glucosyltransferase 2 | 0/59 | 42/68 | 22/58 | 32/60 |
| MAM domain containing glycosylphosphatidylinositol anchor [illegible] | 0/52 | 31/60 | 30/59 | 22/47 |

**TABLE S3**

| Gene full name | Location | Chromosome | Position | Reference allele | Alternate allele | Tissue found in | Father |
|---|---|---|---|---|---|---|---|
| | intron 1 | 15 | 66926125 | A | C | All three | 0/42 |
| CD33 molecule | intron 7 | 19 | 51739471 | G | T | All three | 0/50 |
| T-box 1 | intron 8 | 22 | 19768601 | G | A | All three | 0/55 |

| Gene full name | Mother | Proband Blood | Proband Skin | Proband Diaphragm |
|---|---|---|---|---|
| | 0/54 | 14/36 | 36/71 | 20/45 |
| CD33 molecule | 0/62 | 20/49 | 38/61 | 18/47 |
| T-box 1 | 0/50 | 20/40 | 28/54 | 20/41 |

**TABLE S3**

| CDH Proband Family | Sex | Gene | Gene name | Location | Chromosome | Position |
|---|---|---|---|---|---|---|
| 411 | Male | KIF1B | kinesin family member 1B | intron 32 | 1 | 10405687 |
| 411 | Male | ZRANB3 | zinc finger: RAN-binding domain containing 3 | intron 2 | 2 | 136149974 |
| 411 | Male | SUMF1 | Formylglycine-generating enzyme | intron 8 | 3 | 4114688 |
| 411 | Male | HRH1 | histamine receptor H1 | intron 1 | 3 | 11223160 |
| 411 | male | SYN2 | synapsin II | intron 2 | 3 | 12146936 |
| 411 | Male | ADAMTS9 | \DAM metallopeptidase with thrombospondin type 1 motif: | intron 29 | 3 | 64549044 |
| 411 | Male | FARS2 | phenylalanyl-tRNA synthetase 2: mitochondrial | intron 4 | 6 | 5465769 |
| 411 | Male | AY927641 | | intron 2 | 6 | 88581123 |
| 411 | Male | UBR5 | ubiquitin protein ligase E3 component n-recognin 5 | intron 47 | 8 | 103286470 |
| 411 | Male | ADARB2 | adenosine deaminase: RNA-specific: B2 (non-functional) | intron 1 | 10 | 1769515 |
| 411 | Male | ZNF503-AS2 | ZNF503 antisense RNA 2 | intron 1 | 10 | 77163901 |
| 411 | Male | SBF2 | SET binding factor 2 | intron 16 | 11 | 9924092 |
| 411 | Male | LRRC4C | leucine rich repeat containing 4C | intron 1 | 11 | 41217824 |
| 411 | Male | PLA2G16 | HRAS-like suppressor 3 | intron 2 | 11 | 63371811 |
| 411 | Male | OPCML | opioid binding protein/cell adhesion molecule-like | intron 2 | 11 | 132804548 |
| 411 | Male | PTPRO | protein tyrosine phosphatase. receptor type. O | Intron 21 | 12 | 15734228 |
| 411 | Male | DBX2 | developing brain homeobox 2 | intron 3 | 12 | 45413075 |
| 411 | Male | PTPRR | Protein Tyrosine Phosphatase Receptor Type R | intron 2 | 12 | 71250628 |
| 411 | Male | ANKS1B | ankyrin repeat and sterile alpha motif domain containing 1B | intron 11 | 12 | 99830616 |
| 411 | Male | CRY1 | cryptoome 1 (photolyase-like) | intron 1 | 12 | 107417286 |
| 411 | Male | SETD1B | SET domain containing 1B | intron 16 | 12 | 122267613 |
| 411 | Male | GABRA5 | gamma-aminobutyric acid (GABA) A receptor: alpha 5 | intron 7 | 15 | 27178074 |
| 411 | Male | AK021563 | | intron 1 | 16 | 72735949 |
| 411 | male | NLRP1 | NLR family: pyrin domain containing 1 | intron 7 | 17 | 5442308 |
| 411 | Male | RUNX1 | runt-related transcription factor 1 | intron 4 | 21 | 37165065 |
| 716 | Female | PPP1R12B | protein phosphatase 1: regulatory subunit 12B | intron 11 | 1 | 202411485 |
| 716 | Female | TON1-GTF2A1 | STON1-GTF2A1L readthrough | intron 10 | 2 | 48912220 |
| 716 | Female | GFPT1 | glutamine--fructose-6-phosphate transaminase 1 | intron 3 | 2 | 69591482 |
| 716 | Female | SESTD1 | SEC14 and spectrin domains 1 | intron 1 | 2 | 180104838 |
| 716 | Female | ZNF804A | zinc finger protein 804A | intron 2 | 2 | 185793971 |
| 716 | Female | RQCD1 | RCD1 required for cell differentiation1 homolog (S. pombe) | intron 1 | 2 | 219436257 |

De Novo UTR+Intron Blood

| CDH Proband Family | Reference allele | Alternate allele | Tissue found in | Father | Mother | Proband Blood | Proband Skin | Proband Diaphragm |
|---|---|---|---|---|---|---|---|---|
| 411 | C | T | All three | 0/56 | 0/50 | 46/88 | 27/58 | 41/85 |
| 411 | T | C | All three | 0/51 | 0/54 | 16/29 | 14/30 | 26/46 |
| 411 | G | A | All three | 0/44 | 0/50 | 37/70 | 34/66 | 55/90 |
| 411 | T | A | All three | 0/46 | 0/41 | 24/44 | 33/66 | 21/56 |
| 411 | T | A | Skin+Blood | 0/31 | 0/44 | 9/44 | 21/73 | 5/54 |
| 411 | C | T | All three | 0/52 | 0/58 | 33/70 | 30/71 | 51/83 |
| 411 | G | A | All three | 0/54 | 0/51 | 27/58 | 32/65 | 32/66 |
| 411 | A | G | All three | 0/42 | 0/41 | 33/63 | 29/47 | 28/56 |
| 411 | C | CA | All three | 0/47 | 0/43 | 26/66 | 30/48 | 39/84 |
| 411 | C | T | All three | 0/46 | 0/41 | 29/53 | 30/49 | 29/70 |
| 411 | A | G | All three | 0/43 | 0/38 | 21/65 | 17/60 | 22/74 |
| 411 | A | G | All three | 0/52 | 0/47 | 38/61 | 33/65 | 48/98 |
| 411 | G | A | All three | 0/51 | 0/46 | 36/67 | 33/67 | 38/78 |
| 411 | TAAA | T | All three | 0/43 | 0/39 | 50/95 | 26/54 | 41/86 |
| 411 | C | G | All three | 0/43 | 0/48 | 32/60 | 39/69 | 47/89 |
| 411 | C | A | All three | 0/62 | 0/53 | 6/16 | 9/38 | 23/50 |
| 411 | A | G | All three | 0/57 | 0/47 | 30/43 | 37/69 | 40/73 |
| 411 | GGA | G | Blood | 0/45 | 0/56 | 8/34 | 0/53 | 0/66 |
| 411 | C | G | All three | 0/47 | 0/47 | 29/60 | 44/70 | 35/65 |
| 411 | G | C | All three | 0/61 | 0/45 | 36/63 | 27/55 | 39/78 |
| 411 | C | T | All three | 0/48 | 0/51 | 36/84 | 41/63 | 35/73 |
| 411 | C | G | All three | 0/44 | 0/50 | 20/49 | 31/57 | 41/83 |
| 411 | TG | T | All three | 0/48 | 0/44 | 31/58 | 26/41 | 39/66 |
| 411 | C | T | All three | 0/60 | 0/61 | 29/57 | 32/65 | 34/68 |
| 411 | C | G | All three | 0/58 | 0/40 | 37/68 | 23/60 | 64/94 |
| 716 | G | T | All three | 0/55 | 0/44 | 33/72 | 38/81 | 46/97 |
| 716 | G | T | All three | 0/58 | 0/60 | 34/61 | 34/70 | 34/73 |
| 716 | G | C | All three | 0/55 | 0/54 | 30/59 | 35/74 | 26/61 |
| 716 | A | G | All three | 0/49 | 0/58 | 32/54 | 31/63 | 44/72 |
| 716 | C | T | All three | 0/47 | 0/67 | 22/49 | 36/77 | 34/67 |
| 716 | C | T | All three | 0/63 | 0/59 | 15/37 | 21/47 | 24/50 |

De Novo UTR+Intron Blood

| CDH Proband Family | Sex | Gene | Gene name | Location | Chromosome | Position |
|---|---|---|---|---|---|---|
| 716 | Female | GPR55 | G protein-coupled receptor 55 | intron 1 | 2 | 231787570 |
| 716 | Female | LINC00880 | long intergenic non-protein coding RNA 880 | intron 1 | 3 | 156823715 |
| 716 | Female | ZBBX | zinc finger B-box domain containing | intron 18 | 3 | 167012263 |
| 716 | Female | EIF4G1 | eukaryotic translation initiation factor 4 gamma: 1 | intron 7 | 3 | 184040109 |
| 716 | Female | STX18 | syntaxin 18 | intron 5 | 4 | 4457642 |
| 716 | Female | FLJ13197 | | intron 1 | 4 | 38664228 |
| 716 | Female | STPG2 | sperm-tail PG-rich repeat containing 2 | intron 10 | 4 | 98532709 |
| 716 | Female | CXXC4 | CXXC finger protein 4 | 3 utr | 4 | 105390032 |
| 716 | Female | CDH18 | cadherin 18 | intron 2 | 5 | 20088379 |
| 716 | Female | NR_073113 | | intron 1 | 5 | 43598985 |
| 716 | Female | ANKRD31 | ankyrin repeat domain 31 | intron 4 | 5 | 74504516 |
| 716 | Female | MCC | mutated in colorectal cancers | intron 1 | 5 | 112734744 |
| 716 | Female | KCNIP1 | Kv channel interacting protein 1 | intron 1 | 5 | 170030091 |
| 716 | Female | ANKS1A | ankyrin repeat and sterile alpha motif domain containing 1A | intron 1 | 6 | 34925031 |
| 716 | Female | HMGCLL1 | 3-hydroxymethyl-3-methylglutaryl-CoA lyase-like 1 | intron 2 | 6 | 55434220 |
| 716 | Female | THEMIS | thymocyte selection associated | intron 4 | 6 | 128042029 |
| 716 | Female | AIG1 | androgen-induced 1 | intron 3 | 6 | 143506521 |
| 716 | Female | GRM8 | glutamate receptor: metabotropic 8 | intron 2 | 7 | 126737917 |
| 716 | Female | NEIL2 | nei endonuclease VIII-like 2 (E. coli) | 3 utr | 8 | 11643801 |
| 716 | Female | ANGPT1 | angiopoietin 1 | intron 4 | 8 | 108328503 |
| 716 | Female | KHDRBS3 | omain containing: RNA binding: signal transduction associat | intron 2 | 8 | 136543376 |
| 716 | Female | DOCK8 | dedicator of cytokinesis 8 | intron 18 | 9 | 375626 |
| 716 | Female | ERCC6L2 | oss-complementing rodent repair deficiency: complementat | intron 13 | 9 | 98720169 |
| 716 | Female | MIR3134 | microRNA 3134 | intron 1 | 9 | 114934634 |
| 716 | Female | ADAM12 | ADAM metallopeptidase domain 12 | intron 9 | 10 | 127787410 |
| 716 | female | NELL1 | NEL-like 1 (chicken) | intron 5 | 11 | 20910939 |
| 716 | Female | LRRC4C | leucine rich repeat containing 4C | intron 1 | 11 | 40832247 |
| 716 | Female | CD6 | CD6 molecule | intron 2 | 11 | 60774521 |
| 716 | Female | PDS5B | 5: regulator of cohesion maintenance: homolog B (S. cerevis | intron 1 | 13 | 33169492 |
| 716 | Female | CLYBL | citrate lyase beta like | intron 2 | 13 | 100501587 |
| 716 | Female | abParts | | intron 7 | 15 | 22421168 |

| CDH Proband Family | Reference allele | Alternate allele | Tissue found in | Father | Mother | Proband Blood | Proband Skin | Proband Diaphragm |
|---|---|---|---|---|---|---|---|---|
| 716 | C | A | All three | 0/48 | 0/46 | 37/60 | 22/54 | 32/61 |
| 716 | T | C | All three | 0/40 | 0/59 | 46/82 | 35/76 | 43/90 |
| 716 | G | T | All three | 0/59 | 0/64 | 24/65 | 23/62 | 25/64 |
| 716 | T | A | All three | 0/58 | 0/47 | 39/72 | 34/78 | 45/90 |
| 716 | T | C | All three | 0/40 | 0/42 | 35/67 | 26/59 | 32/64 |
| 716 | T | C | All three | 0/46 | 0/53 | 28/64 | 33/67 | 29/63 |
| 716 | T | C | All three | 0/48 | 0/50 | 32/65 | 39/68 | 35/57 |
| 716 | T | C | All three | 0/56 | 0/47 | 34/71 | 30/65 | 22/61 |
| 716 | T | G | All three | 0/62 | 0/67 | 32/74 | 35/73 | 37/79 |
| 716 | T | C | All three | 0/44 | 0/44 | 36/69 | 33/73 | 39/70 |
| 716 | T | C | All three | 0/47 | 0/54 | 26/60 | 36/69 | 27/63 |
| 716 | G | A | All three | 0/48 | 0/53 | 22/49 | 41/76 | 27/64 |
| 716 | GGATCCCACC | T | All three | 0/51 | 0/66 | 29/59 | 30/76 | 32/58 |
| 716 | T | C | All three | 0/48 | 0/56 | 41/85 | 30/80 | 29/59 |
| 716 | T | C | All three | 0/55 | 0/61 | 30/65 | 41/76 | 34/63 |
| 716 | A | G | All three | 0/43 | 0/48 | 42/71 | 41/75 | 43/68 |
| 716 | C | T | All three | 0/41 | 0/63 | 35/63 | 35/68 | 39/81 |
| 716 | A | G | All three | 0/57 | 0/66 | 46/77 | 25/61 | 35/66 |
| 716 | C | T | All three | 0/59 | 0/55 | 28/55 | 36/69 | 22/54 |
| 716 | T | C | All three | 0/53 | 0/55 | 26/68 | 47/80 | 42/73 |
| 716 | G | T | All three | 0/55 | 0/60 | 40/67 | 23/57 | 34/66 |
| 716 | C | T | All three | 0/51 | 0/57 | 37/66 | 35/64 | 36/68 |
| 716 | T | A | All three | 0/53 | 0/59 | 27/51 | 31/65 | 38/69 |
| 716 | G | A | All three | 0/47 | 0/56 | 41/74 | 35/65 | 28/66 |
| 716 | A | G | All three | 0/61 | 0/60 | 30/71 | 32/72 | 33/68 |
| 716 | A | C | Blood | 0/64 | 0/75 | 19/94 | 4/62 | 4/71 |
| 716 | C | T | All three | 0/49 | 0/46 | 27/61 | 34/62 | 29/57 |
| 716 | G | C | All three | 0/53 | 0/61 | 26/53 | 26/63 | 32/68 |
| 716 | A | G | All three | 0/54 | 0/50 | 35/80 | 24/54 | 39/66 |
| 716 | G | A | All three | 0/52 | 0/53 | 33/59 | 23/58 | 41/73 |
| 716 | C | A | All three | 0/122 | 0/97 | 33/107 | 37/117 | 44/116 |

| CDH Proband Family | Sex | Gene | Gene name | Location | Chromosome | Position |
|---|---|---|---|---|---|---|
| 716 | Female | PEAK1 | pseudopodium-enriched atypical kinase 1 | intron 1 | 15 | 77658986 |
| 716 | Female | ACSBG1 | acyl-CoA synthetase bubblegum family member 1 | intron 2 | 15 | 78499714 |
| 716 | Female | SLCO3A1 | solute carrier organic anion transporter family: member 3A1 | intron 5 | 15 | 92664705 |
| 716 | female | LRRK1 | leucine-rich repeat kinase 1 | 3 utr | 15 | 101611011 |
| 716 | Female | PRKCB | protein kinase C: beta | intron 3 | 16 | 24033310 |
| 716 | Female | CKM | creatine kinase: muscle | intron 4 | 19 | 45818459 |
| 716 | Female | ZNF256 | zinc finger protein 256 | intron 1 | 19 | 58458152 |
| 716 | Female | SIRPA | signal-regulatory protein alpha | intron 2 | 20 | 1890924 |
| 716 | Female | PTPRT | protein tyrosine phosphatase: receptor type: T | intron 2 | 20 | 41496654 |
| 716 | Female | APCDD1L | adenomatosis polyposis coli down-regulated 1-like | intron 1 | 20 | 57048292 |
| 716 | Female | SYCP2 | synaptonemal complex protein 2 | intron 11 | 20 | 58488059 |
| 716 | Female | SGSM1 | small G protein signaling modulator 1 | intron 2 | 22 | 25212931 |
| 716 | Female | TBC1D22A | TBC1 domain family: member 22A | intron 12 | 22 | 47565909 |
| 716 | Female | HEPH | hephaestin | intron 2 | X | 65391234 |
| 809 | Female | SORCS3 | sortilin-related VPS10 domain containing receptor 3 | intron 5 | 1 | 10687743 |
| 809 | Female | AFTPH | aftiphilin | intron 7 | 2 | 64808306 |
| 809 | Female | SFTPB | surfactant protein B | 3 utr | 2 | 85884898 |
| 809 | Female | PLA2R1 | phospholipase A2 receptor 1: 180kDa | intron 11 | 2 | 160851223 |
| 809 | Female | LINC00607 | long intergenic non-protein coding RNA 607 | intron 5 | 2 | 216549095 |
| 809 | Female | SLC9A9 | ier family 9: subfamily A (NHE9: cation proton antiporter 9): | intron 4 | 3 | 143433614 |
| 809 | Female | KCTD8 | potassium channel tetramerization domain containing 8 | intron 1 | 4 | 44359274 |
| 809 | female | EDIL3 | EGF-like repeats and discoidin I-like domains 3 | intron 6 | 5 | 83397673 |
| 809 | Female | FBXL17 | F-box and leucine-rich repeat protein 17 | intron 6 | 5 | 107485066 |
| 809 | Female | GRAMD3 | GRAM domain containing 3 | intron 12 | 5 | 125823743 |
| 809 | Female | EBF1 | early B-cell factor 1 | intron 9 | 5 | 158210489 |
| 809 | Female | TDP2 | tyrosyl-DNA phosphodiesterase 2 | intron 3 | 6 | 24658388 |
| 809 | Female | EYS | eyes shut homolog (Drosophila) | intron 12 | 6 | 65845477 |
| 809 | Female | COL12A1 | collagen: type XII: alpha 1 | intron 45 | 6 | 75829061 |
| 809 | Female | PREP | prolyl endopeptidase | intron 10 | 6 | 105750309 |
| 809 | female | FAM184A | family with sequence similarity 184: member A | intron 1 | 6 | 119395836 |
| 809 | Female | STEAP1B | STEAP family member 1B | intron 5 | 7 | 22475570 |

**TABLE S3**

| CDH Proband Family | Reference allele | Alternate allele | Tissue found in | Father | Mother | Proband Blood | Proband Skin | Proband Diaphragm |
|---|---|---|---|---|---|---|---|---|
| 716 | C | T | All three | 0/50 | 0/50 | 38/63 | 37/69 | 34/58 |
| 716 | T | C | All three | 0/66 | 0/65 | 34/56 | 41/82 | 35/69 |
| 716 | T | C | All three | 0/40 | 0/46 | 39/74 | 35/76 | 41/73 |
| 716 | A | T | All three | 0/64 | 0/53 | 25/55 | 31/66 | 23/65 |
| 716 | C | A | All three | 0/39 | 0/62 | 25/68 | 35/76 | 33/62 |
| 716 | C | T | All three | 0/58 | 0/52 | 30/66 | 34/85 | 36/61 |
| 716 | T | C | All three | 0/50 | 0/55 | 42/85 | 45/87 | 30/72 |
| 716 | T | G | All three | 0/47 | 0/53 | 34/56 | 31/59 | 37/76 |
| 716 | A | G | All three | 0/55 | 0/50 | 31/62 | 42/82 | 32/66 |
| 716 | C | CT | All three | 0/58 | 0/46 | 33/58 | 38/76 | 36/56 |
| 716 | TCA | T | All three | 0/47 | 0/57 | 38/69 | 39/71 | 32/57 |
| 716 | T | C | All three | 0/62 | 0/57 | 39/79 | 39/69 | 34/67 |
| 716 | T | C | All three | 0/54 | 0/47 | 39/67 | 32/61 | 33/68 |
| 716 | T | C | All three | 0/36 | 0/54 | 30/61 | 34/75 | 22/51 |
| 809 | C | T | All three | 0/43 | 0/47 | 44/68 | 30/59 | 36/75 |
| 809 | C | T | All three | 0/42 | 0/55 | 30/68 | 37/72 | 47/87 |
| 809 | T | G | All three | 0/46 | 0/61 | 29/58 | 29/59 | 38/73 |
| 809 | G | A | All three | 0/45 | 0/65 | 39/67 | 37/85 | 44/85 |
| 809 | G | A | All three | 0/50 | 0/64 | 38/85 | 40/69 | 30/66 |
| 809 | T | C | All three | 0/47 | 0/57 | 38/70 | 32/59 | 36/72 |
| 809 | G | A | All three | 0/52 | 0/66 | 45/77 | 35/83 | 32/80 |
| 809 | G | A | All three | 0/62 | 0/81 | 37/96 | 37/82 | 29/57 |
| 809 | G | A | All three | 0/41 | 0/62 | 30/74 | 41/76 | 40/82 |
| 809 | T | C | All three | 0/51 | 0/61 | 41/64 | 35/64 | 35/88 |
| 809 | A | G | All three | 0/42 | 0/53 | 37/78 | 32/74 | 49/88 |
| 809 | T | A | All three | 0/44 | 0/58 | 40/62 | 34/76 | 31/71 |
| 809 | G | T | All three | 0/51 | 0/76 | 37/83 | 28/59 | 36/81 |
| 809 | C | T | All three | 0/44 | 0/59 | 37/89 | 33/65 | 33/68 |
| 809 | T | C | All three | 0/47 | 0/56 | 37/77 | 34/60 | 31/76 |
| 809 | C | G | Skin+Blood | 0/55 | 0/68 | 19/79 | 25/81 | 11/88 |
| 809 | TG | T | All three | 0/45 | 0/34 | 27/58 | 46/81 | 39/71 |

**TABLE S3**

| CDH Proband Family | Sex | Gene | Gene name | Location | Chromosome | Position |
|---|---|---|---|---|---|---|
| 809 | Female | CACNA2D1 | :alcium channel: voltage-dependent: alpha 2/delta subunit 1 | intron 3 | 7 | 81834306 |
| 809 | Female | PTPRD | protein tyrosine phosphatase: receptor type: D | intron 8 | 9 | 9448343 |
| 809 | Female | PTPRD | protein tyrosine phosphatase: receptor type: D | intron 7 | 9 | 9600668 |
| 809 | Female | UGCG | UDP-glucose ceramide glucosyltransferase | intron 1 | 9 | 114666708 |
| 809 | Female | ST5 | suppression of tumorigenicity 5 | intron 1 | 11 | 8822290 |
| 809 | Female | LRRC4C | leucine rich repeat containing 4C | intron 1 | 11 | 40749101 |
| 809 | Female | ALG9 | ALG9: alpha-1:2-mannosyltransferase | intron 9 | 11 | 111723240 |
| 809 | Female | KIRREL3 | kin of IRRE like 3 (Drosophila) | intron 1 | 11 | 126775175 |
| 809 | Female | ATP2A2 | ATPase: Ca++ transporting: cardiac muscle: slow twitch 2 | intron 5 | 12 | 110734883 |
| 809 | Female | NPAS3 | neuronal PAS domain protein 3 | intron 3 | 14 | 33833460 |
| 809 | Female | PRKD2 | Protein Kinase D2 | intron 2 | 19 | 47216261 |
| 967 | Male | SPATA6 | spermatogenesis associated 6 | intron 11 | 1 | 48785688 |
| 967 | Male | LRRC8C | leucine rich repeat containing 8 family: member C | intron 1 | 1 | 90115215 |
| 967 | Male | MERTK | c-mer proto-oncogene tyrosine kinase | intron 8 | 2 | 112744068 |
| 967 | Male | THSD7B | thrombospondin: type I: domain containing 7B | intron 13 | 2 | 138202217 |
| 967 | Male | SCHIP1 | Schwannomin Interacting Protein 1 | intron 2 | 3 | 159529771 |
| 967 | Male | NAALADL2 | N-acetylated alpha-linked acidic dipeptidase-like 2 | intron 9 | 3 | 175240124 |
| 967 | Male | BC005018 | | intron 4 | 4 | 84733093 |
| 967 | Male | PDZD2 | PDZ domain containing 2 | intron 2 | 5 | 31946007 |
| 967 | Male | DQ515898 | | intron 1 | 8 | 128336350 |
| 967 | Male | ANTD3-TMEI | MSANTD3-TMEFF1 readthrough | intron 1 | 9 | 103248140 |
| 967 | Male | NELFB | negative elongation factor complex member B | intron 4 | 9 | 140151750 |
| 967 | Male | RSU1 | Ras suppressor protein 1 | intron 1 | 10 | 16830733 |
| 967 | Male | SUFU | suppressor of fused homolog (Drosophila) | intron 2 | 10 | 104283460 |
| 967 | Male | OR8U8 | olfactory receptor family 8 subfamily U member 8 | intron 1 | 11 | 56202375 |
| 967 | male | DISC1FP1 | DISC1 fusion partner 1 (non-protein coding) | intron 1 | 11 | 90243703 |
| 967 | Male | UGGT2 | UDP-glucose glycoprotein glucosyltransferase 2 | intron 4 | 13 | 96673108 |
| 967 | Male | MDGA2 | IAM domain containing glycosylphosphatidylinositol anchor | intron 1 | 14 | 47897999 |
| 967 | Male | ıCG_2003567 | | intron 1 | 15 | 66926125 |
| 967 | Male | CD33 | CD33 molecule | intron 7 | 19 | 51739471 |
| 967 | Male | TBX1 | T-box 1 | intron 8 | 22 | 19768601 |

**TABLE S3**

| CDH Proband Family | Reference allele | Alternate allele | Tissue found in | Father | Mother | Proband Blood | Proband Skin | Proband Diaphragm |
|---|---|---|---|---|---|---|---|---|
| 809 | G | A | All three | 0/54 | 0/65 | 28/56 | 32/69 | 25/55 |
| 809 | G | T | All three | 0/47 | 0/65 | 39/66 | 40/63 | 40/77 |
| 809 | G | A | All three | 0/44 | 0/67 | 45/76 | 41/79 | 35/66 |
| 809 | C | T | All three | 0/37 | 0/54 | 47/87 | 37/76 | 38/63 |
| 809 | T | G | All three | 0/50 | 0/54 | 32/73 | 43/78 | 38/79 |
| 809 | AT | A | All three | 0/38 | 0/61 | 49/85 | 30/59 | 43/69 |
| 809 | G | A | All three | 0/52 | 0/55 | 30/58 | 32/60 | 37/71 |
| 809 | G | A | All three | 0/60 | 0/64 | 31/61 | 35/67 | 36/73 |
| 809 | C | T | All three | 0/42 | 0/69 | 26/67 | 37/67 | 36/60 |
| 809 | A | G | All three | 0/42 | 0/52 | 39/71 | 27/67 | 30/60 |
| 809 | T | [illegible]TTTTCTTTTT[illegible] | All three | 0/48 | 0/69 | 15/30 | 13/36 | 29/43 |
| 967 | G | A | All three | 0/47 | 0/75 | 18/38 | 25/55 | 27/48 |
| 967 | C | T | All three | 0/47 | 0/58 | 29/52 | 23/54 | 35/52 |
| 967 | C | T | All three | 0/69 | 0/60 | 28/67 | 38/78 | 16/48 |
| 967 | CCT | C | All three | 0/54 | 0/55 | 21/40 | 25/65 | 14/30 |
| 967 | T | TTG | All three | 0/56 | 0/69 | 26/40 | 35/59 | 19/28 |
| 967 | C | T | All three | 0/47 | 0/62 | 33/63 | 22/67 | 24/57 |
| 967 | C | T | All three | 0/42 | 0/53 | 27/49 | 25/55 | 24/45 |
| 967 | T | C | All three | 0/43 | 0/43 | 22/49 | 26/53 | 29/47 |
| 967 | T | C | All three | 0/52 | 0/64 | 24/52 | 27/66 | 13/36 |
| 967 | T | G | All three | 0/54 | 0/59 | 23/43 | 39/65 | 16/46 |
| 967 | C | T | All three | 0/54 | 0/44 | 26/57 | 31/69 | 20/40 |
| 967 | G | A | All three | 0/44 | 0/58 | 35/64 | 36/65 | 22/48 |
| 967 | G | A | All three | 0/53 | 0/57 | 20/43 | 26/64 | 24/50 |
| 967 | A | G | All three | 0/42 | 0/49 | 23/49 | 34/65 | 12/36 |
| 967 | TATGC | T | All three | 1/25 | 0/43 | 12/22 | 10/24 | 13/16 |
| 967 | A | G | All three | 0/50 | 0/59 | 42/68 | 22/58 | 32/60 |
| 967 | T | C | All three | 0/62 | 0/52 | 31/60 | 30/59 | 22/47 |
| 967 | A | C | All three | 0/42 | 0/54 | 14/36 | 36/71 | 20/45 |
| 967 | G | T | All three | 0/50 | 0/62 | 20/49 | 38/61 | 18/47 |
| 967 | G | A | All three | 0/55 | 0/50 | 20/40 | 28/54 | 20/41 |

**TABLE S3**

De Novo Intergenic Diaphragm

| CDH Proband Family | Sex | Location | Chromosome | Position | Reference allele | Alternate allele | Tissue found in | Father |
|---|---|---|---|---|---|---|---|---|
| 411 | Male | Intergenic | 2 | 6448411 | G | A | All three | 0/51 |
| 411 | Male | Intergenic | 2 | 15979430 | C | T | All three | 0/46 |
| 411 | Male | Intergenic | 2 | 23607772 | C | A | All three | 0/34 |
| 411 | Male | Intergenic | 2 | 23607823 | G | A | All three | 0/34 |
| 411 | Male | Intergenic | 2 | 48488807 | CTTTGAAAG | T | All three | 0/55 |
| 411 | male | Intergenic | 4 | 12542452 | ACAAAAAT | A | All three | 0/61 |
| 411 | Male | Intergenic | 4 | 40268400 | A | G | All three | 0/59 |
| 411 | Male | Intergenic | 4 | 130485536 | C | T | All three | 0/39 |
| 411 | Male | Intergenic | 5 | 30972421 | G | T | All three | 0/54 |
| 411 | Male | Intergenic | 5 | 43876751 | A | T | All three | 0/43 |
| 411 | Male | Intergenic | 7 | 121415957 | T | G | All three | 0/42 |
| 411 | Male | Intergenic | 8 | 49656228 | G | C | All three | 0/45 |
| 411 | Male | Intergenic | 9 | 24573667 | A | C | All three | 0/57 |
| 411 | Male | Intergenic | 10 | 45696971 | C | T | All three | 0/44 |
| 411 | male | Intergenic | 10 | 107240148 | T | C | All three | 0/54 |
| 411 | Male | Intergenic | 11 | 48522238 | T | C | All three | 0/53 |
| 411 | Male | Intergenic | 11 | 68646620 | CCT | C | Diaphragm | 0/44 |
| 411 | Male | Intergenic | 12 | 51286910 | T | C | All three | 0/46 |
| 411 | Male | Intergenic | 13 | 63562946 | G | GA | All three | 0/63 |
| 411 | Male | Intergenic | 15 | 38941044 | C | T | All three | 0/45 |
| 411 | Male | Intergenic | 15 | 46407038 | C | T | All three | 0/55 |
| 411 | Male | Intergenic | 16 | 75543055 | C | A | All three | 0/46 |
| 411 | Male | Intergenic | 17 | 404337 | C | T | All three | 0/55 |
| 411 | Male | Intergenic | 20 | 5206920 | C | A | All three | 0/45 |
| 411 | Male | Intergenic | X | 50778530 | ACATTTCTCA | G | Diaphragm | 0/18 |
| 411 | Male | Intergenic | X | 117269505 | CGT | C | Diaphragm | 0/43 |
| 411 | Male | Intergenic | Y | 23424401 | T | C | All three | 0/37 |
| 716 | Female | Intergenic | 1 | 25520943 | G | A | All three | 0/53 |
| 716 | Female | Intergenic | 1 | 108550027 | G | A | All three | 0/55 |
| 716 | Female | Intergenic | 1 | 188119862 | C | T | All three | 0/60 |

**TABLE S3**

| CDH Proband Family | Mother | Proband Blood | Proband Skin | Proband Diaphragm |
|---|---|---|---|---|
| 411 | 0/49 | 29/59 | 21/54 | 51/89 |
| 411 | 0/38 | 16/47 | 23/50 | 36/63 |
| 411 | 0/39 | 16/26 | 25/40 | 23/56 |
| 411 | 0/47 | 16/21 | 17/25 | 17/49 |
| 411 | 0/46 | 23/50 | 22/60 | 32/70 |
| 411 | 0/67 | 23/54 | 23/50 | 32/86 |
| 411 | 0/56 | 36/75 | 30/55 | 38/78 |
| 411 | 0/55 | 34/67 | 23/58 | 36/75 |
| 411 | 0/52 | 28/65 | 34/65 | 41/99 |
| 411 | 0/43 | 37/78 | 32/59 | 21/55 |
| 411 | 0/50 | 25/62 | 26/61 | 42/81 |
| 411 | 0/47 | 31/57 | 21/58 | 42/75 |
| 411 | 0/45 | 24/69 | 18/48 | 32/78 |
| 411 | 0/59 | 33/70 | 31/65 | 29/60 |
| 411 | 0/58 | 32/79 | 25/58 | 33/71 |
| 411 | 0/41 | 26/61 | 24/69 | 35/85 |
| 411 | 0/50 | 0/53 | 0/44 | 7/35 |
| 411 | 0/47 | 57/97 | 34/58 | 35/80 |
| 411 | 0/51 | 19/48 | 20/40 | 33/68 |
| 411 | 0/48 | 28/58 | 31/72 | 37/75 |
| 411 | 0/43 | 34/58 | 22/57 | 36/77 |
| 411 | 0/43 | 22/56 | 30/53 | 41/79 |
| 411 | 0/50 | 36/73 | 28/49 | 38/77 |
| 411 | 0/43 | 26/57 | 26/67 | 35/77 |
| 411 | 0/39 | 0/25 | 0/19 | 16/71 |
| 411 | 0/46 | 0/35 | 0/27 | 7/35 |
| 411 | 0/0 | 35/35 | 35/36 | 31/31 |
| 716 | 0/58 | 24/50 | 40/84 | 23/51 |
| 716 | 0/58 | 38/66 | 36/64 | 33/66 |
| 716 | 0/65 | 27/61 | 38/66 | 23/56 |

**TABLE S3**

| CDH Proband Family | Sex | Location | Chromosome | Position | Reference allele | Alternate allele | Tissue found in | Father |
|---|---|---|---|---|---|---|---|---|
| 716 | female | Intergenic | 1 | 190952170 | A | G | Diaphragm+Skin | 0/20 |
| 716 | Female | Intergenic | 1 | 219133910 | G | A | All three | 0/54 |
| 716 | Female | Intergenic | 1 | 241538742 | A | G | All three | 0/64 |
| 716 | Female | Intergenic | 2 | 60131252 | T | C | All three | 0/56 |
| 716 | Female | Intergenic | 2 | 237733339 | C | T | All three | 0/48 |
| 716 | Female | Intergenic | 2 | 241273592 | G | T | All three | 0/48 |
| 716 | Female | Intergenic | 3 | 31317266 | C | T | All three | 0/65 |
| 716 | Female | Intergenic | 3 | 70707930 | T | C | All three | 0/51 |
| 716 | Female | Intergenic | 3 | 87755171 | C | T | All three | 0/51 |
| 716 | Female | Intergenic | 3 | 96134789 | G | A | All three | 0/58 |
| 716 | Female | Intergenic | 3 | 116969700 | C | T | All three | 0/61 |
| 716 | Female | Intergenic | 4 | 76738805 | A | C | All three | 0/55 |
| 716 | Female | Intergenic | 4 | 90360867 | C | G | All three | 0/45 |
| 716 | Female | Intergenic | 4 | 92765694 | T | A | All three | 0/55 |
| 716 | Female | Intergenic | 4 | 165797047 | C | T | All three | 0/49 |
| 716 | Female | Intergenic | 5 | 13079144 | A | T | All three | 0/45 |
| 716 | Female | Intergenic | 5 | 92381616 | C | G | All three | 0/62 |
| 716 | Female | Intergenic | 6 | 18809404 | A | C | All three | 0/66 |
| 716 | Female | Intergenic | 6 | 50605045 | A | G | All three | 0/57 |
| 716 | Female | Intergenic | 6 | 69243495 | C | G | All three | 0/41 |
| 716 | Female | Intergenic | 6 | 78773081 | C | T | All three | 0/62 |
| 716 | Female | Intergenic | 7 | 2657275 | C | T | All three | 0/47 |
| 716 | Female | Intergenic | 7 | 6718899 | A | G | All three | 0/68 |
| 716 | Female | Intergenic | 7 | 19285491 | C | T | All three | 0/50 |
| 716 | Female | Intergenic | 7 | 96000892 | G | T | All three | 0/46 |
| 716 | Female | Intergenic | 8 | 23549713 | A | G | All three | 0/50 |
| 716 | Female | Intergenic | 8 | 50342185 | C | A | All three | 0/55 |
| 716 | Female | Intergenic | 8 | 53690025 | G | A | All three | 0/65 |
| 716 | Female | Intergenic | 8 | 121394186 | C | G | All three | 0/43 |
| 716 | Female | Intergenic | 9 | 37870079 | C | A | All three | 0/52 |

| CDH Proband Family | Mother | Proband Blood | Proband Skin | Proband Diaphragm |
|---|---|---|---|---|
| 716 | 0/44 | 3/25 | 7/27 | 4/16 |
| 716 | 0/50 | 37/69 | 28/56 | 29/66 |
| 716 | 0/58 | 40/85 | 46/78 | 33/67 |
| 716 | 0/57 | 28/55 | 43/90 | 49/79 |
| 716 | 0/51 | 29/68 | 32/76 | 39/74 |
| 716 | 0/54 | 42/89 | 30/77 | 26/55 |
| 716 | 0/56 | 34/61 | 26/59 | 39/77 |
| 716 | 0/50 | 26/62 | 32/61 | 35/66 |
| 716 | 0/59 | 35/82 | 34/67 | 38/66 |
| 716 | 0/60 | 31/58 | 46/81 | 23/58 |
| 716 | 0/72 | 35/79 | 37/71 | 23/52 |
| 716 | 0/56 | 26/54 | 21/45 | 45/67 |
| 716 | 0/61 | 29/59 | 25/53 | 29/60 |
| 716 | 0/46 | 35/69 | 40/63 | 31/63 |
| 716 | 0/57 | 39/68 | 34/69 | 39/72 |
| 716 | 0/59 | 27/60 | 36/63 | 28/51 |
| 716 | 0/59 | 40/72 | 27/57 | 32/57 |
| 716 | 0/68 | 30/68 | 33/77 | 22/47 |
| 716 | 0/53 | 39/75 | 28/58 | 36/56 |
| 716 | 0/61 | 26/53 | 31/72 | 35/69 |
| 716 | 0/58 | 38/79 | 36/62 | 38/76 |
| 716 | 0/57 | 30/58 | 23/54 | 29/67 |
| 716 | 0/55 | 34/75 | 35/78 | 34/68 |
| 716 | 0/63 | 36/70 | 38/73 | 41/77 |
| 716 | 0/52 | 23/83 | 19/64 | 29/71 |
| 716 | 0/47 | 17/55 | 34/68 | 37/76 |
| 716 | 0/55 | 21/47 | 42/77 | 48/85 |
| 716 | 0/54 | 25/44 | 26/66 | 30/58 |
| 716 | 0/43 | 20/50 | 38/72 | 31/54 |
| 716 | 0/52 | 29/68 | 31/71 | 35/66 |

De Novo Intergenic Diaphragm

| CDH Proband Family | Sex | Location | Chromosome | Position | Reference allele | Alternate allele | Tissue found in | Father |
|---|---|---|---|---|---|---|---|---|
| 716 | Female | Intergenic | 9 | 89980836 | A | G | Diaphragm | 0/52 |
| 716 | Female | Intergenic | 9 | 92940311 | C | T | All three | 0/62 |
| 716 | Female | Intergenic | 10 | 21531548 | A | T | All three | 0/61 |
| 716 | female | Intergenic | 10 | 78610342 | GACAA | G | All three | 0/79 |
| 716 | Female | Intergenic | 10 | 102805027 | C | T | All three | 0/49 |
| 716 | Female | Intergenic | 10 | 111937754 | A | G | All three | 0/55 |
| 716 | Female | Intergenic | 11 | 68910808 | C | T | All three | 0/52 |
| 716 | Female | Intergenic | 11 | 68917454 | A | T | All three | 0/63 |
| 716 | Female | Intergenic | 11 | 85940710 | C | T | All three | 0/52 |
| 716 | Female | Intergenic | 12 | 2894972 | G | A | All three | 0/71 |
| 716 | Female | Intergenic | 12 | 20253658 | G | A | All three | 0/60 |
| 716 | Female | Intergenic | 12 | 61786788 | C | T | All three | 0/56 |
| 716 | female | Intergenic | 12 | 73894281 | T | C | All three | 0/67 |
| 716 | Female | Intergenic | 12 | 81133670 | T | C | All three | 0/44 |
| 716 | Female | Intergenic | 12 | 128640569 | T | C | All three | 0/60 |
| 716 | Female | Intergenic | 13 | 38770384 | A | T | All three | 0/56 |
| 716 | Female | Intergenic | 14 | 94631627 | C | T | All three | 0/47 |
| 716 | Female | Intergenic | 14 | 98723131 | C | T | All three | 0/57 |
| 716 | Female | Intergenic | 15 | 53418167 | A | G | All three | 0/59 |
| 716 | Female | Intergenic | 18 | 26978994 | C | T | All three | 0/53 |
| 716 | Female | Intergenic | 18 | 48313755 | G | T | All three | 0/60 |
| 716 | Female | Intergenic | 18 | 63851839 | G | C | All three | 0/55 |
| 716 | Female | Intergenic | 19 | 12407418 | A | C | All three | 0/78 |
| 716 | Female | Intergenic | 20 | 40498713 | A | T | All three | 0/41 |
| 716 | Female | Intergenic | 21 | 26441156 | G | T | All three | 0/53 |
| 716 | Female | Intergenic | 21 | 29172156 | A | T | All three | 0/59 |
| 716 | Female | Intergenic | X | 39689788 | C | G | All three | 0/31 |
| 716 | Female | Intergenic | X | 65140289 | G | A | All three | 0/25 |
| 716 | Female | Intergenic | X | 82221769 | C | T | All three | 0/30 |
| 716 | Female | Intergenic | X | 88110135 | C | T | All three | 0/34 |

**TABLE S3**

| CDH Proband Family | Mother | Proband Blood | Proband Skin | Proband Diaphragm |
|---|---|---|---|---|
| 716 | 0/59 | 0/54 | 7/59 | 18/78 |
| 716 | 0/56 | 28/66 | 37/73 | 38/77 |
| 716 | 0/56 | 33/61 | 37/73 | 30/61 |
| 716 | 0/80 | 38/79 | 35/88 | 45/91 |
| 716 | 0/49 | 40/77 | 45/80 | 36/73 |
| 716 | 0/47 | 38/77 | 43/81 | 43/68 |
| 716 | 0/59 | 34/72 | 36/71 | 27/59 |
| 716 | 0/52 | 40/72 | 35/69 | 36/65 |
| 716 | 0/53 | 28/63 | 38/76 | 31/57 |
| 716 | 0/58 | 47/80 | 31/61 | 38/76 |
| 716 | 0/64 | 39/77 | 29/50 | 43/61 |
| 716 | 0/74 | 36/76 | 34/67 | 36/61 |
| 716 | 0/77 | 43/81 | 40/73 | 33/71 |
| 716 | 0/63 | 37/65 | 38/72 | 30/70 |
| 716 | 0/64 | 22/57 | 38/73 | 35/76 |
| 716 | 0/42 | 28/57 | 30/63 | 32/61 |
| 716 | 0/61 | 30/55 | 31/66 | 23/44 |
| 716 | 0/55 | 34/65 | 32/66 | 30/67 |
| 716 | 0/60 | 24/67 | 30/66 | 35/66 |
| 716 | 0/49 | 28/62 | 34/74 | 38/74 |
| 716 | 0/53 | 35/64 | 33/71 | 40/74 |
| 716 | 0/54 | 38/76 | 41/83 | 33/67 |
| 716 | 0/74 | 46/68 | 28/54 | 44/62 |
| 716 | 0/46 | 28/68 | 37/68 | 25/59 |
| 716 | 0/58 | 27/62 | 41/73 | 28/59 |
| 716 | 0/62 | 44/74 | 34/84 | 30/58 |
| 716 | 0/57 | 31/66 | 24/54 | 34/81 |
| 716 | 0/47 | 40/79 | 32/68 | 31/70 |
| 716 | 0/55 | 39/68 | 37/69 | 34/54 |
| 716 | 0/63 | 35/64 | 36/69 | 35/68 |

TABLE S3

De Novo Intergenic Diaphragm

| CDH Proband Family | Sex | Location | Chromosome | Position | Reference allele | Alternate allele | Tissue found in | Father |
|---|---|---|---|---|---|---|---|---|
| 716 | Female | Intergenic | X | 89554886 | C | T | All three | 0/25 |
| 716 | Female | Intergenic | X | 105821982 | G | T | All three | 0/35 |
| 809 | Female | Intergenic | 1 | 107022943 | G | T | Diaphragm | 0/38 |
| 809 | Female | Intergenic | 1 | 159684393 | C | T | Diaphragm | 0/36 |
| 809 | Female | Intergenic | 1 | 229943691 | T | A | All three | 0/41 |
| 809 | Female | Intergenic | 1 | 248631521 | G | A | All three | 0/47 |
| 809 | female | Intergenic | 2 | 52087068 | T | A | Diaphragm+Blood | 1/49 |
| 809 | Female | Intergenic | 2 | 53069390 | G | A | Diaphragm | 0/44 |
| 809 | Female | Intergenic | 2 | 106320315 | C | G | Diaphragm | 0/36 |
| 809 | Female | Intergenic | 2 | 118456374 | T | C | All three | 0/46 |
| 809 | Female | Intergenic | 2 | 126870837 | T | A | Diaphragm | 0/46 |
| 809 | Female | Intergenic | 2 | 170283195 | C | T | Diaphragm | 0/47 |
| 809 | Female | Intergenic | 2 | 184254426 | A | T | Diaphragm | 0/42 |
| 809 | Female | Intergenic | 2 | 185100776 | C | A | Diaphragm | 0/44 |
| 809 | Female | Intergenic | 2 | 212004910 | C | A | Diaphragm | 0/46 |
| 809 | Female | Intergenic | 3 | 74259109 | A | G | All three | 0/39 |
| 809 | Female | Intergenic | 3 | 74801621 | G | A | Diaphragm | 0/42 |
| 809 | Female | Intergenic | 3 | 96502668 | T | A | Diaphragm | 0/45 |
| 809 | Female | Intergenic | 3 | 118151660 | G | T | Diaphragm | 0/58 |
| 809 | Female | Intergenic | 3 | 162105735 | C | G | Diaphragm | 0/43 |
| 809 | Female | Intergenic | 3 | 172322331 | C | T | Diaphragm | 0/48 |
| 809 | Female | Intergenic | 4 | 8715501 | G | A | All three | 0/44 |
| 809 | Female | Intergenic | 4 | 23219247 | C | T | Diaphragm | 0/48 |
| 809 | Female | Intergenic | 4 | 30105753 | G | T | Diaphragm | 0/35 |
| 809 | Female | Intergenic | 4 | 59397258 | C | T | All three | 0/48 |
| 809 | Female | Intergenic | 4 | 64527308 | T | G | Diaphragm | 0/42 |
| 809 | Female | Intergenic | 4 | 118404267 | C | T | Diaphragm | 0/50 |
| 809 | Female | Intergenic | 4 | 139479057 | C | G | Diaphragm | 0/43 |
| 809 | Female | Intergenic | 4 | 176517204 | G | T | Diaphragm | 0/49 |
| 809 | Female | Intergenic | 5 | 4375226 | C | T | Diaphragm | 0/42 |

**TABLE S3**

| CDH Proband Family | Mother | Proband Blood | Proband Skin | Proband Diaphragm |
|---|---|---|---|---|
| 716 | 0/44 | 34/72 | 25/57 | 35/82 |
| 716 | 0/48 | 35/68 | 39/66 | 33/61 |
| 809 | 0/60 | 0/68 | 0/57 | 22/90 |
| 809 | 0/58 | 0/59 | 0/65 | 18/68 |
| 809 | 0/51 | 27/73 | 43/70 | 29/61 |
| 809 | 0/52 | 26/60 | 35/61 | 40/78 |
| 809 | 0/88 | 19/64 | 4/48 | 37/79 |
| 809 | 0/60 | 0/52 | 0/53 | 15/69 |
| 809 | 0/65 | 0/60 | 0/62 | 24/77 |
| 809 | 0/56 | 26/65 | 35/71 | 39/71 |
| 809 | 0/56 | 0/53 | 0/42 | 20/73 |
| 809 | 0/50 | 0/41 | 0/51 | 18/69 |
| 809 | 0/59 | 0/65 | 0/57 | 31/87 |
| 809 | 0/62 | 0/58 | 0/61 | 24/89 |
| 809 | 0/71 | 0/67 | 0/68 | 27/74 |
| 809 | 0/59 | 34/70 | 33/59 | 30/64 |
| 809 | 0/52 | 0/54 | 0/58 | 23/73 |
| 809 | 0/59 | 0/64 | 0/59 | 24/82 |
| 809 | 0/72 | 0/62 | 0/64 | 19/85 |
| 809 | 0/57 | 0/66 | 0/72 | 25/65 |
| 809 | 0/48 | 0/52 | 0/48 | 24/73 |
| 809 | 0/58 | 42/69 | 40/76 | 33/73 |
| 809 | 0/73 | 0/51 | 0/79 | 13/65 |
| 809 | 0/49 | 0/41 | 0/50 | 24/75 |
| 809 | 0/77 | 41/70 | 43/67 | 31/58 |
| 809 | 0/64 | 0/59 | 0/51 | 22/70 |
| 809 | 0/79 | 0/60 | 0/66 | 22/69 |
| 809 | 0/65 | 0/54 | 0/54 | 16/74 |
| 809 | 0/63 | 0/65 | 0/59 | 24/74 |
| 809 | 0/64 | 0/68 | 0/53 | 20/76 |

**TABLE S3**

| CDH Proband Family | Sex | Location | Chromosome | Position | Reference allele | Alternate allele | Tissue found in | Father |
|---|---|---|---|---|---|---|---|---|
| 809 | Female | Intergenic | 5 | 13302657 | G | A | Diaphragm | 0/46 |
| 809 | Female | Intergenic | 5 | 24990895 | C | T | Diaphragm | 0/44 |
| 809 | Female | Intergenic | 5 | 45771487 | G | A | Diaphragm | 0/39 |
| 809 | Female | Intergenic | 5 | 117288082 | G | T | Diaphragm | 0/43 |
| 809 | female | Intergenic | 5 | 132509196 | T | C | All three | 0/59 |
| 809 | Female | Intergenic | 5 | 144084315 | G | GA | All three | 0/40 |
| 809 | Female | Intergenic | 5 | 161028079 | G | A | Diaphragm | 0/53 |
| 809 | Female | Intergenic | 5 | 163064447 | G | A | Diaphragm | 0/46 |
| 809 | Female | Intergenic | 6 | 84499729 | T | C | All three | 0/43 |
| 809 | Female | Intergenic | 7 | 46969823 | T | A | All three | 0/39 |
| 809 | Female | Intergenic | 7 | 56693729 | A | T | Diaphragm | 0/45 |
| 809 | Female | Intergenic | 7 | 79427437 | G | A | Diaphragm | 0/43 |
| 809 | Female | Intergenic | 7 | 93895381 | G | T | Diaphragm | 0/40 |
| 809 | Female | Intergenic | 7 | 155849943 | G | T | Diaphragm | 0/42 |
| 809 | Female | Intergenic | 8 | 24834454 | C | T | All three | 0/50 |
| 809 | Female | Intergenic | 8 | 30173080 | C | T | Diaphragm | 0/56 |
| 809 | Female | Intergenic | 8 | 60040127 | C | T | All three | 0/47 |
| 809 | Female | Intergenic | 9 | 24832115 | G | A | Diaphragm | 0/45 |
| 809 | Female | Intergenic | 10 | 4733230 | A | G | All three | 0/44 |
| 809 | female | Intergenic | 10 | 9768107 | G | C | Diaphragm | 0/68 |
| 809 | Female | Intergenic | 10 | 29015321 | T | C | Diaphragm | 0/49 |
| 809 | Female | Intergenic | 10 | 52428713 | T | C | Diaphragm | 0/44 |
| 809 | Female | Intergenic | 10 | 91874832 | T | A | Diaphragm | 0/55 |
| 809 | Female | Intergenic | 11 | 15343128 | AT | A | All three | 0/39 |
| 809 | Female | Intergenic | 11 | 37638703 | G | C | All three | 0/55 |
| 809 | Female | Intergenic | 11 | 38424427 | C | A | Diaphragm | 0/40 |
| 809 | Female | Intergenic | 11 | 39193391 | C | T | Diaphragm | 0/40 |
| 809 | Female | Intergenic | 11 | 61888225 | G | A | Diaphragm | 0/44 |
| 809 | Female | Intergenic | 11 | 73309897 | C | A | Diaphragm | 0/35 |
| 809 | Female | Intergenic | 11 | 112170508 | T | C | All three | 0/45 |

De Novo Intergenic Diaphragm

| CDH Proband Family | Mother | Proband Blood | Proband Skin | Proband Diaphragm |
|---|---|---|---|---|
| 809 | 0/68 | 0/53 | 0/52 | 21/69 |
| 809 | 0/60 | 0/50 | 0/53 | 20/71 |
| 809 | 0/56 | 0/55 | 0/53 | 18/67 |
| 809 | 0/67 | 0/56 | 0/75 | 20/88 |
| 809 | 0/73 | 46/74 | 38/78 | 41/89 |
| 809 | 0/66 | 14/70 | 27/53 | 12/58 |
| 809 | 0/63 | 0/66 | 0/53 | 22/65 |
| 809 | 0/57 | 0/61 | 0/56 | 16/73 |
| 809 | 0/50 | 48/75 | 29/59 | 42/91 |
| 809 | 0/57 | 39/82 | 31/49 | 35/73 |
| 809 | 0/65 | 0/75 | 0/62 | 18/76 |
| 809 | 0/49 | 0/56 | 0/57 | 28/88 |
| 809 | 0/55 | 0/58 | 0/56 | 23/72 |
| 809 | 0/68 | 0/60 | 0/50 | 21/70 |
| 809 | 0/62 | 14/29 | 13/29 | 18/35 |
| 809 | 0/56 | 0/66 | 0/59 | 23/63 |
| 809 | 0/55 | 27/65 | 37/80 | 43/88 |
| 809 | 0/58 | 0/51 | 0/53 | 19/66 |
| 809 | 0/59 | 34/60 | 21/73 | 21/58 |
| 809 | 0/82 | 0/71 | 0/75 | 24/68 |
| 809 | 0/81 | 0/63 | 0/66 | 15/71 |
| 809 | 0/65 | 0/52 | 0/49 | 23/69 |
| 809 | 0/63 | 0/51 | 0/55 | 22/75 |
| 809 | 0/64 | 35/78 | 34/67 | 42/70 |
| 809 | 0/44 | 38/70 | 35/71 | 27/55 |
| 809 | 0/73 | 0/55 | 0/55 | 15/62 |
| 809 | 0/57 | 0/50 | 0/52 | 17/84 |
| 809 | 0/64 | 0/72 | 0/59 | 17/81 |
| 809 | 0/47 | 0/48 | 0/46 | 23/77 |
| 809 | 0/67 | 30/55 | 35/67 | 23/69 |

**TABLE S3**

| CDH Proband Family | Sex | Location | Chromosome | Position | Reference allele | Alternate allele | Tissue found in | Father |
|---|---|---|---|---|---|---|---|---|
| 809 | Female | Intergenic | 11 | 123118311 | G | C | Diaphragm | 0/40 |
| 809 | Female | Intergenic | 12 | 19007497 | G | A | All three | 0/40 |
| 809 | Female | Intergenic | 12 | 110872268 | A | C | All three | 0/55 |
| 809 | Female | Intergenic | 12 | 113182276 | C | T | Diaphragm | 0/45 |
| 809 | Female | Intergenic | 12 | 113896611 | C | A | Diaphragm | 0/47 |
| 809 | Female | Intergenic | 13 | 26669122 | G | T | All three | 0/45 |
| 809 | female | Intergenic | 13 | 35502281 | A | T | Diaphragm | 0/45 |
| 809 | Female | Intergenic | 13 | 69271119 | T | C | All three | 0/43 |
| 809 | Female | Intergenic | 14 | 34632380 | C | T | All three | 0/48 |
| 809 | Female | Intergenic | 15 | 47132793 | C | T | All three | 0/44 |
| 809 | Female | Intergenic | 16 | 1290232 | C | T | All three | 0/24 |
| 809 | Female | Intergenic | 16 | 5365225 | C | T | Diaphragm | 0/54 |
| 809 | Female | Intergenic | 16 | 49287467 | C | A | Diaphragm | 0/45 |
| 809 | Female | Intergenic | 17 | 14274664 | T | A | All three | 0/41 |
| 809 | Female | Intergenic | 17 | 21723862 | T | A | Diaphragm | 0/47 |
| 809 | Female | Intergenic | 18 | 37760582 | G | A | Diaphragm | 0/43 |
| 809 | Female | Intergenic | 18 | 61185856 | A | T | Diaphragm | 0/45 |
| 809 | Female | Intergenic | 18 | 68778863 | C | T | Diaphragm | 0/48 |
| 809 | Female | Intergenic | 19 | 28034043 | A | T | Diaphragm | 0/55 |
| 809 | Female | Intergenic | 19 | 33836836 | G | A | Diaphragm | 0/43 |
| 809 | Female | Intergenic | 21 | 18844527 | G | C | All three | 0/56 |
| 809 | Female | Intergenic | 21 | 20700243 | C | T | All three | 0/44 |
| 809 | Female | Intergenic | 21 | 28403305 | T | A | All three | 0/60 |
| 809 | Female | Intergenic | X | 455196 | C | T | All three | 0/49 |
| 809 | Female | Intergenic | X | 18180676 | G | C | Diaphragm | 0/27 |
| 809 | Female | Intergenic | X | 35733096 | C | T | Diaphragm | 0/34 |
| 809 | female | Intergenic | X | 66349522 | C | A | Diaphragm | 0/34 |
| 809 | Female | Intergenic | X | 69433109 | C | A | Diaphragm | 0/28 |
| 809 | Female | Intergenic | X | 80911975 | G | A | Diaphragm | 0/30 |
| 809 | Female | Intergenic | X | 84394392 | A | G | Diaphragm | 0/26 |

**TABLE S3**

| CDH Proband Family | Mother | Proband Blood | Proband Skin | Proband Diaphragm |
|---|---|---|---|---|
| 809 | 0/56 | 0/50 | 0/56 | 12/58 |
| 809 | 0/64 | 40/73 | 32/76 | 36/66 |
| 809 | 0/58 | 25/50 | 32/67 | 43/81 |
| 809 | 0/61 | 0/55 | 0/59 | 24/87 |
| 809 | 0/66 | 0/52 | 0/58 | 23/92 |
| 809 | 0/65 | 30/70 | 34/51 | 38/71 |
| 809 | 0/60 | 0/58 | 0/62 | 26/98 |
| 809 | 0/49 | 39/74 | 33/74 | 34/71 |
| 809 | 0/55 | 36/63 | 28/57 | 26/58 |
| 809 | 0/55 | 32/63 | 32/66 | 43/91 |
| 809 | 0/46 | 34/52 | 32/50 | 31/56 |
| 809 | 0/77 | 0/64 | 0/50 | 20/66 |
| 809 | 0/62 | 0/53 | 0/46 | 27/82 |
| 809 | 0/62 | 43/70 | 34/73 | 38/70 |
| 809 | 0/54 | 0/44 | 0/44 | 16/72 |
| 809 | 0/54 | 0/54 | 0/40 | 23/85 |
| 809 | 0/72 | 0/56 | 0/50 | 12/57 |
| 809 | 0/59 | 0/57 | 0/55 | 27/65 |
| 809 | 0/73 | 0/65 | 0/70 | 25/78 |
| 809 | 0/73 | 0/53 | 0/59 | 20/88 |
| 809 | 0/57 | 26/69 | 35/68 | 38/83 |
| 809 | 0/66 | 40/73 | 39/66 | 36/76 |
| 809 | 0/65 | 33/65 | 29/60 | 30/59 |
| 809 | 0/65 | 36/69 | 26/59 | 27/75 |
| 809 | 0/60 | 0/51 | 0/49 | 16/60 |
| 809 | 0/53 | 0/53 | 0/50 | 26/75 |
| 809 | 0/73 | 0/70 | 0/71 | 21/82 |
| 809 | 0/48 | 0/58 | 0/48 | 23/69 |
| 809 | 0/67 | 0/60 | 0/62 | 25/74 |
| 809 | 0/61 | 0/58 | 0/78 | 23/79 |

**TABLE S3**

| CDH Proband Family | Sex | Location | Chromosome | Position | Reference allele | Alternate allele | Tissue found in | Father |
|---|---|---|---|---|---|---|---|---|
| 809 | Female | Intergenic | X | 88218323 | C | A | Diaphragm | 0/21 |
| 809 | Female | Intergenic | X | 89570314 | G | A | Diaphragm | 0/34 |
| 809 | Female | Intergenic | X | 94175321 | C | A | Diaphragm | 0/20 |
| 809 | Female | Intergenic | X | 125153870 | T | G | Diaphragm | 0/34 |
| 809 | Female | Intergenic | X | 145921844 | G | T | Diaphragm | 0/34 |
| 967 | Male | Intergenic | 2 | 58718375 | T | C | All three | 0/50 |
| 967 | Male | Intergenic | 2 | 121280003 | G | C | All three | 0/41 |
| 967 | Male | Intergenic | 2 | 158071768 | G | A | All three | 0/42 |
| 967 | Male | Intergenic | 2 | 205132597 | C | A | All three | 0/54 |
| 967 | Male | Intergenic | 3 | 42294565 | G | A | All three | 0/45 |
| 967 | Male | Intergenic | 3 | 75864306 | C | T | All three | 0/53 |
| 967 | Male | Intergenic | 3 | 80519543 | T | G | All three | 0/51 |
| 967 | Male | Intergenic | 4 | 3793114 | T | C | All three | 0/46 |
| 967 | Male | Intergenic | 4 | 45403029 | A | G | All three | 0/43 |
| 967 | Male | Intergenic | 6 | 148384255 | T | C | All three | 0/49 |
| 967 | Male | Intergenic | 7 | 42518541 | G | A | All three | 0/59 |
| 967 | Male | Intergenic | 8 | 18093640 | C | T | Diaphragm | 0/52 |
| 967 | Male | Intergenic | 8 | 29283236 | C | T | All three | 0/49 |
| 967 | Male | Intergenic | 8 | 35788470 | T | C | All three | 0/46 |
| 967 | Male | Intergenic | 8 | 109375130 | G | A | All three | 0/51 |
| 967 | Male | Intergenic | 9 | 87643765 | A | G | All three | 0/52 |
| 967 | Male | Intergenic | 10 | 9433109 | CCTT | C | All three | 0/57 |
| 967 | Male | Intergenic | 10 | 20971557 | T | G | All three | 0/55 |
| 967 | Male | Intergenic | 10 | 35283939 | A | G | All three | 0/52 |
| 967 | Male | Intergenic | 10 | 114676907 | A | AT | All three | 0/50 |
| 967 | Male | Intergenic | 11 | 35124820 | C | T | All three | 0/58 |
| 967 | Male | Intergenic | 11 | 80173545 | G | A | All three | 0/51 |
| 967 | male | Intergenic | 14 | 24359361 | G | A | Diaphragm | 0/58 |
| 967 | Male | Intergenic | 14 | 101628801 | GA | G | All three | 0/49 |
| 967 | male | Intergenic | 15 | 25739074 | C | T | Diaphragm+Skin | 0/64 |

**TABLE S3**

De Novo Intergenic Diaphragm

| CDH Proband Family | Mother | Proband Blood | Proband Skin | Proband Diaphragm |
|---|---|---|---|---|
| 809 | 0/58 | 0/52 | 0/52 | 19/59 |
| 809 | 0/67 | 0/46 | 0/50 | 13/65 |
| 809 | 0/74 | 0/64 | 0/54 | 24/72 |
| 809 | 0/56 | 0/68 | 0/59 | 14/70 |
| 809 | 0/66 | 0/50 | 0/62 | 28/78 |
| 967 | 0/48 | 32/57 | 40/79 | 24/52 |
| 967 | 0/50 | 19/41 | 31/64 | 19/35 |
| 967 | 0/66 | 19/47 | 34/67 | 33/58 |
| 967 | 0/58 | 27/52 | 29/64 | 29/45 |
| 967 | 0/51 | 34/53 | 28/60 | 25/35 |
| 967 | 0/53 | 33/64 | 37/75 | 19/43 |
| 967 | 0/56 | 31/56 | 40/83 | 38/61 |
| 967 | 0/51 | 30/51 | 40/79 | 16/34 |
| 967 | 0/36 | 30/54 | 18/34 | 15/38 |
| 967 | 0/56 | 26/47 | 38/74 | 25/48 |
| 967 | 0/54 | 34/62 | 35/75 | 27/52 |
| 967 | 0/61 | 0/56 | 0/46 | 7/34 |
| 967 | 0/38 | 28/60 | 34/60 | 15/36 |
| 967 | 0/43 | 30/59 | 34/71 | 30/61 |
| 967 | 0/52 | 34/84 | 31/54 | 33/58 |
| 967 | 0/46 | 25/54 | 20/49 | 34/46 |
| 967 | 0/63 | 31/62 | 36/67 | 14/36 |
| 967 | 0/47 | 31/54 | 39/64 | 18/42 |
| 967 | 0/53 | 17/38 | 35/68 | 13/32 |
| 967 | 0/61 | 13/29 | 20/35 | 11/20 |
| 967 | 0/62 | 30/60 | 28/51 | 22/44 |
| 967 | 0/59 | 22/50 | 32/68 | 25/50 |
| 967 | 0/79 | 10/58 | 12/72 | 9/43 |
| 967 | 0/58 | 37/63 | 32/69 | 31/52 |
| 967 | 0/83 | 6/60 | 16/82 | 15/58 |

**TABLE S3**

| CDH Proband Family | Sex | Location | Chromosome | Position | Reference allele | Alternate allele | Tissue found in | Father |
|---|---|---|---|---|---|---|---|---|
| 967 | Male | Intergenic | 15 | 73336781 | C | A | All three | 0/49 |
| 967 | Male | Intergenic | 17 | 39613324 | T | C | All three | 0/41 |
| 967 | Male | Intergenic | 17 | 51582010 | A | G | All three | 0/41 |
| 967 | Male | Intergenic | 22 | 25083212 | TACTC | T | All three | 0/47 |
| 967 | Male | Intergenic | X | 35867136 | G | A | All three | 0/23 |
| 967 | male | Intergenic | X | 68010485 | C | T | Diaphragm+Skin | 0/29 |
| 967 | Male | Intergenic | X | 139661068 | A | G | Diaphragm | 0/23 |
| 967 | Male | Intergenic | Y | 13317016 | G | A | All three | 0/36 |
| 967 | Male | Intergenic | Y | 17597506 | G | GA | All three | 0/32 |

**TABLE S3**

| CDH Proband Family | Mother | Proband Blood | Proband Skin | Proband Diaphragm |
|---|---|---|---|---|
| 967 | 0/48 | 24/59 | 34/61 | 30/57 |
| 967 | 0/64 | 32/58 | 40/67 | 23/52 |
| 967 | 0/47 | 31/53 | 37/67 | 22/38 |
| 967 | 0/64 | 23/60 | 32/56 | 24/48 |
| 967 | 0/47 | 29/29 | 37/37 | 13/13 |
| 967 | 0/60 | 2/30 | 18/40 | 5/22 |
| 967 | 0/52 | 0/33 | 0/35 | 5/18 |
| 967 | 0/28 | 23/36 | 33/48 | 16/38 |
| 967 | 0/0 | 24/25 | 27/29 | 15/15 |

**TABLE S3**

| CDH Proband Family | Sex | Location | Chromosome | Position | Reference allele | Alternate allele | Tissue found in | Father |
|---|---|---|---|---|---|---|---|---|
| 411 | Male | Intergenic | 2 | 6448411 | G | A | All three | 0/51 |
| 411 | Male | Intergenic | 2 | 15979430 | C | T | All three | 0/46 |
| 411 | Male | Intergenic | 2 | 23607772 | C | A | All three | 0/34 |
| 411 | Male | Intergenic | 2 | 23607823 | G | A | All three | 0/34 |
| 411 | Male | Intergenic | 2 | 48488807 | [illegible]CTTTGAAAG[illegible] | T | All three | 0/55 |
| 411 | male | Intergenic | 4 | 12542452 | ACAAAAAT | A | All three | 0/61 |
| 411 | Male | Intergenic | 4 | 40268400 | A | G | All three | 0/59 |
| 411 | Male | Intergenic | 4 | 130485536 | C | T | All three | 0/39 |
| 411 | Male | Intergenic | 4 | 135941340 | ATAATACATG | T | Skin | 0/36 |
| 411 | Male | Intergenic | 5 | 30972421 | G | T | All three | 0/54 |
| 411 | Male | Intergenic | 5 | 43876751 | A | T | All three | 0/43 |
| 411 | Male | Intergenic | 7 | 121415957 | T | G | All three | 0/42 |
| 411 | Male | Intergenic | 8 | 49656228 | G | C | All three | 0/45 |
| 411 | Male | Intergenic | 9 | 24573667 | A | C | All three | 0/57 |
| 411 | Male | Intergenic | 10 | 45696971 | C | T | All three | 0/44 |
| 411 | male | Intergenic | 10 | 107240148 | T | C | All three | 0/54 |
| 411 | Male | Intergenic | 11 | 48522238 | T | C | All three | 0/53 |
| 411 | Male | Intergenic | 12 | 51286910 | T | C | All three | 0/46 |
| 411 | Male | Intergenic | 13 | 63562946 | G | GA | All three | 0/63 |
| 411 | Male | Intergenic | 15 | 38941044 | C | T | All three | 0/45 |
| 411 | Male | Intergenic | 15 | 46407038 | C | T | All three | 0/55 |
| 411 | Male | Intergenic | 16 | 75543055 | C | A | All three | 0/46 |
| 411 | Male | Intergenic | 17 | 404337 | C | T | All three | 0/55 |
| 411 | Male | Intergenic | 20 | 5206920 | C | A | All three | 0/45 |
| 411 | Male | Intergenic | 20 | 48832820 | CAG | C | Skin | 0/45 |
| 411 | Male | Intergenic | X | 3355438 | CTG | C | Skin | 0/38 |
| 411 | Male | Intergenic | Y | 21994006 | CAG | C | Skin | 0/29 |
| 411 | Male | Intergenic | Y | 23424401 | T | C | All three | 0/37 |
| 716 | Female | Intergenic | 1 | 25520943 | G | A | All three | 0/53 |
| 716 | Female | Intergenic | 1 | 108550027 | G | A | All three | 0/55 |

De Novo Intergenic Skin

| CDH Proband Family | Mother | Proband Blood | Proband Skin | Proband Diaphragm |
|---|---|---|---|---|
| 411 | 0/49 | 29/59 | 21/54 | 51/89 |
| 411 | 0/38 | 16/47 | 23/50 | 36/63 |
| 411 | 0/39 | 16/26 | 25/40 | 23/56 |
| 411 | 0/47 | 16/21 | 17/25 | 17/49 |
| 411 | 0/46 | 23/50 | 22/60 | 32/70 |
| 411 | 0/67 | 23/54 | 23/50 | 32/86 |
| 411 | 0/56 | 36/75 | 30/55 | 38/78 |
| 411 | 0/55 | 34/67 | 23/58 | 36/75 |
| 411 | 0/46 | 0/49 | 19/86 | 0/40 |
| 411 | 0/52 | 28/65 | 34/65 | 41/99 |
| 411 | 0/43 | 37/78 | 32/59 | 21/55 |
| 411 | 0/50 | 25/62 | 26/61 | 42/81 |
| 411 | 0/47 | 31/57 | 21/58 | 42/75 |
| 411 | 0/45 | 24/69 | 18/48 | 32/78 |
| 411 | 0/59 | 33/70 | 31/65 | 29/60 |
| 411 | 0/58 | 32/79 | 25/58 | 33/71 |
| 411 | 0/41 | 26/61 | 24/69 | 35/85 |
| 411 | 0/47 | 57/97 | 34/58 | 35/80 |
| 411 | 0/51 | 19/48 | 20/40 | 33/68 |
| 411 | 0/48 | 28/58 | 31/72 | 37/75 |
| 411 | 0/43 | 34/58 | 22/57 | 36/77 |
| 411 | 0/43 | 22/56 | 30/53 | 41/79 |
| 411 | 0/50 | 36/73 | 28/49 | 38/77 |
| 411 | 0/43 | 26/57 | 26/67 | 35/77 |
| 411 | 0/44 | 0/45 | 6/27 | 0/62 |
| 411 | 0/56 | 0/30 | 4/18 | 0/44 |
| 411 | 0/0 | 4/44 | 6/24 | 0/36 |
| 411 | 0/0 | 35/35 | 35/36 | 31/31 |
| 716 | 0/58 | 24/50 | 40/84 | 23/51 |
| 716 | 0/58 | 38/66 | 36/64 | 33/66 |

**TABLE S3**

| CDH Proband Family | Sex | Location | Chromosome | Position | Reference allele | Alternate allele | Tissue found in | Father |
|---|---|---|---|---|---|---|---|---|
| 716 | Female | Intergenic | 1 | 188119862 | C | T | All three | 0/60 |
| 716 | female | Intergenic | 1 | 190952170 | A | G | Diaphragm+Skin | 0/20 |
| 716 | Female | Intergenic | 1 | 219133910 | G | A | All three | 0/54 |
| 716 | Female | Intergenic | 1 | 241538742 | A | G | All three | 0/64 |
| 716 | Female | Intergenic | 2 | 60131252 | T | C | All three | 0/56 |
| 716 | Female | Intergenic | 2 | 237733339 | C | T | All three | 0/48 |
| 716 | Female | Intergenic | 2 | 241273592 | G | T | All three | 0/48 |
| 716 | Female | Intergenic | 3 | 31317266 | C | T | All three | 0/65 |
| 716 | Female | Intergenic | 3 | 70707930 | T | C | All three | 0/51 |
| 716 | Female | Intergenic | 3 | 87755171 | C | T | All three | 0/51 |
| 716 | Female | Intergenic | 3 | 96134789 | G | A | All three | 0/58 |
| 716 | Female | Intergenic | 3 | 116969700 | C | T | All three | 0/61 |
| 716 | Female | Intergenic | 4 | 76738805 | A | C | All three | 0/55 |
| 716 | Female | Intergenic | 4 | 90360867 | C | G | All three | 0/45 |
| 716 | Female | Intergenic | 4 | 92765694 | T | A | All three | 0/55 |
| 716 | Female | Intergenic | 4 | 165797047 | C | T | All three | 0/49 |
| 716 | Female | Intergenic | 5 | 13079144 | A | T | All three | 0/45 |
| 716 | Female | Intergenic | 5 | 57641900 | G | T | Skin | 0/48 |
| 716 | Female | Intergenic | 5 | 92381616 | C | G | All three | 0/62 |
| 716 | Female | Intergenic | 6 | 18809404 | A | C | All three | 0/66 |
| 716 | Female | Intergenic | 6 | 50605045 | A | G | All three | 0/57 |
| 716 | Female | Intergenic | 6 | 69243495 | C | G | All three | 0/41 |
| 716 | Female | Intergenic | 6 | 78773081 | C | T | All three | 0/62 |
| 716 | Female | Intergenic | 7 | 2657275 | C | T | All three | 0/47 |
| 716 | Female | Intergenic | 7 | 6718899 | A | G | All three | 0/68 |
| 716 | Female | Intergenic | 7 | 19285491 | C | T | All three | 0/50 |
| 716 | Female | Intergenic | 7 | 96000892 | G | T | All three | 0/46 |
| 716 | Female | Intergenic | 8 | 23549713 | A | G | All three | 0/50 |
| 716 | Female | Intergenic | 8 | 50342185 | C | A | All three | 0/55 |
| 716 | Female | Intergenic | 8 | 53690025 | G | A | All three | 0/65 |

**TABLE S3**

| CDH Proband Family | Mother | Proband Blood | Proband Skin | Proband Diaphragm |
|---|---|---|---|---|
| 716 | 0/65 | 27/61 | 38/66 | 23/56 |
| 716 | 0/44 | 3/25 | 7/27 | 4/16 |
| 716 | 0/50 | 37/69 | 28/56 | 29/66 |
| 716 | 0/58 | 40/85 | 46/78 | 33/67 |
| 716 | 0/57 | 28/55 | 43/90 | 49/79 |
| 716 | 0/51 | 29/68 | 32/76 | 39/74 |
| 716 | 0/54 | 42/89 | 30/77 | 26/55 |
| 716 | 0/56 | 34/61 | 26/59 | 39/77 |
| 716 | 0/50 | 26/62 | 32/61 | 35/66 |
| 716 | 0/59 | 35/82 | 34/67 | 38/66 |
| 716 | 0/60 | 31/58 | 46/81 | 23/58 |
| 716 | 0/72 | 35/79 | 37/71 | 23/52 |
| 716 | 0/56 | 26/54 | 21/45 | 45/67 |
| 716 | 0/61 | 29/59 | 25/53 | 29/60 |
| 716 | 0/46 | 35/69 | 40/63 | 31/63 |
| 716 | 0/57 | 39/68 | 34/69 | 39/72 |
| 716 | 0/59 | 27/60 | 36/63 | 28/51 |
| 716 | 0/53 | 0/56 | 14/68 | 0/48 |
| 716 | 0/59 | 40/72 | 27/57 | 32/57 |
| 716 | 0/68 | 30/68 | 33/77 | 22/47 |
| 716 | 0/53 | 39/75 | 28/58 | 36/56 |
| 716 | 0/61 | 26/53 | 31/72 | 35/69 |
| 716 | 0/58 | 38/79 | 36/62 | 38/76 |
| 716 | 0/57 | 30/58 | 23/54 | 29/67 |
| 716 | 0/55 | 34/75 | 35/78 | 34/68 |
| 716 | 0/63 | 36/70 | 38/73 | 41/77 |
| 716 | 0/52 | 23/83 | 19/64 | 29/71 |
| 716 | 0/47 | 17/55 | 34/68 | 37/76 |
| 716 | 0/55 | 21/47 | 42/77 | 48/85 |
| 716 | 0/54 | 25/44 | 26/66 | 30/58 |

De Novo Intergenic Skin

| CDH Proband Family | Sex | Location | Chromosome | Position | Reference allele | Alternate allele | Tissue found in | Father |
|---|---|---|---|---|---|---|---|---|
| 716 | Female | Intergenic | 8 | 121394186 | C | G | All three | 0/43 |
| 716 | Female | Intergenic | 9 | 37870079 | C | A | All three | 0/52 |
| 716 | Female | Intergenic | 9 | 92940311 | C | T | All three | 0/62 |
| 716 | Female | Intergenic | 10 | 21531548 | A | T | All three | 0/61 |
| 716 | female | Intergenic | 10 | 78610342 | GACAA | G | All three | 0/79 |
| 716 | Female | Intergenic | 10 | 102805027 | C | T | All three | 0/49 |
| 716 | Female | Intergenic | 10 | 111937754 | A | G | All three | 0/55 |
| 716 | Female | Intergenic | 11 | 68910808 | C | T | All three | 0/52 |
| 716 | Female | Intergenic | 11 | 68917454 | A | T | All three | 0/63 |
| 716 | Female | Intergenic | 11 | 85940710 | C | T | All three | 0/52 |
| 716 | Female | Intergenic | 12 | 2894972 | G | A | All three | 0/71 |
| 716 | Female | Intergenic | 12 | 20253658 | G | A | All three | 0/60 |
| 716 | Female | Intergenic | 12 | 61786788 | C | T | All three | 0/56 |
| 716 | female | Intergenic | 12 | 73894281 | T | C | All three | 0/67 |
| 716 | Female | Intergenic | 12 | 81133670 | T | C | All three | 0/44 |
| 716 | Female | Intergenic | 12 | 128640569 | T | C | All three | 0/60 |
| 716 | Female | Intergenic | 13 | 38770384 | A | T | All three | 0/56 |
| 716 | Female | Intergenic | 13 | 53424768 | G | T | Skin | 0/49 |
| 716 | Female | Intergenic | 14 | 94631627 | C | T | All three | 0/47 |
| 716 | Female | Intergenic | 14 | 98723131 | C | T | All three | 0/57 |
| 716 | Female | Intergenic | 15 | 53418167 | A | G | All three | 0/59 |
| 716 | Female | Intergenic | 18 | 26978994 | C | T | All three | 0/53 |
| 716 | Female | Intergenic | 18 | 48313755 | G | T | All three | 0/60 |
| 716 | Female | Intergenic | 18 | 63851839 | G | C | All three | 0/55 |
| 716 | Female | Intergenic | 19 | 583802 | C | T | Skin | 0/53 |
| 716 | Female | Intergenic | 19 | 12407418 | A | C | All three | 0/78 |
| 716 | Female | Intergenic | 20 | 40498713 | A | T | All three | 0/41 |
| 716 | Female | Intergenic | 21 | 26441156 | G | T | All three | 0/53 |
| 716 | Female | Intergenic | 21 | 29172156 | A | T | All three | 0/59 |
| 716 | Female | Intergenic | X | 39689788 | C | G | All three | 0/31 |

TABLE S3

De Novo Intergenic Skin

| CDH Proband Family | Mother | Proband Blood | Proband Skin | Proband Diaphragm |
|---|---|---|---|---|
| 716 | 0/43 | 20/50 | 38/72 | 31/54 |
| 716 | 0/52 | 29/68 | 31/71 | 35/66 |
| 716 | 0/56 | 28/66 | 37/73 | 38/77 |
| 716 | 0/56 | 33/61 | 37/73 | 30/61 |
| 716 | 0/80 | 38/79 | 35/88 | 45/91 |
| 716 | 0/49 | 40/77 | 45/80 | 36/73 |
| 716 | 0/47 | 38/77 | 43/81 | 43/68 |
| 716 | 0/59 | 34/72 | 36/71 | 27/59 |
| 716 | 0/52 | 40/72 | 35/69 | 36/65 |
| 716 | 0/53 | 28/63 | 38/76 | 31/57 |
| 716 | 0/58 | 47/80 | 31/61 | 38/76 |
| 716 | 0/64 | 39/77 | 29/50 | 43/61 |
| 716 | 0/74 | 36/76 | 34/67 | 36/61 |
| 716 | 0/77 | 43/81 | 40/73 | 33/71 |
| 716 | 0/63 | 37/65 | 38/72 | 30/70 |
| 716 | 0/64 | 22/57 | 38/73 | 35/76 |
| 716 | 0/42 | 28/57 | 30/63 | 32/61 |
| 716 | 0/48 | 0/44 | 11/53 | 0/45 |
| 716 | 0/61 | 30/55 | 31/66 | 23/44 |
| 716 | 0/55 | 34/65 | 32/66 | 30/67 |
| 716 | 0/60 | 24/67 | 30/66 | 35/66 |
| 716 | 0/49 | 28/62 | 34/74 | 38/74 |
| 716 | 0/53 | 35/64 | 33/71 | 40/74 |
| 716 | 0/54 | 38/76 | 41/83 | 33/67 |
| 716 | 0/42 | 0/43 | 14/60 | 0/50 |
| 716 | 0/74 | 46/68 | 28/54 | 44/62 |
| 716 | 0/46 | 28/68 | 37/68 | 25/59 |
| 716 | 0/58 | 27/62 | 41/73 | 28/59 |
| 716 | 0/62 | 44/74 | 34/84 | 30/58 |
| 716 | 0/57 | 31/66 | 24/54 | 34/81 |

De Novo Intergenic Skin

| CDH Proband Family | Sex | Location | Chromosome | Position | Reference allele | Alternate allele | Tissue found in | Father |
|---|---|---|---|---|---|---|---|---|
| 716 | Female | Intergenic | X | 65140289 | G | A | All three | 0/25 |
| 716 | Female | Intergenic | X | 82221769 | C | T | All three | 0/30 |
| 716 | Female | Intergenic | X | 88110135 | C | T | All three | 0/34 |
| 716 | Female | Intergenic | X | 89554886 | C | T | All three | 0/25 |
| 716 | Female | Intergenic | X | 105821982 | G | T | All three | 0/35 |
| 809 | Female | Intergenic | 1 | 229943691 | T | A | All three | 0/41 |
| 809 | Female | Intergenic | 1 | 248631521 | G | A | All three | 0/47 |
| 809 | Female | Intergenic | 2 | 52678867 | T | G | Skin | 0/39 |
| 809 | Female | Intergenic | 2 | 81121345 | A | G | Skin | 0/47 |
| 809 | Female | Intergenic | 2 | 111473446 | G | A | Skin | 0/43 |
| 809 | Female | Intergenic | 2 | 115175328 | G | A | Skin | 0/43 |
| 809 | Female | Intergenic | 2 | 118456374 | T | C | All three | 0/46 |
| 809 | Female | Intergenic | 2 | 125969667 | T | A | Skin | 0/48 |
| 809 | female | Intergenic | 2 | 165063447 | G | A | Skin+Blood | 0/64 |
| 809 | Female | Intergenic | 2 | 214077949 | ATTAC | A | Skin | 0/41 |
| 809 | Female | Intergenic | 2 | 235795397 | C | T | Skin | 0/40 |
| 809 | Female | Intergenic | 3 | 5446160 | G | A | Skin | 0/39 |
| 809 | female | Intergenic | 3 | 35466073 | C | T | Skin+Blood | 0/53 |
| 809 | Female | Intergenic | 3 | 74259109 | A | G | All three | 0/39 |
| 809 | Female | Intergenic | 3 | 99014368 | G | C | Skin | 0/48 |
| 809 | Female | Intergenic | 3 | 117747604 | G | T | Skin | 0/46 |
| 809 | Female | Intergenic | 3 | 137898799 | C | CAT | Skin | 0/44 |
| 809 | female | Intergenic | 4 | 4112769 | C | T | Skin | 0/52 |
| 809 | Female | Intergenic | 4 | 8715501 | G | A | All three | 0/44 |
| 809 | female | Intergenic | 4 | 12826901 | A | T | Skin | 0/54 |
| 809 | Female | Intergenic | 4 | 30258855 | G | T | Skin | 0/44 |
| 809 | Female | Intergenic | 4 | 36849853 | G | A | Skin | 0/62 |
| 809 | Female | Intergenic | 4 | 59397258 | C | T | All three | 0/48 |
| 809 | Female | Intergenic | 4 | 75368244 | G | T | Skin | 0/45 |
| 809 | Female | Intergenic | 4 | 187684926 | T | C | Skin | 0/41 |

De Novo Intergenic Skin

| CDH Proband Family | Mother | Proband Blood | Proband Skin | Proband Diaphragm |
|---|---|---|---|---|
| 716 | 0/47 | 40/79 | 32/68 | 31/70 |
| 716 | 0/55 | 39/68 | 37/69 | 34/54 |
| 716 | 0/63 | 35/64 | 36/69 | 35/68 |
| 716 | 0/44 | 34/72 | 25/57 | 35/82 |
| 716 | 0/48 | 35/68 | 39/66 | 33/61 |
| 809 | 0/51 | 27/73 | 43/70 | 29/61 |
| 809 | 0/52 | 26/60 | 35/61 | 40/78 |
| 809 | 0/66 | 0/63 | 18/59 | 0/64 |
| 809 | 0/61 | 0/60 | 18/61 | 0/49 |
| 809 | 0/53 | 0/47 | 16/73 | 0/53 |
| 809 | 0/67 | 0/49 | 17/57 | 0/62 |
| 809 | 0/56 | 26/65 | 35/71 | 39/71 |
| 809 | 0/47 | 0/61 | 18/65 | 0/61 |
| 809 | 0/78 | 19/83 | 26/93 | 12/82 |
| 809 | 0/63 | 0/54 | 17/58 | 0/54 |
| 809 | 0/72 | 0/58 | 23/69 | 0/67 |
| 809 | 0/62 | 0/61 | 22/67 | 0/62 |
| 809 | 0/66 | 19/89 | 19/68 | 11/80 |
| 809 | 0/59 | 34/70 | 33/59 | 30/64 |
| 809 | 0/73 | 0/54 | 25/76 | 0/70 |
| 809 | 0/59 | 0/60 | 19/61 | 0/63 |
| 809 | 0/61 | 0/61 | 15/50 | 0/56 |
| 809 | 1/74 | 0/79 | 24/67 | 0/91 |
| 809 | 0/58 | 42/69 | 40/76 | 33/73 |
| 809 | 1/54 | 0/60 | 20/51 | 0/66 |
| 809 | 0/61 | 0/49 | 22/62 | 0/45 |
| 809 | 0/66 | 0/75 | 19/54 | 0/66 |
| 809 | 0/77 | 41/70 | 43/67 | 31/58 |
| 809 | 0/66 | 0/81 | 29/70 | 0/67 |
| 809 | 0/59 | 0/51 | 24/79 | 0/56 |

**TABLE S3**

| CDH Proband Family | Sex | Location | Chromosome | Position | Reference allele | Alternate allele | Tissue found in | Father |
|---|---|---|---|---|---|---|---|---|
| 809 | female | Intergenic | 5 | 132509196 | T | C | All three | 0/59 |
| 809 | Female | Intergenic | 5 | 144084315 | G | GA | All three | 0/40 |
| 809 | Female | Intergenic | 5 | 144352082 | A | T | Skin | 0/47 |
| 809 | Female | Intergenic | 5 | 175131810 | G | T | Skin | 0/43 |
| 809 | Female | Intergenic | 6 | 9617082 | G | T | Skin | 0/38 |
| 809 | Female | Intergenic | 6 | 57726168 | G | A | Skin | 0/45 |
| 809 | Female | Intergenic | 6 | 84499729 | T | C | All three | 0/43 |
| 809 | Female | Intergenic | 6 | 118084618 | A | T | Skin | 0/55 |
| 809 | Female | Intergenic | 6 | 142319520 | A | G | Skin | 0/40 |
| 809 | Female | Intergenic | 7 | 46969823 | T | A | All three | 0/39 |
| 809 | Female | Intergenic | 7 | 134456152 | T | A | Skin | 0/44 |
| 809 | Female | Intergenic | 8 | 24834454 | C | T | All three | 0/50 |
| 809 | Female | Intergenic | 8 | 29547994 | C | A | Skin | 0/41 |
| 809 | Female | Intergenic | 8 | 51862888 | C | T | Skin | 0/41 |
| 809 | Female | Intergenic | 8 | 60040127 | C | T | All three | 0/47 |
| 809 | Female | Intergenic | 8 | 112091216 | T | A | Skin | 0/48 |
| 809 | Female | Intergenic | 8 | 135335680 | C | T | Skin | 0/48 |
| 809 | female | Intergenic | 8 | 138869304 | T | G | Skin+Blood | 1/62 |
| 809 | Female | Intergenic | 9 | 23952880 | C | A | Skin | 0/41 |
| 809 | Female | Intergenic | 9 | 30064518 | T | C | Skin | 0/43 |
| 809 | Female | Intergenic | 9 | 90691068 | C | T | Skin | 0/38 |
| 809 | Female | Intergenic | 9 | 99837526 | G | A | Skin | 0/38 |
| 809 | Female | Intergenic | 10 | 4733230 | A | G | All three | 0/44 |
| 809 | Female | Intergenic | 10 | 42969976 | G | T | Skin | 0/44 |
| 809 | female | Intergenic | 10 | 47607914 | G | A | Skin | 0/72 |
| 809 | Female | Intergenic | 10 | 58006434 | C | A | Skin | 0/41 |
| 809 | Female | Intergenic | 10 | 66550830 | T | C | Skin | 0/46 |
| 809 | Female | Intergenic | 11 | 15343128 | AT | A | All three | 0/39 |
| 809 | Female | Intergenic | 11 | 37638703 | G | C | All three | 0/55 |
| 809 | Female | Intergenic | 11 | 80793572 | G | A | Skin | 0/42 |

TABLE S3

De Novo Intergenic Skin

| CDH Proband Family | Mother | Proband Blood | Proband Skin | Proband Diaphragm |
|---|---|---|---|---|
| 809 | 0/73 | 46/74 | 38/78 | 41/89 |
| 809 | 0/66 | 14/70 | 27/53 | 12/58 |
| 809 | 0/51 | 0/63 | 19/76 | 0/52 |
| 809 | 0/65 | 0/52 | 17/83 | 0/56 |
| 809 | 0/59 | 0/56 | 19/61 | 0/55 |
| 809 | 0/50 | 0/66 | 21/68 | 0/66 |
| 809 | 0/50 | 48/75 | 29/59 | 42/91 |
| 809 | 0/63 | 0/63 | 30/80 | 0/60 |
| 809 | 0/62 | 0/57 | 17/65 | 0/68 |
| 809 | 0/57 | 39/82 | 31/49 | 35/73 |
| 809 | 0/51 | 0/65 | 22/75 | 0/61 |
| 809 | 0/62 | 14/29 | 13/29 | 18/35 |
| 809 | 0/52 | 0/53 | 20/75 | 0/57 |
| 809 | 0/50 | 0/59 | 28/70 | 0/61 |
| 809 | 0/55 | 27/65 | 37/80 | 43/88 |
| 809 | 0/73 | 0/57 | 21/72 | 0/57 |
| 809 | 0/56 | 0/49 | 14/70 | 0/57 |
| 809 | 0/68 | 23/77 | 26/74 | 12/74 |
| 809 | 0/56 | 0/62 | 24/62 | 0/54 |
| 809 | 0/67 | 0/53 | 18/71 | 0/43 |
| 809 | 0/58 | 0/53 | 17/58 | 0/49 |
| 809 | 0/59 | 0/48 | 16/52 | 0/56 |
| 809 | 0/59 | 34/60 | 21/73 | 21/58 |
| 809 | 0/43 | 0/62 | 20/68 | 0/56 |
| 809 | 0/81 | 0/89 | 22/66 | 0/70 |
| 809 | 0/58 | 0/48 | 24/90 | 0/47 |
| 809 | 0/61 | 0/50 | 20/79 | 0/48 |
| 809 | 0/64 | 35/78 | 34/67 | 42/70 |
| 809 | 0/44 | 38/70 | 35/71 | 27/55 |
| 809 | 0/57 | 0/72 | 18/61 | 0/53 |

**TABLE S3**

| CDH Proband Family | Sex | Location | Chromosome | Position | Reference allele | Alternate allele | Tissue found in | Father |
|---|---|---|---|---|---|---|---|---|
| 809 | Female | Intergenic | 11 | 112170508 | T | C | All three | 0/45 |
| 809 | Female | Intergenic | 11 | 116677972 | C | T | Skin | 0/48 |
| 809 | Female | Intergenic | 12 | 19007497 | G | A | All three | 0/40 |
| 809 | Female | Intergenic | 12 | 71618878 | G | A | Skin | 0/47 |
| 809 | Female | Intergenic | 12 | 82273075 | G | A | Skin | 0/49 |
| 809 | Female | Intergenic | 12 | 82620453 | G | A | Skin | 0/49 |
| 809 | Female | Intergenic | 12 | 110872268 | A | C | All three | 0/55 |
| 809 | Female | Intergenic | 13 | 25735290 | C | T | Skin | 0/54 |
| 809 | Female | Intergenic | 13 | 26669122 | G | T | All three | 0/45 |
| 809 | Female | Intergenic | 13 | 57081014 | T | G | Skin | 0/52 |
| 809 | Female | Intergenic | 13 | 62968767 | G | A | Skin | 0/45 |
| 809 | Female | Intergenic | 13 | 69271119 | T | C | All three | 0/43 |
| 809 | Female | Intergenic | 13 | 80553307 | T | C | Skin | 0/39 |
| 809 | Female | Intergenic | 14 | 34632380 | C | T | All three | 0/48 |
| 809 | Female | Intergenic | 14 | 47032376 | T | C | Skin | 0/38 |
| 809 | Female | Intergenic | 14 | 67992825 | G | T | Skin | 0/47 |
| 809 | Female | Intergenic | 15 | 47132793 | C | T | All three | 0/44 |
| 809 | Female | Intergenic | 16 | 1290232 | C | T | All three | 0/24 |
| 809 | Female | Intergenic | 16 | 9322325 | G | A | Skin | 0/46 |
| 809 | Female | Intergenic | 16 | 13426591 | C | G | Skin | 0/44 |
| 809 | Female | Intergenic | 16 | 76211179 | C | A | Skin | 0/41 |
| 809 | Female | Intergenic | 17 | 14274664 | T | A | All three | 0/41 |
| 809 | Female | Intergenic | 17 | 52194918 | G | T | Skin | 0/51 |
| 809 | Female | Intergenic | 17 | 52357758 | A | T | Skin | 0/47 |
| 809 | Female | Intergenic | 18 | 12059828 | T | G | Skin | 0/52 |
| 809 | Female | Intergenic | 18 | 25950365 | G | T | Skin | 0/46 |
| 809 | Female | Intergenic | 18 | 49692265 | G | A | Skin | 0/43 |
| 809 | Female | Intergenic | 18 | 70286882 | C | T | Skin | 0/42 |
| 809 | Female | Intergenic | 20 | 38137398 | G | A | Skin | 0/45 |
| 809 | Female | Intergenic | 20 | 44965441 | C | A | Skin | 0/44 |

De Novo Intergenic Skin

| CDH Proband Family | Mother | Proband Blood | Proband Skin | Proband Diaphragm |
|---|---|---|---|---|
| 809 | 0/67 | 30/55 | 35/67 | 23/69 |
| 809 | 0/62 | 0/69 | 20/63 | 0/69 |
| 809 | 0/64 | 40/73 | 32/76 | 36/66 |
| 809 | 0/69 | 0/61 | 18/77 | 0/49 |
| 809 | 0/63 | 0/66 | 23/73 | 0/55 |
| 809 | 0/56 | 0/53 | 20/67 | 0/61 |
| 809 | 0/58 | 25/50 | 32/67 | 43/81 |
| 809 | 0/54 | 0/61 | 19/63 | 0/62 |
| 809 | 0/65 | 30/70 | 34/51 | 38/71 |
| 809 | 0/66 | 0/60 | 26/77 | 0/64 |
| 809 | 0/57 | 0/45 | 18/78 | 0/49 |
| 809 | 0/49 | 39/74 | 33/74 | 34/71 |
| 809 | 0/61 | 0/59 | 17/64 | 0/60 |
| 809 | 0/55 | 36/63 | 28/57 | 26/58 |
| 809 | 0/57 | 0/58 | 22/68 | 0/57 |
| 809 | 0/62 | 0/54 | 21/59 | 0/62 |
| 809 | 0/55 | 32/63 | 32/66 | 43/91 |
| 809 | 0/46 | 34/52 | 32/50 | 31/56 |
| 809 | 0/62 | 0/68 | 21/82 | 0/59 |
| 809 | 0/65 | 0/60 | 16/65 | 0/62 |
| 809 | 0/52 | 0/58 | 15/60 | 0/54 |
| 809 | 0/62 | 43/70 | 34/73 | 38/70 |
| 809 | 0/70 | 0/55 | 21/70 | 0/53 |
| 809 | 0/57 | 0/53 | 22/72 | 0/63 |
| 809 | 0/80 | 0/70 | 22/67 | 0/70 |
| 809 | 0/72 | 0/57 | 23/75 | 0/63 |
| 809 | 0/54 | 0/70 | 20/68 | 0/65 |
| 809 | 0/48 | 0/45 | 28/86 | 0/58 |
| 809 | 0/59 | 0/58 | 20/71 | 0/60 |
| 809 | 0/58 | 0/50 | 24/70 | 0/69 |

**TABLE S3**

| CDH Proband Family | Sex | Location | Chromosome | Position | Reference allele | Alternate allele | Tissue found in | Father |
|---|---|---|---|---|---|---|---|---|
| 809 | female | Intergenic | 20 | 46859236 | C | T | Skin+Blood | 0/58 |
| 809 | Female | Intergenic | 20 | 53041363 | T | C | Skin | 0/36 |
| 809 | Female | Intergenic | 21 | 16549913 | C | A | Skin | 0/41 |
| 809 | Female | Intergenic | 21 | 18844527 | G | C | All three | 0/56 |
| 809 | Female | Intergenic | 21 | 20700243 | C | T | All three | 0/44 |
| 809 | Female | Intergenic | 21 | 28403305 | T | A | All three | 0/60 |
| 809 | Female | Intergenic | X | 455196 | C | T | All three | 0/49 |
| 809 | Female | Intergenic | X | 21286505 | T | C | Skin | 0/26 |
| 809 | Female | Intergenic | X | 89240172 | A | T | Skin | 0/28 |
| 809 | Female | Intergenic | X | 120808757 | C | A | Skin | 0/21 |
| 809 | Female | Intergenic | X | 127078243 | T | G | Skin | 0/23 |
| 809 | Female | Intergenic | X | 145177018 | T | A | Skin | 0/28 |
| 967 | Male | Intergenic | 1 | 163727963 | ATCAGGTGG | G | Skin | 0/28 |
| 967 | Male | Intergenic | 2 | 58718375 | T | C | All three | 0/50 |
| 967 | Male | Intergenic | 2 | 121280003 | G | C | All three | 0/41 |
| 967 | Male | Intergenic | 2 | 158071768 | G | A | All three | 0/42 |
| 967 | Male | Intergenic | 2 | 205132597 | C | A | All three | 0/54 |
| 967 | Male | Intergenic | 3 | 42294565 | G | A | All three | 0/45 |
| 967 | Male | Intergenic | 3 | 75864306 | C | T | All three | 0/53 |
| 967 | Male | Intergenic | 3 | 80519543 | T | G | All three | 0/51 |
| 967 | Male | Intergenic | 4 | 3793114 | T | C | All three | 0/46 |
| 967 | Male | Intergenic | 4 | 45403029 | A | G | All three | 0/43 |
| 967 | Male | Intergenic | 6 | 70264933 | ATC | A | Skin | 0/60 |
| 967 | Male | Intergenic | 6 | 148384255 | T | C | All three | 0/49 |
| 967 | Male | Intergenic | 7 | 42518541 | G | A | All three | 0/59 |
| 967 | Male | Intergenic | 8 | 29283236 | C | T | All three | 0/49 |
| 967 | Male | Intergenic | 8 | 35788470 | T | C | All three | 0/46 |
| 967 | Male | Intergenic | 8 | 109375130 | G | A | All three | 0/51 |
| 967 | Male | Intergenic | 8 | 137476647 | C | T | Skin | 0/44 |
| 967 | Male | Intergenic | 9 | 87643765 | A | G | All three | 0/52 |

**TABLE S3**

| CDH Proband Family | Mother | Proband Blood | Proband Skin | Proband Diaphragm |
|---|---|---|---|---|
| 809 | 0/62 | 20/81 | 24/64 | 12/99 |
| 809 | 0/40 | 0/57 | 31/69 | 0/62 |
| 809 | 0/55 | 0/65 | 20/69 | 0/68 |
| 809 | 0/57 | 26/69 | 35/68 | 38/83 |
| 809 | 0/66 | 40/73 | 39/66 | 36/76 |
| 809 | 0/65 | 33/65 | 29/60 | 30/59 |
| 809 | 0/65 | 36/69 | 26/59 | 27/75 |
| 809 | 0/56 | 0/54 | 31/77 | 0/49 |
| 809 | 0/46 | 0/40 | 20/74 | 0/47 |
| 809 | 0/60 | 0/60 | 16/59 | 0/58 |
| 809 | 0/51 | 0/54 | 19/80 | 0/57 |
| 809 | 0/58 | 0/54 | 24/77 | 0/60 |
| 967 | 0/27 | 0/38 | 14/70 | 0/24 |
| 967 | 0/48 | 32/57 | 40/79 | 24/52 |
| 967 | 0/50 | 19/41 | 31/64 | 19/35 |
| 967 | 0/66 | 19/47 | 34/67 | 33/58 |
| 967 | 0/58 | 27/52 | 29/64 | 29/45 |
| 967 | 0/51 | 34/53 | 28/60 | 25/35 |
| 967 | 0/53 | 33/64 | 37/75 | 19/43 |
| 967 | 0/56 | 31/56 | 40/83 | 38/61 |
| 967 | 0/51 | 30/51 | 40/79 | 16/34 |
| 967 | 0/36 | 30/54 | 18/34 | 15/38 |
| 967 | 0/60 | 0/61 | 8/40 | 0/42 |
| 967 | 0/56 | 26/47 | 38/74 | 25/48 |
| 967 | 0/54 | 34/62 | 35/75 | 27/52 |
| 967 | 0/38 | 28/60 | 34/60 | 15/36 |
| 967 | 0/43 | 30/59 | 34/71 | 30/61 |
| 967 | 0/52 | 34/84 | 31/54 | 33/58 |
| 967 | 0/53 | 0/64 | 18/73 | 0/53 |
| 967 | 0/46 | 25/54 | 20/49 | 34/46 |

De Novo Intergenic Skin

**TABLE S3**

| CDH Proband Family | Sex | Location | Chromosome | Position | Reference allele | Alternate allele | Tissue found in | Father |
|---|---|---|---|---|---|---|---|---|
| 967 | Male | Intergenic | 10 | 9433109 | CCTT | C | All three | 0/57 |
| 967 | Male | Intergenic | 10 | 20971557 | T | G | All three | 0/55 |
| 967 | Male | Intergenic | 10 | 35283939 | A | G | All three | 0/52 |
| 967 | Male | Intergenic | 10 | 114676907 | A | AT | All three | 0/50 |
| 967 | Male | Intergenic | 11 | 35124820 | C | T | All three | 0/58 |
| 967 | Male | Intergenic | 11 | 80173545 | G | A | All three | 0/51 |
| 967 | Male | Intergenic | 14 | 101628801 | GA | G | All three | 0/49 |
| 967 | male | Intergenic | 15 | 25739074 | C | T | Diaphragm+Skin | 0/64 |
| 967 | Male | Intergenic | 15 | 73336781 | C | A | All three | 0/49 |
| 967 | Male | Intergenic | 17 | 39613324 | T | C | All three | 0/41 |
| 967 | Male | Intergenic | 17 | 51582010 | A | G | All three | 0/41 |
| 967 | Male | Intergenic | 22 | 25083212 | TACTC | T | All three | 0/47 |
| 967 | Male | Intergenic | X | 35867136 | G | A | All three | 0/23 |
| 967 | male | Intergenic | X | 68010485 | C | T | Diaphragm+Skin | 0/29 |
| 967 | Male | Intergenic | X | 90667481 | T | TA | Skin | 0/36 |
| 967 | Male | Intergenic | X | 128842994 | A | G | Skin | 0/38 |
| 967 | Male | Intergenic | X | 137019473 | C | G | Skin | 0/33 |
| 967 | Male | Intergenic | Y | 13317016 | G | A | All three | 0/36 |
| 967 | Male | Intergenic | Y | 17597506 | G | GA | All three | 0/32 |

**TABLE S3**

| CDH Proband Family | Mother | Proband Blood | Proband Skin | Proband Diaphragm |
|---|---|---|---|---|
| 967 | 0/63 | 31/62 | 36/67 | 14/36 |
| 967 | 0/47 | 31/54 | 39/64 | 18/42 |
| 967 | 0/53 | 17/38 | 35/68 | 13/32 |
| 967 | 0/61 | 13/29 | 20/35 | 11/19 |
| 967 | 0/62 | 30/60 | 28/51 | 22/44 |
| 967 | 0/59 | 22/50 | 32/68 | 25/50 |
| 967 | 0/58 | 37/63 | 32/69 | 31/52 |
| 967 | 0/83 | 6/60 | 16/82 | 15/58 |
| 967 | 0/48 | 24/59 | 34/61 | 30/57 |
| 967 | 0/64 | 32/58 | 40/67 | 23/52 |
| 967 | 0/47 | 31/53 | 37/67 | 22/38 |
| 967 | 0/64 | 23/60 | 32/56 | 24/48 |
| 967 | 0/47 | 29/29 | 37/37 | 13/13 |
| 967 | 0/60 | 2/30 | 18/40 | 5/22 |
| 967 | 4/59 | 0/27 | 5/25 | 0/17 |
| 967 | 0/44 | 0/32 | 10/44 | 0/21 |
| 967 | 0/50 | 0/32 | 5/25 | 0/25 |
| 967 | 0/28 | 23/36 | 33/48 | 16/38 |
| 967 | 0/0 | 24/25 | 27/29 | 15/15 |

**TABLE S3**

| CDH Proband Family | Sex | Location | Chromosome | Position | Reference allele | Alternate allele | Tissue found in | Father |
|---|---|---|---|---|---|---|---|---|
| 411 | Male | Intergenic | 2 | 6448411 | G | A | All three | 0/51 |
| 411 | Male | Intergenic | 2 | 15979430 | C | T | All three | 0/46 |
| 411 | Male | Intergenic | 2 | 23607772 | C | A | All three | 0/34 |
| 411 | Male | Intergenic | 2 | 23607823 | G | A | All three | 0/34 |
| 411 | Male | Intergenic | 2 | 48488807 | CTTTGAAAG | T | All three | 0/55 |
| 411 | male | Intergenic | 4 | 12542452 | ACAAAAAT | A | All three | 0/61 |
| 411 | Male | Intergenic | 4 | 40268400 | A | G | All three | 0/59 |
| 411 | Male | Intergenic | 4 | 130485536 | C | T | All three | 0/39 |
| 411 | Male | Intergenic | 5 | 30972421 | G | T | All three | 0/54 |
| 411 | Male | Intergenic | 5 | 43876751 | A | T | All three | 0/43 |
| 411 | Male | Intergenic | 7 | 63891470 | TTGGCCGGG | A | Blood | 0/44 |
| 411 | Male | Intergenic | 7 | 121415957 | T | G | All three | 0/42 |
| 411 | Male | Intergenic | 8 | 49656228 | G | C | All three | 0/45 |
| 411 | Male | Intergenic | 9 | 24573667 | A | C | All three | 0/57 |
| 411 | Male | Intergenic | 10 | 45696971 | C | T | All three | 0/44 |
| 411 | male | Intergenic | 10 | 107240148 | T | C | All three | 0/54 |
| 411 | Male | Intergenic | 11 | 48522238 | T | C | All three | 0/53 |
| 411 | Male | Intergenic | 12 | 51286910 | T | C | All three | 0/46 |
| 411 | Male | Intergenic | 13 | 63562946 | G | GA | All three | 0/63 |
| 411 | Male | Intergenic | 15 | 38941044 | C | T | All three | 0/45 |
| 411 | Male | Intergenic | 15 | 46407038 | C | T | All three | 0/55 |
| 411 | Male | Intergenic | 16 | 75543055 | C | A | All three | 0/46 |
| 411 | Male | Intergenic | 17 | 404337 | C | T | All three | 0/55 |
| 411 | Male | Intergenic | 20 | 5206920 | C | A | All three | 0/45 |
| 411 | Male | Intergenic | Y | 23424401 | T | C | All three | 0/37 |
| 716 | Female | Intergenic | 1 | 25520943 | G | A | All three | 0/53 |
| 716 | Female | Intergenic | 1 | 108550027 | G | A | All three | 0/55 |
| 716 | Female | Intergenic | 1 | 188119862 | C | T | All three | 0/60 |
| 716 | Female | Intergenic | 1 | 219133910 | G | A | All three | 0/54 |
| 716 | Female | Intergenic | 1 | 241538742 | A | G | All three | 0/64 |

**TABLE S3**

| CDH Proband Family | Mother | Proband Blood | Proband Skin | Proband Diaphragm |
|---|---|---|---|---|
| 411 | 0/49 | 29/59 | 21/54 | 51/89 |
| 411 | 0/38 | 16/47 | 23/50 | 36/63 |
| 411 | 0/39 | 16/26 | 25/40 | 23/56 |
| 411 | 0/47 | 16/21 | 17/25 | 17/49 |
| 411 | 0/46 | 23/50 | 22/60 | 32/70 |
| 411 | 0/67 | 23/54 | 23/50 | 32/86 |
| 411 | 0/56 | 36/75 | 30/55 | 38/78 |
| 411 | 0/55 | 34/67 | 23/58 | 36/75 |
| 411 | 0/52 | 28/65 | 34/65 | 41/99 |
| 411 | 0/43 | 37/78 | 32/59 | 21/55 |
| 411 | 0/35 | 17/67 | 0/57 | 4/82 |
| 411 | 0/50 | 25/62 | 26/61 | 42/81 |
| 411 | 0/47 | 31/57 | 21/58 | 42/75 |
| 411 | 0/45 | 24/69 | 18/48 | 32/78 |
| 411 | 0/59 | 33/70 | 31/65 | 29/60 |
| 411 | 0/58 | 32/79 | 25/58 | 33/71 |
| 411 | 0/41 | 26/61 | 24/69 | 35/85 |
| 411 | 0/47 | 57/97 | 34/58 | 35/80 |
| 411 | 0/51 | 19/48 | 20/40 | 33/68 |
| 411 | 0/48 | 28/58 | 31/72 | 37/75 |
| 411 | 0/43 | 34/58 | 22/57 | 36/77 |
| 411 | 0/43 | 22/56 | 30/53 | 41/79 |
| 411 | 0/50 | 36/73 | 28/49 | 38/77 |
| 411 | 0/43 | 26/57 | 26/67 | 35/77 |
| 411 | 0/0 | 35/35 | 35/36 | 31/31 |
| 716 | 0/58 | 24/50 | 40/84 | 23/51 |
| 716 | 0/58 | 38/66 | 36/64 | 33/66 |
| 716 | 0/65 | 27/61 | 38/66 | 23/56 |
| 716 | 0/50 | 37/69 | 28/56 | 29/66 |
| 716 | 0/58 | 40/85 | 46/78 | 33/67 |

**TABLE S3**

| CDH Proband Family | Sex | Location | Chromosome | Position | Reference allele | Alternate allele | Tissue found in | Father |
|---|---|---|---|---|---|---|---|---|
| 716 | Female | Intergenic | 2 | 60131252 | T | C | All three | 0/56 |
| 716 | Female | Intergenic | 2 | 237733339 | C | T | All three | 0/48 |
| 716 | Female | Intergenic | 2 | 241273592 | G | T | All three | 0/48 |
| 716 | Female | Intergenic | 3 | 31317266 | C | T | All three | 0/65 |
| 716 | Female | Intergenic | 3 | 70707930 | T | C | All three | 0/51 |
| 716 | Female | Intergenic | 3 | 87755171 | C | T | All three | 0/51 |
| 716 | Female | Intergenic | 3 | 96134789 | G | A | All three | 0/58 |
| 716 | Female | Intergenic | 3 | 116969700 | C | T | All three | 0/61 |
| 716 | Female | Intergenic | 4 | 76738805 | A | C | All three | 0/55 |
| 716 | Female | Intergenic | 4 | 90360867 | C | G | All three | 0/45 |
| 716 | Female | Intergenic | 4 | 92765694 | T | A | All three | 0/55 |
| 716 | Female | Intergenic | 4 | 165797047 | C | T | All three | 0/49 |
| 716 | Female | Intergenic | 5 | 13079144 | A | T | All three | 0/45 |
| 716 | Female | Intergenic | 5 | 92381616 | C | G | All three | 0/62 |
| 716 | Female | Intergenic | 6 | 18809404 | A | C | All three | 0/66 |
| 716 | Female | Intergenic | 6 | 50605045 | A | G | All three | 0/57 |
| 716 | Female | Intergenic | 6 | 69243495 | C | G | All three | 0/41 |
| 716 | Female | Intergenic | 6 | 78773081 | C | T | All three | 0/62 |
| 716 | Female | Intergenic | 7 | 2657275 | C | T | All three | 0/47 |
| 716 | Female | Intergenic | 7 | 6718899 | A | G | All three | 0/68 |
| 716 | Female | Intergenic | 7 | 19285491 | C | T | All three | 0/50 |
| 716 | Female | Intergenic | 7 | 96000892 | G | T | All three | 0/46 |
| 716 | Female | Intergenic | 8 | 23549713 | A | G | All three | 0/50 |
| 716 | Female | Intergenic | 8 | 50342185 | C | A | All three | 0/55 |
| 716 | Female | Intergenic | 8 | 53690025 | G | A | All three | 0/65 |
| 716 | Female | Intergenic | 8 | 121394186 | C | G | All three | 0/43 |
| 716 | Female | Intergenic | 9 | 37870079 | C | A | All three | 0/52 |
| 716 | Female | Intergenic | 9 | 92940311 | C | T | All three | 0/62 |
| 716 | Female | Intergenic | 10 | 21531548 | A | T | All three | 0/61 |
| 716 | female | Intergenic | 10 | 78610342 | GACAA | G | All three | 0/79 |

**TABLE S3**

| CDH Proband Family | Mother | Proband Blood | Proband Skin | Proband Diaphragm |
|---|---|---|---|---|
| 716 | 0/57 | 28/55 | 43/90 | 49/79 |
| 716 | 0/51 | 29/68 | 32/76 | 39/74 |
| 716 | 0/54 | 42/89 | 30/77 | 26/55 |
| 716 | 0/56 | 34/61 | 26/59 | 39/77 |
| 716 | 0/50 | 26/62 | 32/61 | 35/66 |
| 716 | 0/59 | 35/82 | 34/67 | 38/66 |
| 716 | 0/60 | 31/58 | 46/81 | 23/58 |
| 716 | 0/72 | 35/79 | 37/71 | 23/52 |
| 716 | 0/56 | 26/54 | 21/45 | 45/67 |
| 716 | 0/61 | 29/59 | 25/53 | 29/60 |
| 716 | 0/46 | 35/69 | 40/63 | 31/63 |
| 716 | 0/57 | 39/68 | 34/69 | 39/72 |
| 716 | 0/59 | 27/60 | 36/63 | 28/51 |
| 716 | 0/59 | 40/72 | 27/57 | 32/57 |
| 716 | 0/68 | 30/68 | 33/77 | 22/47 |
| 716 | 0/53 | 39/75 | 28/58 | 36/56 |
| 716 | 0/61 | 26/53 | 31/72 | 35/69 |
| 716 | 0/58 | 38/79 | 36/62 | 38/76 |
| 716 | 0/57 | 30/58 | 23/54 | 29/67 |
| 716 | 0/55 | 34/75 | 35/78 | 34/68 |
| 716 | 0/63 | 36/70 | 38/73 | 41/77 |
| 716 | 0/52 | 23/83 | 19/64 | 29/71 |
| 716 | 0/47 | 17/55 | 34/68 | 37/76 |
| 716 | 0/55 | 21/47 | 42/77 | 48/85 |
| 716 | 0/54 | 25/44 | 26/66 | 30/58 |
| 716 | 0/43 | 20/50 | 38/72 | 31/54 |
| 716 | 0/52 | 29/68 | 31/71 | 35/66 |
| 716 | 0/56 | 28/66 | 37/73 | 38/77 |
| 716 | 0/56 | 33/61 | 37/73 | 30/61 |
| 716 | 0/80 | 38/79 | 35/88 | 45/91 |

**TABLE S3**

| CDH Proband Family | Sex | Location | Chromosome | Position | Reference allele | Alternate allele | Tissue found in | Father |
|---|---|---|---|---|---|---|---|---|
| 716 | Female | Intergenic | 10 | 102805027 | C | T | All three | 0/49 |
| 716 | Female | Intergenic | 10 | 111937754 | A | G | All three | 0/55 |
| 716 | Female | Intergenic | 11 | 68910808 | C | T | All three | 0/52 |
| 716 | Female | Intergenic | 11 | 68917454 | A | T | All three | 0/63 |
| 716 | Female | Intergenic | 11 | 85940710 | C | T | All three | 0/52 |
| 716 | Female | Intergenic | 12 | 2894972 | G | A | All three | 0/71 |
| 716 | Female | Intergenic | 12 | 20253658 | G | A | All three | 0/60 |
| 716 | Female | Intergenic | 12 | 61786788 | C | T | All three | 0/56 |
| 716 | female | Intergenic | 12 | 73894281 | T | C | All three | 0/67 |
| 716 | Female | Intergenic | 12 | 81133670 | T | C | All three | 0/44 |
| 716 | Female | Intergenic | 12 | 128640569 | T | C | All three | 0/60 |
| 716 | Female | Intergenic | 13 | 38770384 | A | T | All three | 0/56 |
| 716 | Female | Intergenic | 14 | 94631627 | C | T | All three | 0/47 |
| 716 | Female | Intergenic | 14 | 98723131 | C | T | All three | 0/57 |
| 716 | Female | Intergenic | 15 | 53418167 | A | G | All three | 0/59 |
| 716 | Female | Intergenic | 18 | 26978994 | C | T | All three | 0/53 |
| 716 | Female | Intergenic | 18 | 48313755 | G | T | All three | 0/60 |
| 716 | Female | Intergenic | 18 | 63851839 | G | C | All three | 0/55 |
| 716 | Female | Intergenic | 19 | 12407418 | A | C | All three | 0/78 |
| 716 | Female | Intergenic | 20 | 40498713 | A | T | All three | 0/41 |
| 716 | Female | Intergenic | 21 | 26441156 | G | T | All three | 0/53 |
| 716 | Female | Intergenic | 21 | 29172156 | A | T | All three | 0/59 |
| 716 | Female | Intergenic | X | 39689788 | C | G | All three | 0/31 |
| 716 | Female | Intergenic | X | 65140289 | G | A | All three | 0/25 |
| 716 | Female | Intergenic | X | 82221769 | C | T | All three | 0/30 |
| 716 | Female | Intergenic | X | 88110135 | C | T | All three | 0/34 |
| 716 | Female | Intergenic | X | 89554886 | C | T | All three | 0/25 |
| 716 | Female | Intergenic | X | 105821982 | G | T | All three | 0/35 |
| 809 | Female | Intergenic | 1 | 229943691 | T | A | All three | 0/41 |
| 809 | Female | Intergenic | 1 | 248631521 | G | A | All three | 0/47 |

**TABLE S3**

| CDH Proband Family | Mother | Proband Blood | Proband Skin | Proband Diaphragm |
|---|---|---|---|---|
| 716 | 0/49 | 40/77 | 45/80 | 36/73 |
| 716 | 0/47 | 38/77 | 43/81 | 43/68 |
| 716 | 0/59 | 34/72 | 36/71 | 27/59 |
| 716 | 0/52 | 40/72 | 35/69 | 36/65 |
| 716 | 0/53 | 28/63 | 38/76 | 31/57 |
| 716 | 0/58 | 47/80 | 31/61 | 38/76 |
| 716 | 0/64 | 39/77 | 29/50 | 43/61 |
| 716 | 0/74 | 36/76 | 34/67 | 36/61 |
| 716 | 0/77 | 43/81 | 40/73 | 33/71 |
| 716 | 0/63 | 37/65 | 38/72 | 30/70 |
| 716 | 0/64 | 22/57 | 38/73 | 35/76 |
| 716 | 0/42 | 28/57 | 30/63 | 32/61 |
| 716 | 0/61 | 30/55 | 31/66 | 23/44 |
| 716 | 0/55 | 34/65 | 32/66 | 30/67 |
| 716 | 0/60 | 24/67 | 30/66 | 35/66 |
| 716 | 0/49 | 28/62 | 34/74 | 38/74 |
| 716 | 0/53 | 35/64 | 33/71 | 40/74 |
| 716 | 0/54 | 38/76 | 41/83 | 33/67 |
| 716 | 0/74 | 46/68 | 28/54 | 44/62 |
| 716 | 0/46 | 28/68 | 37/68 | 25/59 |
| 716 | 0/58 | 27/62 | 41/73 | 28/59 |
| 716 | 0/62 | 44/74 | 34/84 | 30/58 |
| 716 | 0/57 | 31/66 | 24/54 | 34/81 |
| 716 | 0/47 | 40/79 | 32/68 | 31/70 |
| 716 | 0/55 | 39/68 | 37/69 | 34/54 |
| 716 | 0/63 | 35/64 | 36/69 | 35/68 |
| 716 | 0/44 | 34/72 | 25/57 | 35/82 |
| 716 | 0/48 | 35/68 | 39/66 | 33/61 |
| 809 | 0/51 | 27/73 | 43/70 | 29/61 |
| 809 | 0/52 | 26/60 | 35/61 | 40/78 |

**TABLE S3**

| CDH Proband Family | Sex | Location | Chromosome | Position | Reference allele | Alternate allele | Tissue found in | Father |
|---|---|---|---|---|---|---|---|---|
| 809 | female | Intergenic | 2 | 52087068 | T | A | Diaphragm+Blood | 1/49 |
| 809 | Female | Intergenic | 2 | 118456374 | T | C | All three | 0/46 |
| 809 | female | Intergenic | 2 | 165063447 | G | A | Skin+Blood | 0/64 |
| 809 | female | Intergenic | 3 | 35466073 | C | T | Skin+Blood | 0/53 |
| 809 | Female | Intergenic | 3 | 74259109 | A | G | All three | 0/39 |
| 809 | Female | Intergenic | 4 | 8715501 | G | A | All three | 0/44 |
| 809 | Female | Intergenic | 4 | 59397258 | C | T | All three | 0/48 |
| 809 | female | Intergenic | 5 | 132509196 | T | C | All three | 0/59 |
| 809 | Female | Intergenic | 5 | 144084315 | G | GA | All three | 0/40 |
| 809 | Female | Intergenic | 6 | 84499729 | T | C | All three | 0/43 |
| 809 | Female | Intergenic | 7 | 46969823 | T | A | All three | 0/39 |
| 809 | Female | Intergenic | 8 | 24834454 | C | T | All three | 0/50 |
| 809 | Female | Intergenic | 8 | 60040127 | C | T | All three | 0/47 |
| 809 | female | Intergenic | 8 | 138869304 | T | G | Skin+Blood | 1/62 |
| 809 | Female | Intergenic | 10 | 4733230 | A | G | All three | 0/44 |
| 809 | Female | Intergenic | 11 | 15343128 | AT | A | All three | 0/39 |
| 809 | Female | Intergenic | 11 | 37638703 | G | C | All three | 0/55 |
| 809 | Female | Intergenic | 11 | 112170508 | T | C | All three | 0/45 |
| 809 | Female | Intergenic | 12 | 19007497 | G | A | All three | 0/40 |
| 809 | Female | Intergenic | 12 | 110872268 | A | C | All three | 0/55 |
| 809 | Female | Intergenic | 13 | 26669122 | G | T | All three | 0/45 |
| 809 | Female | Intergenic | 13 | 69271119 | T | C | All three | 0/43 |
| 809 | Female | Intergenic | 14 | 34632380 | C | T | All three | 0/48 |
| 809 | Female | Intergenic | 15 | 47132793 | C | T | All three | 0/44 |
| 809 | Female | Intergenic | 16 | 1290232 | C | T | All three | 0/24 |
| 809 | Female | Intergenic | 17 | 14274664 | T | A | All three | 0/41 |
| 809 | female | Intergenic | 20 | 46859236 | C | T | Skin+Blood | 0/58 |
| 809 | Female | Intergenic | 21 | 18844527 | G | C | All three | 0/56 |
| 809 | Female | Intergenic | 21 | 20700243 | C | T | All three | 0/44 |
| 809 | Female | Intergenic | 21 | 28403305 | T | A | All three | 0/60 |

De Novo Intergenic Blood

**TABLE S3**

| CDH Proband Family | Mother | Proband Blood | Proband Skin | Proband Diaphragm |
|---|---|---|---|---|
| 809 | 0/88 | 19/64 | 4/52 | 37/79 |
| 809 | 0/56 | 26/65 | 35/71 | 39/71 |
| 809 | 0/78 | 19/83 | 26/93 | 12/82 |
| 809 | 0/66 | 19/89 | 19/68 | 11/80 |
| 809 | 0/59 | 34/70 | 33/59 | 30/64 |
| 809 | 0/58 | 42/69 | 40/76 | 33/73 |
| 809 | 0/77 | 41/70 | 43/67 | 31/58 |
| 809 | 0/73 | 46/74 | 38/78 | 41/89 |
| 809 | 0/66 | 14/70 | 27/53 | 12/58 |
| 809 | 0/50 | 48/75 | 29/59 | 42/91 |
| 809 | 0/57 | 39/82 | 31/49 | 35/73 |
| 809 | 0/62 | 14/29 | 13/29 | 18/35 |
| 809 | 0/55 | 27/65 | 37/80 | 43/88 |
| 809 | 0/68 | 23/77 | 26/74 | 12/74 |
| 809 | 0/59 | 34/60 | 21/73 | 21/58 |
| 809 | 0/64 | 35/78 | 34/67 | 42/70 |
| 809 | 0/44 | 38/70 | 35/71 | 27/55 |
| 809 | 0/67 | 30/55 | 35/67 | 23/69 |
| 809 | 0/64 | 40/73 | 32/76 | 36/66 |
| 809 | 0/58 | 25/50 | 32/67 | 43/81 |
| 809 | 0/65 | 30/70 | 34/51 | 38/71 |
| 809 | 0/49 | 39/74 | 33/74 | 34/71 |
| 809 | 0/55 | 36/63 | 28/57 | 26/58 |
| 809 | 0/55 | 32/63 | 32/66 | 43/91 |
| 809 | 0/46 | 34/52 | 32/50 | 31/56 |
| 809 | 0/62 | 43/70 | 34/73 | 38/70 |
| 809 | 0/62 | 20/81 | 24/64 | 12/99 |
| 809 | 0/57 | 26/69 | 35/68 | 38/83 |
| 809 | 0/66 | 40/73 | 39/66 | 36/76 |
| 809 | 0/65 | 33/65 | 29/60 | 30/59 |

**TABLE S3**

| CDH Proband Family | Sex | Location | Chromosome | Position | Reference allele | Alternate allele | Tissue found in | Father |
|---|---|---|---|---|---|---|---|---|
| 809 | Female | Intergenic | X | 455196 | C | T | All three | 0/49 |
| 967 | Male | Intergenic | 2 | 58718375 | T | C | All three | 0/50 |
| 967 | Male | Intergenic | 2 | 121280003 | G | C | All three | 0/41 |
| 967 | Male | Intergenic | 2 | 158071768 | G | A | All three | 0/42 |
| 967 | Male | Intergenic | 2 | 205132597 | C | A | All three | 0/54 |
| 967 | Male | Intergenic | 3 | 42294565 | G | A | All three | 0/45 |
| 967 | Male | Intergenic | 3 | 75864306 | C | T | All three | 0/53 |
| 967 | Male | Intergenic | 3 | 80519543 | T | G | All three | 0/51 |
| 967 | Male | Intergenic | 4 | 3793114 | T | C | All three | 0/46 |
| 967 | Male | Intergenic | 4 | 45403029 | A | G | All three | 0/43 |
| 967 | Male | Intergenic | 6 | 148384255 | T | C | All three | 0/49 |
| 967 | Male | Intergenic | 7 | 42518541 | G | A | All three | 0/59 |
| 967 | Male | Intergenic | 8 | 29283236 | C | T | All three | 0/49 |
| 967 | Male | Intergenic | 8 | 35788470 | T | C | All three | 0/46 |
| 967 | Male | Intergenic | 8 | 109375130 | G | A | All three | 0/51 |
| 967 | Male | Intergenic | 9 | 87643765 | A | G | All three | 0/52 |
| 967 | Male | Intergenic | 10 | 9433109 | CCTT | C | All three | 0/57 |
| 967 | Male | Intergenic | 10 | 20971557 | T | G | All three | 0/55 |
| 967 | Male | Intergenic | 10 | 35283939 | A | G | All three | 0/52 |
| 967 | Male | Intergenic | 10 | 114676907 | A | AT | All three | 0/50 |
| 967 | Male | Intergenic | 11 | 35124820 | C | T | All three | 0/58 |
| 967 | Male | Intergenic | 11 | 80173545 | G | A | All three | 0/51 |
| 967 | Male | Intergenic | 14 | 101628801 | GA | G | All three | 0/49 |
| 967 | Male | Intergenic | 15 | 73336781 | C | A | All three | 0/49 |
| 967 | Male | Intergenic | 17 | 39613324 | T | C | All three | 0/41 |
| 967 | Male | Intergenic | 17 | 51582010 | A | G | All three | 0/41 |
| 967 | Male | Intergenic | 22 | 25083212 | TACTC | T | All three | 0/47 |
| 967 | Male | Intergenic | X | 25378699 | CAT | C | Blood | 0/34 |
| 967 | Male | Intergenic | X | 35867136 | G | A | All three | 0/23 |
| 967 | Male | Intergenic | Y | 13317016 | G | A | All three | 0/36 |

De Novo Intergenic Blood

| CDH Proband Family | Mother | Proband Blood | Proband Skin | Proband Diaphragm |
|---|---|---|---|---|
| 809 | 0/65 | 36/69 | 26/59 | 27/75 |
| 967 | 0/48 | 32/57 | 40/79 | 24/52 |
| 967 | 0/50 | 19/41 | 31/64 | 19/35 |
| 967 | 0/66 | 19/47 | 34/67 | 33/58 |
| 967 | 0/58 | 27/52 | 29/64 | 29/45 |
| 967 | 0/51 | 34/53 | 28/60 | 25/35 |
| 967 | 0/53 | 33/64 | 37/75 | 19/43 |
| 967 | 0/56 | 31/56 | 40/83 | 38/61 |
| 967 | 0/51 | 30/51 | 40/79 | 16/34 |
| 967 | 0/36 | 30/54 | 18/34 | 15/38 |
| 967 | 0/56 | 26/47 | 38/74 | 25/48 |
| 967 | 0/54 | 34/62 | 35/75 | 27/52 |
| 967 | 0/38 | 28/60 | 34/60 | 15/36 |
| 967 | 0/43 | 30/59 | 34/71 | 30/61 |
| 967 | 0/52 | 34/84 | 31/54 | 33/58 |
| 967 | 0/46 | 25/54 | 20/49 | 34/46 |
| 967 | 0/63 | 31/62 | 36/67 | 14/36 |
| 967 | 0/47 | 31/54 | 39/64 | 18/42 |
| 967 | 0/53 | 17/38 | 35/68 | 13/32 |
| 967 | 0/61 | 13/29 | 20/35 | 43789 |
| 967 | 0/62 | 30/60 | 28/51 | 22/44 |
| 967 | 0/59 | 22/50 | 32/68 | 25/50 |
| 967 | 0/58 | 37/63 | 32/69 | 31/52 |
| 967 | 0/48 | 24/59 | 34/61 | 30/57 |
| 967 | 0/64 | 32/58 | 40/67 | 23/52 |
| 967 | 0/47 | 31/53 | 37/67 | 22/38 |
| 967 | 0/64 | 23/60 | 32/56 | 24/48 |
| 967 | 0/58 | 5/17 | 0/38 | 0/20 |
| 967 | 0/47 | 29/29 | 37/37 | 13/13 |
| 967 | 0/28 | 23/36 | 33/48 | 16/38 |

**TABLE S3**

| CDH Proband Family | Sex | Location | Chromosome | Position | Reference allele | Alternate allele | Tissue found in | Father |
|---|---|---|---|---|---|---|---|---|
| 967 | Male | Intergenic | Y | 17597506 | G | GA | All three | 0/32 |

De Novo Intergenic Blood

| CDH Proband Family | Mother | Proband Blood | Proband Skin | Proband Diaphragm |
|---|---|---|---|---|
| 967 | 0/0 | 24/25 | 27/29 | 15/15 |

### TABLE S4

| CDH Proband | Sex | Gene Name | Gene Full Name | Location in Gene | Chromosome | Chromosome Position HG19 | Reference Allele | Alternate Allele | Parent found in | VAAST Score |
|---|---|---|---|---|---|---|---|---|---|---|
| 411 | Male | UPF3A | Regulator of nonsense mediated mRNA decay | Exon 5 | 13 | 115052021 | C | A | Father | 26.69342 |
| 411 | Male | UPF3A | Regulator of nonsense mediated mRNA decay | Exon 8 | 13 | 115064333 | A | G | Mother | 26.69342 |
| 716 | Female | ALG2 | Alpha-1,3/1,6-mannosyltransferase | Exon 1 | 9 | 101983960 | C | G | Father | 26.08287 |
| 716 | Female | ALG2 | Alpha-1,3/1,6-mannosyltransferase | Exon 1 | 9 | 101984010 | G | A | Mother | 26.08287 |
| 716 | Female | HRC | Sarcoplasmic reticulum histidine-rich calcium-binding protein | Exon 1 | 19 | 49657569 | T | A | Mother | 28.64456 |
| 716 | Female | HRC | Sarcoplasmic reticulum histidine-rich calcium-binding protein | Exon 1 | 19 | 49657221 | T | C | Father | 28.64456 |
| 716 | Female | AHNAK | Neuroblast differentiation-associated protein AHNAK | Exon 5 | 11 | 62292741 | C | G | Mother | 26.46372 |
| 716 | Female | AHNAK | Neuroblast differentiation-associated protein AHNAK | Exon 5 | 11 | 62293913 | T | G | Mother | 26.46372 |
| 716 | Female | MYO1H | Unconventional Myosin IH | Exon 3 | 12 | 109834353 | GT | G | Father | 30.64254 |
| 716 | Female | MYO1H | Unconventional Myosin IH | Exon 23 | 12 | 109877506 | C | T | Mother | 30.64254 |
| 716 | Female | MPEG1 | Macrophage-expressed gene 1 | Exon 1 | 11 | 58980189 | GT | GCC | Mother | 29.24719 |
| 716 | Female | MPEG1 | Macrophage-expressed gene 1 | Exon 1 | 11 | 58979643 | A | C | Father | 29.24719 |
| 716 | Female | CCDC136 | coiled-coil domain containing 136 | Exon 13 | 7 | 128452280 | G | A | Mother | 37.81706 |
| 716 | Female | CCDC136 | coiled-coil domain containing 136 | Exon 10 | 7 | 128447502 | GAGCAGGACCT | A | Father | 37.81706 |
| 716 | Female | NOP9 | Nucleolar protein 9 | Exon 2 | 14 | 24769849 | A | AGAGGAG | Father | 29.46885 |
| 716 | Female | NOP9 | Nucleolar protein 9 | Exon 8 | 14 | 24773448 | C | T | Mother | 29.46885 |
| 716 | Female | ZNF646 | Zinc finger protein 646 | Exon 2 | 16 | 31088738 | G | A | Mother | 29.06799 |
| 716 | Female | ZNF646 | Zinc finger protein 646 | Exon 2 | 16 | 31090928 | C | T | Father | 29.06799 |
| 967 | Male | MYOF | Myoferlin | Exon 18 | 10 | 95148899 | T | A | Father | 40.01234 |
| 967 | Male | MYOF | Myoferlin | Exon 7 | 10 | 95168549 | C | CA | Mother | 40.01234 |
| 967 | Male | SACS | Sacsin | Exon 10 | 13 | 23910966 | T | C | Mother | 34.80669 |
| 967 | Male | SACS | Sacsin | Exon 10 | 13 | 23904275 | TCA | T | Father | 34.80669 |
| 967 | Male | ADAMTS2 | A disintegrin and metalloproteinase with thrombospondin motifs 2 | Exon 1 | 5 | 178772241 | G | GGCGGCAGGA | Both | 40.76989 |
| 967 | Male | SH3PXD2A | SH3 and PX domain-containing protein 2A | Exon 14 | 10 | 105361796 | G | T | Father | 30.6648 |
| 967 | Male | SH3PXD2A | SH3 and PX domain-containing protein 2A | Exon 14 | 10 | 105362735 | G | A | Mother | 30.6648 |

**TABLE S4**

| CDH Proband | VAAST p value | VAAST Impact | PROVEAN Prediction | ExAC Pli 1= intolerant 0= tolerant of haploinsufficency | gnomAD frequency | CADD score, 10= top 10% damaging mutations, 40= 0.01% damaging | Protein Amino Acid Change | TPM RNA-Seq PPFs | Rank |
|---|---|---|---|---|---|---|---|---|---|
| 411 | 0.025559 | missense | Neutral | 0 | 0.00008506 | 11.39 | p.Thr183Asn | 75.82683746 | 4 |
| **411** | **0.025559** | **missense** | **Deleterious** | | **0.00004338** | **25.7** | **p.Lys289Glu** | **75.82683746** | |
| **716** | **0.005192** | **missense** | **Deleterious** | 0.02 | **0.0001388** | **27.6** | **p.Asp73His** | **90.3247514** | 1 |
| **716** | **0.005192** | **missense** | **Deleterious** | | **0.0008662** | **32** | **p.Pro56Leu** | **90.3247514** | |
| **716** | **0.00639** | **missense** | **Deleterious** | 0 | **0.000439** | **17.69** | **p.Asp309Val** | **35.7001661** | 1 |
| **716** | **0.00639** | **missense** | **Deleterious** | | **0.0000875** | **16.41** | **p.Glu425Gly** | **35.7001661** | |
| **716** | **0.038738** | **missense** | **Deleterious** | 0.34 | **0.000003978** | **23.6** | **p.Asp3050His** | **45.74418807** | 2 |
| **716** | **0.038738** | **missense** | **Deleterious** | | **0.001128** | **25.5** | **p.Asp2659Ala** | **45.74418807** | |
| **716** | **0.000799** | **frameshift_variant** | **(predicted null)** | 0 | **0.00008452** | **NA** | **p.Asp137ThrfsT** | **36.39642641** | 2 |
| **716** | **0.000799** | **missense** | **Deleterious** | | **0.006348** | **27.4** | **p.Arg773Trp** | **36.39642641** | |
| **716** | **0.001597** | **frameshift_variant** | **(predicted null)** | 0 | **Not found in gno** | **NA** | **p.Trp51AlafsTer** | **21.31529659** | 3 |
| 716 | 0.001597 | missense | Neutral | | 0.00293 | 3.585 | p.Ser232Arg | 21.31529659 | |
| **716** | 0.000399 | missense | Neutral | 0 | 0.00006029 | 14.79 | p.Gly819Ser | 25.94114519 | 4 |
| **716** | **0.000399** | **inframe_deletion** | **Deleterious** | **0** | **0.00001378** | **NA** | **p.Gln507_Glu51** | **25.94114519** | |
| 716 | 0.001597 | inframe_insertion | Neutral | 0 | 0.3478 | NA | p.Glu168_Glu16 | 39.05250943 | 4 |
| **716** | **0.001597** | **missense** | **Deleterious** | | **0.001724** | **25.9** | **p.Arg538Cys** | **39.05250943** | |
| 716 | 0.000399 | missense | Neutral | 0.01 | 0.0003458 | 24.9 | p.Asp365Asn | 39.47584 | 4 |
| **716** | **0.000399** | **missense** | **Deleterious** | | **0.0002126** | **33** | **p.Arg1095Cys** | **39.47584** | |
| 967 | 0.000399 | missense | Neutral | 0 | 0.0004856 | 24.2 | p.Glu490Val | 28.96335117 | 3 |
| **967** | **0.000399** | **frameshift_variant** | **(predicted null)** | | **0.00003568** | **NA** | **p.Asp242Ter** | **28.96335117** | |
| 967 | 0.000399 | missense | Neutral | 0 | 0.00008853 | 24.5 | p.Glu2350Gly | 10.85410849 | 3 |
| **967** | **0.000399** | **frameshift_variant&stop_lost** | **(predicted null)** | | **0.00002795** | **NA** | **p.Ter4580Lysfs** | **10.85410849** | |
| 967 | 0.000399 | inframe_insertion | Neutral | **0.99** | 0.01099 | NA | p.Leu27_Pro29d | 38.9203826 | 4 |
| **967** | **0.000399** | **missense** | **Deleterious** | 0.23 | **0.00001075** | **29.1** | **p.Pro1032His** | **136.7142617** | 4 |
| 967 | 0.000399 | missense | Neutral | | 0.00006717 | 19.76 | p.Thr719Ile | 136.7142617 | |

### TABLE S4

| CDH Proband | Sex | Gene Name | Location in Gene | Chromosome | Position | Reference Allele | Alternative Allele | Parent found in | VAAST Score | VAAST p value | VAAST Impact | PROVEAN Functional Prediction |
|---|---|---|---|---|---|---|---|---|---|---|---|---|
| 411 | Male | RPS27AP18 | Intron 1 | 5 | 137602877 | CTT | C | Father | 42.296146 | 0.000399 | intron_variant | |
| 411 | Male | RPS27AP18 | Intron 1 | 5 | 137602877 | CTT | CT | Mother | 42.296146 | 0.000399 | intron_variant | |
| 411 | Male | KRT18 | Exon 1 | 12 | 53343084 | G | T | Mother | 34.909542 | 0.000399 | missense | Deleterious |
| 411 | Male | KRT18 | Exon 1 | 12 | 53342968 | C | T | Father | 34.909542 | 0.000399 | missense | Neutral |
| 411 | Male | LRRTM4 | Exon 4 | 2 | 76975931 | G | C | Both | 33.070103 | 0.000399 | missense | Neutral |
| 411 | Male | POM121L7P | Exon 1 | 22 | 21481561 | A | G | Mother | 32.301556 | 0.000399 | missense | Deleterious |
| 411 | Male | POM121L7P | Exon 1 | 22 | 21481720 | G | C | Father | 32.301556 | 0.000399 | missense | Neutral |
| 411 | Male | HLA-B | Exon 2 | 6 | 31324489 | C | A | Mother | 29.024973 | 0.000399 | missense | Deleterious |
| 411 | Male | HLA-B | Exon 2 | 6 | 31324725 | G | A | Father | 29.024973 | 0.000399 | missense | Deleterious |
| 411 | Male | LCE4A | Exon 1 | 1 | 152681680 | C | CTGGGGGCT | Both | 26.935133 | 0.006789 | inframe_insertion | Neutral |
| 411 | Male | **UPF3A** | Exon 5 | 13 | 115052021 | C | A | Father | 26.69342 | 0.025559 | missense | Neutral |
| 411 | Male | **UPF3A** | Exon 8 | 13 | 115064333 | A | G | Mother | 26.69342 | 0.025559 | missense | Deleterious |
| 411 | Male | AC092953.1 | Intron 2 | 3 | 182673378 | GT | G | Father | 26.253948 | 0.005591 | intron_variant | |
| 411 | Male | AC092953.1 | Intron 2 | 3 | 182673717 | C | CT | Mother | 26.253948 | 0.005591 | intron_variant | |
| 411 | Male | HLA-DRB1 | Exon 2 | 6 | 32552128 | TCCC | T | Mother | 25.645077 | 0.002396 | inframe_deletion | Deleterious |
| 411 | Male | HLA-DRB1 | Exon 3 | 6 | 32549584 | C | CT | Father | 25.645077 | 0.002396 | frameshift_variant | |
| 411 | Male | HRNR | Exon 3 | 1 | 152187310 | C | CATGT | Mother | 25.0231 | 0.044728 | frameshift_variant | |
| 411 | Male | HRNR | Exon 3 | 1 | 152190945 | C | T | Father | 25.0231 | 0.044728 | missense | Neutral |
| 411 | Male | HLA-DRB5 | Exon 2 | 6 | 32489925 | C | T | Mother | 24.031416 | 0.006789 | missense | Deleterious |
| 411 | Male | HLA-DRB5 | Exon 2 | 6 | 32489933 | T | A | Father | 24.031416 | 0.006789 | missense | Neutral |
| 411 | Male | OR2T8 | Exon 1 | 1 | 248085124 | G | A | Father | 23.806671 | 0.011182 | missense | Deleterious |
| 411 | Male | OR2T8 | Exon 1 | 1 | 248084470 | C | CG | Mother | 23.806671 | 0.011182 | frameshift_variant | (predicted null) |
| 411 | Male | HLA-A | Exon 3 | 6 | 29911115 | G | GAA | Mother | 22.901964 | 0.020767 | frameshift_variant | |
| 411 | Male | AC008686.1 | | 19 | 13899041 | CT | C | Mother | 22.889645 | 0.042332 | frameshift_variant | (predicted null) |
| 411 | Male | GOLGA8S | Exon 18 | 15 | 23610226 | A | G | Father | 21.057671 | 0.046725 | missense | Neutral |
| 411 | Male | GOLGA8S | Exon 13 | 15 | 23608318 | C | A | Mother | 21.057671 | 0.046725 | missense | Neutral |
| 716 | Female | C11orf40 | Exon 1 | 11 | 4599017 | T | C | Father | 37.952408 | 0.000399 | missense | Deleterious |
| 716 | Female | C11orf40 | Exon 4 | 11 | 4592706 | T | TAC | Mother | 37.952408 | 0.000399 | frameshift_variant | |
| 716 | Female | CCDC136 | Exon 13 | 7 | 128452280 | G | A | Mother | 37.817055 | 0.000399 | missense | Neutral |
| 716 | Female | **CCDC136** | Exon 10 | 7 | 128447502 | GCAGGACC | A | Father | 37.817055 | 0.000399 | inframe_deletion | Deleterious |
| 716 | Female | CCDC66 | Exon 13 | 3 | 56650051 | A | ACTT | Both | 36.696259 | 0.000399 | inframe_insertion | Neutral |
| 716 | Female | MUC12 | Exon 2 | 7 | 100635591 | C | A | Both | 35.780857 | 0.000399 | missense | Neutral |
| 716 | Female | **MYO1H** | Exon 3 | 12 | 109834353 | GT | G | Father | 30.642544 | 0.000799 | frameshift_variant | (predicted null) |

**TABLE S4**

| CDH Proband | ExAC Pli 1= intolerant 0= tolerant of haploinsufficency | gnomAD Frequency | TPM RNA-Seq PPFs | Expressed in PPFs? | Highly mutable gene family? | FLAGS gene? | Multi-allelic in gnomAD? | GENCODE annotation of no-coding genes https://www.gencodegenes.org/ | Why included or excluded as gene potentially contributing to CDH |
|---|---|---|---|---|---|---|---|---|---|
| 411 | NA | 0.2248 | 0 | No | | | Yes | Pseudogene | Excluded: multi-allelic and pseudogene, no PPF expression |
| 411 | NA | Not found in gnomad | 0 | No | | | Yes | Pseudogene | |
| 411 | 0.62 | 0.01272 | 44.53403597 | Yes | Mutable locus according to MY Yandell lab, unpub | | Yes | | Excluded: multi-allelic and mutable locus |
| 411 | 0.62 | 0.004992 | 44.53403597 | Yes | Mutable locus according to MY Yandell lab, unpub | | Yes | | |
| 411 | 0.75 | 0.0002355 | 0.536857763 | Minimal | | | | | Excluded: minimal PPF expression |
| 411 | NA | 0.1893 | 0 | No | | | | Pseudogene | Excluded: no PPF expression |
| 411 | NA | 0.1047 | 0 | No | | | Yes | Pseudogene | |
| 411 | 0 | 0.04477 | No mouse ortholog | NA | Highly mutable, Lenz et al. 2016 Mol. Biol. Evol. | | Yes | | Excluded: multiallelic and mutable locus |
| 411 | 0 | 0.00128 | No mouse ortholog | NA | Highly mutable, Lenz et al. 2016 Mol. Biol. Evol. | | | | |
| 411 | 0 | 0.7575 | 0 | No | | | Yes | | Excluded: multi-allelic, no PPF expression |
| 411 | 0 | 0.00008506 | 75.82683746 | Yes | | | | | **Included: 1 rare missense predicted deletereious by Provean allele, expressed in PPFs BUT other allele rare predicted neutral missense allele. 4th rank** |
| 411 | 0 | 0.00004338 | 75.82683746 | Yes | | | | | |
| 411 | NA | 0.08975 | 0 | No | | | Yes | Pseudogene | Excluded: intron variant minimal PPF expression |
| 411 | NA | 0.5216 | 0 | No | | | Yes | Pseudogene | |
| 411 | 0 | 0.003851 | No mouse ortholog | NA | Highly mutable, Lenz et al. 2016 Mol. Biol. Evol. | | Yes | | Excluded: highly mutable and multi-allelic |
| 411 | 0 | 0.02 | No mouse ortholog | NA | Highly mutable, Lenz et al. 2016 Mol. Biol. Evol. | | Yes | | |
| 411 | 0 | 0.0005634 | 0 | No | | Yes | | | Excluded: FLAGS gene, multi-allelic, and no PPF expression |
| 411 | 0 | 0.2298 | 0 | No | | Yes | Yes | | |
| 411 | 0 | 0.01428 | No mouse ortholog | NA | Highly mutable, Lenz et al. 2016 Mol. Biol. Evol. | | Yes | | Excluded: highly mutable and multi-allelic |
| 411 | 0 | 0.03155 | No mouse ortholog | NA | Highly mutable, Lenz et al. 2016 Mol. Biol. Evol. | | Yes | | |
| 411 | 0.72 | Not found in gnomad | 0 | No | | | | | Excluded: multi-allelic and no PPF expression |
| 411 | 0.72 | 0.0008493 | 0 | No | | | Yes | | |
| 411 | 0.13 | 0.02532 | No mouse ortholog | NA | Highly mutable, Lenz et al. 2016 Mol. Biol. Evol. | | Yes | | Excluded: highly mutable and multi-allelic |
| 411 | NA | Not found in gnomad | 0 | No | | | | LncRNA | Excluded: no PPF expression |
| 411 | 0 | 0.003815 | 61.33785 | Yes | | | | Noncoding RNA | Excluded: both alleles predived neutral by Provean and 1 allele > .01 gnomAD frequency |
| 411 | 0 | 0.03139 | 61.33785 | Yes | | | | Noncoding RNA | |
| 716 | 0 | 0.00002368 | No mouse ortholog | NA | | | | | Excluded: multi-allelic, 1 allele >> 0.01 gnomAD frequency |
| 716 | 0 | 0.2999 | No mouse ortholog | NA | | | Yes | | |
| 716 | 0 | 0.00006029 | 25.94114519 | Yes | | | | | **Included: 1 rare in frame deletion and predicted deleterious by Provean allele, expressed in PPFs BUT other allele rare neutral missense allele. 4th rank** |
| 716 | 0 | 0.00001378 | 25.94114519 | Yes | | | | | |
| 716 | 0 | 0.4272 | 31.78998157 | Yes | | | | | Excluded: allele >> 0.01gnomAD frequency |
| 716 | NA | 0.4973 | No mouse ortholog | NA | Mutable locus according to MY Yandell lab, unpub | | Yes | | Excluded: highly mutatable, multi-allelic |
| 716 | 0 | 0.00008452 | 36.39642641 | Yes | | | Yes | | **Included: 1 rare frameshift (null) allele, 1** |

### TABLE S4

| CDH Proband | Sex | Gene Name | Location in Gene | Chromosome | Position | Reference Allele | Alternative Allele | Parent found in | VAAST Score | VAAST p value | VAAST Impact | PROVEAN Functional Prediction |
|---|---|---|---|---|---|---|---|---|---|---|---|---|
| 716 | Female | **MYO1H** | Exon 23 | 12 | 109877506 | C | T | Mother | 30.642544 | 0.000799 | missense | Deleterious |
| 716 | Female | CTAGE8 | Exon 1 | 7 | 143964381 | T | C | Mother | 30.100883 | 0.000399 | missense | Neutral |
| 716 | Female | CTAGE8 | Exon 1 | 7 | 143964126 | T | C | Father | 30.100883 | 0.000399 | missense | Deleterious |
| 716 | Female | NOP9 | Exon 2 | 14 | 24769849 | A | AGAGGAG | Father | 29.468845 | 0.001597 | inframe_insertion | Neutral |
| 716 | Female | **NOP9** | Exon 8 | 14 | 24773448 | C | T | Mother | 29.468845 | 0.001597 | missense | Deleterious |
| 716 | Female | **MPEG1** | Exon 1 | 11 | 58980189 | G | GCC | Mother | 29.247192 | 0.001597 | frameshift_variant | (predicted null) |
| 716 | Female | MPEG1 | Exon 1 | 11 | 58979643 | A | C | Father | 29.247192 | 0.001597 | missense | Neutral |
| 716 | Female | ZNF646 | Exon 2 | 16 | 31088738 | G | A | Mother | 29.067986 | 0.000399 | missense | Neutral |
| 716 | Female | **ZNF646** | Exon 2 | 16 | 31090928 | C | T | Father | 29.067986 | 0.000399 | missense | Deleterious |
| 716 | Female | **HRC** | Exon 1 | 19 | 49657569 | T | A | Mother | 28.644558 | 0.00639 | missense | Deleterious |
| 716 | Female | **HRC** | Exon 1 | 19 | 49657221 | T | C | Father | 28.644558 | 0.00639 | missense | Deleterious |
| 716 | Female | HLA-B | Exon 2 | 6 | 31324528 | G | GGA | Mother | 26.501488 | 0.004393 | frameshift_variant | NA |
| 716 | Female | HLA-B | Exon 3 | 6 | 31324025 | G | A | Father | 26.501488 | 0.004393 | missense | Neutral |
| 716 | Female | **AHNAK** | Exon 5 | 11 | 62292741 | C | G | Mother | 26.463722 | 0.038738 | missense | Deleterious |
| 716 | Female | **AHNAK** | Exon 5 | 11 | 62293913 | T | G | Mother | 26.463722 | 0.038738 | missense | Deleterious |
| 716 | Female | **ALG2** | Exon 1 | 9 | 101983960 | C | G | Father | 26.082867 | 0.005192 | missense | Deleterious |
| 716 | Female | **ALG2** | Exon 1 | 9 | 101984010 | G | A | Mother | 26.082867 | 0.005192 | missense | Deleterious |
| 716 | Female | FRG1BP | Exon 8 | 20 | 29632673 | T | C | Father | 24.940754 | 0.045527 | missense | Deleterious |
| 716 | Female | FRG1BP | Exon 8 | 20 | 29632662 | G | C | Mother | 24.940754 | 0.045527 | missense | Neutral |
| 716 | Female | PLD2 | Exon 10 | 17 | 4714225 | G | A | Father | 24.710567 | 0.042732 | missense | Neutral |
| 716 | Female | PLD2 | Exon 25 | 17 | 4726052 | G | C | Mother | 24.710567 | 0.042732 | missense | Neutral |
| 716 | Female | PRAMEF1 | Exon 4 | 1 | 12855895 | A | G | Mother | 22.769529 | 0.045128 | missense | Neutral |
| 716 | Female | PRAMEF1 | Exon 4 | 1 | 12856105 | C | G | Father | 22.769529 | 0.045128 | stop_gained | (predicted null) |
| 716 | Female | CHP2 | Exon 5 | 16 | 23767738 | G | T | Both | 22.554684 | 0.033147 | missense | Deleterious |
| 809 | Female | HLA-A | Exon 2 | 6 | 29910779 | G | GCTCC | Both | 41.939743 | 0.000399 | frameshift_variant | (predicted null) |
| 809 | Female | AL138799.4 | | 1 | 81979731 | TCCTTTGG[illegible] | C | Both | 40.185852 | 0.000399 | frameshift_variant | |
| 809 | Female | FOXD4L5 | Exon 1 | 9 | 70176851 | A | G | Father | 39.515709 | 0.000399 | missense | Neutral |
| 809 | Female | FOXD4L5 | Exon 1 | 9 | 70176768 | AG | A | Mother | 39.515709 | 0.000399 | frameshift_variant | |
| 809 | Female | NBPF9 | Exon 88 | 1 | 145349714 | G | A | Mother | 39.404705 | 0.000399 | missense | Neutral |

### TABLE S4

| CDH Proband | ExAC Pli 1= intolerant 0= tolerant of haploinsufficency | gnomAD Frequency | TPM RNA-Seq PPFs | Expressed in PPFs? | Highly mutable gene family? | FLAGS gene? | Multi-allelic in gnomAD? | GENCODE annotation of no-coding genes https://www.gencodegenes.org/ | Why included or excluded as gene potentially contributing to CDH |
|---|---|---|---|---|---|---|---|---|---|
| 716 | 0 | 0.006348 | 36.39642641 | Yes | | | Yes | | **rare missense and predicted deleterious by Provean allele, expressed in PPFs, BUT alleles in multi-allelic gnomAD region. 2nd rank** |
| 716 | NA | 0.1207 | 0.12 | Minimal | | | | | Excluded: both variants > 0.01 in gnomAD, one predicted neutral by Provean, and minimal PPD expression |
| 716 | NA | 0.01583 | 0.12 | Minimal | | | | | |
| 716 | 0 | 0.3478 | 39.05250943 | Yes | | | Yes | | **Included: 1 rare missense and predicted deleterious by Provean allele, expressed in PPFs BUT other allele common predicted neutral in frame insertion allele. 4th rank** |
| 716 | 0 | 0.001724 | 39.05250943 | Yes | | | Yes | | |
| 716 | 0 | Not found in gnomad | 21.31529659 | Yes | | | | | **Included: 1 frameshift (null) rare allele, expressed in PPFs BUT other allele rare predicted neutral by Provean missense allele. 3rd rank** |
| 716 | 0 | 0.00293 | 21.31529659 | Yes | | | | | |
| 716 | 0.01 | 0.0003458 | 39.47584 | Yes | | | | | **Included: 1 rare missense and predicted deleterious by Provean allele, expressed in PPFs BUT other allele rare predicted neutral missense allele. 4th rank** |
| 716 | 0.01 | 0.0002126 | 39.47584 | Yes | | | Yes (1 other predicted damaging allele) | | |
| 716 | 0 | 0.000439 | 35.7001661 | Yes | | | | | **Included: 2 rare, missense, and predicted deleterious by Provean alleles, expressed in PPFs. 1st rank** |
| 716 | 0 | 0.0000875 | 35.7001661 | Yes | | | | | |
| 716 | 0 | 0.03249 | No mouse ortholog | NA | Highly mutable, Lenz et al. 2016 Mol. Biol. Evol. | | Yes | | Excluded: highly mutable and multi-allelic |
| 716 | 0 | 0.1785 | No mouse ortholog | NA | Highly mutable, Lenz et al. 2016 Mol. Biol. Evol. | | Yes | | |
| 716 | 0.34 | 0.000003978 | 45.74418807 | Yes | | Yes | | | **Included: 2 rare missense and predicted deleterious by Provean alleles, expressed in PPFs BUT gene FLAGS gene as mutable. 2nd rank.** |
| 716 | 0.34 | 0.001128 | 45.74418807 | Yes | | Yes | | | |
| 716 | 0.02 | 0.0001388 | 90.3247514 | Yes | | | | | **Included: 2 rare, missense, and predicted deleterious by Provean alleles, expressed in PPFs. 1st rank** |
| 716 | 0.02 | 0.0008662 | 90.3247514 | Yes | | | | | |
| 716 | 0 | 0.0000589 | 0 | No | | | | Pseudogene | Excluded: pseudogene, multi-allelic, and not expressed in PPFs |
| 716 | 0 | 0.04104 | 0 | No | | | Yes | Pseudogene | |
| 716 | 0 | 0.003188 | 12.36657083 | Yes | | | | | Excluded: 2 missense but neutral variants |
| 716 | 0 | Not found in gnomad | 12.36657083 | Yes | | | | | |
| 716 | 0.1 | 0.002484 | 0 | No | | | | | Predicted null allele expressed at > 0.01 frequency in gnomAD, and not expressed in PPFs |
| 716 | 0.1 | 0.01956 | 0 | No | | | | | |
| 716 | 0.01 | 0.002696 | 0.367621499 | Minimal | | | Yes | | Excluded: minimally expressed in PPFs |
| 809 | 0.13 | 0.1635 | No mouse ortholog | NA | Highly mutable, Lenz et al. 2016 Mol. Biol. Evol. | | Yes | | Excluded: highly mutable and multi-allelic |
| 809 | NA | 0.07625 | No mouse ortholog | NA | | | | lncRNA | Excluded: allele expressed at > 0.01 frequency in gnomAD |
| 809 | NA | 0.2413 | 0.05 | Minimal | | | | | Excluded: allele expressed at >> 0.01 frequency in gnomAD and minimally expressed in PPFs |
| 809 | NA | 0.3831 | 0.05 | Minimal | | | | | |
| 809 | NA | 0.1188 | 0 | No | | | | | Excluded: rare premature stop allele, but |

### TABLE S4

| CDH Proband | Sex | Gene Name | Location in Gene | Chromosome | Position | Reference Allele | Alternative Allele | Parent found in | VAAST Score | VAAST p value | VAAST Impact | PROVEAN Functional Prediction |
|---|---|---|---|---|---|---|---|---|---|---|---|---|
| 809 | Female | NBPF9 | Intron 21 | 1 | 145311135 | C | T | Father | 39.404705 | 0.000399 | stop_gained | |
| 809 | Female | CNTNAP3B | Exon 22 | 9 | 43915558 | G | C | Father | 35.723511 | 0.000399 | missense | Neutral |
| 809 | Female | CNTNAP3B | Exon 22 | 9 | 43915557 | C | T | Mother | 35.723511 | 0.000399 | missense | Neutral |
| 809 | Female | POM121L7P | Exon 1 | 22 | 21481150 | C | T | Father | 33.905251 | 0.000399 | missense | Neutral |
| 809 | Female | POM121L7P | Exon 1 | 22 | 21481208 | G | A | Mother | 33.905251 | 0.000399 | missense | Neutral |
| 809 | Female | PABPC3 | Exon 1 | 13 | 25670801 | T | TG | Mother | 30.636066 | 0.000399 | frameshift_variant | (predicted null) |
| 809 | Female | PABPC3 | Exon 1 | 13 | 25671941 | A | C | Father | 30.636066 | 0.000399 | missense | Deleterious |
| 809 | Female | TEX15 | Exon 3 | 8 | 30694432 | G | A | Mother | 27.399559 | 0.005192 | missense | Neutral |
| 809 | Female | TEX15 | Exon 3 | 8 | 30694423 | C | T | Father | 27.399559 | 0.005192 | missense | Neutral |
| 809 | Female | FTMT | Exon 1 | 5 | 121187773 | G | T | Mother | 25.929003 | 0.00599 | missense | Neutral |
| 809 | Female | FTMT | Exon 1 | 5 | 121188325 | G | T | Father | 25.929003 | 0.00599 | missense | Neutral |
| 809 | Female | LINC01551 | Exon 3 | 14 | 29261309 | A | AAAC | Mother | 24.375664 | 0.033147 | inframe_insertion | Neutral |
| 809 | Female | LINC01551 | Exon 3 | 14 | 29261309 | A | AAC | Father | 24.375664 | 0.033147 | frameshift_variant | (predicted null) |
| 967 | Male | LNP1 | Exon 3 | 3 | 100170600 | A | CCAGGAATT | Both | 41.939743 | 0.000399 | inframe_insertion | Deleterious |
| 967 | Male | ADAMTS2 | Exon 1 | 5 | 178772241 | G | GCGGCAGG | Both | 40.769886 | 0.000399 | inframe_insertion | Neutral |
| 967 | Male | MYOF | Exon 18 | 10 | 95148899 | T | A | Father | 40.012344 | 0.000399 | missense | Neutral |
| 967 | Male | **MYOF** | Exon 7 | 10 | 95168549 | C | CA | Mother | 40.012344 | 0.000399 | frameshift_variant | (predicted null) |
| 967 | Male | POM121L7P | Exon 1 | 22 | 21481354 | G | C | Father | 39.526966 | 0.000399 | missense | Neutral |
| 967 | Male | POM121L7P | Exon 1 | 22 | 21481518 | CT | C | Mother | 39.526966 | 0.000399 | frameshift_variant | (predicted null) |
| 967 | Male | DUX4L4 | Exon 1 | 4 | 191003195 | C | T | Father | 34.866108 | 0.000399 | missense | Deleterious |
| 967 | Male | DUX4L4 | Exon 1 | 4 | 191003212 | C | T | Mother | 34.866108 | 0.000399 | missense | Neutral |
| 967 | Male | SACS | Exon 10 | 13 | 23910966 | T | C | Mother | 34.806686 | 0.000399 | missense | Neutral |
| 967 | Male | **SACS** | Exon 10 | 13 | 23904275 | TCA | T | Father | 34.806686 | 0.000399 | frameshift_variant&stop_lost | (predicted null) |
| 967 | Male | CLEC18B | Exon 3 | 16 | 74452114 | A | G | Mother | 34.245117 | 0.000399 | upstream_gene_variant | Neutral |
| 967 | Male | CLEC18B | Exon 9 | 16 | 74444863 | G | A | Father | 34.245117 | 0.000399 | missense | Neutral |
| 967 | Male | NBPF10 | Intron 68 | 1 | 145368630 | G | T | Father | 32.5159 | 0.001597 | missense | Neutral |
| 967 | Male | NBPF10 | Exon 4 | 1 | 145297661 | C | A | Mother | 32.5159 | 0.001597 | missense | Neutral |
| 967 | Male | PARP4 | Exon 33 | 13 | 25000617 | C | T | Mother | 30.722176 | 0.001997 | missense | Neutral |
| 967 | Male | PARP4 | Exon 31 | 13 | 25009117 | G | T | Father | 30.722176 | 0.001997 | missense | Neutral |
| 967 | Male | **SH3PXD2A** | Exon 14 | 10 | 105361796 | G | T | Father | 30.664799 | 0.000399 | missense | Deleterious |
| 967 | Male | SH3PXD2A | Exon 14 | 10 | 105362735 | G | A | Mother | 30.664799 | 0.000399 | missense | Neutral |
| 967 | Male | GRIN3B | Exon 1 | 19 | 1000448 | G | CGCTGTGGC | Father | 29.922745 | 0.001597 | inframe_insertion | Neutral |
| 967 | Male | GRIN3B | Exon 3 | 19 | 1004896 | G | GCGTT | Mother | 29.922745 | 0.001597 | frameshift_variant | |
| 967 | Male | HLA-A | Exon 2 | 6 | 29910779 | G | GCTCC | Father | 29.53603 | 0.000399 | frameshift_variant | (predicted null) |
| 967 | Male | HLA-A | Exon 3 | 6 | 29911240 | T | A | Father | 29.53603 | 0.000399 | Nonsense | |

### TABLE S4

| CDH Proband | ExAC Pli 1= intolerant 0= tolerant of haploinsufficency | gnomAD Frequency | TPM RNA-Seq PPFs | Expressed in PPFs? | Highly mutable gene family? | FLAGS gene? | Multi-allelic in gnomAD? | GENCODE annotation of no-coding genes https://www.gencodegenes.org/ | Why included or excluded as gene potentially contributing to CDH |
|---|---|---|---|---|---|---|---|---|---|
| 809 | NA | 0.00006187 | 0 | No | | | Yes | | other allele missense and predicted neutral by Provean, minimally expressed in PPFs |
| 809 | NA | 0.1221 | 0 | No | | | | | Excluded: common missense alleles predicted by Provean and not expressed in PPFs |
| 809 | NA | 0.2926 | 0 | No | | | Yes | | |
| 809 | NA | 0.001166 | 0 | No | | | | Pseudogene | Excluded: pseudogene |
| 809 | NA | 0.06326 | 0 | No | | | | Pseudogene | |
| 809 | 0 | 0.2585 | 0.03673923 | Minimal | | | Yes | | Excluded: one null, but common allele, other allele deleterious, minimally expressed in PPFs |
| 809 | 0 | 0.002715 | 0.03673923 | Minimal | | | | | |
| 809 | 0 | 0.00522 | 2.69477027 | Minimal | | | | | Excluded: missense, but predicted neutral by Provean alleles, minimally expressed in PPFs |
| 809 | 0 | 0.00001194 | 2.69477027 | Minimal | | | | | |
| 809 | 0.08 | Not found in gnomad | 0 | No | | | | | Excluded: missense, but predicted neutral by Provean alleles, not expressed in PPFs |
| 809 | 0.08 | 0.004263 | 0 | No | | | Yes | | |
| 809 | NA | 0.4238 | No mouse ortholog | NA | | | Yes | lncRNA | Excluded: both alleles > 0.01 frequency in gnomAD |
| 809 | NA | 0.09915 | No mouse ortholog | NA | | | Yes | lncRNA | |
| 967 | 0 | Not found in gnomad | 0 | No | | | | | Excluded: deleterious alleles, but not expressed in PPFs |
| 967 | **0.99** | 0.01099 | 38.9203826 | Yes | | | | | **Included: 2 rare in-frame insertion alleles with high Pli score BUT insertion predicted by Provean to be neutral. 4th rank** |
| 967 | 0 | 0.0004856 | 28.96335117 | Yes | | | Yes | | **Included: 1 frameshift (null) rare allele, expressed in PPFs BUT other allele rare predicted neutral by Provean missense allele. 3rd rank** |
| 967 | 0 | 0.00003568 | 28.96335117 | Yes | | | | | |
| 967 | NA | 0.004121 | No mouse ortholog | NA | | | | Pseudogene | Excluded: pseudogene |
| 967 | NA | 0.001337 | No mouse ortholog | NA | | | | Pseudogene | |
| 967 | NA | 0.003953 | 0 | No | | | Yes | Pseudogene | Excluded: pseudogene and multi-allelic |
| 967 | NA | 0.06725 | 0 | No | | | Yes | Pseudogene | |
| 967 | 0 | 0.00008853 | 10.85410849 | Yes | | | | | **Included: 1 frameshift (null) rare allele, expressed in PPFs BUT other allele rare predicted neutral by Provean missense allele. 3rd rank** |
| 967 | 0 | 0.00002795 | 10.85410849 | Yes | | Yes | Yes | | |
| 967 | NA | 0.003991 | 0.03444378 | Minimal | | | | | Excluded: both alleles predicted neutral by Provean and minimally expressed in PPFs |
| 967 | NA | 0.008186 | 0.03444378 | Minimal | | | | | |
| 967 | 0 | 0.0004153 | 0 | No | | | Yes | | Excluded: both alleles predicted neutral by Provean and not expressed in PPFs |
| 967 | 0 | 0.005416 | 0 | No | | | Yes | | |
| 967 | 0 | 0.0001703 | 4.725922336 | Minimal | | | Yes | | Excluded: both alleles predicted neutral by Provean and minimally expressed in PPFs |
| 967 | 0 | Not found in gnomad | 4.725922336 | Minimal | | | | | |
| 967 | 0.23 | 0.00001075 | 136.7142617 | Yes | | | | | **Included: 1 rare missense and predicted by Provean deleterious allele, expressed in PPFs BUT other allele rare predicted neutral missense allele. 4th** |
| 967 | 0.23 | 0.00006717 | 136.7142617 | Yes | | | | | |
| 967 | 0 | 0.0003975 | 3.039457967 | Minimal | | | | | Excluded: predicted neutral by Provean or common allele, minimally expressed in |
| 967 | 0 | 0.2481 | 3.039457967 | Minimal | | | Yes | | |
| 967 | 0.13 | 0.1635 | 0 | No | Highly mutable, Lenz et al. 2016 Mol. Biol. Evol. | | Yes | | Excluded: highly mutable and multi-allelic |
| 967 | 0.13 | 0.2131 | 0 | No | Highly mutable, Lenz et al. 2016 Mol. Biol. Evol. | | Yes | | |

**TABLE S4**

| CDH Proband | Sex | Gene Name | Location in Gene | Chromosome | Position | Reference Allele | Alternative Allele | Parent found in | VAAST Score | VAAST p value | VAAST Impact | PROVEAN Functional Prediction |
|---|---|---|---|---|---|---|---|---|---|---|---|---|
| 967 | Male | C4orf17 | Exon 5 | 4 | 100451040 | GGATGACA | A | Father | 28.450815 | 0.001198 | frameshift_variant | |
| 967 | Male | C4orf17 | Exon 7 | 4 | 100460420 | T | TG | Mother | 28.450815 | 0.001198 | frameshift_variant | |
| 967 | Male | ECSIT | Exon 5 | 19 | 11618827 | G | T | Mother | 27.276222 | 0.003594 | missense | Neutral |
| 967 | Male | ECSIT | Exon 8 | 19 | 11617052 | C | T | Father | 27.276222 | 0.003594 | missense | Neutral |
| 967 | Male | MUC17 | Exon 3 | 7 | 100680243 | C | T | Mother | 26.722765 | 0.043131 | missense | Neutral |
| 967 | Male | MUC17 | Exon 3 | 7 | 100678821 | G | A | Father | 26.722765 | 0.043131 | missense | Neutral |
| 967 | Male | HLA-B | Exon 2 | 6 | 31324601 | C | CG | Both | 26.501488 | 0.004393 | frameshift_variant | (predicted null) |
| 967 | Male | DACT2 | Exon 4 | 6 | 168709386 | C | T | Mother | 23.919924 | 0.038738 | missense | Neutral |
| 967 | Male | DACT2 | Exon 4 | 6 | 168709234 | G | CCCTGCCGG | Father | 23.919924 | 0.038738 | inframe_insertion | Deleterious |
| 967 | Male | ZNF594 | Exon 2 | 17 | 5085480 | C | T | Mother | 23.159637 | 0.041933 | missense | Neutral |
| 967 | Male | ZNF594 | Exon 2 | 17 | 5085877 | GCT | G | Father | 23.159637 | 0.041933 | frameshift_variant | (predicted null) |
| 967 | Male | C22orf42 | Exon 5 | 22 | 32547523 | A | T | Both | 23.149113 | 0.034345 | missense | Neutral |
| 967 | Male | ZFP36L2 | Exon 2 | 2 | 43452665 | C | T | Mother | 22.470539 | 0.037141 | missense | Neutral |
| 967 | Male | ZFP36L2 | Exon 2 | 2 | 43452608 | A | T | Father | 22.470539 | 0.037141 | missense | Neutral |
| 967 | Male | CR936218.1 | | 17 | 44112733 | G | A | Both | 20.695335 | 0.033546 | missense | |

**TABLE S4**

| CDH Proband | ExAC Pli 1= intolerant 0= tolerant of haploinsufficency | gnomAD Frequency | TPM RNA-Seq PPFs | Expressed in PPFs? | Highly mutable gene family? | FLAGS gene? | Multi-allelic in gnomAD? | GENCODE annotation of no-coding genes https://www.gencodegenes.org/ | Why included or excluded as gene potentially contributing to CDH |
|---|---|---|---|---|---|---|---|---|---|
| 967 | 0 | 0.00482 | No mouse ortholog | NA | | | | | Excluded: no infomration to include as likely candidate gene |
| 967 | 0 | 0.0006901 | No mouse ortholog | NA | | | | | |
| 967 | 0.02 | 0.0003429 | 41.57275224 | Yes | | | Yes | | Excluded: 2 missense but neutral variants |
| 967 | 0.02 | 0.000004082 | 41.57275224 | Yes | | | | | |
| 967 | NA | 0.0046 | No mouse ortholog | NA | Mutable locus according to MY Yandell lab, unpub | Yes | | | Excluded: highly mutable, FLAGS gene, multi-allelic |
| 967 | NA | 0.002613 | No mouse ortholog | NA | Mutable locus according to MY Yandell lab, unpub | Yes | Yes | | |
| 967 | 0 | 0.1152 | No mouse ortholog | NA | Highly mutable, Lenz et al. 2016 Mol. Biol. Evol. | | Yes | | Excluded: highly mutable and multi-allelic |
| 967 | NA | 0.0000485 | 12.80168554 | Yes | | | Yes | | Excluded: only 1 neutral missense allele and 1 deleterious by > 0.01 gnomAD frequency |
| 967 | NA | 0.03964 | 12.80168554 | Yes | | | Yes | | |
| 967 | 0 | 0.0003373 | 0 | No | | | | | Excluded: predicted null allele and neutral missense allele by Provean, but no expression in PPFs |
| 967 | 0 | 0.004689 | 0 | No | | | Yes | | |
| 967 | 0 | 0.002148 | No mouse ortholog | NA | | | | | Excluded: missense allele predicted neutral by Provean |
| 967 | 0.46 | 0.00009252 | 36.52641388 | Yes | | | Yes | | Excluded: 2 missense but predicted neutral by Provean |
| 967 | 0.46 | 0.001987 | 36.52641388 | Yes | | | Yes | | |
| 967 | NA | 0.02682 | 0 | No | | | | lncRNA | Excluded: allele expressed at > 0.01 frequency in gnomAD |

### TABLE S5

| CDH Proband | Chromosome | Start | End | Length of Variant | Genotype: 0 = reference, 1 = alternate SV | Reference split reads | Alternative split reads | Reference discordant pairs | Alternative discordant pairs | Length and translocated location | Mutation type | Gene location | Repeats overlapping SV | Allele frequency in Abel *et al.* 2018 |
|---|---|---|---|---|---|---|---|---|---|---|---|---|---|---|
| 411 | 8 | 11596931 | 11597274 | 343 | 0/1 | 41 | 8 | 30 | 7 | <DEL> | Inherited from father | **Gata4 intron 2** | | 0.01946 |
| 411 | 8 | 19055260 | 19055530 | 270 | 0/1 | 55 | 24 | 41 | 7 | <DEL> | Inherited from father | Loc100128993 intron 3 | | 0.08774 |
| 411 | 8 | 117241911 | 117243103 | 1192 | 0/1 | 51 | 18 | 27 | 12 | <DEL> | Inherited from father | Linc00536 intron 4 | | Not found |
| 716 | 5 | 15719465 | 15720950 | 1485 | 0/1 | 43 | 21 | 34 | 22 | <DEL> | Inherited from father | Fbxl7 intron 2 | MiRb (smaller than SV) | 0.09933 |
| 716 | 12 | 2952933 | 2953348 | 415 | 0/1 | 54 | 10 | 36 | 20 | <DEL> | Inherited from father | Loc100507424 intron 2 | | 0.093 |
| 809 | 4 | 141633311 | 141633603 | 292 | 0/1 | 49 | 26 | 58 | 9 | <DEL> | Inherited from father | TBC1D9 intron 1 | | Not found |
| 967 | 6 | 167543015 | 167544428 | 1413 | 0/1 | 40 | 4 | 30 | 16 | <DEL> | Inherited from father | Ccr6 intron 1 | L3 (smaller than SV) | 0.01874 |
| 967 | 8 | 11596931 | 11597274 | 343 | 0/1 | 22 | 9 | 31 | 18 | <DEL> | Inherited from mother | **Gata4 intron 2** | | 0.01946 |
| 967 | 8 | 103053960 | 103056716 | 2756 | 0/1 | 54 | 19 | 39 | 19 | <DEL> | Inherited from mother | Ncald intron 1 | LTR78 (smaller than SV) | 0.01057 |

### TABLE S5

| CDH Proband | Chromosome | Start | End | Length of Variant | Genotype: 0 = reference, 1 = alternate SV | Reference split reads | Alternative split reads | Reference discordant pairs | Alternative discordant pairs | Length and translocated location | Mutation type | Gene location | Repeats overlapping SV | Allele frequency in Abel *et al.* 2018 | Filtering reason |
|---|---|---|---|---|---|---|---|---|---|---|---|---|---|---|---|
| 411 | 1 | 180749772 | 180755393 | 5621 | 0/1 | 52 | 19 | 29 | 23 | <DEL> | Inherited from father | XPR1 intron 2 | Multiple | 0.006326 | Overlaps repetitive regions |
| 411 | 1 | 229236567 | 229243367 | 6800 | 0/1 | 98 | 24 | 57 | 28 | <DUP> | Inherited from father | Intergenic | Multiple | 0.006018 | Intergenic |
| 411 | 2 | 233292854 | 233296570 | 3716 | 0/1 | 44 | 19 | 28 | 14 | <DEL> | Inherited from father | Intergenic | Multiple | 0.007591 | Intergenic |
| 411 | 3 | 133137976 | 133140341 | 2365 | 0/1 | 107 | 17 | 59 | 14 | <DUP> | Inherited from mom | Bfsp2 intron 1 | Multiple | 3.42E-05 | Overlaps repetitive regions |
| 411 | 4 | 147953942 | 147953942 | 0 | 0/1 | 70 | 11 | 44 | 16 | N]20:54914679] | Inherited from mom | Intergenic | | 0.09037 | Intergenic |
| 411 | 4 | 152778524 | 152778992 | 468 | 0/1 | 50 | 14 | 30 | 10 | <DEL> | Inherited from mom | Intergenic | LTR78 | 0.07529 | Intergenic |
| 411 | 6 | 773629 | 774802 | 1173 | 0/1 | 54 | 16 | 30 | 14 | <DEL> | Hom in father, het in proband | Intergenic | Multiple | 0.09365 | Parent homozygous for SV |
| 411 | 8 | 11596931 | 11597274 | 343 | 0/1 | 41 | 8 | 30 | 7 | <DEL> | Inherited from father | **Gata4 intron 2** | | 0.01946 | |
| 411 | 8 | 19055260 | 19055530 | 270 | 0/1 | 55 | 24 | 41 | 7 | <DEL> | Inherited from father | Loc100128993 intron 3 | | 0.08774 | |
| 411 | 8 | 117241911 | 117243103 | 1192 | 0/1 | 51 | 18 | 27 | 12 | <DEL> | Inherited from father | Linc00536 intron 4 | | Not found | |
| 411 | 12 | 10407411 | 10407721 | 310 | 0/1 | 53 | 13 | 35 | 13 | <DEL> | Inherited from mother | Intergenic | AluSx, L1PREC2 | 0.06131 | Intergenic |
| 411 | 12 | 13725909 | 13726553 | 644 | 0/1 | 39 | 20 | 22 | 23 | <DEL> | Inherited from mother | GRIN2B intron 9 | MiRb, AluY | 0.09225 | Overlaps repetitive regions |
| 411 | 12 | 21861577 | 21861929 | 352 | 0/1 | 40 | 17 | 24 | 18 | <DEL> | Inherited from father | Intergenic | MiR | 0.05799 | Intergenic |
| 716 | 1 | 184278739 | 184279041 | 302 | 0/1 | 33 | 19 | 39 | 18 | <DEL> | Inherited from father | Intergenic | AluYa5 | 0.08254 | Intergenic |
| 716 | 1 | 195013812 | 195019758 | 5946 | 0/1 | 40 | 25 | 28 | 27 | <DEL> | Inherited from mother | Intergenic | Multiple | 0.004035 | Intergenic |
| 716 | 4 | 152790165 | 152794721 | 4556 | 0/1 | 55 | 16 | 37 | 22 | <DEL> | Mother hom, proband het | Intergenic | Multiple | 0.06162 | Parent homozygous for SV |
| 716 | 4 | 152891448 | 152893680 | 2232 | 0/1 | 54 | 16 | 38 | 13 | <DEL> | Inherited from father | Intergenic | Multiple | 0.004719 | Intergenic |
| 716 | 4 | 154263656 | 154264646 | 990 | 0/1 | 37 | 16 | 32 | 19 | <DEL> | Inherited from father | Intergenic | AluSq, MER83 | Not found | Intergenic |
| 716 | 5 | 15719465 | 15720950 | 1485 | 0/1 | 43 | 21 | 34 | 22 | <DEL> | Inherited from father | Fbxl7 intron 2 | b (smaller than | 0.09933 | |
| 716 | 6 | 169506281 | 169521151 | 14870 | 0/1 | 51 | 15 | 41 | 25 | <DEL> | Inherited from mother | Intergenic | Multiple | 0.0003761 | Intergenic |
| 716 | 8 | 19786243 | 19792427 | 6184 | 0/1 | 45 | 24 | 32 | 29 | <DEL> | Inherited from mother | Intergenic | L1PA4 | 0.0002052 | Intergenic |
| 716 | 8 | 22982301 | 22988246 | 5945 | 0/1 | 59 | 15 | 42 | 20 | <DEL> | Inherited from mother | Intergenic | Multiple | Not found | Intergenic |
| 716 | 8 | 100158249 | 100158249 | 0 | 0/1 | 71 | 23 | 50 | 26 | N]8:100157812] | Inherited from father | Vps13b intron 13 | L1MD2 | Not found | Overlaps repetitive regions |
| 716 | 12 | 2952933 | 2953348 | 415 | 0/1 | 54 | 10 | 36 | 20 | <DEL> | Inherited from father | Loc100507424 intron 2 | | 0.093 | |
| 716 | 12 | 30237244 | 30243566 | 6322 | 0/1 | 76 | 18 | 49 | 25 | <DEL> | Inherited from father | Intergenic | Multiple | 0.09341 | Intergenic |
| 716 | 14 | 90991749 | 90992338 | 589 | 0/1 | 50 | 28 | 34 | 19 | <DEL> | Mother hom, proband het | Intergenic | | 0.02435 | Parent homozygous for SV |
| 716 | 15 | 97680857 | 97681219 | 362 | 0/1 | 47 | 22 | 38 | 21 | <DEL> | Inherited from father | Intergenic | MSTA | Not found | Intergenic |
| 716 | 22 | 17107192 | 17108407 | 1215 | 0/1 | 65 | 19 | 46 | 28 | <DEL> | Inherited from father | Tptep1 intron 4 | MSTB1, L1P3 | 0.05167 | Overlaps repetitive regions |
| 716 | 22 | 24876431 | 24877004 | 573 | 0/1 | 45 | 20 | 35 | 20 | <DEL> | Inherited from mother | Adora2a-as1 intron 1 | AluSz, MiR | 3.42E-05 | Overlaps repetitive regions |
| 809 | 1 | 187466727 | 187466727 | 0 | 0/1 | 77 | 15 | 52 | 11 | N]1:187464828] | Inherited from father | Intergenic | | Not found | Intergenic |
| 809 | 2 | 234631493 | 234633986 | 2493 | 0/1 | 52 | 22 | 45 | 33 | <DEL> | Inherited from mother | Ugt1a4 intron 1 | Multiple | 0.07782 | Overlaps repetitive regions |
| 809 | 4 | 141633311 | 141633603 | 292 | 0/1 | 49 | 26 | 58 | 9 | <DEL> | Inherited from father | TBC1D9 intron 1 | | Not found | |
| 809 | 4 | 141926940 | 141927241 | 301 | 0/1 | 45 | 12 | 41 | 14 | <DEL> | Inherited from father | RNF150 intron 1 | AluYa5 | 0.07885 | Overlaps repetitive regions |
| 809 | 4 | 147953942 | 147953942 | 0 | 0/1 | 90 | 6 | 74 | 17 | N]20:54914679] | Inherited from mother | Intergenic | | 0.09037 | Intergenic |
| 809 | 4 | 150789801 | 150789801 | 0 | 0/1 | 14 | 12 | 9 | 7 | ]14:96048906]N | Inherited from mother | Intergenic | | Not found | Intergenic |
| 809 | 4 | 152778524 | 152778992 | 468 | 0/1 | 61 | 23 | 49 | 25 | <DEL> | Inherited from mother | Intergenic | LTR78 | 0.07529 | Intergenic |
| 809 | 5 | 2563610 | 2564403 | 793 | 0/1 | 48 | 16 | 42 | 22 | <DEL> | Inherited from father | Intergenic | L2a, Tigger15a | Not found | Intergenic |
| 809 | 6 | 2568178 | 2571929 | 3751 | 0/1 | 58 | 20 | 45 | 34 | <DEL> | Inherited from mother | Intergenic | Multiple | 0.07471 | Intergenic |
| 809 | 6 | 162725094 | 162725913 | 819 | 0/1 | 59 | 21 | 44 | 29 | <DEL> | Inherited from father | Park2 intron 2 | MC4a, MER102 | 0.04308 | Overlaps repetitive regions |
| 809 | 8 | 114056495 | 114058772 | 2277 | 0/1 | 44 | 18 | 37 | 23 | <DEL> | Mother hom, proband het | Csmd3 intron 5 | Multiple | 0.07293 | Parent homozygous for SV |
| 809 | 8 | 115293003 | 115293414 | 411 | 0/1 | 56 | 16 | 46 | 16 | <DEL> | Inherited from mother | Intergenic | | 0.09364 | Intergenic |
| 809 | 12 | 17956553 | 17965645 | 9092 | 0/1 | 108 | 18 | 78 | 27 | <DUP> | Inherited from mother | Intergenic | Multiple | 0.01655 | Intergenic |
| 809 | 15 | 90059372 | 90061759 | 2387 | 0/1 | 45 | 9 | 34 | 32 | <DEL> | Inherited from father | LINC00928 | Multiple | Not found | Overlaps repetitive regions |
| 967 | 1 | 187466727 | 187466727 | 0 | 0/1 | 56 | 21 | 35 | 23 | N]1:187464828] | Inherited from father | Intergenic | | Not found | Intergenic |
| 967 | 1 | 218417913 | 218421598 | 3685 | 0/1 | 60 | 17 | 37 | 22 | <DEL> | Inherited from mother | Intergenic | Multiple | 0.006804 | Intergenic |
| 967 | 2 | 236742160 | 236742737 | 577 | 0/1 | 28 | 23 | 24 | 21 | <DEL> | Inherited from father | Agap1 intron 9 | L2a, MER112 | 0.002428 | Overlaps repetitive regions |
| 967 | 4 | 147953942 | 147953942 | 0 | 0/1 | 64 | 11 | 44 | 13 | N]20:54914679] | Inherited from mother | Intergenic | | 0.09037 | Intergenic |
| 967 | 5 | 5107853 | 5109888 | 2035 | 0/1 | 47 | 13 | 32 | 19 | <DEL> | Inherited from mother | Intergenic | Multiple | 0.01764 | Intergenic |
| 967 | 5 | 8554417 | 8559174 | 4757 | 0/1 | 38 | 20 | 34 | 24 | <DEL> | Inherited from father | Intergenic | Multiple | Not found | Intergenic |
| 967 | 5 | 13053738 | 13059901 | 6163 | 0/1 | 40 | 18 | 30 | 16 | <DEL> | Inherited from father | Intergenic | Multiple | 0.007249 | Intergenic |
| 967 | 5 | 17711472 | 17719560 | 8088 | 0/1 | 52 | 7 | 35 | 16 | <DEL> | Inherited from father | Intergenic | Multiple | 6.84E-05 | Intergenic |
| 967 | 6 | 162725094 | 162725913 | 819 | 0/1 | 35 | 15 | 30 | 17 | <DEL> | Inherited from father | Park2 intron 2 | MER102b | 0.04308 | Overlaps repetitive regions |
| 967 | 6 | 167543015 | 167544428 | 1413 | 0/1 | 40 | 4 | 30 | 16 | <DEL> | Inherited from father | Ccr6 intron 1 | (smaller than S | 0.01874 | |
| 967 | 6 | 168792011 | 168792011 | 0 | 0/1 | 44 | 13 | 32 | 16 | N]6:168792882] | Inherited from father | Intergenic | | 0.07632 | Intergenic |
| 967 | 8 | 10291569 | 10292936 | 1367 | 0/1 | 45 | 13 | 35 | 15 | <DEL> | Inherited from mother | Intergenic | L2b, L1MC1 | 0.005437 | Intergenic |
| 967 | 8 | 11596931 | 11597274 | 343 | 0/1 | 22 | 9 | 31 | 18 | <DEL> | Inherited from mother | **Gata4 intron 2** | | 0.01946 | |
| 967 | 8 | 21500259 | 21500761 | 502 | 0/1 | 36 | 16 | 28 | 19 | <DEL> | Inherited from father | Intergenic | AluSz6 | 0.0002393 | Intergenic |
| 967 | 8 | 23000479 | 23000793 | 314 | 0/1 | 85 | 18 | 65 | 19 | <DEL> | Inherited from father | Tnfrsf10d intron 7 | AluY | Not found | Overlaps repetitive regions |
| 967 | 8 | 24442378 | 24447067 | 4689 | 0/1 | 32 | 8 | 20 | 15 | <DEL> | Inherited from father | Intergenic | Multiple | 0.01778 | Intergenic |
| 967 | 8 | 26856384 | 26858022 | 1638 | 0/1 | 87 | 24 | 55 | 17 | <DUP> | Inherited from mother | Intergenic | L1ME1, LTR8A | 0.05591 | Intergenic |
| 967 | 8 | 103053960 | 103056716 | 2756 | 0/1 | 54 | 19 | 39 | 19 | <DEL> | Inherited from mother | Ncald intron 1 | 78 (smaller than | 0.01057 | |
| 967 | 8 | 104906386 | 104912697 | 6311 | 0/1 | 51 | 17 | 29 | 19 | <DEL> | Inherited from father | Rims2 intron 4 | Multiple | 0.01392 | Overlaps repetitive regions |
| 967 | 8 | 106219051 | 106223429 | 4378 | 0/1 | 31 | 14 | 27 | 13 | <DEL> | Inherited from father | Intergenic | Multiple | 6.84E-05 | Intergenic |
| 967 | 8 | 106247733 | 106248024 | 291 | 0/1 | 48 | 16 | 37 | 22 | <DEL> | Inherited from mother | Intergenic | L1MDa | 0.02132 | Intergenic |
| 967 | 8 | 106796533 | 106797305 | 772 | 0/1 | 31 | 26 | 24 | 37 | <DEL> | Father hom, proband het | Zfpm2 intron 5 | MiR, L2 | Not found | Parent homozygous for SV |
| 967 | 8 | 114040467 | 114046766 | 6299 | 0/1 | 37 | 22 | 27 | 20 | <DEL> | Inherited from mother | Csmd3 intron 5 | Multiple | 0.02011 | Overlaps repetitive regions |
| 967 | 8 | 114912823 | 114913530 | 707 | 0/1 | 52 | 7 | 31 | 12 | <DEL> | Inherited from mother | Intergenic | AluY | 0.06319 | Intergenic |
| 967 | 8 | 115293003 | 115293414 | 411 | 0/1 | 38 | 10 | 27 | 16 | <DEL> | Inherited from father | Intergenic | | 0.09364 | Intergenic |
| 967 | 9 | 8640762 | 8641661 | 899 | 0/1 | 69 | 10 | 36 | 21 | <DEL> | Inherited from father | Ptprd intron 12 | | 0.08948 | Intergenic |
| 967 | 11 | 128311492 | 128316095 | 4603 | 0/1 | 38 | 12 | 25 | 12 | <DEL> | Inherited from mother | Intergenic | Multiple | 0.0611 | Intergenic |
| 967 | 12 | 11258537 | 11258838 | 301 | 0/1 | 41 | 17 | 36 | 17 | <DEL> | Father hom, proband het | Intergenic | AluYk12 | Not found | Parent homozygous for SV |
| 967 | 12 | 11341901 | 11342212 | 311 | 0/1 | 33 | 13 | 30 | 15 | <DEL> | Father hom, proband het | Intergenic | L1M5, AluJb | Not found | Parent homozygous for SV |
| 967 | 12 | 12252827 | 12254646 | 1819 | 0/1 | 35 | 20 | 30 | 19 | <DEL> | Inherited from mother | Intergenic | Multiple | 0.05642 | Intergenic |
| 967 | 12 | 21861577 | 21861929 | 352 | 0/1 | 47 | 14 | 28 | 18 | <DEL> | Inherited from mother | Intergenic | MiR | 0.05799 | Intergenic |
| 967 | 12 | 25285421 | 25285878 | 457 | 0/1 | 40 | 26 | 32 | 26 | <DEL> | Inherited from father | Casc1 intron 9 | L1MC4a | 0.05403 | Overlaps repetitive regions |
| 967 | 14 | 106723481 | 106723835 | 354 | 0/1 | 90 | 17 | 59 | 17 | <DEL> | Inherited from mother | Intergenic | | 0.03649 | Intergenic |
| 967 | 15 | 93839703 | 93839703 | 0 | 0/1 | 75 | 12 | 50 | 14 | [2:228189984[N | Inherited from father | Intergenic | L1ME4a | Not found | Intergenic |
| 967 | 15 | 98532622 | 98536469 | 3847 | 0/1 | 42 | 7 | 35 | 21 | <DEL> | Inherited from father | Intergenic | Multiple | 0.0516 | Intergenic |
| 967 | 22 | 17107192 | 17108407 | 1215 | 0/1 | 50 | 16 | 35 | 16 | <DEL> | Inherited from father | Tptep1 intron 4 | Multiple | 0.05167 | Overlaps repetitive regions |
| 967 | 22 | 22906059 | 22909454 | 3395 | 0/1 | 36 | 12 | 30 | 9 | <DEL> | Father hom, proband het | Loc648691 exon 3 | AluSx, MLT1E1A | 0.03085 | Parent homozygous for SV |

### TABLE S5

| CDH Proband | Chromosome | Start | End | Length of Variant | Genotype: 0 = reference, 1 = alternate SV | Reference split reads | Alternative split reads | Reference discordant pairs | Alternative discordant pairs | Length and translocated location | Mutation type | Gene location | Repeats overlapping SV | Allele frequency in Abel *et al.* 2018 | Filtering reason |
|---|---|---|---|---|---|---|---|---|---|---|---|---|---|---|---|
| 411 | 13 | 86977496 | 86977496 | 0 | 0/1 | 53 | 21 | 0 | 0 | N[6:115243059[ | De Novo (in all three proband samples) | Intergenic | L1PA12 | 0.2683 | Common in Abel *et al.* 2018, intergenic |
| 411 | 17 | 80856815 | 80856996 | 181 | 0/1 | 126 | 14 | 48 | 6 | <DUP> | De Novo (in all three proband samples) | TBCD intron 17 | None | 0.1517 | Common in Abel et al. 2018 |
| 716 | 10 | 4454891 | 4454891 | 0 | 0/1 | 28 | 10 | 0 | 0 | N]X:94367322] | De Novo (in all three proband samples) | Intergenic | (TA)n | 0.01238 | Intergenic |
| 809 | 18 | 64889326 | 64891140 | 1814 | 0/1 | 62 | 23 | 52 | 30 | <DEL> | De Novo (in all three proband samples) | Intergenic | AluSx | Not found | Intergenic |
